# Loss of characteristic species across German federal states detected by repeated mapping of protected habitats

**DOI:** 10.1101/2025.02.27.640325

**Authors:** Lina Lüttgert, Samuel Heisterkamp, Florian Jansen, Rico Kaufmann, Simon Kellner, Reinhard Arnold Klenke, Silke Lütt, Gunnar Seidler, Axel Wedler, Ronja Wörmann, Helge Bruelheide

## Abstract

Identifying the winners and losers of biodiversity change within different habitat types requires systematic monitoring. While such data are still lacking in Germany, species trends could be derived from previously untapped sources. Here, we derive temporal trends in plant species from data of repeated habitat mapping programs of three German states from 1977-2021, both across all habitat types per state and within habitat types. Consistently negative trends were found across all states for species preferring heaths and semi-natural grasslands, moist to wet grasslands, and coastal and marine habitats, including many endangered species. Consistently positive trends were found for species preferring scrubs, copses and field hedges, and for non-native species. Trends within habitat types showed negative trends for species characteristic of those habitat types. While trends varied among states, the overall patterns were very similar. This points to ongoing habitat degradation and common drivers of biodiversity change in Germany.

## Introduction

In response to the ongoing biodiversity loss and habitat degradation in Europe, with the EU Biodiversity Strategy and Nature Restoration Law the European Union set targets to protect 30% of its terrestrial and marine area and to restore at least 30% of degraded habitats by 2030 (European Commission, 2022; European Commission & Directorate-General for Environment, 2021). Further targets include halting or even reversing negative trends in both protected habitat types and plant species. To efficiently direct these nature conservation efforts, identifying degraded habitats as well as the winners and losers of habitat degradation is essential (Kühl et al., 2020).

Past decades’ environmental change did not threaten species randomly, with especially habitat specialists showing declines (Britton et al., 2009; Diekmann et al., 2019; Heinrichs & Schmidt, 2017). For plants, these are often species typical of nutrient-poor habitats due to habitat loss and nutrient enrichment (Klinkovská et al., 2024; Sperle & Bruelheide, 2021). Losers are also often rare and endangered species (Kempel et al., 2020; Metzing et al., 2018). In contrast, the set of plant species that have benefited from environmental change includes common and widespread generalist, often nutrient-demanding species (Glaser et al., 2024; Hallman et al., 2022; Heinrichs & Schmidt, 2017; Kindlund & Tyler, 2023; Staude et al., 2023). Trends are particularly positive for woody species (Buitenwerf et al., 2018; Hallman et al., 2022; Wieczorkowski & Lehmann, 2022) and non-native species (Eichenberg et al., 2021; Hallman et al., 2022; Kindlund & Tyler, 2023).

How many and what type of species increase or decrease depends on the spatial and temporal scale the observations are made, the region and the habitat types studied (Eichenberg et al., 2021; Gonzalez et al., 2023; Prévosto et al., 2011; Vellend et al., 2017). While studies based on a fine spatial grain, such as permanent vegetation plots on a few square meters, are sensitive to detect changes, they are mostly not spatially representative and exclude sites that have experienced habitat type transition and destruction (Gonzalez et al., 2016; Jandt et al., 2022). In contrast, studies on a coarse spatial grain, for example based on grid-cells of several square kilometres, can only detect strong trends and do not allow to distinguish between habitat types (e.g. Eichenberg et al., 2021; Rich & Woodruff, 1996). It is the intermediate scale, i.e. of habitats within a landscape, that would provide sufficient sensitivity and representativeness. However, such studies at the regional scale are notoriously scarce (Chase et al., 2019; Naaf & Wulf, 2010). This is particularly unfortunate because at this scale most conservation measures are implemented (Erasmus et al., 1999; Ferrier, 2002; Naaf & Wulf, 2010). Similar to the spatial scale, the temporal scale will influence the trends observed, with short-term studies expected to result in different and/or weaker trends than long-term studies due to short-term fluctuations and time lags (Magurran et al., 2010; Skálová et al., 2022). Still, long-term studies are rare. Furthermore, we lack studies that allow comparing trends between different habitat types. Thus, in an ideal world, systematic plant monitoring would cover all habitat types, be geographically representative, and extend into the last century (Buckland & Johnston, 2017; Gonzalez et al., 2023). While such systematic data are not available in any country, some data sources may come close, although they are based on programs that were never intended to provide species trends.

Such data might be extracted from habitat mapping programs, which have been established decades ago in many countries with the goal of nature conservation and landscape planning (Bunce et al., 2012; European Environment Agency & Museum national d’Histoire naturelle, 2014; Lengyel et al., 2008). In Germany, mapping campaigns have been carried out by all federal states over the past decades, in some of them also repeatedly (Kaiser et al., 2013). They include wall-to-wall mapping of all protected habitat types through field surveys, including records of plant species occurrences in most habitat sites. The programs thus provide species occurrence information at the habitat scale across large regions and habitat types. More recently, these data have been used to derive species trends, but only to a limited extend for individual federal states (Bruelheide et al., 2020; Jansen et al., 2020; Lüttgert et al., 2022; Lüttgert et al., 2024). A main obstacle preventing data integration across federal states has been the federal structure of the nature conservation administration in Germany, which resulted in differences in mapping keys, habitat definitions and database structures. We also expected that conditions across states might be idiosyncratic due to differences in regional history of land use, habitat types and environmental conditions (Hallman et al., 2022; Prévosto et al., 2011; Timmermann et al., 2015).

Here, we employ data from the habitat mapping programs of three federal states (hereafter states) in Germany to derive and compare their plant species trends. We chose the states Schleswig-Holstein, Hamburg, and Baden-Württemberg because they are representative of the German landscape but also because they had repeated mapping data available for at least two time periods within 1977-2021. We calculated species trends both across all habitat types of each state and within different habitat types, and tested whether specific species groups tended to increase or decrease over time. We then compared the consistency of trends among states based on both individual species trends and group trends.

While the three states differ in location (northern vs. southern, coastal vs. more continental), geography (lowland vs. upland) and urbanization level, they have all experienced intensification of agricultural land use, increases of build-up area, and abandonment of traditional land uses. Thus, we expected that these drivers would result in common patterns of species change. In particular, we hypothesized to encounter

1) a common set of losers across the states, including species of nutrient-poor habitats (bogs, semi-natural grasslands), and endangered species;
2) a common set of winners across the states, including shrubs, ruderals, and neophytes (non-native species introduced after 1492);
3) declines in habitat-characteristic species within their preferred habitat type across all states.

## Methods

### Habitat mapping

The three German federal states Schleswig-Holstein (SH), Hamburg (HH), and Baden-Württemberg (BW) together cover an area of 52 307 km^2^, which is 14.6% of Germany (Statistisches Bundesamt, 2024; SH: 15 804 km^2^, HH: 755 km^2^, and BW: 35 748 km^2^; Figure S1.1). All three states have been mapping their protected habitat types for decades, starting in the 1970s. While in BW only protected habitat types have been mapped, in SH also other “potentially valuable” sites were included, while in HH also non-protected habitat types have been mapped since the mid-1990s. Most habitat sites have been remapped at least twice by 2021 (except in some districts for the open land mapping in BW). Habitat mapping includes the recording of both habitat types and plant species at each site. However, complete species lists were not mandatory. Thus, species lists are mostly incomplete, with the focus on species that are endangered, characteristic for a particular habitat type, or dominant at a particular site.

### Spatial data processing

All states have digitized at least one version of old and all new mapping data, with each habitat site represented by a polygon to which metadata and species lists were assigned. Hereafter, we refer to a mapped unit as a polygon, even if a mapped unit sometimes consists of several different polygons in proximity if they all belong to the same habitat type. The number and size of polygons differed between states and mapping periods (Table S1.1). We processed the spatial data separately by federal state. For each state, we intersected polygons from the first digitally available mapping campaign with polygons from the most recent mapping campaign in ArcGIS 10.5 (ESRI, 2016). Only intersecting polygons were used for trend analyses. Thus, polygons that were either not remapped or not previously mapped were excluded. Further data cleaning was slightly adapted for each state, according to the underlying mapping schemes and data (see Supplementary Methods for SH, Lüttgert et al. (2022) for HH, and Lüttgert et al. (2024) for BW for explicit cleaning steps). Generally, we excluded small intersections considered as mapping and digitization inaccuracies (less than 5% of each or either polygon, depending on state), polygons which had not been remapped to a proportion of a set threshold (50% for SH, 95% for HH, and 75% for BW), and which were not accompanied by a species list. We divided data into two time intervals each, which differed slightly between the states depending on the available data: *t*_1_: 1977-2005 and *t*_2_: 2007-2021 for SH, *t*_1_: 1979-1994 and *t*_2_: 1995-2017 for HH, and *t*_1_: 1989-2005 and *t*_2_: 2006-2021 for BW. Time spans between resurveys of polygons ranged from 6 to 42 years, with a median of 32, 22, and 19 years for SH, HH, and BW, respectively (Figure S1.2).

### Plant occurrence data

Taxonomy was harmonized using GermanSL 1.5. (Jansen & Dengler, 2008, 2010), aggregating species to the section level and merging some taxa further onto the genus level. We excluded mosses, lichen, algae, most hybrids and cultivated forms, as well as some species known to have been falsely identified. 2212 species remained after harmonization, with 1301, 1287, and 1977 species recorded in SH, HH, and BW, respectively. 998 species were recorded in all states. Since the ratio of species to polygons was relatively high in HH, we drew species accumulation curves for all states, expecting slower accumulation of species for HH. However, while the curve was slightly steeper for HH, the asymptote was approached in all states (Figure S1.3).

### Habitat type data

We used information on habitat types per polygon to assign preferred habitat types to species and to calculate species trends within habitat types. Habitat type categories differed between states and mapping periods. Thus, we assigned all habitat types to 14 broadly defined habitat type groups in accordance with all habitat mapping keys, including 646, 475, and 248 detailed habitat types from SH, HH, and BW, respectively (Tables S1.2 & S1.3). Some groups were only recorded in some states. Since a polygon could have multiple habitat types assigned to it, we only used the habitat type with the highest cover (at least 51%) of each polygon.

### Preferred habitat types, Red List and non-native status

We assigned a preferred habitat type to each species by applying the Φ coefficient, i.e. the fidelity of each species x habitat type combination (Chytrý et al., 2002; Equation 1).

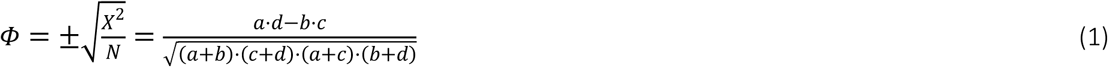

The fidelity Φ of a species to a specific habitat type is calculated using the Χ^2^ statistic for a 2 x 2 contingency table and is dependent on the total number of observations *N*, with *a* the number of occurrences of the species within the habitat type, *b* the number of occurrences of the species outside the habitat type, *c* the number of absences of the species within the habitat type, and *d* the number of absences of the species in all other habitat types. Φ ranges from −1 to 1 for species perfectly avoiding a habitat type or species perfectly bound to a habitat type, respectively. We used all polygons from all states together (including polygons that did not intersect) to calculate across-state fidelities for all species x habitat type combinations. Fidelities are therefore biased towards the larger states as they had more polygons available. We considered the habitat type showing the highest fidelity to each species as its preferred habitat type. Species that prefer a habitat type can also be seen as characteristic for that habitat type. However, we did not assign a type to species whose highest Φ value was below the median of all species’ highest Φ value (0.0073). This resulted in the removal of species with only few observations from the list of habitat-specific species.

We assigned Red List (endangerment) and non-native status in Germany to all species, based on the Red List of Germany (Metzing et al., 2018) and BiolFlor (Klotz et al., 2002), respectively. Non-native species were separated into archaeophytes, i.e. non-natives introduced before 1492, and neophytes, i.e. non-natives introduced after 1492. We manually assigned a few species’ non-native status based on information from Floraweb (www.floraweb.de; Buttler et al., 2018; Wisskirchen & Haeupler, 1998). We did not assign a status to species aggregates for which the lower taxonomic levels had different status.

### Trend calculation and metrics

Polygons from one time interval were often overlapping with several polygons from the other time interval, caused by habitat change or changes in the detail in which habitat types were mapped. Thus, we compared each *t*_1_-polygon’s species list with the merged species lists of all its intersecting polygons from *t*_2_ (hereinafter *t*_1_ → *t*_2_), and each *t*_2_-polygon’s species list with the merged species lists of all its intersecting polygons from *t*_1_ (hereinafter *t*_2_ → *t*_1_). We derived species trends using both *t*_1_ → *t*_2_ and *t*_2_ → *t*_1_ separately. Because we often compare one species list from one time period with several species lists from the other time period, changes derived from *t*_1_ → *t*_2_ are biased towards species gains and changes derived from *t*_2_ → *t*_1_ are biased towards species losses. Thus, to make sure to only report robust trends that can be compared between states, we here only report trends that were significant and consistent for both approaches *t*_1_ → *t*_2_ and *t*_2_ → *t*_1_. For those cases, we report the mean trends derived from both approaches and common statistical values.

We calculated temporal species trends separately for each state, using two change metrics each: relative change in frequency and probability of occurrence. For frequency trends, we calculated the change in presence/absence for each species in all its previously and/or recently occupied polygons from *t*_1_ towards *t*_2_ (−1, 0, or 1). The mean of those changes per species represented the relative change across each state. Trends ranged from −1 to 1, with a value of −1 indicating that a species was lost in all previously occupied polygons and 1 that a species was new to all its polygons.

To account for incomplete species observations, we additionally estimated species change in probability of occurrence, using Beals’ index of sociological favourability (Beals, 1984). We estimated the probability of each species to occur in each polygon, based on a species’ co-occurrence information with all species that were recorded in that polygon (Equation 2).

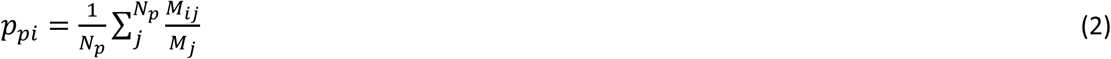

The probability *p_pi_* for species *i* to occur in polygon *p* is calculated from its joint occurrences *M_ij_* with all species *j* of the number of species in that polygon *N_p_*, divided by the number of polygons *M_j_* in which species *j* is present. *M_ij_* values were derived from a co-occurrence matrix of all species across all polygons of a state (including polygons that did not intersect). We then calculated the change in probability of occurrence (hereinafter Beals) from *t*_1_ to *t*_2_ for each polygon *p* x species *i* combination and calculated each species’ mean change across each state. Because Beals values are generally lower for co-occurrence matrices that are based on a higher number of polygons, the large state BW had lower Beals values, and hence, also Beals change values were lower than in the other states (Figure S1.4). To make the values comparable between the states, we standardized the Beals trends per species and state by dividing them by the standard deviation of species’ Beals trends of each state.

### Trends within habitat types

In addition to deriving trends across a state, we also derived trends within different habitat types. We used the same habitat types that we used for assigning species to their preferred habitat types. We grouped the data by using the habitat type of the first mapping of a habitat site, regardless of whether a polygon’s habitat type changed over time. For each recorded habitat type and species combination we derived trends in both frequency and in Beals as described above for the trends across the states. Again, we standardized the mean Beals change values per species, habitat type and state by dividing them by the standard deviation of the mean trends of each state.

### Statistical analyses

To test for significance of each species’ frequency trend, we used two-tailed binomial tests (i.e. sign tests). We first tested significance for the approaches *t*_1_ → *t*_2_ and *t*_2_ → *t*_1_ separately while accounting for duplicated usage of polygons within *t*_1_ → *t*_2_ and within *t*_2_ → *t*_1_ by reducing the degrees of freedom accordingly. For species with significant trends for both *t*_1_ → *t*_2_ and *t*_2_ → *t*_1_, we calculated their mean trends as well as common p-values from both approaches, also with two-tailed binomial tests. For this, we used the mean numbers of occurrences from both approaches, i.e. the mean number of increases, mean number of decreases, and mean number of cases without a change.

To test for significance of each species’ Beals trend, we used two-tailed t-tests, again first testing significance for the approaches *t*_1_ → *t*_2_ and *t*_2_ → *t*_1_ separately. We corrected p-values by using the number of polygons in which a species actually occurred instead of the number of all polygons. We corrected for duplicated usage of polygons within *t*_1_ → *t*_2_ and within *t*_2_ → *t*_1_ and additionally applied Holm adjustment of significance levels for testing of multiple species. For species with significant trends for both *t*_1_ → *t*_2_ and *t*_2_ → *t*_1_, we calculated their mean trend as well as common p-values and 95% confidence intervals from both approaches. For this, we used the mean trend, mean number of occurrences, and mean variance from both approaches to derive common standard errors and t-values.

For each state, we used Spearman rank correlations to compare frequency and Beals trends using all species with a significant trend for both metrics. We additionally compared single species trends between two states each by using Spearman rank correlations for each metric.

Next, for each state and change metric, we tested if specific species groups rather increased or decreased. We grouped species based on their preferred habitat type, Red List, and non-native status. We only used species with a significant trend in the respective change metric and tested for significance using two-tailed Wilcoxon signed rank tests. We also produced trend heatmaps for species grouped by state and preferred habitat type, including dendrograms for states and for preferred habitat types.

We additionally calculated each species mean trend across all states by taking the average from the three state trends. Again, we did so for frequency and Beals trends. Based on those mean trends, we tested again if specific species groups (preferred habitat type, Red List, non-native status) rather increased or decreased using two-tailed Wilcoxon signed rank tests.

For frequency and Beals trends within habitat types, we used the same statistical approach as we did for the trends across the states. For each state and change metric, we then tested whether all species within a given habitat type tended to show positive or negative trends. We only used species x habitat type combinations with a significant trend in the respective change metric and tested for significance using two-tailed Wilcoxon signed rank tests.

The significance for all statistical tests was determined at p < 0.05.

If not stated otherwise, we used R 4.3.2 (R Core Team, 2023) for all analyses. We used the packages data.table, dplyr, ggdist, ggnewscale, ggplot2, ggpubr, gplots, reshape2, sf, sjPlot, stringr, and vegdata.

## Results

### Trends across each state

We found significant Beals trends for 389, 319, and 809 species in SH, HH, and BW, respectively (Table S2). Overall, 105 species displayed a significant Beals trend in all states (Table S1.4). Significant trends in frequency were encountered for 136, 122, and 404 species in SH, HH, and BW, respectively (Table S3). Only nine species showed a significant frequency trend in all states.

A total of 96, 93, and 307 species showed a significant trend in both frequency and Beals in SH, HH, and BW, respectively (Figures S1.5-S1.7). Frequency and Beals trends in each state were only moderately correlated (Spearman rank correlation: r_s_ = 0.53 for SH, r_s_ = 0.53 for HH, and r_s_ = 0.26 for BW, all p < 0.001).

Trends of species that were significant in two states were positively correlated between states. These Spearman rank correlations between states were generally higher for Beals trends, with the highest for trends between SH and HH (r_s_ = 0.82 with n = 137 for SH and HH, r_s_ = 0.68 with n = 251 for SH and BW, r_s_ = 0.71 with n = 237 for BW and HH, all p < 0.001; Figure S1.8 a-c). For frequency, Spearman rank correlations were lower, especially for trends between SH and BW (r_s_ = 0.81 with p <0.001 and n = 23 for SH and HH, r_s_ = 0.39 with p = 0.011 and n = 42 for SH and BW, r_s_ = 0.58 with p < 0.001 and n = 58 for BW and HH; Figure S1.8 d-f).

### Preferential habitat types

Species preferring scrubs, copses and field hedges mainly increased in their probability of occurrence in all states (Figure 1, Table S1.5). By contrast, species preferring moist to wet grasslands, heaths, inland dunes and semi-natural grasslands, as well as coastal and marine habitats mostly decreased in their probability of occurrence in all states. For species preferring other habitat types, Beals trends were not significant in all states (Figure 1, Table S1.5). Species of bogs, transition mires, marshes and fens showed negative trends in both SH and BW. Species of mesic grasslands as well as species of moist to wet forests showed negative trends in both HH and BW. Comparing the direction and magnitudes of significant Beals trends between states, based on their preferred habitat type, revealed no case of opposing trends (Figure 2).

**Figure 1.**
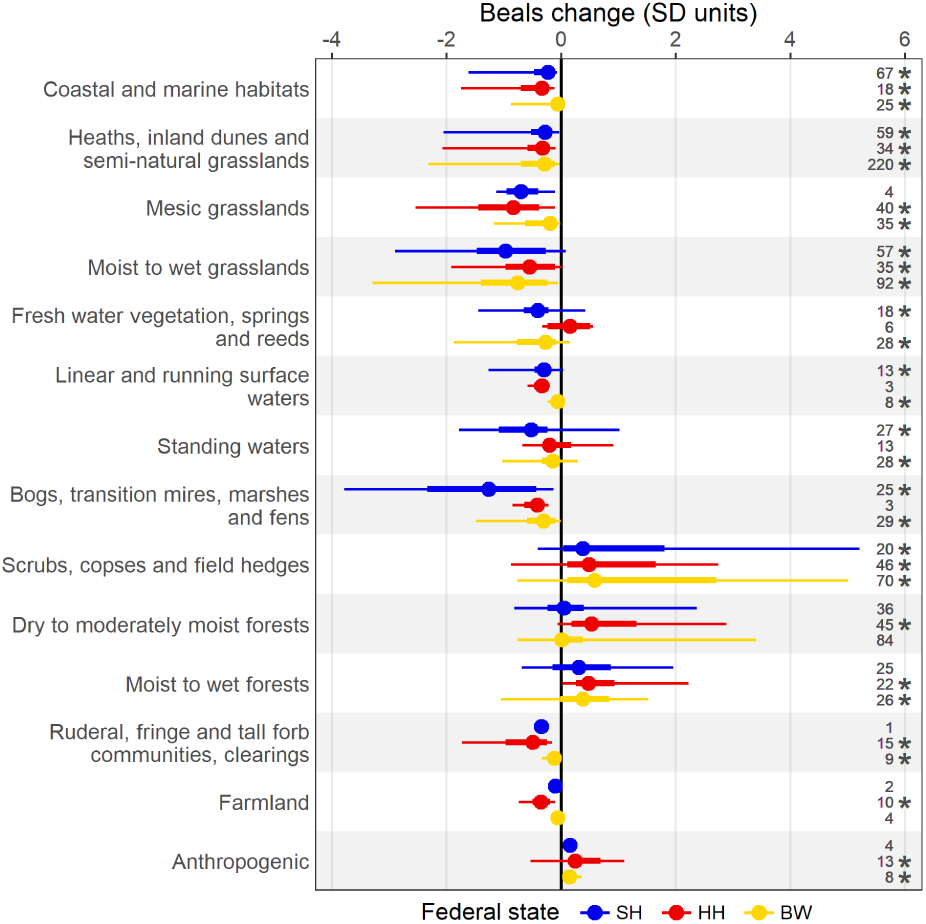
Beals trends of species grouped by their preferred habitat type separately by state. Points, thick and thin whiskers show the median, 50% and 95% data range. Only species are included per group that showed a significant trend and that had a preferred habitat type assigned. *n* is given for number of species with a significant trend that are included in each combination of state and preferred habitat type. Asterisks indicate whether a group’s trend deviated from zero change according to Wilcoxon signed rank tests. Habitat types correspond to the following EUNIS habitat types (European Nature Information System; European Environment Agency; Moss (2008); 2021 version if not stated otherwise): Coastal and marine habitats (N1, N2, N3), Heaths, inland dunes and semi-natural grasslands (S4, R1), Mesic grasslands (R2), Moist to wet grasslands (R3), Fresh water vegetation, springs and reeds (C3, D5; based on EUNIS 2012 version), Linear and running surface waters (C2, C3; based on EUNIS 2012 version), Standing waters (C1, C3; based on EUNIS 2012 version), Bogs, transition mires, marshes and fens (D1, D2, D4, D5; based on EUNIS 2012 version), Scrubs, copses and field hedges (S3, T4, V4), Dry to moderately moist forests (T1, T3, V6), Moist to wet forests (T1, T3), Ruderal, fringe and tall forb communities, clearings (V3, T4, R5), Farmland (V1, V5, V6), Anthropogenic (V2, V3).

**Figure 2.**
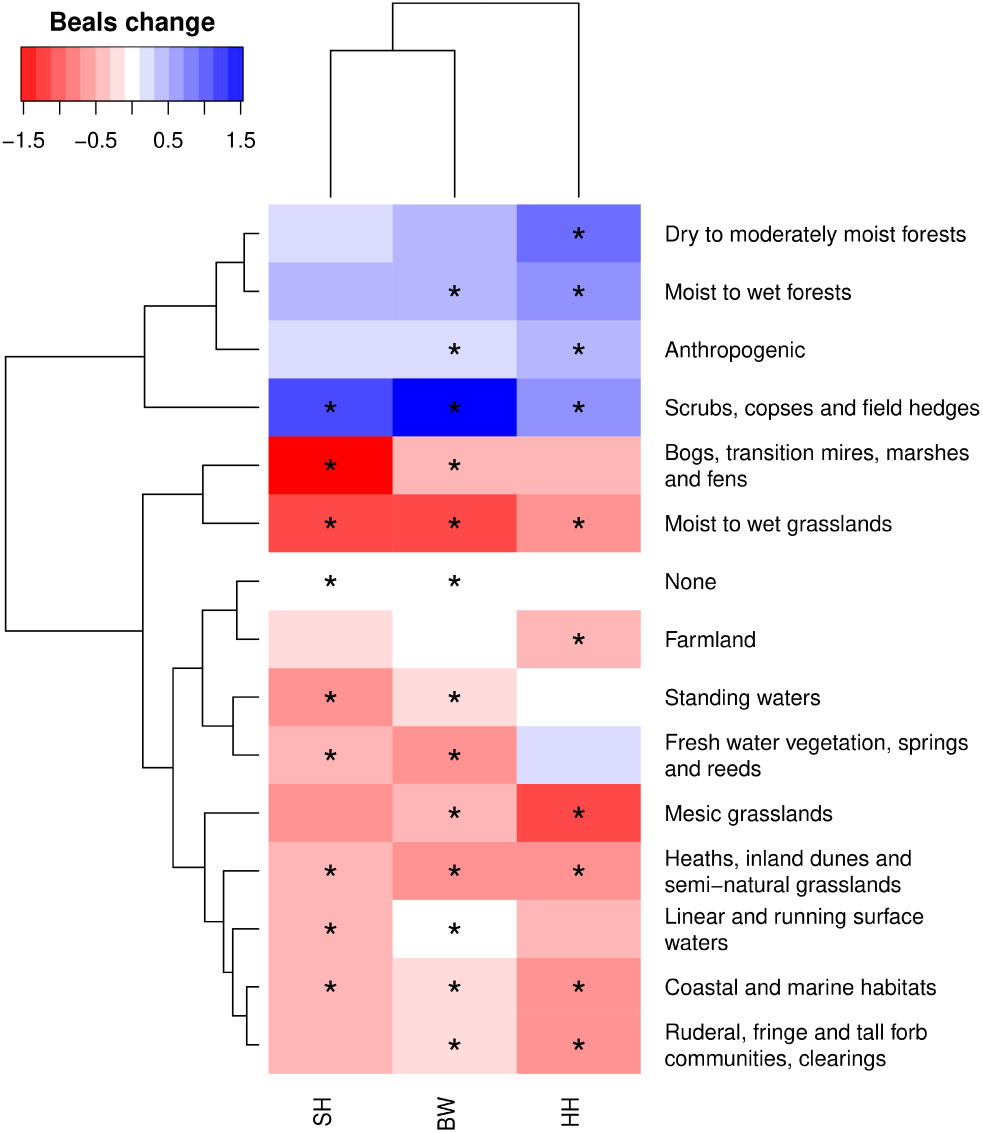
Mean Beals trends of species grouped by state and by their preferred habitat type. Only species were included in a group that showed a significant trend in the respective state (but not necessarily in all states). Asterisks indicate whether a group’s trend deviated from zero change according to Wilcoxon signed rank tests.

Similarly, out of the 105 species with a significant Beals trend in all states, only very few showed opposing trends between states (Figure S1.9). Mean species trends across states revealed relatively many winner species preferring scrubs, copses and field hedges, moist to wet forests and dry to moderately moist forests (Figure 3, Table S1.5). In contrast, loser species often preferred moist to wet grasslands, heaths, inland dunes and semi-natural grasslands, and coastal and marine habitats. In contrast to the Beals trends, no clear species frequency trends per preferred habitat type were encountered across all states (Figure S1.10).

**Figure 3.**
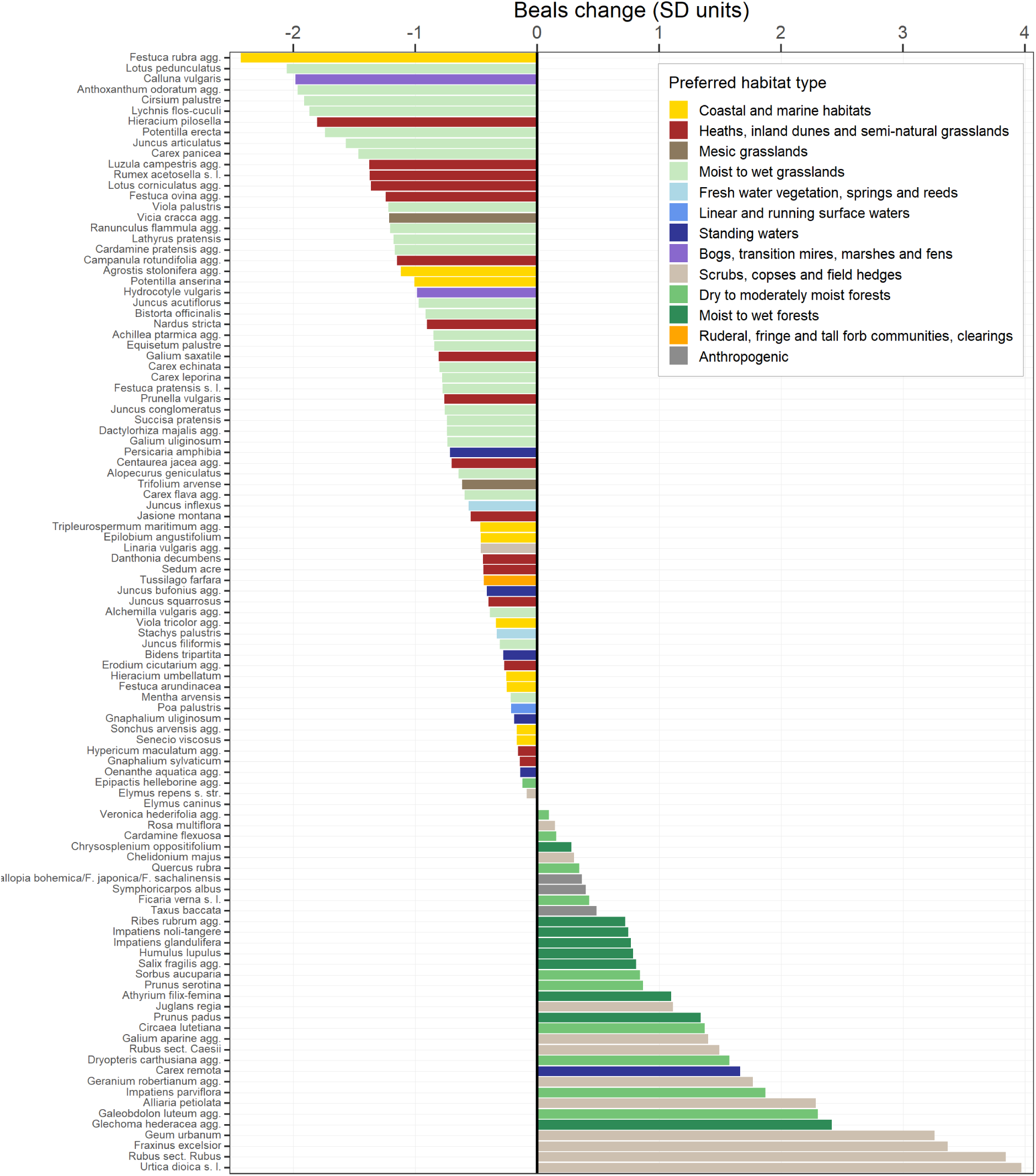
Mean Beals trend of species across all states, derived from the three trends of each state. Only species are included that showed a significant trend in all states (105). Colours indicate the species’ preferred habitat type.

### Non-native and Red List status

Neophytes increased in their occurrence probability in all states, however this was only significant for SH and BW (Figure 4a, Table S1.6). Native species on average decreased in their occurrence probability in all states. Archaeophytes showed negative trends in both BW and HH, while in SH no significant overall trend was found for this group. Trend patterns concerning those three status groups were generally similar for frequency and Beals (Figure 4a, Figure S1.11a).

**Figure 4.**
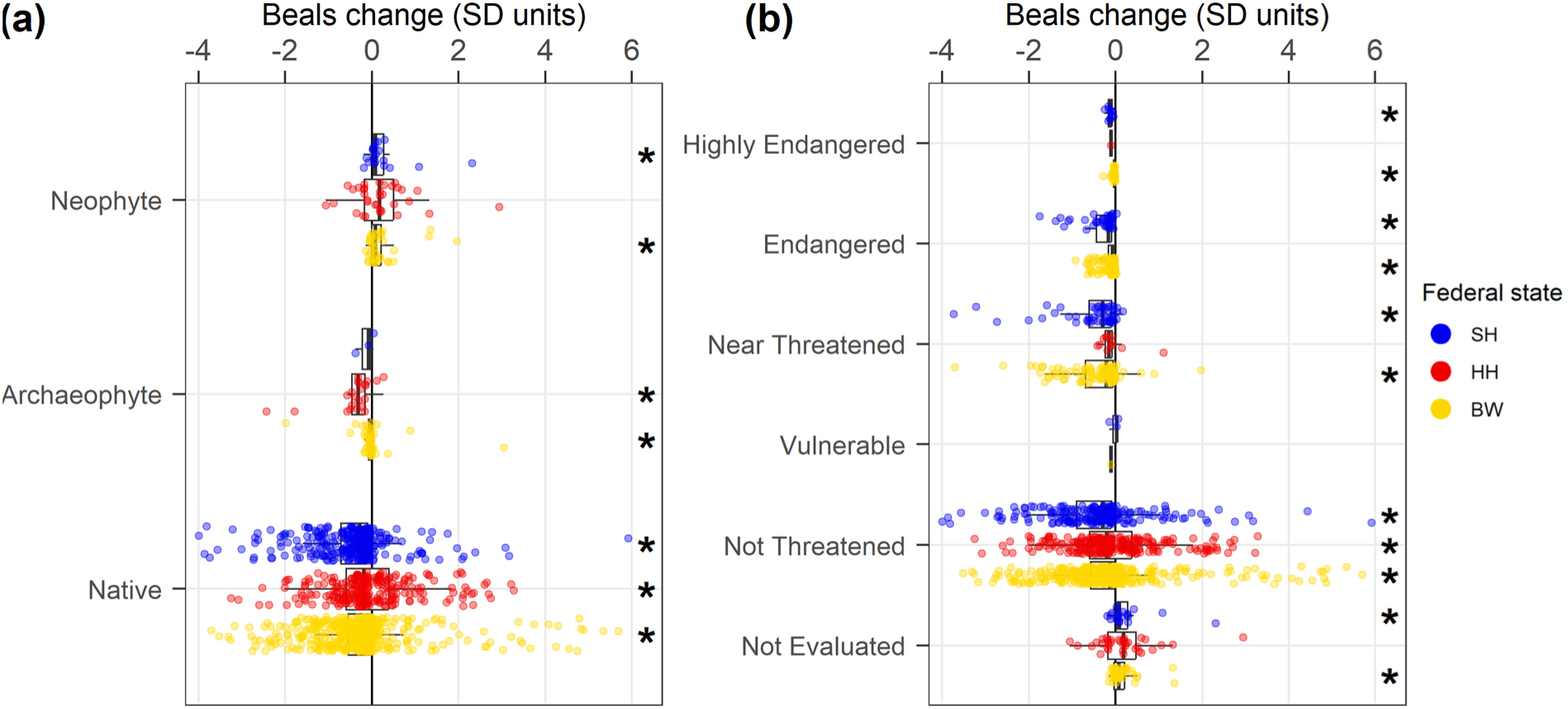
Beals trends grouped by states and **(a)** non-native and **(b)** Red List status. **(a)** Archaeophytes = non-natives introduced before 1492, neophytes = non-natives introduced after 1492. **(b)** There were no species with a significant trend of the Red List categories “extinct or lost” or “threatened with extinction”. Asterisks indicate whether a group’s trend deviated from zero change according to Wilcoxon signed rank tests. Status according to German wide lists (Buttler et al., 2018; Klotz et al., 2002; Metzing et al., 2018; Wisskirchen & Haeupler, 1998).

Regarding the Red List status, highly endangered, endangered, and near threatened species decreased in their occurrence probability in SH and BW (Figure 4b, Table S1.7). In addition, non-threatened species decreased in their occurrence probability in all states. There were only very few significant group trends regarding endangered species’ frequency (Figure S1.11b).

### Trends within habitat types

We found significant Beals trends for 1380, 67, and 1458 species x habitat type combinations for SH, HH, and BW, respectively. Frequency trends were significant for 600, 131, and 776 species x habitat type combinations for SH, HH, and BW, respectively. Only five combinations showed significant trends in all states, both concerning Beals and frequency trends.

Species trends within habitats types showed overall more positive trends for frequency compared with Beals (Figure 5, Figure S1.12). We found a prevalence of species to significantly decrease in Beals within two, zero, and five habitat types in SH, HH, and BW respectively, while a prevalence to significantly increase was only encountered within one habitat type in HH (dry to moderately moist forests). Concerning frequency, we found a prevalence of species to significantly decline only within one habitat type in BW, while a prevalence of species to significantly increase within eight, six, and four habitat types in SH, HH, and BW, respectively.

**Figure 5.**
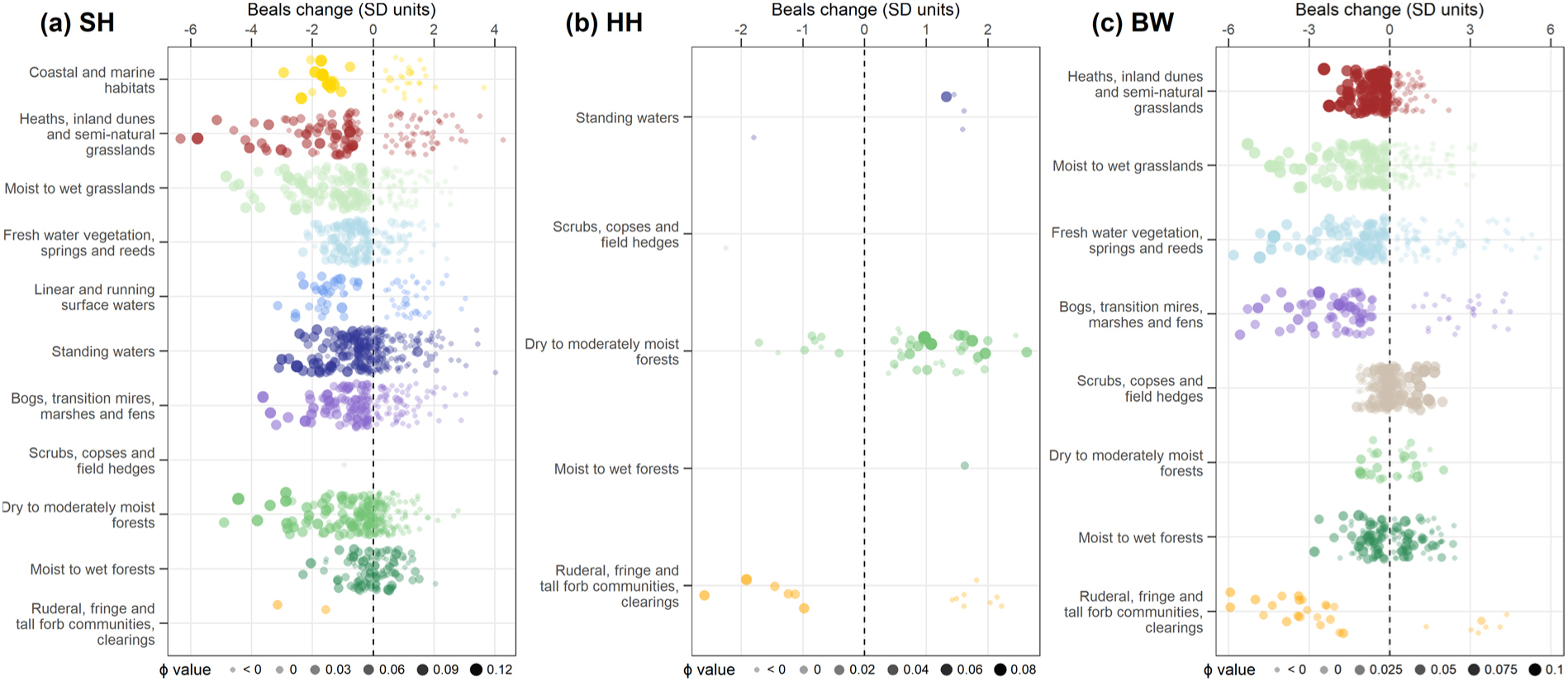
Beals trends within habitat types for **(a)** SH, **(b)** HH, and **(c)** BW. Species’ preferences to each habitat type (Φ value) are indicated by size and opacity. All Φ values below 0 are grouped into the category < 0. Only species are included that showed a significant trend. Note that not all habitat types were mapped in all states.

Especially species that preferred the respective habitat type showed decreases in their probability of occurrence in SH and BW (Figure 5). This pattern was not found for most trends in frequency or in HH (Figure S1.12, Figure 5). Species of scrubs, copses and field hedges as well as forests increased within several habitat types, especially in SH and BW (Figures S1.13 & S1.14). Within dry to moderately moist forests, species preferring these forest types showed mostly negative trends in SH, mixed trends in BW, and positive trends in HH (Figure 5).

## Discussion

We found consistent trends in probability of occurrence for most species groups across all three federal states. Similarly, the trends for the same species were largely consistent across states. Both approaches identified a set of loser species typically found in heaths, inland dunes and semi-natural grassland, wet to moist grassland, or coastal and marine habitats. In contrast, winners were species characteristic of scrubs, copses and field hedges.

Declines in species of heaths, inland dunes and semi-natural grasslands confirm our expectation that in particular species that favour nutrient-poor conditions are suffering from losses. These species have shown declines in many habitats across Europe and are typically threatened by management abandonment and increases in nutrients (Diekmann et al., 2014; Fagúndez, 2013; Jandt et al., 2011; Klinkovská et al., 2024; Tyler et al., 2020). Similarly, species characteristic of bogs, transition mires, marshes and fens, which where almost exclusively species of nutrient-poor habitat types, showed overall negative trends in BW and SH. However, this pattern was not significant in HH. This might be due to the limited number of bogs and mires remaining at the beginning of habitat mapping in this state, resulting in a rather small number of typical species present. Following habitat destruction, climate change, and eutrophication, declines of typical bog and mire species have also been found in the black forest region of Baden-Württemberg and elsewhere in Europe (Finderup Nielsen et al., 2021; Hallman et al., 2022; Kindlund & Tyler, 2023; Sperle & Bruelheide, 2021; Tyler et al., 2020).

Species of meadows and pastures may also be negatively affected by increases in nutrients and by an overall intensification of management (Newbold et al., 2015; Wesche et al., 2012). Indeed, the biggest losers included species of moist to wet grasslands in SH and BW and species of mesic grasslands in HH. These results are consistent with declines of these species in other regions of Germany (Diekmann et al., 2019; Wesche et al., 2012).

Dune and coastal habitats have one of the worst conservation statuses in Europe, with a similar situation in Germany (European Environment Agency, 2020; Finck et al., 2017; Hodapp et al., 2024). In SH, especially grey dunes, coastal dune heaths and moist dune valleys are in a bad condition (Landesamt für Landwirtschaft, Umwelt und ländliche Räume, 2020). Species characteristic of coastal and marine habitats generally declined in all states, including in HH and BW, which have no marine coastline and only few brackish water bodies. The latter was possible because this category included species common in both coastal and non-coastal habitat types, such as *Festuca rubra* agg. Those species were likely often assigned as coastal species because the protected coastal sites had more often complete species lists available, including non-characteristic species, compared with other habitat sites. However, these declining species share their tolerance to salt, drought, extreme temperatures, nutrient deficiency, and disturbance (Martínez et al., 2013). In the Netherlands, changes in drainage regimes, grazing pressure, and atmospheric nitrogen deposition led to a loss of characteristic species in coastal dunes over 50 years (Kuiters et al., 2009). Similar reasons for the decline of species characteristic of coastal and marine habitats are likely for Germany. In addition, human outdoor-activities (including trampling), pollution, and modifications of the coastline by recreational structures and dykes threat these species at the coast state SH (Heinze et al., 2019). As most saltmarshes had to be excluded from the analyses (Supplementary Methods), many coastal sites at the North Sea are missing from the analysis of SH. Trends in those mostly protected areas, which are generally in a good condition (Landesamt für Landwirtschaft, Umwelt und ländliche Räume, 2020), might be more stable than in the sites used for our analysis.

Endangered species showed the most pronounced declines in BW and SH. The lack of significant negative trends for those species in HH can be explained by fewer endangered species left in the city. A key finding was the overall decrease in the occurrence probability of non-threatened species in all states. This underpins recent observations that biodiversity decline not only affects endangered plant species but also previously moderately common species (Jansen et al., 2020).

In accordance with our expectations, the set of across-states winner species included species of scrubs, copses and field hedges, with also forest species showing mostly positive trends. Furthermore, we found increases of woody plants within several different habitat types in SH and BW. These positive trends imply a woody encroachment across the landscape, which has also been observed in many other formerly extensively used habitats (mainly grasslands and heathlands) in Europe (Buitenwerf et al., 2018; European Environment Agency, 2017; Navarro & Pereira, 2015; Prévosto et al., 2011). Forest species have also been found to increase across Germany (Jandt et al., 2022) and in the German biodiversity assessment (Müller et al., 2024).

As expected, neophytes showed significant increases for BW and SH in accordance with positive trends of non-native species reported across Germany in other studies (Eichenberg et al., 2021). In HH, the lack of an overall significant trend might be due to a long history of neophyte introductions in HH, with some long-established neophyte species nowadays being similarly negatively affected by land use changes as native ones. The spread of neophytes might have been more pronounced in close-to-nature habitat types over the last decades, which are more abundant in BW and SH.

We found no opposing significant trends in occurrence probability of species groups in the different states. Furthermore, species showing a significant trend in two respective states had generally positively correlated trends. Overall, the largely consistent trends of species groups across the states suggest common drivers of biodiversity change across the studied parts of Germany. Thus, on a national scale, these common drivers appear to override regional and local differences in land use history, habitat types and environmental conditions. We expect other parts of Germany to show similar species trends, as despite some regional differences, they generally share a similar history of land use change, including agricultural intensification, shrub encroachment, climate change, nitrogen deposition, and urban expansion (Finck et al., 2017).

Consistent trends were also found for trait-based species groups within different regions and semi-natural habitat types in Denmark, indicating that common drivers led to more fertile and less disturbed conditions across the landscape (Timmermann et al., 2015). Similarly, mostly consistent trends were found across different parishes in Sweden, with some dissimilarities due to different environmental or land use changes (Hallman et al., 2022).

In line with our expectations, trends within habitat types in SH and BW showed that especially species that preferred a certain habitat type decreased in occurrence probability within that habitat type. The decline of characteristic species has been a common pattern found in many different habitat types across Europe over the past decades (Britton et al., 2009; Heinrichs & Schmidt, 2017; Klinkovská et al., 2023; Kuiters et al., 2009). However, this pattern was not encountered for trends in frequency or trends in HH. The latter is probably due to a limited amount of data for HH. Generally, we found higher losses indicated by the probability of species to occur compared with their actual losses in frequency. As frequencies are based on the raw number of observations, they suffer from a bias towards positive change brought about by the higher number of polygons in the second time interval. This bias was accompanied by uncertainties caused by incomplete species lists, which we tried to account for by only reporting robust trends. However, this conservative approach excluded many trends in frequency. Still, we included the frequency metric in our analysis to demonstrate its limits when being applied to habitat mapping data. Instead, Beals occurrence probabilities offer a more reliable change metric. In addition, as Beals occurrence probabilities are based on all present species at a site, they can be interpreted as an early warning of species loss in response to habitat degradation (Bruelheide et al., 2020). Thus, it seems that while a few characteristic species might still persist in degraded sites across the states (no frequency loss), conditions are not favourable anymore (negative Beals trends caused by the absence of typically co-occurring species) and species might after an extinction debt be completely lost from many sites (Hylander & Ehrlen, 2013; Kuussaari et al., 2009).

While forest species increased in many uncharacteristic habitat types in our investigated states, the situation of those species within dry to moderately moist forests seems to differ between states. We found mostly negative trends of those forest species in SH, mixed trends in BW, but positive trends in HH. Thus, while species turnover seems to occur in all states, the trend goes towards less characteristic species in dry to moderately moist forests in SH but indicates recovery of characteristic species in dry to moderately moist forests in HH from being more intensively used in the past. This could be further explained by a higher proportion of forests that are protected in HH (90%) compared with the other states (59% and 79% for SH and BW, respectively, based on the polygons used for analysis).

While common challenges of habitat mapping data for trend analysis can be overcome by appropriate data cleaning and applying metrics that account for incomplete species records, some limitations remain. First, we want to stress that frequency trends have to be taken with caution given their susceptibility to incomplete species recordings, while Beals trends are generally more robust (Bruelheide et al., 2020). Second, while our study included sites that underwent habitat transition over the study period, those were almost exclusively sites that kept their protection status and had species lists available from both time intervals. Thus, severely degraded sites were mostly excluded. Third, it is not possible to derive trends for most rare species, given their incomplete representation in the co-occurrence matrices used for Beals (Bruelheide et al., 2021). In conclusion, given the bias towards positive trends, difficulties to detect trends for most rare species, and the exclusion of most severely degraded sites, our estimates are highly conservative and underestimate the amount and magnitude of negative trends in all states. The “true” trends can be expected to be even worse. Still, given the consistency of species group trends that were analysed independently across the three federal states and with other studies, the overall trends we derived can be considered robust.

Despite its heterogeneous quality, we demonstrated that habitat mapping data can be used to determine the winner and loser species of last decades’ biodiversity change. The mostly consistent trends of species groups we found across the three federal states point to common drivers of biodiversity change in different regions of Germany. Identifying those exact drivers for different habitat types needs so far unavailable fine-scale data for multiple drivers, especially concerning land management and interventions, nutrient enrichment and hydrological changes. Furthermore, to analyse changes for very rare species or changes in species richness and composition, we need systematic monitoring efforts of species occurrences across habitat types and regions. Our trends are mostly consistent with findings from local-scale analyses derived from other data sources and regions in Europe. Our analyses demonstrate the importance of including the landscape scale and not only to rely on trends on coarser scales, as this is the relevant scale at which conservation measures are applied to achieve the goals set by, for example, the EU Biodiversity Strategy.

## Acknowledgements

We are grateful for all the surveyors, coordinators, and other people involved in the decades-long habitat mapping programs in the states Schleswig-Holstein, Hamburg and Baden-Württemberg. This study is part of the work of the sMon project (Trend analysis of biodiversity data in Germany) of the German Centre for Integrative Biodiversity Research (iDiv) Halle-Jena-Leipzig. sMon appreciates funding from the German Research Foundation (DFG FZT 118, project number 202548816). L.L. acknowledges support by the Graduate scholarship program of Saxony-Anhalt. Projekt DEAL enabled and organized Open Access funding. The authors declare no conflicts of interest.

## Data Accessibility Statement

Polygon data are publicly available via https://umweltanwendungen.schleswig-holstein.de/fachauswertungweb/ for the current mappings in SH, via https://suche.transparenz.hamburg.de/dataset/biotopkataster-hamburg9 for both old and current mappings in HH, and via https://udo.lubw.baden-wuerttemberg.de/public/index.xhtml for the current mappings in BW. As all data belong to the federal state agencies, the full dataset is in parts restricted to be published. This concerns data from previous mappings for the open land for BW and observations of endangered species in BW. All other data are currently in the process of being archived in the iDiv Biodiversity Data Portal (iBDP, https://idata.idiv.de/) and will be publicly available upon acceptance.

