## Appendix S1 for "Loss of characteristic species across German federal states detected by repeated mapping of protected habitats"

**Associated paper**: Loss of characteristic species across German federal states detected by repeated mapping of protected habitats

**Authors**: Lina Lüttgert, Samuel Heisterkamp, Florian Jansen, Rico Kaufmann, Simon Kellner, Reinhard Arnold Klenke, Silke Lütt, Gunnar Seidler, Axel Wedler, Ronja Wörmann, Helge Bruelheide

**Supplementary Methods – Data cleaning of SH habitat mapping data**

This part provides additional information on the data cleaning specific to the habitat mapping data of the federal state Schleswig-Holstein (SH).

Habitat mapping in SH was mainly carried out in two mapping campaigns, with the first one running from 1978-1993 and the recent one from 2014-2020. However, we also included data coming from side mapping projects, which resulted in a total time interval from 1977-2021. We divided the data into two time intervals: *t*_1_ from 1977-2005 and *t*_2_ from 2007-2021, with no data available for 2006.

While as for most federal states, the habitat mapping in SH includes mapping all protected habitat types, in SH also other ‘potentially valuable’ and some other unprotected habitat types are mapped. However, species lists are usually only compiled for protected habitat types. Thus, only 45% of our intersecting polygons had a species list attached.

Next to the two main mapping campaigns, data was available from the additional mapping projects for saltmarshes (Salzwiesen-Kartierung), lakes (Seen-Kartierung), high-nature value grasslands (Wertgrünland-Kartierung) and islands (Inselkartierung). They all followed the same mapping keys and thus were generally included in our analysis. However, we excluded polygons from the saltmarshes project, as here often the same species list is attached to many polygons (>50) and thus we would have treated many different sites as if they were one.

After intersecting all (remaining) polygons, we excluded intersections covering less than 5% of the area of both polygons. This deviates from the methods for BW and HH, where only intersections covering less than 5% of the area of either polygon were excluded. However, for SH this stricter method would exclude many cases where one large *t*_1_ polygon intersects with many small *t*_2_ polygons.

For the same reason, we set the proportion of a polygon’s area that had to intersect with polygons from the other time interval to 50%. Since all habitat sites have been remapped, the risk to include not yet remapped polygons is negligible (in contrast to BW for example). For BW and HH we set this threshold to 75% and 95% respectively. Already using 50% as a threshold for SH excluded many intersections, especially since no longer protected sites mostly did not have a species list attached for the *t*_2_ record. Thus, all once but no more protected habitats’ species lists could not be compared to anything in *t*_2_ and thus those probable species losses could not go into analyses. For approach *t*_1_ → *t*_2_, only 76% of intersections (and 66% of *t*_1_ polygons) remained after deleting polygons that have not been remapped with a species list by at least 50% of their area. For approach *t*_2_ → *t*_1_, 84% of intersections (and 83% of *t*_2_ polygons) remained.

There were cases where a polygon of one time interval intersected with several neighbouring polygons from several years from the other time interval. This was the case because 1) some polygons (mainly from *t*_1_) lay on the border of two districts and those districts were mapped in different years in the other time interval (mainly *t*_2_); 2) of different mapping projects that ran in different years; and 3) case-specific remappings of sites. Thus, for those cases, we always compared a polygon from one year with several polygons from several years.

While for the species analysis across all polygons we included polygons with no or insufficient habitat type information, we had to reduce the dataset to polygons with sufficient habitat type information for trends within habitat types and the assignment of species’ preferred habitat types. Habitat mapping keys changed over time, with formerly 69 and recently 577 habitat types (Landesamt für Landwirtschaft Umwelt und ländliche Räume des Landes Schleswig-Holstein (LLUR), 2017; Landesamt für Naturschutz und Landschaftspflege Schleswig-Holstein, 1991). To make the types between time intervals, but also between all three states comparable, we aggregated all 646 detailed types to 12 broad habitat type groups. There were no species lists available for the two additional groups out of the 14 that were used for analysis, i.e. for anthropogenic habitats and farmland. See table “habitat_types_all_states.csv” for all assignments. Generally, the mapping key includes two types of habitat types, one is based on the structure of a location and one is based on the vegetation of a site. Polygons that have habitat types from both categories assigned can have a summed cover of all habitat types of up to 200%. Cases with a summed cover of >200% were excluded if the mapped habitat types were from different broad habitat type groups, otherwise the cover was set to 100%. Polygons without a main habitat type covering min. 51% of its area had to be excluded.

For taxonomic harmonization, all species were aggregated to the section level and cleaned in accordance with the species lists from the other two states. For SH, this resulted in 1301 species.

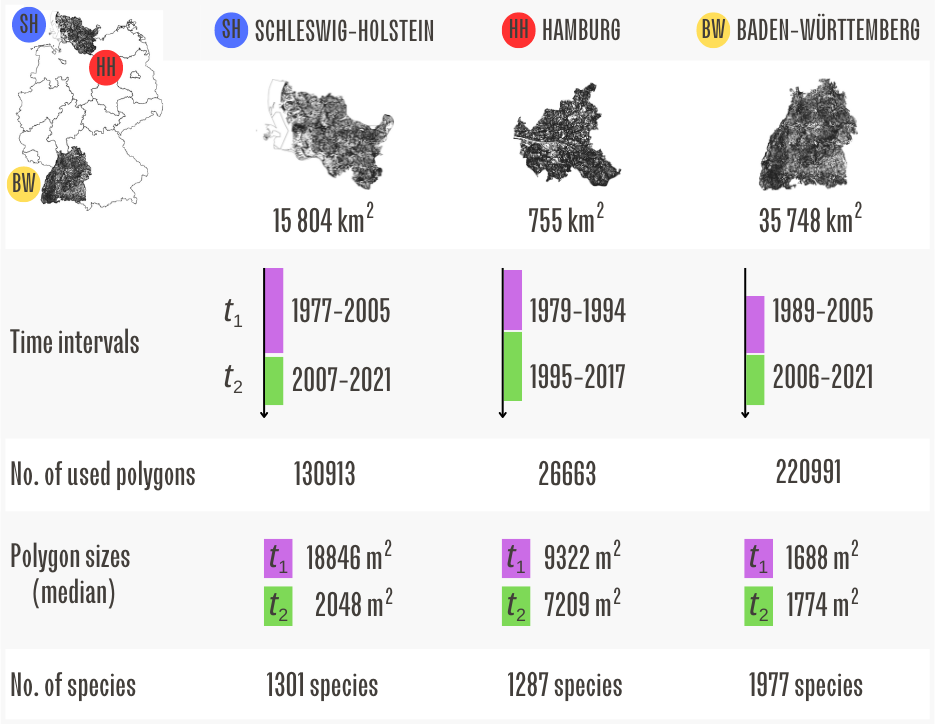
 **Figure S1.1** Overview of the three federal states used for analysis. Data for area of states is based on Statistisches Bundesamt 2023. All information on the mapping data is based on the adjusted data set.

**Table S1.1** Polygon numbers and sizes for the three states and two time intervals.

| **Federal state** | **Time** | **Number polygons** | **Polygon area [m^2^]** | | | |
| --- | --- | --- | --- | --- | --- | --- |
|  |  |  | **mean** | **median** | **min** | **max** |
| SH | both | 130913 | 18661 | 2547 | 1 | 29025586 |
|  | *t*_1_ | 13435 | 105953 | 18846 | 2 | 28816471 |
|  | *t*_2_ | 117478 | 8590 | 2048 | 1 | 29025586 |
| HH | both | 26663 | 21170 | 7421 | 1 | 2111000 |
|  | *t*_1_ | 2769 | 30974 | 9322 | 1 | 2025152 |
|  | *t*_2_ | 23894 | 20034 | 7209 | 12 | 2111000 |
| BW | both | 220991 | 6898 | 1731 | 2 | 7403074 |
|  | *t*_1_ | 108865 | 7182 | 1688 | 5 | 6999119 |
|  | *t*_2_ | 112222 | 6626 | 1774 | 2 | 7403074 |


Notes: Based on the complete cleaned data set, which still includes non-intersecting polygons as they were included for
the calculation of the Beals co-occurrence matrix and species’ habitat type preferences.

**(a)	(b)	(c)**

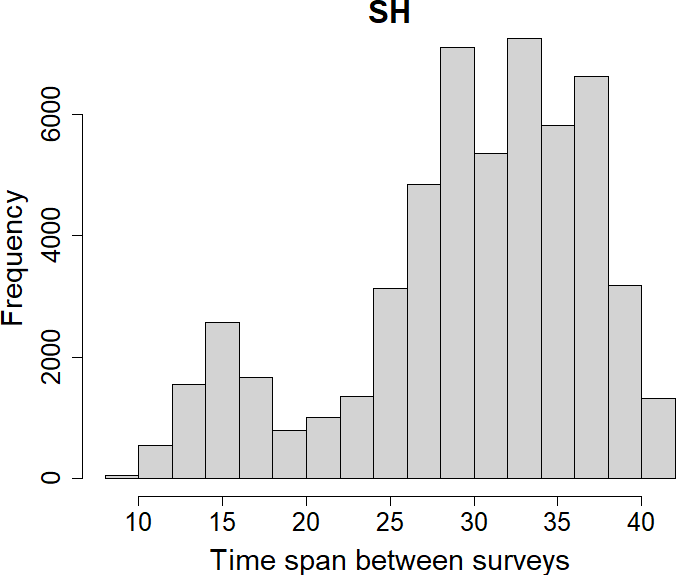

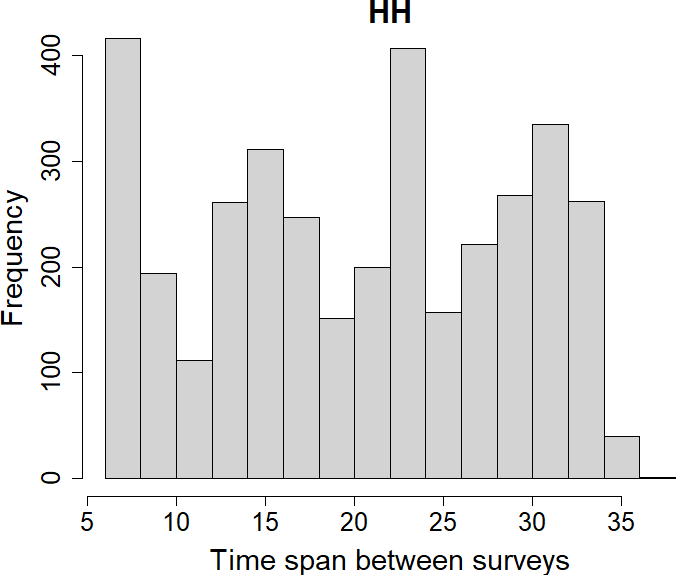

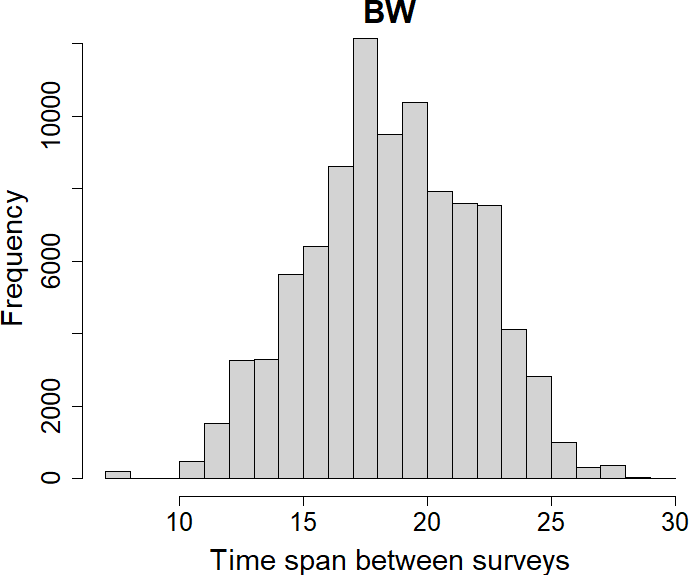

**Figure S1.2** Time spans between surveys *t*_1_ and *t*_2_ for **(a)** SH, **(b)** HH and **(c)** BW. Based on intersection data used for trend analysis. For SH, in cases where polygons of several years intersected with one polygon of the other time interval, mean values of those mapped polygons’ years were calculated.

**(a)	(b)	(c)	(d)**

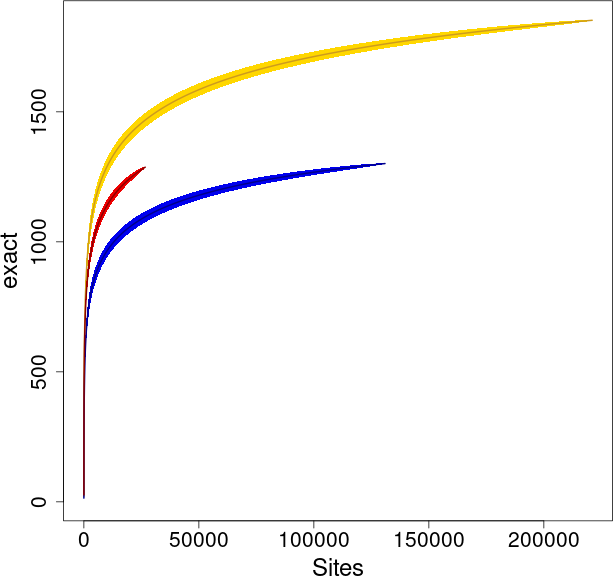

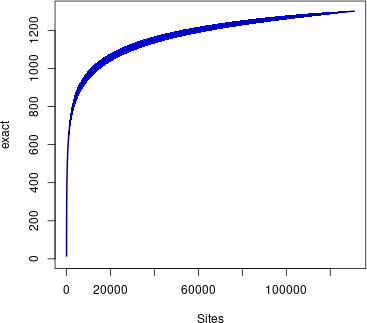

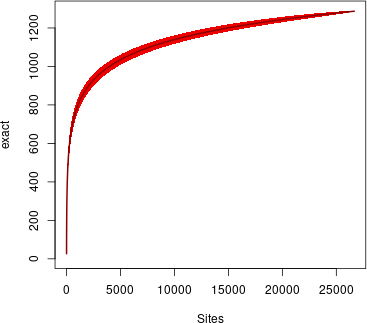

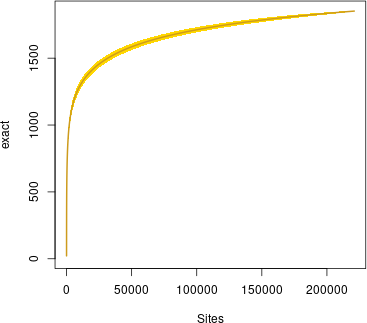

**Figure S1.3** Accumulation curve of species recorded in polygons for **(a)** all states, separated by colour with SH in blue, HH in red and BW in yellow; and separately for **(b)** SH, **(c)** HH, **(d)** BW. Sites on the x-axis refer to the number of polygons sampled and exact on the y-axis refers to the number of species recorded.

**Table S1.2** The 14 habitat type groups used for analysis.

| **Habitat type group** | **SH** | **HH** | **BW** |
| --- | --- | --- | --- |
| Coastal and marine habitats | x |  |  |
| Heaths, inland dunes and semi-natural grasslands | x | x | x |
| Mesic grasslands | x | x | x |
| Moist to wet grasslands | x | x | x |
| Fresh water vegetation, springs and reeds | x |  | x |
| Linear and running surface waters | x | x |  |
| Standing waters | x | x |  |
| Bogs, transition mires, marshes and fens | x | x | x |
| Scrubs, copses and field hedges | x | x | x |
| Dry to moderately moist forests | x | x | x |
| Moist to wet forests | x | x | x |
| Ruderal, fringe and tall forb communities, clearings | x | x | x |
| Farmland |  | x |  |
| Anthropogenic |  | x |  |

.
 Notes: With “x” in the columns SH, HH, and BW indicating that a group was mapped in the respective federal state.

**Table S1.3** All habitat types included per habitat type group in each state. For a table including the German habitat type names see https://doi.org/10.6084/m9.figshare.28507244.v1.

| **Code** | **Group** | **State** | **Time** |
| --- | --- | --- | --- |
| AA | Farmland | SH | 1 |
| AB | Anthropogenic | SH | 1 |
| AF | Dry to moderately moist forests | SH | 1 |
| AG | Mesic grasslands | SH | 1 |
| AK | Dry to moderately moist forests | SH | 1 |
| AP | Dry to moderately moist forests | SH | 1 |
| AR | Ruderal, fringe and tall forb communities, clearings | SH | 1 |
| AV | Anthropogenic | SH | 1 |
| DB | Heaths, inland dunes and semi-natural grasslands | SH | 1 |
| DG | Coastal and marine habitats | SH | 1 |
| DH | Coastal and marine habitats | SH | 1 |
| DK | Coastal and marine habitats | SH | 1 |
| DN | Coastal and marine habitats | SH | 1 |
| FA | Linear and running surface waters | SH | 1 |
| FB | Linear and running surface waters | SH | 1 |
| FF | Linear and running surface waters | SH | 1 |
| FQ | Fresh water vegetation, springs and reeds | SH | 1 |
| GA | Mesic grasslands | SH | 1 |
| GC | Heaths, inland dunes and semi-natural grasslands | SH | 1 |
| GF | Moist to wet grasslands | SH | 1 |
| GH | Ruderal, fringe and tall forb communities, clearings | SH | 1 |
| GM | Heaths, inland dunes and semi-natural grasslands | SH | 1 |
| GP | Ruderal, fringe and tall forb communities, clearings | SH | 1 |
| GS | Bogs, transition mires, marshes and fens | SH | 1 |
| KA | Coastal and marine habitats | SH | 1 |
| KB | Coastal and marine habitats | SH | 1 |
| KD | Coastal and marine habitats | SH | 1 |
| KH | Coastal and marine habitats | SH | 1 |
| KO | Coastal and marine habitats | SH | 1 |
| KS | Coastal and marine habitats | SH | 1 |
| KV | Coastal and marine habitats | SH | 1 |
| KW | Coastal and marine habitats | SH | 1 |
| MB | Moist to wet forests | SH | 1 |
| MH | Bogs, transition mires, marshes and fens | SH | 1 |
| MM | Bogs, transition mires, marshes and fens | SH | 1 |
| MS | Bogs, transition mires, marshes and fens | SH | 1 |
| MT | Bogs, transition mires, marshes and fens | SH | 1 |
| MZ | Bogs, transition mires, marshes and fens | SH | 1 |
| SD | Standing waters | SH | 1 |
| SG | Standing waters | SH | 1 |
| SK | Standing waters | SH | 1 |
| SL | Standing waters | SH | 1 |
| SM | Standing waters | SH | 1 |
| SS | Coastal and marine habitats | SH | 1 |
| ST | Standing waters | SH | 1 |
| SW | Standing waters | SH | 1 |
| UF | Coastal and marine habitats | SH | 1 |
| VA | Scrubs, copses and field hedges | SH | 1 |
| VG | Fresh water vegetation, springs and reeds | SH | 1 |
| VH | Bogs, transition mires, marshes and fens | SH | 1 |
| VQ | Fresh water vegetation, springs and reeds | SH | 1 |
| VR | Fresh water vegetation, springs and reeds | SH | 1 |
| WA | Moist to wet forests | SH | 1 |
| WB | Moist to wet forests | SH | 1 |
| WE | Moist to wet forests | SH | 1 |
| WG | Scrubs, copses and field hedges | SH | 1 |
| WH | Scrubs, copses and field hedges | SH | 1 |
| WK | Dry to moderately moist forests | SH | 1 |
| WL | Dry to moderately moist forests | SH | 1 |
| WM | Dry to moderately moist forests | SH | 1 |
| WN | Dry to moderately moist forests | SH | 1 |
| WQ | Dry to moderately moist forests | SH | 1 |
| WR | NA | SH | 1 |
| BM | Bogs, transition mires, marshes and fens | SH | 1 |
| F | Linear and running surface waters | SH | 1 |
| GMma | Mesic grasslands | SH | 1 |
| GH/AR | Ruderal, fringe and tall forb communities, clearings | SH | 1 |
| WAs | Moist to wet forests | SH | 1 |
| (GP) | Ruderal, fringe and tall forb communities, clearings | SH | 1 |
| AA | Farmland | SH | 2 |
| AG | Farmland | SH | 2 |
| AO | Farmland | SH | 2 |
| AB | Farmland | SH | 2 |
| SKv | Anthropogenic | SH | 2 |
| SVs | Anthropogenic | SH | 2 |
| SXb | Anthropogenic | SH | 2 |
| SXr | Anthropogenic | SH | 2 |
| WFn | Dry to moderately moist forests | SH | 2 |
| GA | Mesic grasslands | SH | 2 |
| GAe | Mesic grasslands | SH | 2 |
| GAy | Mesic grasslands | SH | 2 |
| GY | NA | SH | 2 |
| GYy | Mesic grasslands | SH | 2 |
| WMx | Dry to moderately moist forests | SH | 2 |
| WLx | Dry to moderately moist forests | SH | 2 |
| RHg | Ruderal, fringe and tall forb communities, clearings | SH | 2 |
| RHn | Ruderal, fringe and tall forb communities, clearings | SH | 2 |
| RHy | Ruderal, fringe and tall forb communities, clearings | SH | 2 |
| SVx | Ruderal, fringe and tall forb communities, clearings | SH | 2 |
| SLl | Ruderal, fringe and tall forb communities, clearings | SH | 2 |
| SLf | Ruderal, fringe and tall forb communities, clearings | SH | 2 |
| SVu | Anthropogenic | SH | 2 |
| SVp | Anthropogenic | SH | 2 |
| SVy | Anthropogenic | SH | 2 |
| SVt | Anthropogenic | SH | 2 |
| TBc | Heaths, inland dunes and semi-natural grasslands | SH | 2 |
| TBe | Heaths, inland dunes and semi-natural grasslands | SH | 2 |
| TBa | Heaths, inland dunes and semi-natural grasslands | SH | 2 |
| TBd | Heaths, inland dunes and semi-natural grasslands | SH | 2 |
| KDo | Coastal and marine habitats | SH | 2 |
| KDv | Coastal and marine habitats | SH | 2 |
| KDm | Coastal and marine habitats | SH | 2 |
| KDw | Coastal and marine habitats | SH | 2 |
| KDl | Coastal and marine habitats | SH | 2 |
| KDr | Coastal and marine habitats | SH | 2 |
| KDn | Coastal and marine habitats | SH | 2 |
| KDs | Coastal and marine habitats | SH | 2 |
| KDg | Coastal and marine habitats | SH | 2 |
| KDe | Coastal and marine habitats | SH | 2 |
| KDc | Coastal and marine habitats | SH | 2 |
| KHr | Coastal and marine habitats | SH | 2 |
| KHh | Coastal and marine habitats | SH | 2 |
| KHs | Coastal and marine habitats | SH | 2 |
| KHt | Coastal and marine habitats | SH | 2 |
| KHq | Coastal and marine habitats | SH | 2 |
| KHp | Coastal and marine habitats | SH | 2 |
| KHx | Coastal and marine habitats | SH | 2 |
| KHg | Coastal and marine habitats | SH | 2 |
| KDy | Coastal and marine habitats | SH | 2 |
| XSw | Coastal and marine habitats | SH | 2 |
| KDx | Coastal and marine habitats | SH | 2 |
| KP | Coastal and marine habitats | SH | 2 |
| KPc | Coastal and marine habitats | SH | 2 |
| KPi | Coastal and marine habitats | SH | 2 |
| KPl | Coastal and marine habitats | SH | 2 |
| KPr | Coastal and marine habitats | SH | 2 |
| KPy | Coastal and marine habitats | SH | 2 |
| KM | Coastal and marine habitats | SH | 2 |
| KMy | Coastal and marine habitats | SH | 2 |
| KMf | Coastal and marine habitats | SH | 2 |
| KMr | Coastal and marine habitats | SH | 2 |
| KMm | Coastal and marine habitats | SH | 2 |
| KMt | Coastal and marine habitats | SH | 2 |
| KMh | Coastal and marine habitats | SH | 2 |
| KMw | Coastal and marine habitats | SH | 2 |
| KMb | Coastal and marine habitats | SH | 2 |
| KMe | Coastal and marine habitats | SH | 2 |
| FFa | Linear and running surface waters | SH | 2 |
| FBa | Linear and running surface waters | SH | 2 |
| FBf | Linear and running surface waters | SH | 2 |
| FBg | Linear and running surface waters | SH | 2 |
| FBn | Linear and running surface waters | SH | 2 |
| FBt | Linear and running surface waters | SH | 2 |
| FBx | Linear and running surface waters | SH | 2 |
| FUb | Linear and running surface waters | SH | 2 |
| FUg | Linear and running surface waters | SH | 2 |
| FLf | Linear and running surface waters | SH | 2 |
| FLr | Linear and running surface waters | SH | 2 |
| FLs | Linear and running surface waters | SH | 2 |
| FLw | Linear and running surface waters | SH | 2 |
| FLy | Linear and running surface waters | SH | 2 |
| FLg | Linear and running surface waters | SH | 2 |
| FGg | Linear and running surface waters | SH | 2 |
| FFf | Linear and running surface waters | SH | 2 |
| FFg | Linear and running surface waters | SH | 2 |
| FFn | Linear and running surface waters | SH | 2 |
| FFt | Linear and running surface waters | SH | 2 |
| FFx | Linear and running surface waters | SH | 2 |
| YQ | Fresh water vegetation, springs and reeds | SH | 2 |
| GM | Mesic grasslands | SH | 2 |
| GMf | Mesic grasslands | SH | 2 |
| GMm | Mesic grasslands | SH | 2 |
| GMt | Mesic grasslands | SH | 2 |
| HOm | Mesic grasslands | SH | 2 |
| TF | Heaths, inland dunes and semi-natural grasslands | SH | 2 |
| TFd | Heaths, inland dunes and semi-natural grasslands | SH | 2 |
| TFg | Heaths, inland dunes and semi-natural grasslands | SH | 2 |
| TFn | Heaths, inland dunes and semi-natural grasslands | SH | 2 |
| TFt | Heaths, inland dunes and semi-natural grasslands | SH | 2 |
| TH | Heaths, inland dunes and semi-natural grasslands | SH | 2 |
| THd | Heaths, inland dunes and semi-natural grasslands | SH | 2 |
| THs | Heaths, inland dunes and semi-natural grasslands | SH | 2 |
| THx | Heaths, inland dunes and semi-natural grasslands | SH | 2 |
| THg | Heaths, inland dunes and semi-natural grasslands | SH | 2 |
| THt | Heaths, inland dunes and semi-natural grasslands | SH | 2 |
| THw | Heaths, inland dunes and semi-natural grasslands | SH | 2 |
| GN | Moist to wet grasslands | SH | 2 |
| GNp | Moist to wet grasslands | SH | 2 |
| GNa | Moist to wet grasslands | SH | 2 |
| GNb | Moist to wet grasslands | SH | 2 |
| GNm | Moist to wet grasslands | SH | 2 |
| GNr | Moist to wet grasslands | SH | 2 |
| GNh | Moist to wet grasslands | SH | 2 |
| GF | Moist to wet grasslands | SH | 2 |
| GFb | Moist to wet grasslands | SH | 2 |
| GFc | Moist to wet grasslands | SH | 2 |
| GFf | Moist to wet grasslands | SH | 2 |
| GFr | Moist to wet grasslands | SH | 2 |
| GYj | Moist to wet grasslands | SH | 2 |
| GYn | Moist to wet grasslands | SH | 2 |
| GYf | Moist to wet grasslands | SH | 2 |
| RH | Ruderal, fringe and tall forb communities, clearings | SH | 2 |
| RHw | Ruderal, fringe and tall forb communities, clearings | SH | 2 |
| RHu | Ruderal, fringe and tall forb communities, clearings | SH | 2 |
| RHs | Ruderal, fringe and tall forb communities, clearings | SH | 2 |
| RHf | Ruderal, fringe and tall forb communities, clearings | SH | 2 |
| RHm | Ruderal, fringe and tall forb communities, clearings | SH | 2 |
| RHr | Ruderal, fringe and tall forb communities, clearings | SH | 2 |
| RHt | Ruderal, fringe and tall forb communities, clearings | SH | 2 |
| RHx | Ruderal, fringe and tall forb communities, clearings | SH | 2 |
| RHp | Ruderal, fringe and tall forb communities, clearings | SH | 2 |
| TRb | Heaths, inland dunes and semi-natural grasslands | SH | 2 |
| TRm | Heaths, inland dunes and semi-natural grasslands | SH | 2 |
| TRn | Heaths, inland dunes and semi-natural grasslands | SH | 2 |
| TRj | Heaths, inland dunes and semi-natural grasslands | SH | 2 |
| TRo | Heaths, inland dunes and semi-natural grasslands | SH | 2 |
| TRs | Heaths, inland dunes and semi-natural grasslands | SH | 2 |
| TRh | Heaths, inland dunes and semi-natural grasslands | SH | 2 |
| TRy | Heaths, inland dunes and semi-natural grasslands | SH | 2 |
| TR | Heaths, inland dunes and semi-natural grasslands | SH | 2 |
| RP | Ruderal, fringe and tall forb communities, clearings | SH | 2 |
| RPa | Ruderal, fringe and tall forb communities, clearings | SH | 2 |
| RPr | Ruderal, fringe and tall forb communities, clearings | SH | 2 |
| NS | Bogs, transition mires, marshes and fens | SH | 2 |
| NSa | Bogs, transition mires, marshes and fens | SH | 2 |
| NSb | Bogs, transition mires, marshes and fens | SH | 2 |
| NSj | Bogs, transition mires, marshes and fens | SH | 2 |
| NSf | Bogs, transition mires, marshes and fens | SH | 2 |
| NSc | Bogs, transition mires, marshes and fens | SH | 2 |
| NSr | Bogs, transition mires, marshes and fens | SH | 2 |
| NSy | Bogs, transition mires, marshes and fens | SH | 2 |
| XK | Coastal and marine habitats | SH | 2 |
| XKf | Coastal and marine habitats | SH | 2 |
| XKh | Coastal and marine habitats | SH | 2 |
| XKn | Coastal and marine habitats | SH | 2 |
| XKo | Coastal and marine habitats | SH | 2 |
| XSn | Coastal and marine habitats | SH | 2 |
| XSo | Coastal and marine habitats | SH | 2 |
| KNa | Coastal and marine habitats | SH | 2 |
| KNh | Coastal and marine habitats | SH | 2 |
| KNv | Coastal and marine habitats | SH | 2 |
| KNw | Coastal and marine habitats | SH | 2 |
| KNo | Coastal and marine habitats | SH | 2 |
| Knp | Coastal and marine habitats | SH | 2 |
| KNk | Coastal and marine habitats | SH | 2 |
| KNt | Coastal and marine habitats | SH | 2 |
| KNx | Coastal and marine habitats | SH | 2 |
| KNy | Coastal and marine habitats | SH | 2 |
| KNd | Coastal and marine habitats | SH | 2 |
| KOa | Coastal and marine habitats | SH | 2 |
| KOd | Coastal and marine habitats | SH | 2 |
| KOw | Coastal and marine habitats | SH | 2 |
| KOf | Coastal and marine habitats | SH | 2 |
| KOj | Coastal and marine habitats | SH | 2 |
| KOm | Coastal and marine habitats | SH | 2 |
| KOo | Coastal and marine habitats | SH | 2 |
| KOs | Coastal and marine habitats | SH | 2 |
| KOr | Coastal and marine habitats | SH | 2 |
| KOy | Coastal and marine habitats | SH | 2 |
| KOc | Coastal and marine habitats | SH | 2 |
| KOq | Coastal and marine habitats | SH | 2 |
| KOl | Coastal and marine habitats | SH | 2 |
| KOp | Coastal and marine habitats | SH | 2 |
| KOt | Coastal and marine habitats | SH | 2 |
| KOh | Coastal and marine habitats | SH | 2 |
| KSs | Coastal and marine habitats | SH | 2 |
| KNs | Coastal and marine habitats | SH | 2 |
| KWg | Coastal and marine habitats | SH | 2 |
| KQs | Coastal and marine habitats | SH | 2 |
| KQd | Coastal and marine habitats | SH | 2 |
| KQu | Coastal and marine habitats | SH | 2 |
| KQr | Coastal and marine habitats | SH | 2 |
| KWw | Coastal and marine habitats | SH | 2 |
| KWf | Coastal and marine habitats | SH | 2 |
| MWb | Moist to wet forests | SH | 2 |
| MWk | Moist to wet forests | SH | 2 |
| MWs | Moist to wet forests | SH | 2 |
| MDb | Moist to wet forests | SH | 2 |
| MSr | Bogs, transition mires, marshes and fens | SH | 2 |
| MSz | Bogs, transition mires, marshes and fens | SH | 2 |
| MSs | Bogs, transition mires, marshes and fens | SH | 2 |
| MW | Moist to wet forests | SH | 2 |
| MSy | Bogs, transition mires, marshes and fens | SH | 2 |
| MDm | Bogs, transition mires, marshes and fens | SH | 2 |
| MSt | Bogs, transition mires, marshes and fens | SH | 2 |
| MDe | Bogs, transition mires, marshes and fens | SH | 2 |
| MRe | Bogs, transition mires, marshes and fens | SH | 2 |
| MRb | Bogs, transition mires, marshes and fens | SH | 2 |
| MRm | Bogs, transition mires, marshes and fens | SH | 2 |
| MRs | Bogs, transition mires, marshes and fens | SH | 2 |
| MRt | Bogs, transition mires, marshes and fens | SH | 2 |
| MRj | Bogs, transition mires, marshes and fens | SH | 2 |
| MRy | Bogs, transition mires, marshes and fens | SH | 2 |
| MR | Bogs, transition mires, marshes and fens | SH | 2 |
| MHc | Bogs, transition mires, marshes and fens | SH | 2 |
| MHe | Bogs, transition mires, marshes and fens | SH | 2 |
| MHs | Bogs, transition mires, marshes and fens | SH | 2 |
| MHy | Bogs, transition mires, marshes and fens | SH | 2 |
| MH | Bogs, transition mires, marshes and fens | SH | 2 |
| FSk | Standing waters | SH | 2 |
| FS | Standing waters | SH | 2 |
| FK/FS | Standing waters | SH | 2 |
| KSe | Coastal and marine habitats | SH | 2 |
| KF | Coastal and marine habitats | SH | 2 |
| HRe | Scrubs, copses and field hedges | SH | 2 |
| NSs | Fresh water vegetation, springs and reeds | SH | 2 |
| NHs | Bogs, transition mires, marshes and fens | SH | 2 |
| NHy | Bogs, transition mires, marshes and fens | SH | 2 |
| WQe | Fresh water vegetation, springs and reeds | SH | 2 |
| YQk | Fresh water vegetation, springs and reeds | SH | 2 |
| YQs | Fresh water vegetation, springs and reeds | SH | 2 |
| YQf | Fresh water vegetation, springs and reeds | SH | 2 |
| YQt | Fresh water vegetation, springs and reeds | SH | 2 |
| KRs | Fresh water vegetation, springs and reeds | SH | 2 |
| KRb | Fresh water vegetation, springs and reeds | SH | 2 |
| KRg | Fresh water vegetation, springs and reeds | SH | 2 |
| KRy | Fresh water vegetation, springs and reeds | SH | 2 |
| FWs | Fresh water vegetation, springs and reeds | SH | 2 |
| FWg | Fresh water vegetation, springs and reeds | SH | 2 |
| FWb | Fresh water vegetation, springs and reeds | SH | 2 |
| NR | Fresh water vegetation, springs and reeds | SH | 2 |
| NRa | Fresh water vegetation, springs and reeds | SH | 2 |
| NRc | Fresh water vegetation, springs and reeds | SH | 2 |
| NRs | Fresh water vegetation, springs and reeds | SH | 2 |
| NRr | Fresh water vegetation, springs and reeds | SH | 2 |
| NRg | Fresh water vegetation, springs and reeds | SH | 2 |
| NRb | Fresh water vegetation, springs and reeds | SH | 2 |
| NRy | Fresh water vegetation, springs and reeds | SH | 2 |
| WAm | Moist to wet forests | SH | 2 |
| WAp | Moist to wet forests | SH | 2 |
| WAx | Moist to wet forests | SH | 2 |
| WAq | Moist to wet forests | SH | 2 |
| WAw | Moist to wet forests | SH | 2 |
| WAe | Moist to wet forests | SH | 2 |
| WAy | Moist to wet forests | SH | 2 |
| WA | Moist to wet forests | SH | 2 |
| WBm | Moist to wet forests | SH | 2 |
| WBp | Moist to wet forests | SH | 2 |
| WBx | Moist to wet forests | SH | 2 |
| WBz | Moist to wet forests | SH | 2 |
| WBb | Moist to wet forests | SH | 2 |
| WBe | Moist to wet forests | SH | 2 |
| WBy | Moist to wet forests | SH | 2 |
| WB | Moist to wet forests | SH | 2 |
| WEm | Moist to wet forests | SH | 2 |
| WEp | Moist to wet forests | SH | 2 |
| WEx | Moist to wet forests | SH | 2 |
| WEz | Moist to wet forests | SH | 2 |
| WEw | Moist to wet forests | SH | 2 |
| WEe | Moist to wet forests | SH | 2 |
| WEy | Moist to wet forests | SH | 2 |
| WE | Moist to wet forests | SH | 2 |
| MDg | Scrubs, copses and field hedges | SH | 2 |
| MRg | Scrubs, copses and field hedges | SH | 2 |
| MRw | Scrubs, copses and field hedges | SH | 2 |
| MDw | Scrubs, copses and field hedges | SH | 2 |
| WBw | Moist to wet forests | SH | 2 |
| HWb | Scrubs, copses and field hedges | SH | 2 |
| HWo | Scrubs, copses and field hedges | SH | 2 |
| HWw | Scrubs, copses and field hedges | SH | 2 |
| HWx | Scrubs, copses and field hedges | SH | 2 |
| HWy | Scrubs, copses and field hedges | SH | 2 |
| HW | Scrubs, copses and field hedges | SH | 2 |
| HAo | Scrubs, copses and field hedges | SH | 2 |
| HAx | Scrubs, copses and field hedges | SH | 2 |
| HAy | Scrubs, copses and field hedges | SH | 2 |
| HA | Scrubs, copses and field hedges | SH | 2 |
| HFb | Scrubs, copses and field hedges | SH | 2 |
| HFx | Scrubs, copses and field hedges | SH | 2 |
| HFz | Scrubs, copses and field hedges | SH | 2 |
| HFy | Scrubs, copses and field hedges | SH | 2 |
| HF | Scrubs, copses and field hedges | SH | 2 |
| HR | Scrubs, copses and field hedges | SH | 2 |
| HRo | Scrubs, copses and field hedges | SH | 2 |
| HRy | Scrubs, copses and field hedges | SH | 2 |
| HRx | Scrubs, copses and field hedges | SH | 2 |
| HRn | Scrubs, copses and field hedges | SH | 2 |
| HB | Scrubs, copses and field hedges | SH | 2 |
| HBw | Scrubs, copses and field hedges | SH | 2 |
| HBx | Scrubs, copses and field hedges | SH | 2 |
| HBt | Scrubs, copses and field hedges | SH | 2 |
| HBy | Scrubs, copses and field hedges | SH | 2 |
| WMu | Dry to moderately moist forests | SH | 2 |
| WMs | Dry to moderately moist forests | SH | 2 |
| WL | Dry to moderately moist forests | SH | 2 |
| WLi | Dry to moderately moist forests | SH | 2 |
| WLa | Dry to moderately moist forests | SH | 2 |
| WLs | Dry to moderately moist forests | SH | 2 |
| WLy | Dry to moderately moist forests | SH | 2 |
| WLq | Dry to moderately moist forests | SH | 2 |
| WLb | Dry to moderately moist forests | SH | 2 |
| WLk | Dry to moderately moist forests | SH | 2 |
| WM | Dry to moderately moist forests | SH | 2 |
| WMc | Dry to moderately moist forests | SH | 2 |
| WMe | Dry to moderately moist forests | SH | 2 |
| WMm | Dry to moderately moist forests | SH | 2 |
| WMy | Dry to moderately moist forests | SH | 2 |
| WLt | Dry to moderately moist forests | SH | 2 |
| XHk | Bogs, transition mires, marshes and fens | SH | 2 |
| AAu | Farmland | SH | 2 |
| AAw | Farmland | SH | 2 |
| AAb | Farmland | SH | 2 |
| AAj | Farmland | SH | 2 |
| AAe | Farmland | SH | 2 |
| AAy | Farmland | SH | 2 |
| AGb | Farmland | SH | 2 |
| AGg | Farmland | SH | 2 |
| AGy | Farmland | SH | 2 |
| AOb | Farmland | SH | 2 |
| AOo | Farmland | SH | 2 |
| AOw | Farmland | SH | 2 |
| AOy | Farmland | SH | 2 |
| ABw | Farmland | SH | 2 |
| ABb | Farmland | SH | 2 |
| FK | Standing waters | SH | 2 |
| FSd | Standing waters | SH | 2 |
| FSo | Standing waters | SH | 2 |
| FSi | Standing waters | SH | 2 |
| FSm | Standing waters | SH | 2 |
| FSe | Standing waters | SH | 2 |
| FSx | Standing waters | SH | 2 |
| FSy | Standing waters | SH | 2 |
| FSs | Standing waters | SH | 2 |
| FKd | Standing waters | SH | 2 |
| FKo | Standing waters | SH | 2 |
| FKi | Standing waters | SH | 2 |
| FKm | Standing waters | SH | 2 |
| FKe | Standing waters | SH | 2 |
| FKx | Standing waters | SH | 2 |
| FKy | Standing waters | SH | 2 |
| SVf | Anthropogenic | SH | 2 |
| FLk | Linear and running surface waters | SH | 2 |
| FW | Linear and running surface waters | SH | 2 |
| FGx | Linear and running surface waters | SH | 2 |
| FGy | Linear and running surface waters | SH | 2 |
| FX | Standing waters | SH | 2 |
| FXu | Standing waters | SH | 2 |
| FXx | Standing waters | SH | 2 |
| HEn | Scrubs, copses and field hedges | SH | 2 |
| HEy | Scrubs, copses and field hedges | SH | 2 |
| HEx | Scrubs, copses and field hedges | SH | 2 |
| HGm | Scrubs, copses and field hedges | SH | 2 |
| HGn | Scrubs, copses and field hedges | SH | 2 |
| HGp | Scrubs, copses and field hedges | SH | 2 |
| HGs | Scrubs, copses and field hedges | SH | 2 |
| SZf | Anthropogenic | SH | 2 |
| HGx | Scrubs, copses and field hedges | SH | 2 |
| HGy | Scrubs, copses and field hedges | SH | 2 |
| KFb | Coastal and marine habitats | SH | 2 |
| KFy | Coastal and marine habitats | SH | 2 |
| KGf | Coastal and marine habitats | SH | 2 |
| KGg | Coastal and marine habitats | SH | 2 |
| KGy | Coastal and marine habitats | SH | 2 |
| KN | Coastal and marine habitats | SH | 2 |
| KNp | Coastal and marine habitats | SH | 2 |
| KO | Coastal and marine habitats | SH | 2 |
| KQb | Coastal and marine habitats | SH | 2 |
| KQn | Coastal and marine habitats | SH | 2 |
| KBc | Coastal and marine habitats | SH | 2 |
| KBe | Coastal and marine habitats | SH | 2 |
| KBr | Coastal and marine habitats | SH | 2 |
| KSa | Coastal and marine habitats | SH | 2 |
| KSv | Coastal and marine habitats | SH | 2 |
| KSx | Coastal and marine habitats | SH | 2 |
| KTa | Coastal and marine habitats | SH | 2 |
| KW | Coastal and marine habitats | SH | 2 |
| MAt | Bogs, transition mires, marshes and fens | SH | 2 |
| MDy | Bogs, transition mires, marshes and fens | SH | 2 |
| MSg | Standing waters | SH | 2 |
| SDf | Anthropogenic | SH | 2 |
| SEk | Anthropogenic | SH | 2 |
| SEs | Anthropogenic | SH | 2 |
| SGb | Anthropogenic | SH | 2 |
| SGr | Anthropogenic | SH | 2 |
| SIe | Anthropogenic | SH | 2 |
| SKx | Anthropogenic | SH | 2 |
| SMf | Anthropogenic | SH | 2 |
| SVh | Anthropogenic | SH | 2 |
| SVo | Anthropogenic | SH | 2 |
| SFs | Anthropogenic | SH | 2 |
| WTe | Moist to wet forests | SH | 2 |
| WFm | Dry to moderately moist forests | SH | 2 |
| WMo | Dry to moderately moist forests | SH | 2 |
| WPa | Dry to moderately moist forests | SH | 2 |
| WPb | Dry to moderately moist forests | SH | 2 |
| WPm | Dry to moderately moist forests | SH | 2 |
| WPn | Dry to moderately moist forests | SH | 2 |
| WPp | Dry to moderately moist forests | SH | 2 |
| WPs | Dry to moderately moist forests | SH | 2 |
| WPw | Moist to wet forests | SH | 2 |
| WPx | Dry to moderately moist forests | SH | 2 |
| WPy | Dry to moderately moist forests | SH | 2 |
| XAs | Anthropogenic | SH | 2 |
| XAy | Anthropogenic | SH | 2 |
| XD | Coastal and marine habitats | SH | 2 |
| XDl | Coastal and marine habitats | SH | 2 |
| XDs | Coastal and marine habitats | SH | 2 |
| XKd | Coastal and marine habitats | SH | 2 |
| YQx | Fresh water vegetation, springs and reeds | SH | 2 |
| FLa | Linear and running surface waters | SH | 2 |
| FWo | Linear and running surface waters | SH | 2 |
| FWp | Linear and running surface waters | SH | 2 |
| FWy | Linear and running surface waters | SH | 2 |
| FXt | Standing waters | SH | 2 |
| HEo | Scrubs, copses and field hedges | SH | 2 |
| HEw | Scrubs, copses and field hedges | SH | 2 |
| HGe | Scrubs, copses and field hedges | SH | 2 |
| HOn | Mesic grasslands | SH | 2 |
| HOy | Mesic grasslands | SH | 2 |
| KBp | Coastal and marine habitats | SH | 2 |
| SFb | Anthropogenic | SH | 2 |
| SGe | Anthropogenic | SH | 2 |
| WAn | Moist to wet forests | SH | 2 |
| WPe | Moist to wet forests | SH | 2 |
| WTb | Moist to wet forests | SH | 2 |
| WTn | Moist to wet forests | SH | 2 |
| WTp | Moist to wet forests | SH | 2 |
| WTw | Moist to wet forests | SH | 2 |
| WTx | Moist to wet forests | SH | 2 |
| WTy | Moist to wet forests | SH | 2 |
| KWp | Coastal and marine habitats | SH | 2 |
| A | Farmland | SH | 2 |
| FB | Linear and running surface waters | SH | 2 |
| FF | Linear and running surface waters | SH | 2 |
| FG | Linear and running surface waters | SH | 2 |
| FL | Linear and running surface waters | SH | 2 |
| FU | Linear and running surface waters | SH | 2 |
| FXb | Standing waters | SH | 2 |
| FXk | Linear and running surface waters | SH | 2 |
| FXy | Standing waters | SH | 2 |
| FXz | Standing waters | SH | 2 |
| HE | Scrubs, copses and field hedges | SH | 2 |
| HG | Scrubs, copses and field hedges | SH | 2 |
| HO | Mesic grasslands | SH | 2 |
| KB | Coastal and marine habitats | SH | 2 |
| KD | Coastal and marine habitats | SH | 2 |
| KG | Coastal and marine habitats | SH | 2 |
| KH | Coastal and marine habitats | SH | 2 |
| KQ | Coastal and marine habitats | SH | 2 |
| KR | Fresh water vegetation, springs and reeds | SH | 2 |
| KS | Coastal and marine habitats | SH | 2 |
| KT | Coastal and marine habitats | SH | 2 |
| KTy | Coastal and marine habitats | SH | 2 |
| MA | Bogs, transition mires, marshes and fens | SH | 2 |
| MAf | Bogs, transition mires, marshes and fens | SH | 2 |
| MD | Bogs, transition mires, marshes and fens | SH | 2 |
| MS | Bogs, transition mires, marshes and fens | SH | 2 |
| NH | Bogs, transition mires, marshes and fens | SH | 2 |
| S | Anthropogenic | SH | 2 |
| SB | Anthropogenic | SH | 2 |
| SBe | Anthropogenic | SH | 2 |
| SBf | Anthropogenic | SH | 2 |
| SBg | Anthropogenic | SH | 2 |
| SBy | Anthropogenic | SH | 2 |
| SBz | Anthropogenic | SH | 2 |
| SD | Anthropogenic | SH | 2 |
| SDe | Anthropogenic | SH | 2 |
| SDp | Anthropogenic | SH | 2 |
| SDs | Anthropogenic | SH | 2 |
| SDy | Anthropogenic | SH | 2 |
| SE | Anthropogenic | SH | 2 |
| SEb | Anthropogenic | SH | 2 |
| SEc | Anthropogenic | SH | 2 |
| SEd | Anthropogenic | SH | 2 |
| SEf | Anthropogenic | SH | 2 |
| SEg | Anthropogenic | SH | 2 |
| SEr | Anthropogenic | SH | 2 |
| SEw | Anthropogenic | SH | 2 |
| SEy | Anthropogenic | SH | 2 |
| SF | Anthropogenic | SH | 2 |
| SFf | Anthropogenic | SH | 2 |
| SFm | Anthropogenic | SH | 2 |
| SFw | Anthropogenic | SH | 2 |
| SFx | Anthropogenic | SH | 2 |
| SFy | Anthropogenic | SH | 2 |
| SG | Anthropogenic | SH | 2 |
| SGg | Anthropogenic | SH | 2 |
| SGn | Anthropogenic | SH | 2 |
| SGo | Anthropogenic | SH | 2 |
| SGp | Anthropogenic | SH | 2 |
| SGs | Anthropogenic | SH | 2 |
| SGx | Anthropogenic | SH | 2 |
| SGy | Anthropogenic | SH | 2 |
| SGz | Anthropogenic | SH | 2 |
| SI | Anthropogenic | SH | 2 |
| SIa | Anthropogenic | SH | 2 |
| SId | Anthropogenic | SH | 2 |
| SIf | Anthropogenic | SH | 2 |
| SIg | Anthropogenic | SH | 2 |
| SIi | Anthropogenic | SH | 2 |
| SIk | Anthropogenic | SH | 2 |
| SIp | Anthropogenic | SH | 2 |
| SIv | Anthropogenic | SH | 2 |
| SIy | Anthropogenic | SH | 2 |
| SK | Anthropogenic | SH | 2 |
| SKa | Anthropogenic | SH | 2 |
| SKb | Anthropogenic | SH | 2 |
| SKl | Anthropogenic | SH | 2 |
| SKm | Anthropogenic | SH | 2 |
| SKy | Anthropogenic | SH | 2 |
| SL | Anthropogenic | SH | 2 |
| SLg | Anthropogenic | SH | 2 |
| SLr | Anthropogenic | SH | 2 |
| SLt | Anthropogenic | SH | 2 |
| SLy | Anthropogenic | SH | 2 |
| SM | Anthropogenic | SH | 2 |
| SMd | Anthropogenic | SH | 2 |
| SMk | Anthropogenic | SH | 2 |
| SMr | Anthropogenic | SH | 2 |
| SMt | Anthropogenic | SH | 2 |
| SMy | Anthropogenic | SH | 2 |
| SP | Anthropogenic | SH | 2 |
| SPe | Anthropogenic | SH | 2 |
| SPf | Anthropogenic | SH | 2 |
| SPh | Anthropogenic | SH | 2 |
| SPi | Anthropogenic | SH | 2 |
| SPk | Anthropogenic | SH | 2 |
| SPp | Anthropogenic | SH | 2 |
| SPu | Anthropogenic | SH | 2 |
| SPw | Anthropogenic | SH | 2 |
| SPy | Anthropogenic | SH | 2 |
| SPz | Anthropogenic | SH | 2 |
| SV | Anthropogenic | SH | 2 |
| SVb | Anthropogenic | SH | 2 |
| SVe | Anthropogenic | SH | 2 |
| SVg | Anthropogenic | SH | 2 |
| SVi | Anthropogenic | SH | 2 |
| SVS | Anthropogenic | SH | 2 |
| SVT | Anthropogenic | SH | 2 |
| SX | Anthropogenic | SH | 2 |
| SXa | Anthropogenic | SH | 2 |
| SXk | Anthropogenic | SH | 2 |
| SXs | Anthropogenic | SH | 2 |
| SXx | Anthropogenic | SH | 2 |
| SXy | Anthropogenic | SH | 2 |
| SZ | Anthropogenic | SH | 2 |
| SZg | Anthropogenic | SH | 2 |
| SZh | Anthropogenic | SH | 2 |
| SZk | Anthropogenic | SH | 2 |
| SZs | Anthropogenic | SH | 2 |
| SZy | Anthropogenic | SH | 2 |
| TB | Heaths, inland dunes and semi-natural grasslands | SH | 2 |
| WBn | Moist to wet forests | SH | 2 |
| WEn | Moist to wet forests | SH | 2 |
| WF | Dry to moderately moist forests | SH | 2 |
| WP | NA | SH | 2 |
| WQ | Fresh water vegetation, springs and reeds | SH | 2 |
| WT | Moist to wet forests | SH | 2 |
| WTm | Moist to wet forests | SH | 2 |
| XAb | Anthropogenic | SH | 2 |
| XAw | Anthropogenic | SH | 2 |
| KN/KO | Coastal and marine habitats | SH | 2 |
| Wla | Dry to moderately moist forests | SH | 2 |
| Fse | Standing waters | SH | 2 |
| Wfn | Dry to moderately moist forests | SH | 2 |
| Rhn | Ruderal, fringe and tall forb communities, clearings | SH | 2 |
| A | Ruderal, fringe and tall forb communities, clearings | HH |  |
| AK | Ruderal, fringe and tall forb communities, clearings | HH |  |
| AKF | Ruderal, fringe and tall forb communities, clearings | HH |  |
| AKM | Ruderal, fringe and tall forb communities, clearings | HH |  |
| AKN | Ruderal, fringe and tall forb communities, clearings | HH |  |
| AKT | Ruderal, fringe and tall forb communities, clearings | HH |  |
| AP | Ruderal, fringe and tall forb communities, clearings | HH |  |
| APF | Ruderal, fringe and tall forb communities, clearings | HH |  |
| APM | Ruderal, fringe and tall forb communities, clearings | HH |  |
| APT | Ruderal, fringe and tall forb communities, clearings | HH |  |
| B | Anthropogenic | HH |  |
| BB | Anthropogenic | HH |  |
| BBA | Anthropogenic | HH |  |
| BBG | Anthropogenic | HH |  |
| BBN | Anthropogenic | HH |  |
| BBV | Anthropogenic | HH |  |
| BH | Anthropogenic | HH |  |
| BI | Anthropogenic | HH |  |
| BIG | Anthropogenic | HH |  |
| BII | Anthropogenic | HH |  |
| BM | Anthropogenic | HH |  |
| BML | Anthropogenic | HH |  |
| BMP | Anthropogenic | HH |  |
| BMS | Anthropogenic | HH |  |
| BN | Anthropogenic | HH |  |
| BNA | Anthropogenic | HH |  |
| BNE | Anthropogenic | HH |  |
| BNG | Anthropogenic | HH |  |
| BNN | Anthropogenic | HH |  |
| BNO | Anthropogenic | HH |  |
| BNS | Anthropogenic | HH |  |
| BNV | Anthropogenic | HH |  |
| BR | Anthropogenic | HH |  |
| BRG | Anthropogenic | HH |  |
| BRM | Anthropogenic | HH |  |
| BRN | Anthropogenic | HH |  |
| BRS | Anthropogenic | HH |  |
| BS | Anthropogenic | HH |  |
| BSG | Anthropogenic | HH |  |
| BSK | Anthropogenic | HH |  |
| BSS | Anthropogenic | HH |  |
| BSV | Anthropogenic | HH |  |
| BV | Anthropogenic | HH |  |
| BVD | Anthropogenic | HH |  |
| BVK | Anthropogenic | HH |  |
| BVZ | Anthropogenic | HH |  |
| BZ | Anthropogenic | HH |  |
| BZM | Anthropogenic | HH |  |
| BZN | Anthropogenic | HH |  |
| E | Anthropogenic | HH |  |
| EB | Anthropogenic | HH |  |
| EC | Anthropogenic | HH |  |
| EF | Anthropogenic | HH |  |
| EFA | Anthropogenic | HH |  |
| EFP | Anthropogenic | HH |  |
| EFR | Anthropogenic | HH |  |
| EFW | Anthropogenic | HH |  |
| EH | Anthropogenic | HH |  |
| EHB | Anthropogenic | HH |  |
| EHG | Anthropogenic | HH |  |
| EHH | Anthropogenic | HH |  |
| EHN | Anthropogenic | HH |  |
| EHO | Anthropogenic | HH |  |
| EHP | Anthropogenic | HH |  |
| EHZ | Anthropogenic | HH |  |
| EK | Anthropogenic | HH |  |
| EKA | Anthropogenic | HH |  |
| EKG | Anthropogenic | HH |  |
| EKR | Anthropogenic | HH |  |
| EP | Anthropogenic | HH |  |
| EPA | Anthropogenic | HH |  |
| EPB | Anthropogenic | HH |  |
| EPI | Anthropogenic | HH |  |
| EPK | Anthropogenic | HH |  |
| EPL | Anthropogenic | HH |  |
| EPN | Anthropogenic | HH |  |
| EPW | Anthropogenic | HH |  |
| EPZ | Anthropogenic | HH |  |
| ES | Anthropogenic | HH |  |
| ESB | Anthropogenic | HH |  |
| ESG | Anthropogenic | HH |  |
| ESS | Anthropogenic | HH |  |
| ET | Anthropogenic | HH |  |
| EX | Anthropogenic | HH |  |
| F | Linear and running surface waters | HH |  |
| FB | Linear and running surface waters | HH |  |
| FBA | Linear and running surface waters | HH |  |
| FBM | Linear and running surface waters | HH |  |
| FBR | Linear and running surface waters | HH |  |
| FBS | Linear and running surface waters | HH |  |
| FBT | Linear and running surface waters | HH |  |
| FF | Linear and running surface waters | HH |  |
| FFA | Linear and running surface waters | HH |  |
| FFF | Linear and running surface waters | HH |  |
| FFM | Linear and running surface waters | HH |  |
| FFR | Linear and running surface waters | HH |  |
| FFS | Linear and running surface waters | HH |  |
| FFT | Linear and running surface waters | HH |  |
| FG | Linear and running surface waters | HH |  |
| FGA | Linear and running surface waters | HH |  |
| FGM | Linear and running surface waters | HH |  |
| FGR | Linear and running surface waters | HH |  |
| FGV | Linear and running surface waters | HH |  |
| FGX | Linear and running surface waters | HH |  |
| FH | Linear and running surface waters | HH |  |
| FK | Linear and running surface waters | HH |  |
| FL | Linear and running surface waters | HH |  |
| FLA | Linear and running surface waters | HH |  |
| FLH | Linear and running surface waters | HH |  |
| FLM | Linear and running surface waters | HH |  |
| FLR | Linear and running surface waters | HH |  |
| FQ | Linear and running surface waters | HH |  |
| FQB | Linear and running surface waters | HH |  |
| FQG | Linear and running surface waters | HH |  |
| FQS | Linear and running surface waters | HH |  |
| FS | Linear and running surface waters | HH |  |
| FSO | Linear and running surface waters | HH |  |
| FSV | Linear and running surface waters | HH |  |
| FSW | Linear and running surface waters | HH |  |
| FV | Linear and running surface waters | HH |  |
| FVS | Linear and running surface waters | HH |  |
| FVT | Linear and running surface waters | HH |  |
| FVV | Linear and running surface waters | HH |  |
| FVZ | Linear and running surface waters | HH |  |
| FW | Linear and running surface waters | HH |  |
| FWB | Linear and running surface waters | HH |  |
| FWO | Linear and running surface waters | HH |  |
| FWP | Linear and running surface waters | HH |  |
| FWV | Linear and running surface waters | HH |  |
| FWX | Linear and running surface waters | HH |  |
| FWZ | Linear and running surface waters | HH |  |
| FX | Linear and running surface waters | HH |  |
| G | NA | HH |  |
| GF | Moist to wet grasslands | HH |  |
| GFA | Moist to wet grasslands | HH |  |
| GFC | Moist to wet grasslands | HH |  |
| GFF | Moist to wet grasslands | HH |  |
| GFR | Moist to wet grasslands | HH |  |
| GFS | Moist to wet grasslands | HH |  |
| GI | NA | HH |  |
| GIA | Mesic grasslands | HH |  |
| GIF | Moist to wet grasslands | HH |  |
| GIM | Mesic grasslands | HH |  |
| GIS | Mesic grasslands | HH |  |
| GIW | Mesic grasslands | HH |  |
| GM | Mesic grasslands | HH |  |
| GMG | Mesic grasslands | HH |  |
| GMM | Mesic grasslands | HH |  |
| GMT | Mesic grasslands | HH |  |
| GMW | Mesic grasslands | HH |  |
| GMZ | Mesic grasslands | HH |  |
| GN | Moist to wet grasslands | HH |  |
| GNA | Moist to wet grasslands | HH |  |
| GNF | Moist to wet grasslands | HH |  |
| GNK | Moist to wet grasslands | HH |  |
| GNP | Moist to wet grasslands | HH |  |
| GNR | Moist to wet grasslands | HH |  |
| GW | Mesic grasslands | HH |  |
| H | Scrubs, copses and field hedges | HH |  |
| HE | Scrubs, copses and field hedges | HH |  |
| HEA | Scrubs, copses and field hedges | HH |  |
| HEE | Scrubs, copses and field hedges | HH |  |
| HEG | Scrubs, copses and field hedges | HH |  |
| HF | Scrubs, copses and field hedges | HH |  |
| HFS | Scrubs, copses and field hedges | HH |  |
| HFT | Scrubs, copses and field hedges | HH |  |
| HFZ | Scrubs, copses and field hedges | HH |  |
| HG | Scrubs, copses and field hedges | HH |  |
| HGF | Scrubs, copses and field hedges | HH |  |
| HGM | Scrubs, copses and field hedges | HH |  |
| HGT | Scrubs, copses and field hedges | HH |  |
| HGX | Scrubs, copses and field hedges | HH |  |
| HGZ | Scrubs, copses and field hedges | HH |  |
| HH | Scrubs, copses and field hedges | HH |  |
| HHB | Scrubs, copses and field hedges | HH |  |
| HHM | Scrubs, copses and field hedges | HH |  |
| HHN | Scrubs, copses and field hedges | HH |  |
| HHS | Scrubs, copses and field hedges | HH |  |
| HHX | Scrubs, copses and field hedges | HH |  |
| HM | Scrubs, copses and field hedges | HH |  |
| HR | Scrubs, copses and field hedges | HH |  |
| HRR | Scrubs, copses and field hedges | HH |  |
| HRS | Scrubs, copses and field hedges | HH |  |
| HRX | Scrubs, copses and field hedges | HH |  |
| HS | Scrubs, copses and field hedges | HH |  |
| HSC | Scrubs, copses and field hedges | HH |  |
| HSG | Scrubs, copses and field hedges | HH |  |
| HSZ | Scrubs, copses and field hedges | HH |  |
| HT | Scrubs, copses and field hedges | HH |  |
| HTG | Scrubs, copses and field hedges | HH |  |
| HTL | Scrubs, copses and field hedges | HH |  |
| HTT | Scrubs, copses and field hedges | HH |  |
| HTZ | Scrubs, copses and field hedges | HH |  |
| HU | Scrubs, copses and field hedges | HH |  |
| HUE | Scrubs, copses and field hedges | HH |  |
| HUW | Scrubs, copses and field hedges | HH |  |
| HUZ | Scrubs, copses and field hedges | HH |  |
| HW | Scrubs, copses and field hedges | HH |  |
| HWB | Scrubs, copses and field hedges | HH |  |
| HWD | Scrubs, copses and field hedges | HH |  |
| HWM | Scrubs, copses and field hedges | HH |  |
| HWN | Scrubs, copses and field hedges | HH |  |
| HWS | Scrubs, copses and field hedges | HH |  |
| HWX | Scrubs, copses and field hedges | HH |  |
| L | Farmland | HH |  |
| LA | Farmland | HH |  |
| LAL | Farmland | HH |  |
| LAM | Farmland | HH |  |
| LAS | Farmland | HH |  |
| LB | Farmland | HH |  |
| LG | Farmland | HH |  |
| LGG | Farmland | HH |  |
| LGO | Farmland | HH |  |
| LO | Farmland | HH |  |
| LOA | Farmland | HH |  |
| LOB | Farmland | HH |  |
| LOW | Farmland | HH |  |
| LW | Farmland | HH |  |
| LZ | Farmland | HH |  |
| M | Bogs, transition mires, marshes and fens | HH |  |
| MF | Bogs, transition mires, marshes and fens | HH |  |
| MFF | Bogs, transition mires, marshes and fens | HH |  |
| MFT | Bogs, transition mires, marshes and fens | HH |  |
| MH | Bogs, transition mires, marshes and fens | HH |  |
| MHH | Bogs, transition mires, marshes and fens | HH |  |
| MHR | Bogs, transition mires, marshes and fens | HH |  |
| MM | Bogs, transition mires, marshes and fens | HH |  |
| MMF | Bogs, transition mires, marshes and fens | HH |  |
| MMT | Bogs, transition mires, marshes and fens | HH |  |
| MR | Bogs, transition mires, marshes and fens | HH |  |
| MRR | Bogs, transition mires, marshes and fens | HH |  |
| MRS | Bogs, transition mires, marshes and fens | HH |  |
| MRW | Bogs, transition mires, marshes and fens | HH |  |
| MX | Bogs, transition mires, marshes and fens | HH |  |
| MXA | Bogs, transition mires, marshes and fens | HH |  |
| MXR | Bogs, transition mires, marshes and fens | HH |  |
| N | Bogs, transition mires, marshes and fens | HH |  |
| NA | Bogs, transition mires, marshes and fens | HH |  |
| NAA | Bogs, transition mires, marshes and fens | HH |  |
| NAK | Bogs, transition mires, marshes and fens | HH |  |
| NG | Bogs, transition mires, marshes and fens | HH |  |
| NGB | Bogs, transition mires, marshes and fens | HH |  |
| NGG | Bogs, transition mires, marshes and fens | HH |  |
| NGZ | Bogs, transition mires, marshes and fens | HH |  |
| NH | Bogs, transition mires, marshes and fens | HH |  |
| NHA | Bogs, transition mires, marshes and fens | HH |  |
| NHR | Bogs, transition mires, marshes and fens | HH |  |
| NP | Bogs, transition mires, marshes and fens | HH |  |
| NPA | Bogs, transition mires, marshes and fens | HH |  |
| NPR | Bogs, transition mires, marshes and fens | HH |  |
| NPT | Bogs, transition mires, marshes and fens | HH |  |
| NPZ | Bogs, transition mires, marshes and fens | HH |  |
| NR | Bogs, transition mires, marshes and fens | HH |  |
| NRG | Bogs, transition mires, marshes and fens | HH |  |
| NRR | Bogs, transition mires, marshes and fens | HH |  |
| NRS | Bogs, transition mires, marshes and fens | HH |  |
| NRT | Bogs, transition mires, marshes and fens | HH |  |
| NRW | Bogs, transition mires, marshes and fens | HH |  |
| NRZ | Bogs, transition mires, marshes and fens | HH |  |
| NU | Bogs, transition mires, marshes and fens | HH |  |
| NUB | Bogs, transition mires, marshes and fens | HH |  |
| NUE | Bogs, transition mires, marshes and fens | HH |  |
| NUG | Bogs, transition mires, marshes and fens | HH |  |
| NUW | Bogs, transition mires, marshes and fens | HH |  |
| NUZ | Bogs, transition mires, marshes and fens | HH |  |
| O | Anthropogenic | HH |  |
| OA | Anthropogenic | HH |  |
| OAG | Anthropogenic | HH |  |
| OAS | Anthropogenic | HH |  |
| OAT | Anthropogenic | HH |  |
| OAX | Anthropogenic | HH |  |
| OB | Anthropogenic | HH |  |
| OBK | Anthropogenic | HH |  |
| OBT | Anthropogenic | HH |  |
| OBX | Anthropogenic | HH |  |
| OK | Anthropogenic | HH |  |
| OKL | Anthropogenic | HH |  |
| OKS | Anthropogenic | HH |  |
| OW | Anthropogenic | HH |  |
| OWL | Anthropogenic | HH |  |
| OWS | Anthropogenic | HH |  |
| OWX | Anthropogenic | HH |  |
| OX | Anthropogenic | HH |  |
| S | Standing waters | HH |  |
| SE | Standing waters | HH |  |
| SEA | Standing waters | HH |  |
| SEB | Standing waters | HH |  |
| SED | Standing waters | HH |  |
| SEF | Standing waters | HH |  |
| SEG | Standing waters | HH |  |
| SEN | Standing waters | HH |  |
| SEO | Standing waters | HH |  |
| SEP | Standing waters | HH |  |
| SER | Standing waters | HH |  |
| SES | Standing waters | HH |  |
| SET | Standing waters | HH |  |
| SEW | Standing waters | HH |  |
| SEY | Standing waters | HH |  |
| SEZ | Standing waters | HH |  |
| SG | Standing waters | HH |  |
| SGA | Standing waters | HH |  |
| SGF | Standing waters | HH |  |
| SGN | Standing waters | HH |  |
| SGT | Standing waters | HH |  |
| SGZ | Standing waters | HH |  |
| SO | Standing waters | HH |  |
| SOA | Standing waters | HH |  |
| SOD | Standing waters | HH |  |
| SOG | Standing waters | HH |  |
| SOM | Standing waters | HH |  |
| SON | Standing waters | HH |  |
| SOT | Standing waters | HH |  |
| SOW | Standing waters | HH |  |
| SOZ | Standing waters | HH |  |
| ST | Standing waters | HH |  |
| STA | Standing waters | HH |  |
| STG | Standing waters | HH |  |
| STQ | Standing waters | HH |  |
| STR | Standing waters | HH |  |
| STW | Standing waters | HH |  |
| STZ | Standing waters | HH |  |
| SV | Standing waters | HH |  |
| SVS | Standing waters | HH |  |
| SVT | Standing waters | HH |  |
| SX | Standing waters | HH |  |
| SXA | Standing waters | HH |  |
| SXB | Standing waters | HH |  |
| SXG | Standing waters | HH |  |
| SXK | Standing waters | HH |  |
| SXL | Standing waters | HH |  |
| SXN | Standing waters | HH |  |
| SXP | Standing waters | HH |  |
| SXR | Standing waters | HH |  |
| SXT | Standing waters | HH |  |
| SXY | Standing waters | HH |  |
| SXZ | Standing waters | HH |  |
| T | Heaths, inland dunes and semi-natural grasslands | HH |  |
| TC | Heaths, inland dunes and semi-natural grasslands | HH |  |
| TCF | Heaths, inland dunes and semi-natural grasslands | HH |  |
| TCT | Heaths, inland dunes and semi-natural grasslands | HH |  |
| TD | Heaths, inland dunes and semi-natural grasslands | HH |  |
| TDC | Heaths, inland dunes and semi-natural grasslands | HH |  |
| TDO | Heaths, inland dunes and semi-natural grasslands | HH |  |
| TDS | Heaths, inland dunes and semi-natural grasslands | HH |  |
| TDZ | Heaths, inland dunes and semi-natural grasslands | HH |  |
| TM | Heaths, inland dunes and semi-natural grasslands | HH |  |
| TMA | Heaths, inland dunes and semi-natural grasslands | HH |  |
| TMB | Heaths, inland dunes and semi-natural grasslands | HH |  |
| TMK | Heaths, inland dunes and semi-natural grasslands | HH |  |
| TMS | Heaths, inland dunes and semi-natural grasslands | HH |  |
| TMZ | Heaths, inland dunes and semi-natural grasslands | HH |  |
| TN | Heaths, inland dunes and semi-natural grasslands | HH |  |
| TNF | Heaths, inland dunes and semi-natural grasslands | HH |  |
| TNT | Heaths, inland dunes and semi-natural grasslands | HH |  |
| V | Anthropogenic | HH |  |
| VB | Anthropogenic | HH |  |
| VBB | Anthropogenic | HH |  |
| VBD | Anthropogenic | HH |  |
| VBG | Anthropogenic | HH |  |
| VK | Anthropogenic | HH |  |
| VKH | Anthropogenic | HH |  |
| VKS | Anthropogenic | HH |  |
| VL | Anthropogenic | HH |  |
| VLF | Anthropogenic | HH |  |
| VLH | Anthropogenic | HH |  |
| VLS | Anthropogenic | HH |  |
| VLZ | Anthropogenic | HH |  |
| VS | Anthropogenic | HH |  |
| VSA | Anthropogenic | HH |  |
| VSF | Anthropogenic | HH |  |
| VSL | Anthropogenic | HH |  |
| VSP | Anthropogenic | HH |  |
| VSR | Anthropogenic | HH |  |
| VSS | Anthropogenic | HH |  |
| VSW | Anthropogenic | HH |  |
| VSZ | Anthropogenic | HH |  |
| W | NA | HH |  |
| WB | Moist to wet forests | HH |  |
| WBB | Moist to wet forests | HH |  |
| WBE | Moist to wet forests | HH |  |
| WBX | Moist to wet forests | HH |  |
| WBY | Moist to wet forests | HH |  |
| WC | NA | HH |  |
| WCF | Moist to wet forests | HH |  |
| WCM | Dry to moderately moist forests | HH |  |
| WE | Moist to wet forests | HH |  |
| WEA | Moist to wet forests | HH |  |
| WEQ | Moist to wet forests | HH |  |
| WEZ | Moist to wet forests | HH |  |
| WH | Moist to wet forests | HH |  |
| WHA | Moist to wet forests | HH |  |
| WHB | Moist to wet forests | HH |  |
| WI | Dry to moderately moist forests | HH |  |
| WJ | Dry to moderately moist forests | HH |  |
| WJL | Dry to moderately moist forests | HH |  |
| WJN | Dry to moderately moist forests | HH |  |
| WM | Dry to moderately moist forests | HH |  |
| WMM | Dry to moderately moist forests | HH |  |
| WMS | Dry to moderately moist forests | HH |  |
| WN | NA | HH |  |
| WNF | Dry to moderately moist forests | HH |  |
| WNK | Dry to moderately moist forests | HH |  |
| WNN | Moist to wet forests | HH |  |
| WNZ | Dry to moderately moist forests | HH |  |
| WP | NA | HH |  |
| WPA | Dry to moderately moist forests | HH |  |
| WPB | Dry to moderately moist forests | HH |  |
| WPW | Moist to wet forests | HH |  |
| WPZ | Dry to moderately moist forests | HH |  |
| WQ | NA | HH |  |
| WQF | Moist to wet forests | HH |  |
| WQM | Dry to moderately moist forests | HH |  |
| WQT | Dry to moderately moist forests | HH |  |
| WQZ | Dry to moderately moist forests | HH |  |
| WR | NA | HH |  |
| WS | Moist to wet forests | HH |  |
| WSE | Moist to wet forests | HH |  |
| WSW | Moist to wet forests | HH |  |
| WSZ | Moist to wet forests | HH |  |
| WW | Moist to wet forests | HH |  |
| WWA | Moist to wet forests | HH |  |
| WWT | Moist to wet forests | HH |  |
| WWZ | Moist to wet forests | HH |  |
| WX | Dry to moderately moist forests | HH |  |
| WXE | Dry to moderately moist forests | HH |  |
| WXH | Dry to moderately moist forests | HH |  |
| WXP | Dry to moderately moist forests | HH |  |
| WXR | Dry to moderately moist forests | HH |  |
| WXZ | Dry to moderately moist forests | HH |  |
| WY | Dry to moderately moist forests | HH |  |
| WZ | Dry to moderately moist forests | HH |  |
| WZD | Dry to moderately moist forests | HH |  |
| WZF | Dry to moderately moist forests | HH |  |
| WZK | Dry to moderately moist forests | HH |  |
| WZL | Dry to moderately moist forests | HH |  |
| WZZ | Dry to moderately moist forests | HH |  |
| Y | Anthropogenic | HH |  |
| YD | Anthropogenic | HH |  |
| YDG | Anthropogenic | HH |  |
| YDK | Anthropogenic | HH |  |
| YDR | Anthropogenic | HH |  |
| YDX | Anthropogenic | HH |  |
| YDZ | Anthropogenic | HH |  |
| YF | Anthropogenic | HH |  |
| YFB | Anthropogenic | HH |  |
| YFK | Anthropogenic | HH |  |
| YFP | Anthropogenic | HH |  |
| YFR | Anthropogenic | HH |  |
| YFS | Anthropogenic | HH |  |
| YFV | Anthropogenic | HH |  |
| YFW | Anthropogenic | HH |  |
| YFZ | Anthropogenic | HH |  |
| YM | Anthropogenic | HH |  |
| YMF | Anthropogenic | HH |  |
| YMH | Anthropogenic | HH |  |
| YMN | Anthropogenic | HH |  |
| YMW | Anthropogenic | HH |  |
| YMX | Anthropogenic | HH |  |
| YMZ | Anthropogenic | HH |  |
| Z | Anthropogenic | HH |  |
| ZH | Anthropogenic | HH |  |
| ZHF | Anthropogenic | HH |  |
| ZHN | Anthropogenic | HH |  |
| ZN | Anthropogenic | HH |  |
| ZR | Anthropogenic | HH |  |
| ZRE | Anthropogenic | HH |  |
| ZRT | Anthropogenic | HH |  |
| ZRW | Anthropogenic | HH |  |
| ZS | Anthropogenic | HH |  |
| ZSF | Anthropogenic | HH |  |
| ZSH | Anthropogenic | HH |  |
| ZSN | Anthropogenic | HH |  |
| ZSR | Anthropogenic | HH |  |
| ZSS | Anthropogenic | HH |  |
| ZZ | Anthropogenic | HH |  |
| 3100 | Bogs, transition mires, marshes and fens | BW |  |
| 3110 | Bogs, transition mires, marshes and fens | BW |  |
| 3111 | Bogs, transition mires, marshes and fens | BW |  |
| 3112 | Bogs, transition mires, marshes and fens | BW |  |
| 3120 | Bogs, transition mires, marshes and fens | BW |  |
| 3130 | Bogs, transition mires, marshes and fens | BW |  |
| 3131 | Bogs, transition mires, marshes and fens | BW |  |
| 3132 | Bogs, transition mires, marshes and fens | BW |  |
| 3200 | Bogs, transition mires, marshes and fens | BW |  |
| 3210 | Bogs, transition mires, marshes and fens | BW |  |
| 3211 | Bogs, transition mires, marshes and fens | BW |  |
| 3212 | Bogs, transition mires, marshes and fens | BW |  |
| 3220 | Bogs, transition mires, marshes and fens | BW |  |
| 3221 | Bogs, transition mires, marshes and fens | BW |  |
| 3222 | Bogs, transition mires, marshes and fens | BW |  |
| 3230 | Bogs, transition mires, marshes and fens | BW |  |
| 3231 | Bogs, transition mires, marshes and fens | BW |  |
| 3232 | Bogs, transition mires, marshes and fens | BW |  |
| 3233 | Bogs, transition mires, marshes and fens | BW |  |
| 3300 | NA | BW |  |
| 3310 | Moist to wet grasslands | BW |  |
| 3320 | Moist to wet grasslands | BW |  |
| 3321 | Moist to wet grasslands | BW |  |
| 3322 | Moist to wet grasslands | BW |  |
| 3323 | Moist to wet grasslands | BW |  |
| 3324 | Moist to wet grasslands | BW |  |
| 3330 | Moist to wet grasslands | BW |  |
| 3340 | Mesic grasslands | BW |  |
| 3341 | Mesic grasslands | BW |  |
| 3343 | Mesic grasslands | BW |  |
| 3344 | Mesic grasslands | BW |  |
| 3350 | Mesic grasslands | BW |  |
| 3351 | Mesic grasslands | BW |  |
| 3352 | Mesic grasslands | BW |  |
| 3360 | Mesic grasslands | BW |  |
| 3361 | Mesic grasslands | BW |  |
| 3362 | Mesic grasslands | BW |  |
| 3363 | Mesic grasslands | BW |  |
| 3370 | Mesic grasslands | BW |  |
| 3371 | Mesic grasslands | BW |  |
| 3372 | Mesic grasslands | BW |  |
| 3380 | Mesic grasslands | BW |  |
| 3400 | Fresh water vegetation, springs and reeds | BW |  |
| 3410 | Fresh water vegetation, springs and reeds | BW |  |
| 3411 | Fresh water vegetation, springs and reeds | BW |  |
| 3412 | Fresh water vegetation, springs and reeds | BW |  |
| 3420 | Fresh water vegetation, springs and reeds | BW |  |
| 3421 | Fresh water vegetation, springs and reeds | BW |  |
| 3422 | Fresh water vegetation, springs and reeds | BW |  |
| 3430 | Fresh water vegetation, springs and reeds | BW |  |
| 3431 | Fresh water vegetation, springs and reeds | BW |  |
| 3432 | Fresh water vegetation, springs and reeds | BW |  |
| 3440 | Fresh water vegetation, springs and reeds | BW |  |
| 3450 | Fresh water vegetation, springs and reeds | BW |  |
| 3451 | Fresh water vegetation, springs and reeds | BW |  |
| 3452 | Fresh water vegetation, springs and reeds | BW |  |
| 3453 | Fresh water vegetation, springs and reeds | BW |  |
| 3454 | Fresh water vegetation, springs and reeds | BW |  |
| 3455 | Fresh water vegetation, springs and reeds | BW |  |
| 3456 | Fresh water vegetation, springs and reeds | BW |  |
| 3457 | Fresh water vegetation, springs and reeds | BW |  |
| 3458 | Fresh water vegetation, springs and reeds | BW |  |
| 3459 | Fresh water vegetation, springs and reeds | BW |  |
| 3460 | Fresh water vegetation, springs and reeds | BW |  |
| 3461 | Fresh water vegetation, springs and reeds | BW |  |
| 3462 | Fresh water vegetation, springs and reeds | BW |  |
| 3463 | Fresh water vegetation, springs and reeds | BW |  |
| 3464 | Fresh water vegetation, springs and reeds | BW |  |
| 3465 | Fresh water vegetation, springs and reeds | BW |  |
| 3466 | Fresh water vegetation, springs and reeds | BW |  |
| 3467 | Fresh water vegetation, springs and reeds | BW |  |
| 3468 | Fresh water vegetation, springs and reeds | BW |  |
| 3469 | Fresh water vegetation, springs and reeds | BW |  |
| 3500 | Ruderal, fringe and tall forb communities, clearings | BW |  |
| 3510 | Ruderal, fringe and tall forb communities, clearings | BW |  |
| 3511 | Ruderal, fringe and tall forb communities, clearings | BW |  |
| 3512 | Ruderal, fringe and tall forb communities, clearings | BW |  |
| 3520 | Ruderal, fringe and tall forb communities, clearings | BW |  |
| 3530 | Ruderal, fringe and tall forb communities, clearings | BW |  |
| 3531 | Ruderal, fringe and tall forb communities, clearings | BW |  |
| 3532 | Ruderal, fringe and tall forb communities, clearings | BW |  |
| 3533 | Ruderal, fringe and tall forb communities, clearings | BW |  |
| 3534 | Ruderal, fringe and tall forb communities, clearings | BW |  |
| 3535 | Ruderal, fringe and tall forb communities, clearings | BW |  |
| 3536 | Ruderal, fringe and tall forb communities, clearings | BW |  |
| 3537 | Ruderal, fringe and tall forb communities, clearings | BW |  |
| 3538 | Ruderal, fringe and tall forb communities, clearings | BW |  |
| 3539 | Ruderal, fringe and tall forb communities, clearings | BW |  |
| 3540 | Ruderal, fringe and tall forb communities, clearings | BW |  |
| 3541 | Ruderal, fringe and tall forb communities, clearings | BW |  |
| 3542 | Ruderal, fringe and tall forb communities, clearings | BW |  |
| 3543 | Ruderal, fringe and tall forb communities, clearings | BW |  |
| 3544 | Ruderal, fringe and tall forb communities, clearings | BW |  |
| 3550 | Ruderal, fringe and tall forb communities, clearings | BW |  |
| 3560 | Ruderal, fringe and tall forb communities, clearings | BW |  |
| 3561 | Ruderal, fringe and tall forb communities, clearings | BW |  |
| 3562 | Ruderal, fringe and tall forb communities, clearings | BW |  |
| 3563 | Ruderal, fringe and tall forb communities, clearings | BW |  |
| 3564 | Ruderal, fringe and tall forb communities, clearings | BW |  |
| 3565 | Ruderal, fringe and tall forb communities, clearings | BW |  |
| 3600 | Heaths, inland dunes and semi-natural grasslands | BW |  |
| 3610 | Heaths, inland dunes and semi-natural grasslands | BW |  |
| 3620 | Heaths, inland dunes and semi-natural grasslands | BW |  |
| 3630 | Heaths, inland dunes and semi-natural grasslands | BW |  |
| 3640 | Heaths, inland dunes and semi-natural grasslands | BW |  |
| 3641 | Heaths, inland dunes and semi-natural grasslands | BW |  |
| 3642 | Heaths, inland dunes and semi-natural grasslands | BW |  |
| 3643 | Heaths, inland dunes and semi-natural grasslands | BW |  |
| 3644 | Heaths, inland dunes and semi-natural grasslands | BW |  |
| 3645 | Heaths, inland dunes and semi-natural grasslands | BW |  |
| 3650 | Heaths, inland dunes and semi-natural grasslands | BW |  |
| 3660 | Heaths, inland dunes and semi-natural grasslands | BW |  |
| 3661 | Heaths, inland dunes and semi-natural grasslands | BW |  |
| 3662 | Heaths, inland dunes and semi-natural grasslands | BW |  |
| 3670 | Heaths, inland dunes and semi-natural grasslands | BW |  |
| 4100 | Scrubs, copses and field hedges | BW |  |
| 4110 | Scrubs, copses and field hedges | BW |  |
| 4120 | Scrubs, copses and field hedges | BW |  |
| 4121 | Scrubs, copses and field hedges | BW |  |
| 4122 | Scrubs, copses and field hedges | BW |  |
| 4123 | Scrubs, copses and field hedges | BW |  |
| 4124 | Scrubs, copses and field hedges | BW |  |
| 4125 | Scrubs, copses and field hedges | BW |  |
| 4126 | Scrubs, copses and field hedges | BW |  |
| 4200 | Scrubs, copses and field hedges | BW |  |
| 4210 | Scrubs, copses and field hedges | BW |  |
| 4211 | Scrubs, copses and field hedges | BW |  |
| 4212 | Scrubs, copses and field hedges | BW |  |
| 4213 | Scrubs, copses and field hedges | BW |  |
| 4214 | Scrubs, copses and field hedges | BW |  |
| 4220 | Scrubs, copses and field hedges | BW |  |
| 4221 | Scrubs, copses and field hedges | BW |  |
| 4222 | Scrubs, copses and field hedges | BW |  |
| 4223 | Scrubs, copses and field hedges | BW |  |
| 4224 | Scrubs, copses and field hedges | BW |  |
| 4230 | Scrubs, copses and field hedges | BW |  |
| 4231 | Scrubs, copses and field hedges | BW |  |
| 4232 | Scrubs, copses and field hedges | BW |  |
| 4240 | Scrubs, copses and field hedges | BW |  |
| 4250 | Scrubs, copses and field hedges | BW |  |
| 4251 | Scrubs, copses and field hedges | BW |  |
| 4252 | Scrubs, copses and field hedges | BW |  |
| 4300 | Scrubs, copses and field hedges | BW |  |
| 4310 | Scrubs, copses and field hedges | BW |  |
| 4311 | Scrubs, copses and field hedges | BW |  |
| 4312 | Scrubs, copses and field hedges | BW |  |
| 4313 | Scrubs, copses and field hedges | BW |  |
| 4314 | Scrubs, copses and field hedges | BW |  |
| 4330 | Scrubs, copses and field hedges | BW |  |
| 4350 | Scrubs, copses and field hedges | BW |  |
| 4351 | Scrubs, copses and field hedges | BW |  |
| 4352 | Scrubs, copses and field hedges | BW |  |
| 4353 | Scrubs, copses and field hedges | BW |  |
| 4354 | Scrubs, copses and field hedges | BW |  |
| 4400 | Scrubs, copses and field hedges | BW |  |
| 4410 | Scrubs, copses and field hedges | BW |  |
| 4411 | Scrubs, copses and field hedges | BW |  |
| 4412 | Scrubs, copses and field hedges | BW |  |
| 4420 | Scrubs, copses and field hedges | BW |  |
| 4421 | Scrubs, copses and field hedges | BW |  |
| 4422 | Scrubs, copses and field hedges | BW |  |
| 4430 | Scrubs, copses and field hedges | BW |  |
| 4500 | Scrubs, copses and field hedges | BW |  |
| 4510 | Scrubs, copses and field hedges | BW |  |
| 4511 | Scrubs, copses and field hedges | BW |  |
| 4512 | Scrubs, copses and field hedges | BW |  |
| 4520 | Scrubs, copses and field hedges | BW |  |
| 4540 | Scrubs, copses and field hedges | BW |  |
| 5100 | Moist to wet forests | BW |  |
| 5110 | Moist to wet forests | BW |  |
| 5111 | Moist to wet forests | BW |  |
| 5112 | Moist to wet forests | BW |  |
| 5120 | Moist to wet forests | BW |  |
| 5200 | Moist to wet forests | BW |  |
| 5210 | Moist to wet forests | BW |  |
| 5211 | Moist to wet forests | BW |  |
| 5212 | Moist to wet forests | BW |  |
| 5220 | Moist to wet forests | BW |  |
| 5221 | Moist to wet forests | BW |  |
| 5222 | Moist to wet forests | BW |  |
| 5223 | Moist to wet forests | BW |  |
| 5230 | Moist to wet forests | BW |  |
| 5231 | Moist to wet forests | BW |  |
| 5232 | Moist to wet forests | BW |  |
| 5233 | Moist to wet forests | BW |  |
| 5234 | Moist to wet forests | BW |  |
| 5240 | Moist to wet forests | BW |  |
| 5250 | Moist to wet forests | BW |  |
| 5300 | Dry to moderately moist forests | BW |  |
| 5310 | Dry to moderately moist forests | BW |  |
| 5311 | Dry to moderately moist forests | BW |  |
| 5312 | Dry to moderately moist forests | BW |  |
| 5313 | Dry to moderately moist forests | BW |  |
| 5320 | Dry to moderately moist forests | BW |  |
| 5321 | Dry to moderately moist forests | BW |  |
| 5322 | Dry to moderately moist forests | BW |  |
| 5330 | Dry to moderately moist forests | BW |  |
| 5340 | Dry to moderately moist forests | BW |  |
| 5341 | Dry to moderately moist forests | BW |  |
| 5342 | Dry to moderately moist forests | BW |  |
| 5343 | Dry to moderately moist forests | BW |  |
| 5400 | Dry to moderately moist forests | BW |  |
| 5410 | Dry to moderately moist forests | BW |  |
| 5411 | Dry to moderately moist forests | BW |  |
| 5413 | Dry to moderately moist forests | BW |  |
| 5414 | Dry to moderately moist forests | BW |  |
| 5420 | Dry to moderately moist forests | BW |  |
| 5421 | Dry to moderately moist forests | BW |  |
| 5422 | Dry to moderately moist forests | BW |  |
| 5430 | Dry to moderately moist forests | BW |  |
| 5440 | Dry to moderately moist forests | BW |  |
| 5500 | Dry to moderately moist forests | BW |  |
| 5510 | Dry to moderately moist forests | BW |  |
| 5512 | Dry to moderately moist forests | BW |  |
| 5520 | Dry to moderately moist forests | BW |  |
| 5521 | Dry to moderately moist forests | BW |  |
| 5522 | Dry to moderately moist forests | BW |  |
| 5540 | Dry to moderately moist forests | BW |  |
| 5550 | Dry to moderately moist forests | BW |  |
| 5600 | Dry to moderately moist forests | BW |  |
| 5610 | Dry to moderately moist forests | BW |  |
| 5611 | Dry to moderately moist forests | BW |  |
| 5612 | Dry to moderately moist forests | BW |  |
| 5620 | Dry to moderately moist forests | BW |  |
| 5630 | Dry to moderately moist forests | BW |  |
| 5700 | NA | BW |  |
| 5720 | Moist to wet forests | BW |  |
| 5730 | NA | BW |  |
| 5731 | NA | BW |  |
| 5732 | NA | BW |  |
| 5733 | NA | BW |  |
| 5734 | NA | BW |  |
| 5735 | NA | BW |  |
| 5800 | NA | BW |  |
| 5810 | NA | BW |  |
| 5811 | NA | BW |  |
| 5812 | NA | BW |  |
| 5813 | NA | BW |  |
| 5820 | NA | BW |  |
| 5821 | NA | BW |  |
| 5822 | NA | BW |  |
| 5840 | NA | BW |  |
| 5841 | NA | BW |  |
| 5842 | NA | BW |  |
| 5843 | NA | BW |  |
| 6500 | Mesic grasslands | BW |  |
| 6510 | Mesic grasslands | BW |  |
| 6520 | Mesic grasslands | BW |  |

Notes: The column “Code” indicates the state’s specific habitat type code. Groups correspond to the groups listed in table S1.2. NAs in this column indicate that the habitat type was not used for analysis, mainly because there were no species lists available. The column “Time” for the state SH indicates if this habitat type category was mapped during the first (1) or second (2) time interval.

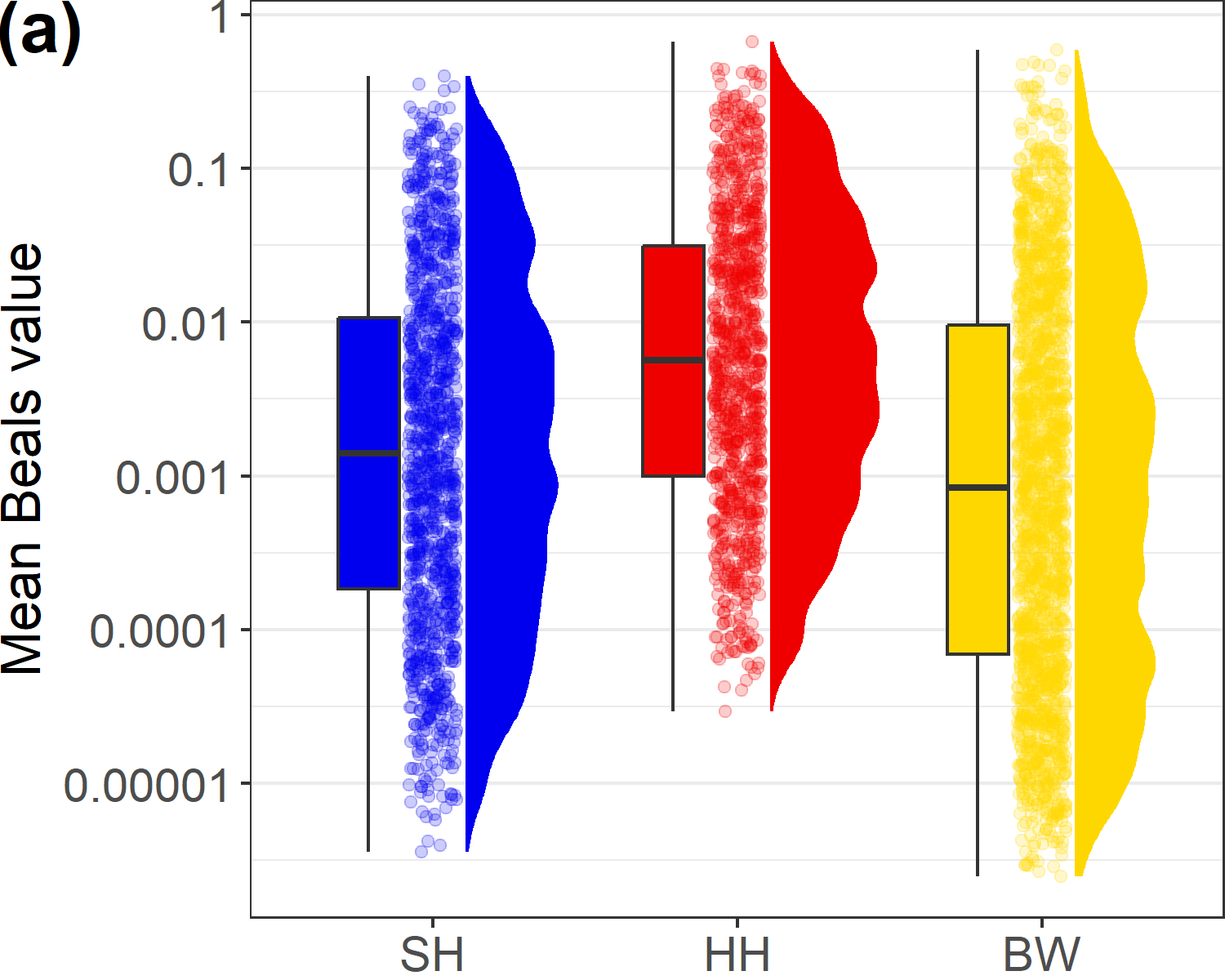

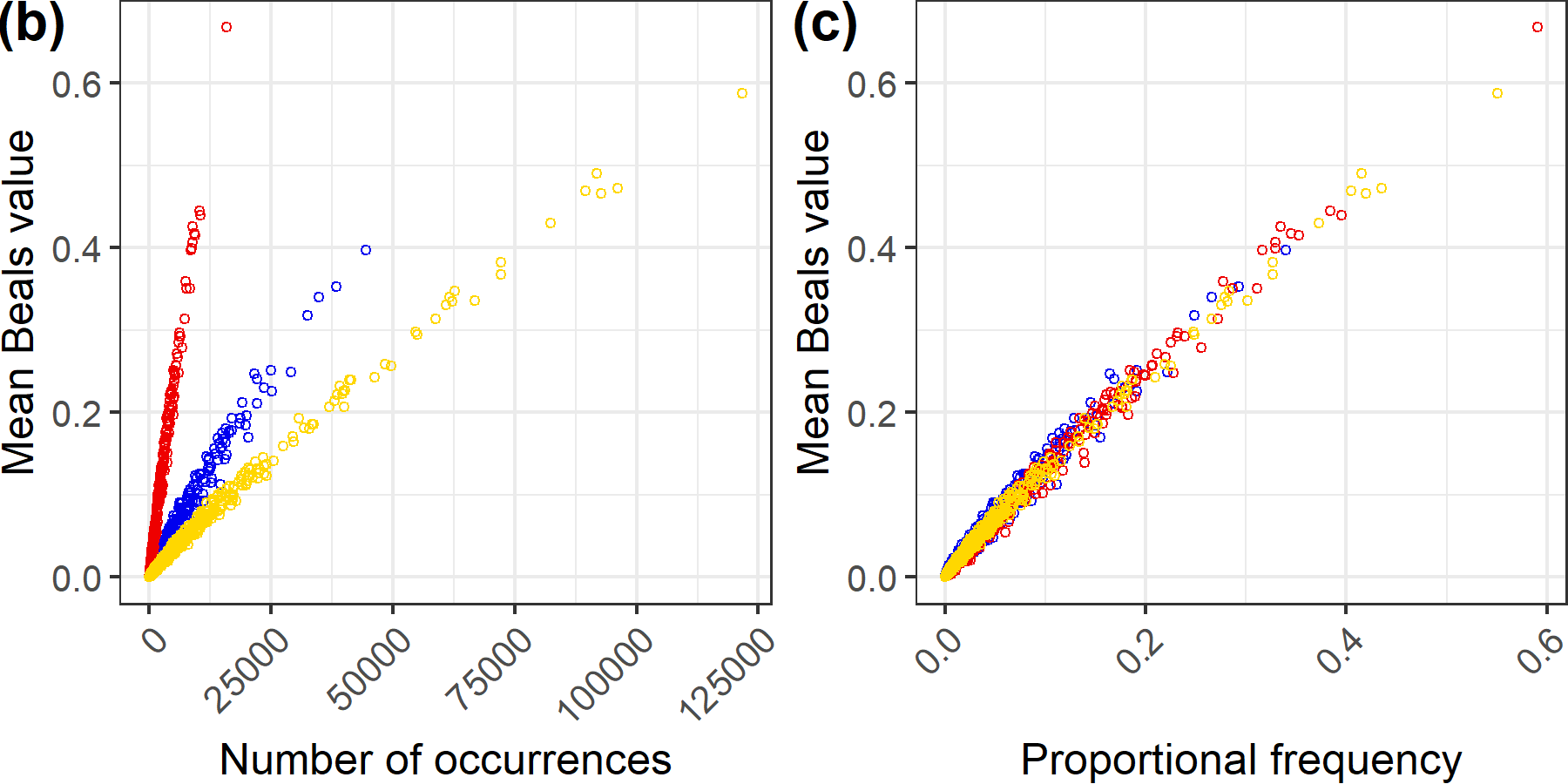

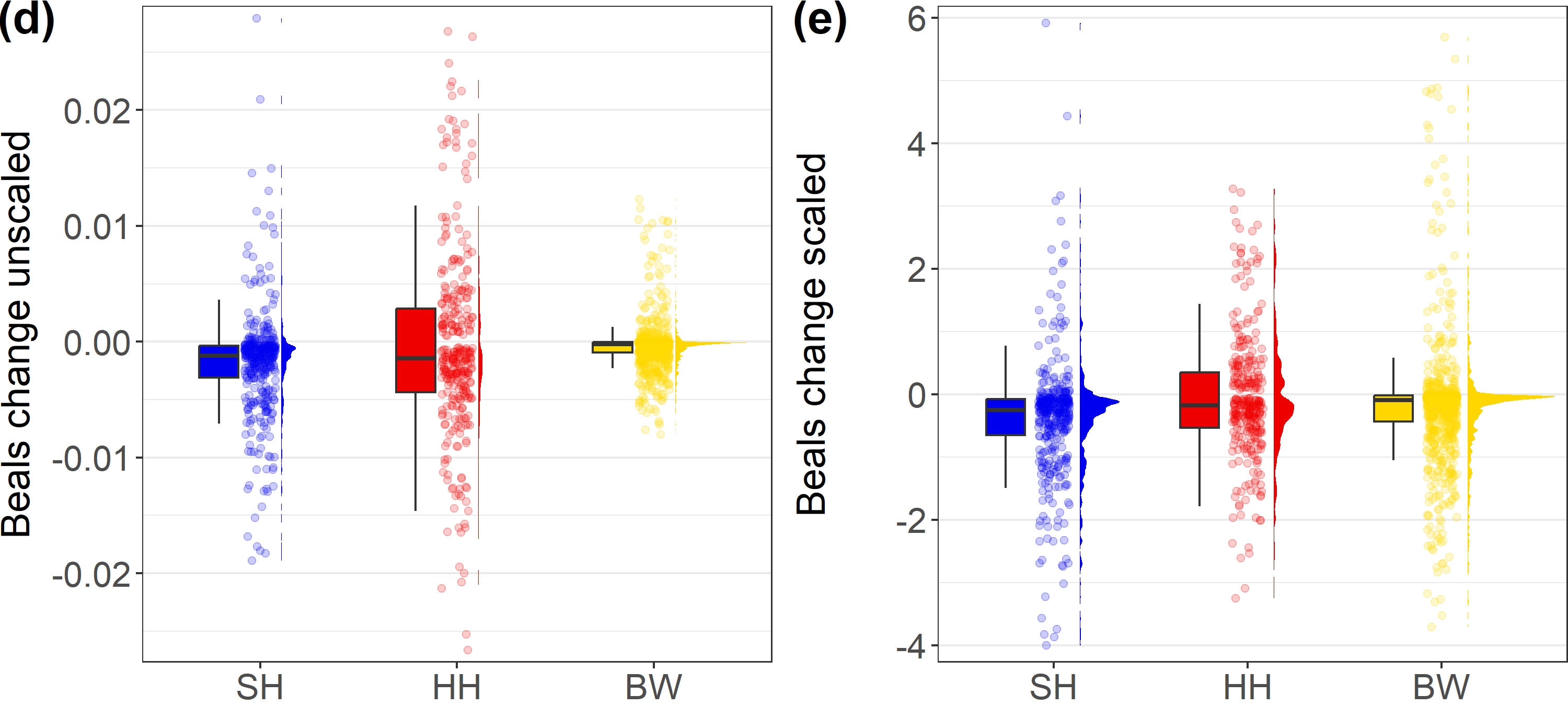

**Figure S1.4** Mean Beals and Beals change values for each species across each state. **(a)** Species mean Beals values, **(b)** species mean Beals values in relation to the number of the species’ occurrences in a state (polygons), **(c)** species mean Beals values in relation to the proportional frequency of each species in a state, **(d)** species mean Beals change values, and **(e)** standardized species mean Beals change values (mean Beals change/sd(mean Beals changes)). Dots for SH in blue, for HH in red and for BW in yellow for all plots.

**Table S1.4** Beals trends of all species that showed a significant trend in all states (105).

| **Species** | **n (SH)** | **Beals trend (SH)** | **t (SH)** | **p (SH)** | **Beals trend scaled (SH)** | **n (HH)** | **Beals trend (HH)** | **t (HH)** | **p (HH)** | **Beals trend scaled (HH)** | **n (BW)** | **Beals trend (BW)** | **t (BW)** | **p (BW)** | **Beals trend scaled (BW)** | **Habitat type preferred** | **Mean Beals trend** |
| --- | --- | --- | --- | --- | --- | --- | --- | --- | --- | --- | --- | --- | --- | --- | --- | --- | --- |
| *Festuca rubra* agg. | 2066 | -0.0189 | -31.73 | <.001 | -4.00 | 405 | -0.0165 | -11.83 | <.001 | -2.01 | 3200 | -0.0028 | -31.04 | <.001 | -1.28 | Coastal and marine habitats | -2.43 |
| *Lotus pedunculatus* | 1538 | -0.0109 | -32.48 | <.001 | -2.30 | 254 | -0.0087 | -8.92 | <.001 | -1.07 | 3692 | -0.006 | -43.22 | <.001 | -2.79 | Moist to wet grasslands | -2.05 |
| *Calluna vulgaris* | 1011 | -0.0183 | -46.43 | <.001 | -3.87 | 110 | -0.0071 | -7.31 | <.001 | -0.87 | 1834 | -0.0026 | -29.45 | <.001 | -1.21 | Bogs, transition mires, marshes and fens | -1.98 |
| *Anthoxanthum odoratum* agg. | 1608 | -0.0143 | -32.57 | <.001 | -3.02 | 192 | -0.009 | -11.10 | <.001 | -1.10 | 2466 | -0.0038 | -44.67 | <.001 | -1.77 | Moist to wet grasslands | -1.96 |
| *Cirsium palustre* | 2401 | -0.0124 | -34.91 | <.001 | -2.62 | 280 | -0.0057 | -6.17 | <.001 | -0.70 | 3916 | -0.0052 | -39.46 | <.001 | -2.41 | Moist to wet grasslands | -1.91 |
| *Lychnis flos-cuculi* | 1180 | -0.0127 | -50.31 | <.001 | -2.70 | 154 | -0.0056 | -8.09 | <.001 | -0.68 | 2890 | -0.0048 | -43.79 | <.001 | -2.23 | Moist to wet grasslands | -1.87 |
| *Hieracium pilosella* | 538 | -0.0095 | -31.78 | <.001 | -2.01 | 64 | -0.0058 | -10.12 | <.001 | -0.70 | 4046 | -0.0058 | -39.06 | <.001 | -2.70 | Heaths, inland dunes and semi-natural grasslands | -1.80 |
| *Potentilla erecta* | 896 | -0.0111 | -63.15 | <.001 | -2.34 | 85 | -0.0032 | -9.68 | <.001 | -0.39 | 3204 | -0.0054 | -44.18 | <.001 | -2.49 | Moist to wet grasslands | -1.74 |
| *Juncus articulatus* | 1290 | -0.0125 | -41.77 | <.001 | -2.64 | 176 | -0.0067 | -8.10 | <.001 | -0.82 | 1543 | -0.0027 | -44.16 | <.001 | -1.25 | Moist to wet grasslands | -1.57 |
| *Carex panicea* | 486 | -0.008 | -69.78 | <.001 | -1.70 | 25 | -0.0009 | -6.91 | <.001 | -0.11 | 2702 | -0.0056 | -45.84 | <.001 | -2.59 | Moist to wet grasslands | -1.47 |
| *Luzula campestris* agg. | 978 | -0.011 | -52.93 | <.001 | -2.33 | 134 | -0.0056 | -10.53 | <.001 | -0.69 | 1674 | -0.0024 | -36.51 | <.001 | -1.11 | Heaths, inland dunes and semi-natural grasslands | -1.38 |
| *Rumex acetosella* s. l. | 866 | -0.0099 | -35.39 | <.001 | -2.09 | 221 | -0.0144 | -13.76 | <.001 | -1.76 | 965 | -0.0006 | -12.38 | <.001 | -0.27 | Heaths, inland dunes and semi-natural grasslands | -1.37 |
| *Lotus corniculatus* agg. | 507 | -0.0054 | -33.28 | <.001 | -1.14 | 84 | -0.0059 | -11.45 | <.001 | -0.72 | 4168 | -0.0048 | -34.89 | <.001 | -2.22 | Heaths, inland dunes and semi-natural grasslands | -1.36 |
| *Festuca ovina* agg. | 448 | -0.0073 | -39.01 | <.001 | -1.55 | 104 | -0.0073 | -10.04 | <.001 | -0.89 | 2152 | -0.0028 | -34.62 | <.001 | -1.30 | Heaths, inland dunes and semi-natural grasslands | -1.24 |
| *Viola palustris* | 764 | -0.0097 | -66.83 | <.001 | -2.06 | 64 | -0.0015 | -6.30 | <.001 | -0.18 | 1558 | -0.0031 | -36.92 | <.001 | -1.42 | Moist to wet grasslands | -1.22 |
| *Ranunculus flammula* agg. | 964 | -0.0082 | -38.13 | <.001 | -1.73 | 158 | -0.0046 | -6.06 | <.001 | -0.56 | 1685 | -0.0029 | -38.47 | <.001 | -1.33 | Moist to wet grasslands | -1.21 |
| *Vicia cracca* agg. | 764 | -0.0055 | -32.07 | <.001 | -1.16 | 306 | -0.0132 | -13.87 | <.001 | -1.61 | 2634 | -0.0019 | -38.03 | <.001 | -0.87 | Mesic grasslands | -1.21 |
| *Lathyrus pratensis* | 986 | -0.0055 | -26.70 | <.001 | -1.16 | 219 | -0.0086 | -10.20 | <.001 | -1.05 | 2433 | -0.0029 | -48.33 | <.001 | -1.33 | Moist to wet grasslands | -1.18 |
| *Cardamine pratensis* agg. | 1324 | -0.01 | -25.69 | <.001 | -2.11 | 194 | -0.005 | -6.26 | <.001 | -0.61 | 1424 | -0.0017 | -37.34 | <.001 | -0.77 | Moist to wet grasslands | -1.17 |
| *Campanula rotundifolia* agg. | 372 | -0.005 | -33.62 | <.001 | -1.06 | 37 | -0.0021 | -9.82 | <.001 | -0.26 | 4148 | -0.0046 | -36.24 | <.001 | -2.12 | Heaths, inland dunes and semi-natural grasslands | -1.15 |
| *Agrostis stolonifera* agg. | 2554 | -0.0063 | -14.77 | <.001 | -1.33 | 558 | -0.0115 | -10.46 | <.001 | -1.41 | 1227 | -0.0013 | -27.79 | <.001 | -0.61 | Coastal and marine habitats | -1.12 |
| *Potentilla anserina* | 1218 | -0.0071 | -22.57 | <.001 | -1.49 | 360 | -0.0102 | -10.91 | <.001 | -1.25 | 585 | -0.0006 | -25.28 | <.001 | -0.28 | Coastal and marine habitats | -1.01 |
| *Hydrocotyle vulgaris* | 929 | -0.0129 | -68.22 | <.001 | -2.73 | 60 | -0.0017 | -6.85 | <.001 | -0.21 | 15 | 0 | -12.42 | <.001 | -0.01 | Bogs, transition mires, marshes and fens | -0.98 |
| *Juncus acutiflorus* | 197 | -0.0018 | -35.85 | <.001 | -0.38 | 48 | -0.0015 | -6.43 | <.001 | -0.19 | 2752 | -0.0051 | -40.61 | <.001 | -2.34 | Moist to wet grasslands | -0.97 |
| *Bistorta officinalis* | 52 | -0.0007 | -28.39 | <.001 | -0.15 | 62 | -0.0015 | -6.14 | <.001 | -0.18 | 3288 | -0.0052 | -44.68 | <.001 | -2.41 | Moist to wet grasslands | -0.91 |
| *Nardus stricta* | 264 | -0.0043 | -48.71 | <.001 | -0.92 | 35 | -0.0031 | -7.06 | <.001 | -0.38 | 1530 | -0.0031 | -34.45 | <.001 | -1.42 | Heaths, inland dunes and semi-natural grasslands | -0.90 |
| *Achillea ptarmica* agg. | 515 | -0.007 | -68.97 | <.001 | -1.48 | 104 | -0.0043 | -13.79 | <.001 | -0.52 | 819 | -0.0012 | -34.58 | <.001 | -0.55 | Moist to wet grasslands | -0.85 |
| *Equisetum palustre* | 1162 | -0.004 | -20.85 | <.001 | -0.85 | 263 | -0.0049 | -6.74 | <.001 | -0.60 | 1643 | -0.0023 | -40.81 | <.001 | -1.08 | Moist to wet grasslands | -0.84 |
| *Galium saxatile* | 738 | -0.0063 | -41.16 | <.001 | -1.33 | 112 | -0.0028 | -6.26 | <.001 | -0.34 | 973 | -0.0016 | -28.33 | <.001 | -0.76 | Heaths, inland dunes and semi-natural grasslands | -0.81 |
| *Carex echinata* | 201 | -0.0032 | -59.99 | <.001 | -0.67 | 21 | -0.0009 | -7.66 | <.001 | -0.11 | 1606 | -0.0035 | -37.46 | <.001 | -1.63 | Moist to wet grasslands | -0.80 |
| *Carex leporina* | 490 | -0.0049 | -38.82 | <.001 | -1.03 | 107 | -0.0048 | -11.87 | <.001 | -0.58 | 982 | -0.0016 | -35.59 | <.001 | -0.72 | Moist to wet grasslands | -0.78 |
| *Festuca pratensis* s. l. | 553 | -0.0046 | -27.75 | <.001 | -0.97 | 191 | -0.0074 | -10.64 | <.001 | -0.91 | 714 | -0.001 | -35.08 | <.001 | -0.45 | Moist to wet grasslands | -0.78 |
| *Juncus conglomeratus* | 524 | -0.0051 | -52.89 | <.001 | -1.09 | 82 | -0.0032 | -10.24 | <.001 | -0.39 | 1048 | -0.0017 | -41.59 | <.001 | -0.79 | Moist to wet grasslands | -0.76 |
| *Prunella vulgaris* | 378 | -0.004 | -39.41 | <.001 | -0.85 | 108 | -0.0049 | -12.40 | <.001 | -0.60 | 1226 | -0.0018 | -46.60 | <.001 | -0.83 | Heaths, inland dunes and semi-natural grasslands | -0.76 |
| *Dactylorhiza majalis* agg. | 294 | -0.0051 | -67.22 | <.001 | -1.08 | 16 | -0.0007 | -6.24 | <.001 | -0.09 | 1059 | -0.0023 | -38.16 | <.001 | -1.04 | Moist to wet grasslands | -0.74 |
| *Galium uliginosum* | 402 | -0.0042 | -51.90 | <.001 | -0.88 | 35 | -0.0009 | -5.88 | <.001 | -0.11 | 1536 | -0.0026 | -40.70 | <.001 | -1.22 | Moist to wet grasslands | -0.74 |
| *Succisa pratensis* | 216 | -0.0043 | -68.17 | <.001 | -0.92 | 11 | -0.0006 | -8.21 | <.001 | -0.07 | 1353 | -0.0027 | -43.53 | <.001 | -1.23 | Moist to wet grasslands | -0.74 |
| *Persicaria amphibia* | 1272 | -0.0048 | -22.20 | <.001 | -1.02 | 250 | -0.0067 | -8.52 | <.001 | -0.82 | 493 | -0.0006 | -24.46 | <.001 | -0.30 | Standing waters | -0.72 |
| *Centaurea jacea* agg. | 146 | -0.0017 | -30.77 | <.001 | -0.36 | 12 | -0.0012 | -7.87 | <.001 | -0.14 | 3258 | -0.0035 | -34.94 | <.001 | -1.60 | Heaths, inland dunes and semi-natural grasslands | -0.70 |
| *Alopecurus geniculatus* | 760 | -0.0057 | -24.91 | <.001 | -1.20 | 115 | -0.0055 | -10.88 | <.001 | -0.67 | 80 | -0.0001 | -12.15 | <.001 | -0.06 | Moist to wet grasslands | -0.64 |
| *Trifolium arvense* | 268 | -0.0042 | -26.80 | <.001 | -0.90 | 59 | -0.0071 | -11.12 | <.001 | -0.87 | 107 | -0.0002 | -8.14 | <.001 | -0.08 | Mesic grasslands | -0.61 |
| *Carex flava* agg. | 152 | -0.0031 | -60.16 | <.001 | -0.65 | 16 | -0.0008 | -8.98 | <.001 | -0.09 | 998 | -0.0022 | -41.60 | <.001 | -1.03 | Moist to wet grasslands | -0.59 |
| *Juncus inflexus* | 326 | -0.0017 | -24.37 | <.001 | -0.36 | 29 | -0.0008 | -6.08 | <.001 | -0.10 | 1996 | -0.0026 | -36.67 | <.001 | -1.22 | Fresh water vegetation, springs and reeds | -0.56 |
| *Jasione montana* | 238 | -0.0052 | -34.23 | <.001 | -1.11 | 24 | -0.0035 | -9.59 | <.001 | -0.42 | 120 | -0.0002 | -10.58 | <.001 | -0.10 | Heaths, inland dunes and semi-natural grasslands | -0.54 |
| *Epilobium angustifolium* | 370 | -0.0014 | -20.04 | <.001 | -0.29 | 272 | -0.0062 | -11.97 | <.001 | -0.76 | 1104 | -0.0007 | -23.70 | <.001 | -0.33 | Coastal and marine habitats | -0.46 |
| *Linaria vulgaris* agg. | 271 | -0.0024 | -27.86 | <.001 | -0.51 | 107 | -0.005 | -13.26 | <.001 | -0.61 | 1532 | -0.0006 | -14.51 | <.001 | -0.26 | Scrubs, copses and field hedges | -0.46 |
| *Tripleurospermum maritimum* agg. | 130 | -0.0012 | -19.13 | <.001 | -0.26 | 102 | -0.009 | -14.75 | <.001 | -1.10 | 46 | -0.0001 | -16.06 | <.001 | -0.04 | Coastal and marine habitats | -0.46 |
| *Danthonia decumbens* | 150 | -0.0024 | -40.47 | <.001 | -0.50 | 26 | -0.0023 | -6.76 | <.001 | -0.28 | 937 | -0.0012 | -25.62 | <.001 | -0.55 | Heaths, inland dunes and semi-natural grasslands | -0.44 |
| *Sedum acre* | 130 | -0.0022 | -26.29 | <.001 | -0.46 | 20 | -0.0024 | -8.36 | <.001 | -0.30 | 745 | -0.0012 | -30.19 | <.001 | -0.57 | Heaths, inland dunes and semi-natural grasslands | -0.44 |
| *Tussilago farfara* | 198 | -0.0016 | -24.27 | <.001 | -0.35 | 148 | -0.0067 | -15.28 | <.001 | -0.82 | 334 | -0.0003 | -15.33 | <.001 | -0.15 | Ruderal, fringe and tall forb communities, clearings | -0.44 |
| *Juncus bufonius* agg. | 308 | -0.0036 | -43.62 | <.001 | -0.77 | 67 | -0.0028 | -9.26 | <.001 | -0.35 | 130 | -0.0003 | -23.44 | <.001 | -0.12 | Standing waters | -0.41 |
| *Juncus squarrosus* | 176 | -0.0031 | -46.17 | <.001 | -0.67 | 24 | -0.0018 | -7.17 | <.001 | -0.21 | 284 | -0.0007 | -27.84 | <.001 | -0.32 | Heaths, inland dunes and semi-natural grasslands | -0.40 |
| *Alchemilla vulgaris* agg. | 110 | -0.0013 | -42.43 | <.001 | -0.27 | 18 | -0.0008 | -7.96 | <.001 | -0.09 | 1198 | -0.0017 | -44.13 | <.001 | -0.79 | Moist to wet grasslands | -0.39 |
| *Viola tricolor* agg. | 126 | -0.0021 | -27.45 | <.001 | -0.44 | 44 | -0.0044 | -12.53 | <.001 | -0.54 | 54 | -0.0001 | -14.85 | <.001 | -0.04 | Coastal and marine habitats | -0.34 |
| *Stachys palustris* | 802 | -0.0025 | -23.02 | <.001 | -0.52 | 214 | -0.0024 | -4.48 | <.001 | -0.29 | 741 | -0.0004 | -13.91 | <.001 | -0.18 | Fresh water vegetation, springs and reeds | -0.33 |
| *Juncus filiformis* | 180 | -0.0027 | -56.96 | <.001 | -0.58 | 17 | -0.0007 | -6.21 | <.001 | -0.08 | 272 | -0.0005 | -30.70 | <.001 | -0.25 | Moist to wet grasslands | -0.30 |
| *Bidens tripartita* | 228 | -0.0027 | -46.44 | <.001 | -0.58 | 73 | -0.0019 | -7.10 | <.001 | -0.23 | 26 | -0.0001 | -11.71 | <.001 | -0.03 | Standing waters | -0.28 |
| *Erodium cicutarium* agg. | 97 | -0.0014 | -25.63 | <.001 | -0.29 | 22 | -0.0035 | -9.90 | <.001 | -0.43 | 83 | -0.0002 | -8.35 | <.001 | -0.09 | Heaths, inland dunes and semi-natural grasslands | -0.27 |
| *Festuca arundinacea* | 371 | -0.0015 | -19.07 | <.001 | -0.32 | 126 | -0.0027 | -8.66 | <.001 | -0.33 | 378 | -0.0002 | -11.74 | <.001 | -0.10 | Coastal and marine habitats | -0.25 |
| *Hieracium umbellatum* | 93 | -0.0025 | -39.83 | <.001 | -0.53 | 20 | -0.001 | -8.01 | <.001 | -0.12 | 369 | -0.0002 | -13.78 | <.001 | -0.10 | Coastal and marine habitats | -0.25 |
| *Mentha arvensis* | 84 | -0.0008 | -32.58 | <.001 | -0.16 | 82 | -0.0031 | -12.47 | <.001 | -0.37 | 234 | -0.0003 | -20.27 | <.001 | -0.12 | Moist to wet grasslands | -0.22 |
| *Poa palustris* | 190 | -0.0012 | -30.71 | <.001 | -0.25 | 138 | -0.0028 | -8.16 | <.001 | -0.34 | 122 | -0.0001 | -10.86 | <.001 | -0.05 | Linear and running surface waters | -0.21 |
| *Gnaphalium uliginosum* | 84 | -0.0009 | -23.99 | <.001 | -0.19 | 37 | -0.0029 | -11.43 | <.001 | -0.35 | 28 | 0 | -8.94 | <.001 | -0.02 | Standing waters | -0.19 |
| *Senecio viscosus* | 59 | -0.0007 | -17.42 | <.001 | -0.15 | 33 | -0.0028 | -11.10 | <.001 | -0.34 | 28 | 0 | -8.39 | <.001 | -0.01 | Coastal and marine habitats | -0.17 |
| *Sonchus arvensis* agg. | 290 | -0.0014 | -16.44 | <.001 | -0.30 | 30 | -0.0014 | -9.12 | <.001 | -0.18 | 48 | 0 | -5.80 | <.001 | -0.02 | Coastal and marine habitats | -0.17 |
| *Hypericum maculatum* agg. | 112 | -0.0008 | -35.56 | <.001 | -0.18 | 37 | -0.0009 | -6.81 | <.001 | -0.11 | 408 | -0.0004 | -22.52 | <.001 | -0.17 | Heaths, inland dunes and semi-natural grasslands | -0.15 |
| *Gnaphalium sylvaticum* | 24 | -0.0004 | -20.50 | <.001 | -0.08 | 44 | -0.0025 | -14.07 | <.001 | -0.31 | 52 | -0.0001 | -14.62 | <.001 | -0.03 | Heaths, inland dunes and semi-natural grasslands | -0.14 |
| *Oenanthe aquatica* agg. | 431 | -0.001 | -12.25 | <.001 | -0.21 | 68 | -0.0015 | -5.21 | <.001 | -0.18 | 22 | 0 | -6.12 | <.001 | -0.02 | Standing waters | -0.14 |
| *Epipactis helleborine* agg. | 194 | -0.0014 | -29.55 | <.001 | -0.29 | 18 | 0.001 | 9.15 | <.001 | 0.13 | 872 | -0.0004 | -15.33 | <.001 | -0.20 | Dry to moderately moist forests | -0.12 |
| *Elymus repens* s. str*.* | 727 | 0.0025 | 21.11 | <.001 | 0.53 | 594 | -0.0134 | -11.34 | <.001 | -1.63 | 3962 | 0.0018 | 27.24 | <.001 | 0.85 | Scrubs, copses and field hedges | -0.09 |
| *Elymus caninus* | 48 | -0.0004 | -20.45 | <.001 | -0.08 | 18 | -0.0006 | -6.66 | <.001 | -0.07 | 616 | 0.0003 | 14.51 | <.001 | 0.13 | Moist to wet forests | -0.01 |
| *Veronica hederifolia* agg. | 196 | 0.0012 | 25.41 | <.001 | 0.25 | 14 | 0.0009 | 8.86 | <.001 | 0.11 | 209 | -0.0001 | -9.72 | <.001 | -0.07 | Dry to moderately moist forests | 0.10 |
| *Rosa multiflora* | 11 | 0.0001 | 16.51 | <.001 | 0.03 | 36 | 0.0018 | 12.64 | <.001 | 0.23 | 326 | 0.0004 | 25.10 | <.001 | 0.19 | Scrubs, copses and field hedges | 0.15 |
| *Cardamine flexuosa* | 370 | 0.0011 | 19.97 | <.001 | 0.23 | 18 | 0.0008 | 6.89 | <.001 | 0.10 | 282 | 0.0003 | 16.31 | <.001 | 0.14 | Dry to moderately moist forests | 0.16 |
| *Chrysosplenium oppositifolium* | 774 | 0.0014 | 11.90 | <.001 | 0.30 | 7 | 0.0012 | 12.31 | <.001 | 0.14 | 1250 | 0.0009 | 16.03 | <.001 | 0.40 | Moist to wet forests | 0.28 |
| *Chelidonium majus* | 104 | 0.0009 | 30.05 | <.001 | 0.19 | 41 | 0.0033 | 13.69 | <.001 | 0.40 | 1488 | 0.0007 | 17.02 | <.001 | 0.32 | Scrubs, copses and field hedges | 0.30 |
| *Quercus rubra* | 208 | 0.0013 | 26.60 | <.001 | 0.27 | 105 | 0.0048 | 11.99 | <.001 | 0.59 | 461 | 0.0004 | 21.35 | <.001 | 0.18 | Dry to moderately moist forests | 0.35 |
| *Fallopia bohemica_Fallopia japonica_Fallopia sachalinensis* | 138 | 0.0008 | 21.52 | <.001 | 0.16 | 208 | 0.0056 | 10.52 | <.001 | 0.69 | 439 | 0.0005 | 32.12 | <.001 | 0.25 | Anthropogenic | 0.37 |
| *Symphoricarpos albus* | 78 | 0.0006 | 28.23 | <.001 | 0.13 | 122 | 0.007 | 14.45 | <.001 | 0.85 | 400 | 0.0005 | 29.71 | <.001 | 0.21 | Anthropogenic | 0.40 |
| *Ficaria verna* s. l. | 2319 | 0.0055 | 22.15 | <.001 | 1.16 | 72 | 0.0046 | 15.85 | <.001 | 0.56 | 2699 | -0.0009 | -12.18 | <.001 | -0.44 | Dry to moderately moist forests | 0.43 |
| *Taxus baccata* | 116 | 0.0008 | 30.14 | <.001 | 0.17 | 100 | 0.0091 | 17.72 | <.001 | 1.11 | 360 | 0.0004 | 14.16 | <.001 | 0.18 | Anthropogenic | 0.49 |
| *Ribes rubrum* agg. | 1397 | 0.0041 | 32.59 | <.001 | 0.87 | 98 | 0.0077 | 19.34 | <.001 | 0.95 | 1218 | 0.0008 | 25.47 | <.001 | 0.35 | Moist to wet forests | 0.72 |
| *Impatiens noli-tangere* | 1702 | 0.0053 | 29.71 | <.001 | 1.13 | 54 | 0.0032 | 11.64 | <.001 | 0.39 | 3682 | 0.0016 | 17.56 | <.001 | 0.72 | Moist to wet forests | 0.75 |
| *Impatiens glandulifera* | 388 | 0.002 | 38.02 | <.001 | 0.42 | 99 | 0.0044 | 14.20 | <.001 | 0.54 | 3699 | 0.0029 | 38.41 | <.001 | 1.35 | Moist to wet forests | 0.77 |
| *Humulus lupulus* | 1240 | 0.0018 | 14.86 | <.001 | 0.39 | 193 | 0.007 | 13.67 | <.001 | 0.86 | 3668 | 0.0024 | 41.05 | <.001 | 1.11 | Moist to wet forests | 0.79 |
| *Salix fragilis* agg. | 1150 | 0.0054 | 38.60 | <.001 | 1.14 | 208 | 0.0038 | 7.52 | <.001 | 0.47 | 5566 | 0.0018 | 18.23 | <.001 | 0.83 | Moist to wet forests | 0.81 |
| *Sorbus aucuparia* | 3506 | 0.0058 | 12.10 | <.001 | 1.23 | 533 | 0.0171 | 10.35 | <.001 | 2.09 | 6278 | -0.0017 | -18.38 | <.001 | -0.79 | Dry to moderately moist forests | 0.85 |
| *Prunus serotina* | 1558 | 0.0051 | 24.79 | <.001 | 1.09 | 255 | 0.0108 | 13.13 | <.001 | 1.32 | 360 | 0.0004 | 21.17 | <.001 | 0.20 | Dry to moderately moist forests | 0.87 |
| *Athyrium filix-femina* | 1942 | 0.0052 | 24.98 | <.001 | 1.10 | 189 | 0.0093 | 15.58 | <.001 | 1.13 | 4008 | 0.0023 | 22.86 | <.001 | 1.08 | Moist to wet forests | 1.10 |
| *Juglans regia* | 21 | 0.0002 | 20.12 | <.001 | 0.03 | 32 | 0.0022 | 12.05 | <.001 | 0.26 | 6497 | 0.0066 | 68.73 | <.001 | 3.04 | Scrubs, copses and field hedges | 1.11 |
| *Prunus padus* | 1417 | 0.0042 | 26.59 | <.001 | 0.89 | 208 | 0.017 | 20.27 | <.001 | 2.08 | 4686 | 0.0023 | 32.55 | <.001 | 1.06 | Moist to wet forests | 1.34 |
| *Circaea lutetiana* | 2698 | 0.0063 | 17.35 | <.001 | 1.34 | 120 | 0.0108 | 19.00 | <.001 | 1.32 | 3252 | 0.0032 | 44.64 | <.001 | 1.47 | Dry to moderately moist forests | 1.38 |
| *Galium aparine* agg*.* | 2587 | 0.0093 | 42.40 | <.001 | 1.97 | 466 | 0.0071 | 9.25 | <.001 | 0.87 | 12650 | 0.003 | 23.39 | <.001 | 1.38 | Scrubs, copses and field hedges | 1.40 |
| *Rubus* sect. *Caesii* | 724 | 0.0016 | 22.10 | <.001 | 0.35 | 101 | 0.0062 | 17.43 | <.001 | 0.76 | 9583 | 0.0073 | 65.89 | <.001 | 3.38 | Scrubs, copses and field hedges | 1.49 |
| *Dryopteris carthusiana* agg. | 4632 | 0.0099 | 22.99 | <.001 | 2.09 | 382 | 0.0151 | 12.31 | <.001 | 1.84 | 3582 | 0.0017 | 18.03 | <.001 | 0.81 | Dry to moderately moist forests | 1.58 |
| *Carex remota* | 2979 | 0.0145 | 43.87 | <.001 | 3.08 | 104 | 0.008 | 17.11 | <.001 | 0.98 | 1942 | 0.002 | 29.95 | <.001 | 0.94 | Standing waters | 1.67 |
| *Geranium robertianum* agg. | 2120 | 0.0076 | 27.42 | <.001 | 1.60 | 90 | 0.0065 | 17.62 | <.001 | 0.80 | 11282 | 0.0063 | 47.14 | <.001 | 2.92 | Scrubs, copses and field hedges | 1.77 |
| *Impatiens parviflora* | 1948 | 0.0109 | 45.78 | <.001 | 2.31 | 483 | 0.0241 | 15.77 | <.001 | 2.94 | 1208 | 0.0008 | 21.84 | <.001 | 0.37 | Dry to moderately moist forests | 1.87 |
| *Alliaria petiolata* | 1180 | 0.0055 | 29.79 | <.001 | 1.15 | 250 | 0.0167 | 19.67 | <.001 | 2.05 | 9350 | 0.0079 | 68.08 | <.001 | 3.66 | Scrubs, copses and field hedges | 2.29 |
| *Galeobdolon luteum* agg. | 3294 | 0.013 | 27.37 | <.001 | 2.75 | 220 | 0.0184 | 20.16 | <.001 | 2.25 | 5666 | 0.0041 | 38.89 | <.001 | 1.92 | Dry to moderately moist forests | 2.31 |
| *Glechoma hederacea* agg. | 4288 | 0.015 | 49.83 | <.001 | 3.17 | 782 | 0.0188 | 18.15 | <.001 | 2.30 | 7063 | 0.0038 | 44.29 | <.001 | 1.78 | Moist to wet forests | 2.42 |
| *Geum urbanum* | 2633 | 0.0083 | 23.78 | <.001 | 1.75 | 348 | 0.0263 | 21.28 | <.001 | 3.22 | 15057 | 0.0104 | 64.45 | <.001 | 4.82 | Scrubs, copses and field hedges | 3.26 |
| *Fraxinus excelsior* | 3544 | 0.01 | 16.82 | <.001 | 2.12 | 444 | 0.0216 | 18.70 | <.001 | 2.64 | 22786 | 0.0115 | 64.85 | <.001 | 5.34 | Scrubs, copses and field hedges | 3.37 |
| *Rubus* sect. *Rubus* | 5653 | 0.0209 | 36.33 | <.001 | 4.43 | 796 | 0.0184 | 12.50 | <.001 | 2.24 | 17402 | 0.0105 | 71.24 | <.001 | 4.86 | Scrubs, copses and field hedges | 3.85 |
| *Urtica dioica* s. l. | 7535 | 0.0279 | 51.70 | <.001 | 5.91 | 1270 | 0.014 | 10.65 | <.001 | 1.72 | 28635 | 0.0093 | 45.61 | <.001 | 4.29 | Scrubs, copses and field hedges | 3.97 |

.
 Notes: Given are raw and scaled Beals trends of species within each state, including n-, t-, and p- values (after Holm adjustment) according to t-tests. In addition, each species preferred habitat
 type and its mean scaled Beals trend across all states, derived from the three trends of each state, are given. Habitat type preferences are based on the fidelity (Φ) of species to habitat types in
 all states taken together. Beals trends were scaled per state as Beals trend/sd(Beals trends).

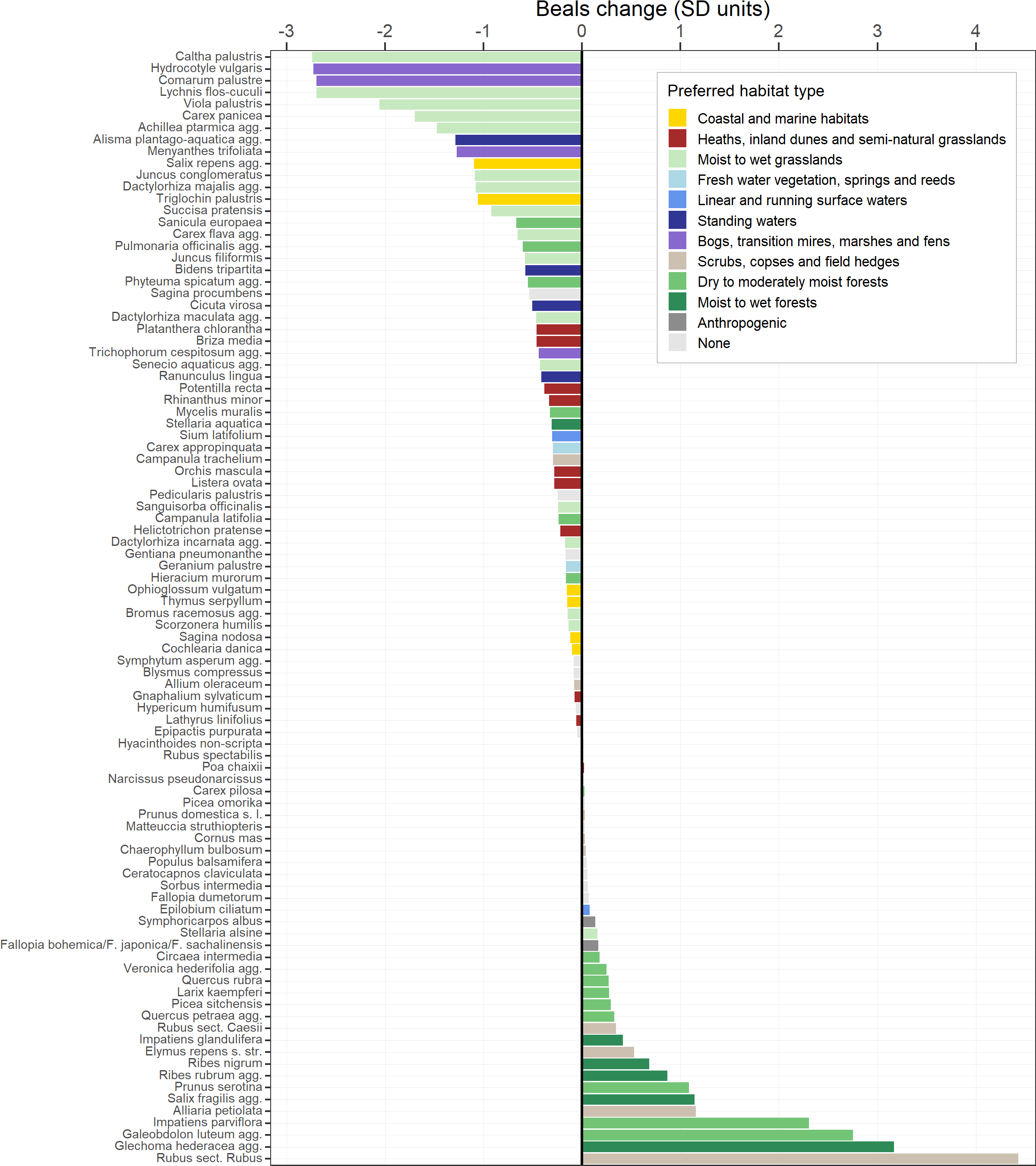

**Figure S1.5** Beals trends of species that showed significant and consistent trends for Beals and frequency in SH (94 species). Colours indicate the species’ preferred habitat type.

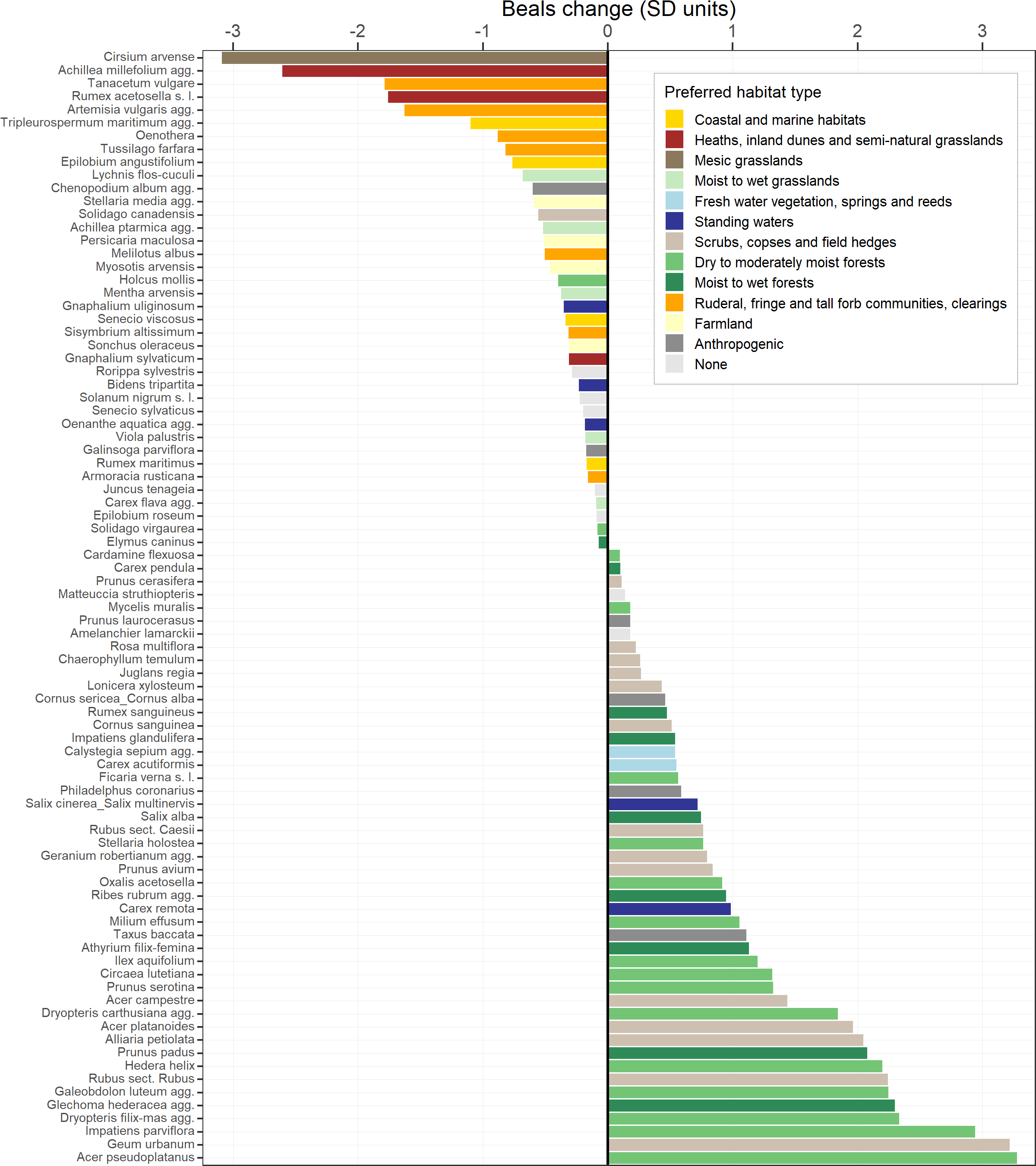

**Figure S1.6** Beals trends of species that showed significant and consistent trends for Beals and frequency in HH (85 species). Colours indicate the species’ preferred habitat type.

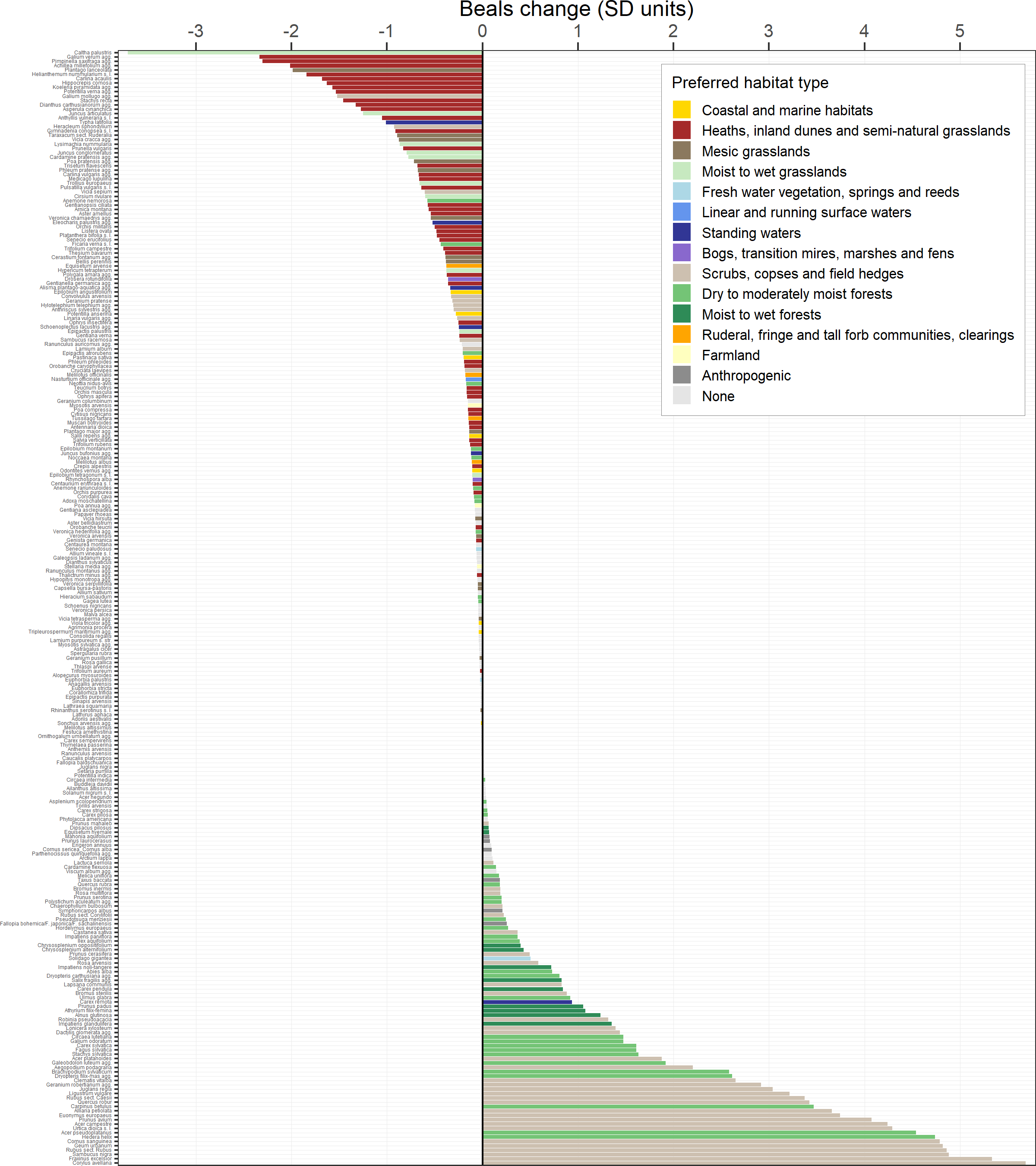

**Figure S1.7** Beals trends of species that showed significant and consistent trends for Beals and frequency in BW (256 species). Colours indicate the species’ preferred habitat type.

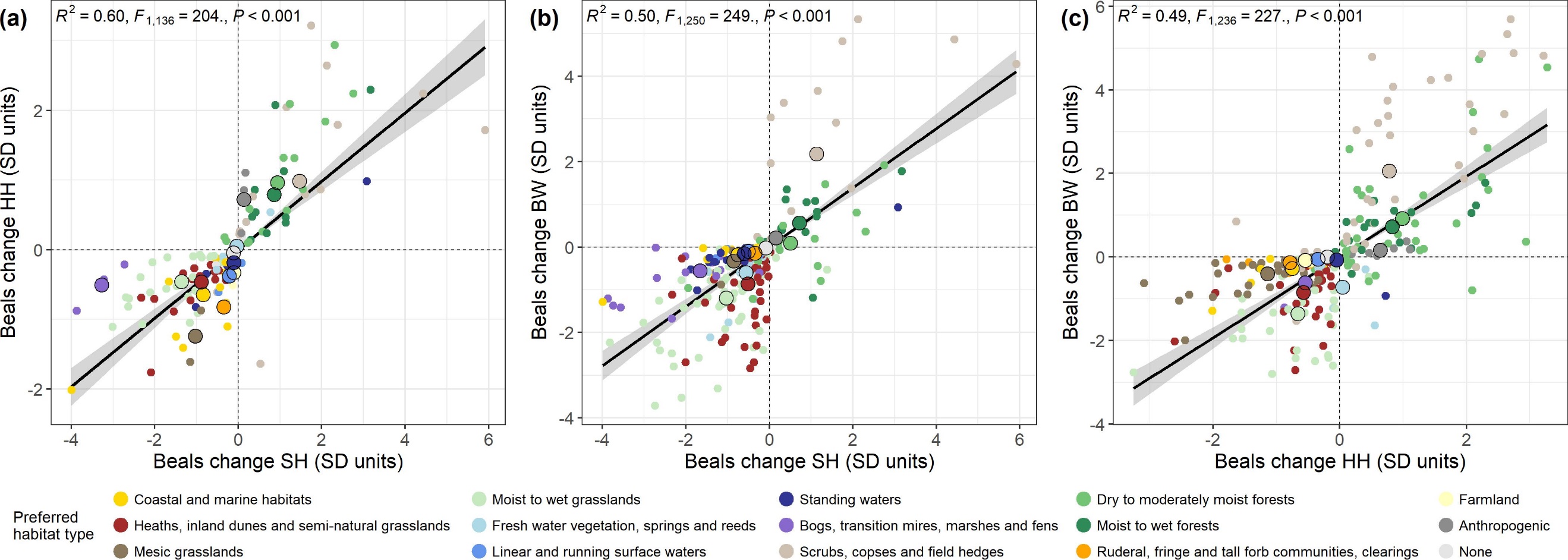

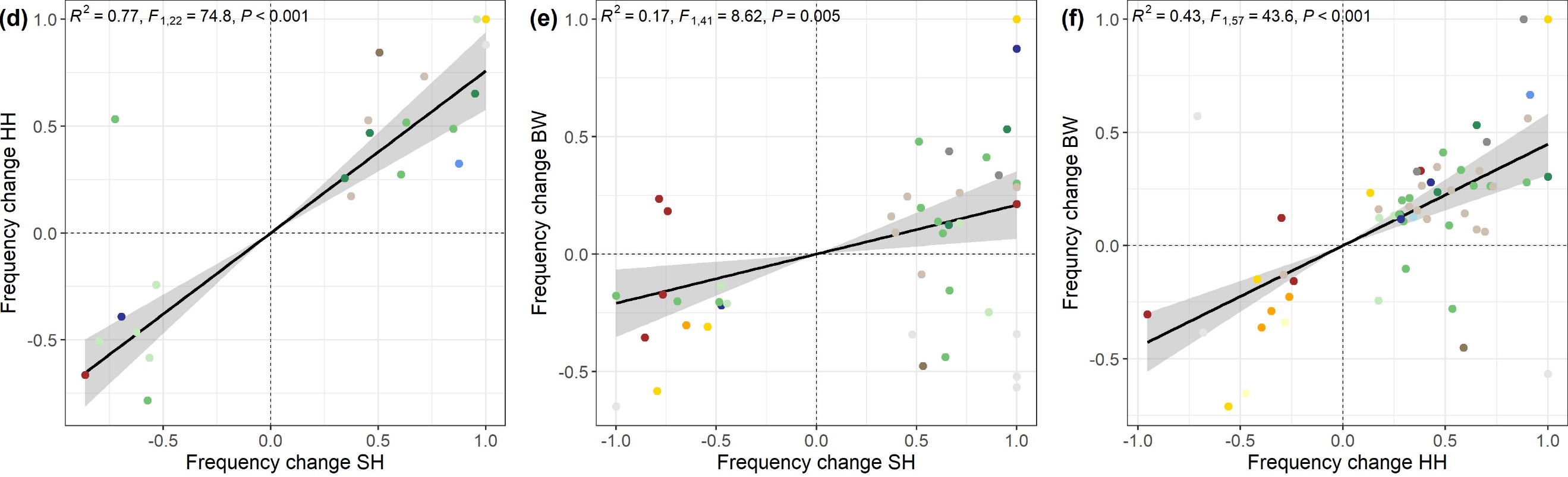

**Figure S1.8** Comparison of species trends in two states each, concerning **(a-c)** Beals and **(d-f)** frequency. Species are coloured by their preferred habitat type. Thick dots in **(a-c)** represent mean trends per species of a preferred habitat type. Only species are included that showed a significant trend in both states each. Regression lines were obtained from a linear model, including 95% confidence intervals.

**Table S1.5** Mean and Median Beals trends of species grouped by their preferred habitat type and by state.

| **Habitat type preferred** | **State** | **n** | **Mean trend** | **Median trend** | **p** |
| --- | --- | --- | --- | --- | --- |
| Anthropogenic | SH | 4 | 0.136 | 0.149 | 0.125 |
| Anthropogenic | HH | 13 | 0.338 | 0.240 | 0.04 |
| Anthropogenic | BW | 8 | 0.163 | 0.140 | 0.008 |
| Anthropogenic | Across | 3 | 0.418 | 0.400 | 0.25 |
| Bogs, transition mires, marshes and fens | SH | 25 | -1.531 | -1.274 | < 0.001 |
| Bogs, transition mires, marshes and fens | HH | 3 | -0.503 | -0.423 | 0.25 |
| Bogs, transition mires, marshes and fens | BW | 29 | -0.463 | -0.311 | < 0.001 |
| Bogs, transition mires, marshes and fens | Across | 2 | -1.484 | -1.484 | 0.5 |
| Coastal and marine habitats | SH | 67 | -0.427 | -0.232 | < 0.001 |
| Coastal and marine habitats | HH | 18 | -0.576 | -0.344 | < 0.001 |
| Coastal and marine habitats | BW | 25 | -0.161 | -0.066 | < 0.001 |
| Coastal and marine habitats | Across | 10 | -0.665 | -0.400 | 0.002 |
| Dry to moderately moist forests | SH | 36 | 0.267 | 0.038 | 0.315 |
| Dry to moderately moist forests | HH | 45 | 0.872 | 0.522 | < 0.001 |
| Dry to moderately moist forests | BW | 84 | 0.339 | 0.002 | 0.06 |
| Dry to moderately moist forests | Across | 11 | 0.887 | 0.845 | 0.003 |
| Farmland | SH | 2 | -0.106 | -0.106 | 0.5 |
| Farmland | HH | 10 | -0.379 | -0.356 | 0.002 |
| Farmland | BW | 4 | -0.078 | -0.068 | 0.125 |
| Fresh water vegetation, springs and reeds | SH | 18 | -0.469 | -0.410 | < 0.001 |
| Fresh water vegetation, springs and reeds | HH | 6 | 0.128 | 0.153 | 0.438 |
| Fresh water vegetation, springs and reeds | BW | 28 | -0.523 | -0.277 | < 0.001 |
| Fresh water vegetation, springs and reeds | Across | 2 | -0.445 | -0.445 | 0.5 |
| Heaths, inland dunes and semi-natural grasslands | SH | 59 | -0.469 | -0.291 | < 0.001 |
| Heaths, inland dunes and semi-natural grasslands | HH | 34 | -0.530 | -0.334 | < 0.001 |
| Heaths, inland dunes and semi-natural grasslands | BW | 220 | -0.546 | -0.298 | < 0.001 |
| Heaths, inland dunes and semi-natural grasslands | Across | 17 | -0.816 | -0.759 | < 0.001 |
| Linear and running surface waters | SH | 13 | -0.454 | -0.305 | < 0.001 |
| Linear and running surface waters | HH | 3 | -0.379 | -0.342 | 0.25 |
| Linear and running surface waters | BW | 8 | -0.092 | -0.063 | 0.008 |
| Linear and running surface waters | Across | 1 | -0.213 | -0.213 | 1 |
| Mesic grasslands | SH | 4 | -0.661 | -0.706 | 0.125 |
| Mesic grasslands | HH | 40 | -1.017 | -0.840 | < 0.001 |
| Mesic grasslands | BW | 35 | -0.391 | -0.200 | < 0.001 |
| Mesic grasslands | Across | 2 | -0.914 | -0.914 | 0.5 |
| Moist to wet forests | SH | 25 | 0.361 | 0.300 | 0.052 |
| Moist to wet forests | HH | 22 | 0.706 | 0.471 | < 0.001 |
| Moist to wet forests | BW | 26 | 0.389 | 0.377 | 0.006 |
| Moist to wet forests | Across | 10 | 0.897 | 0.778 | 0.004 |
| Moist to wet grasslands | SH | 57 | -1.036 | -0.974 | < 0.001 |
| Moist to wet grasslands | HH | 35 | -0.660 | -0.558 | < 0.001 |
| Moist to wet grasslands | BW | 92 | -1.020 | -0.758 | < 0.001 |
| Moist to wet grasslands | Across | 27 | -1.051 | -0.849 | < 0.001 |
| None | SH | 31 | -0.068 | -0.063 | 0.001 |
| None | HH | 16 | -0.089 | -0.114 | 0.051 |
| None | BW | 143 | -0.027 | -0.025 | < 0.001 |
| Ruderal, fringe and tall forb communities, clearings | SH | 1 | -0.348 | -0.348 | 1 |
| Ruderal, fringe and tall forb communities, clearings | HH | 15 | -0.713 | -0.503 | < 0.001 |
| Ruderal, fringe and tall forb communities, clearings | BW | 9 | -0.135 | -0.123 | 0.004 |
| Ruderal, fringe and tall forb communities, clearings | Across | 1 | -0.438 | -0.438 | 1 |
| Scrubs, copses and field hedges | SH | 20 | 1.102 | 0.371 | 0.002 |
| Scrubs, copses and field hedges | HH | 46 | 0.712 | 0.482 | < 0.001 |
| Scrubs, copses and field hedges | BW | 70 | 1.328 | 0.579 | < 0.001 |
| Scrubs, copses and field hedges | Across | 13 | 1.725 | 1.495 | 0.002 |
| Standing waters | SH | 27 | -0.568 | -0.529 | < 0.001 |
| Standing waters | HH | 13 | -0.040 | -0.206 | 0.497 |
| Standing waters | BW | 28 | -0.234 | -0.150 | < 0.001 |
| Standing waters | Across | 6 | -0.010 | -0.233 | 0.438 |

.
 Notes: Only species were included per group that showed a significant trend in the respective state and that had a
 preferred habitat type assigned. The state category “Across” includes all species that showed a significant trend in all
 states and its values refer to the mean and median of those species mean trends across all states, derived from the three
 trends of each state. *n* is given for number of species with a significant trend that are included in each group. p-values
 indicate whether a group’s trend deviated from zero change according to Wilcoxon signed rank tests.

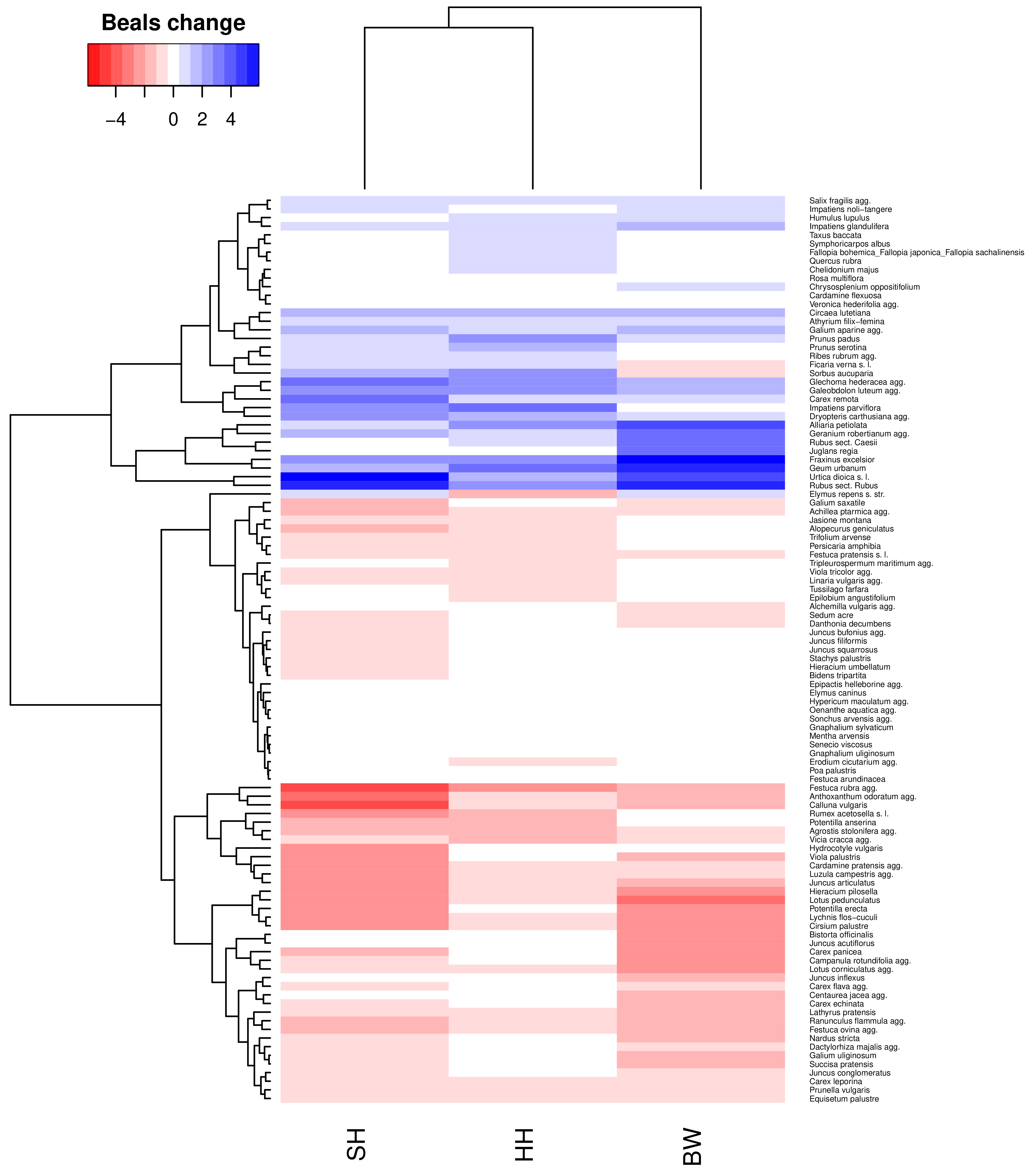

**Figure S1.9** Beals trends in each state of the 105 species that showed a significant trend in all states. Beals trends were scaled per state as Beals trend/sd(Beals trends).

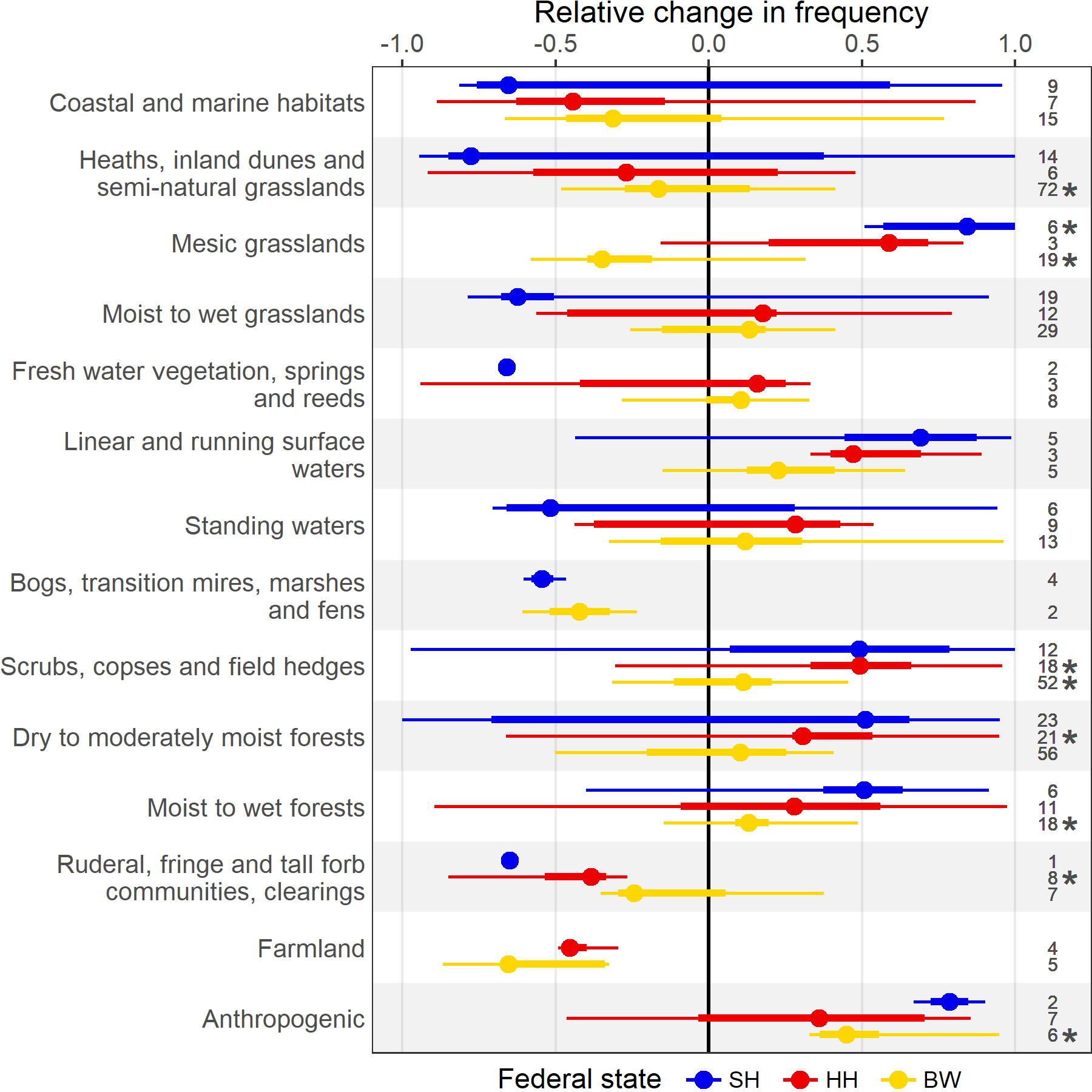

**Figure S1.10** Frequency trends of species grouped by their preferred habitat type separately by state. Points, thick and thin whiskers show the median, 50% and 95% data range. Only species are included per group that showed a significant trend and that had a preferred habitat type assigned. *n* is given for number of species with a significant trend that are included in each combination of state and preferred habitat type. Asterisks indicate whether a group’s trend deviated from zero change according to Wilcoxon signed rank tests. Habitat types correspond to the following EUNIS habitat types (European Nature Information System; European Environment Agency; Moss (2008); 2021 version if not stated otherwise): Coastal and marine habitats (N1, N2, N3), Heaths, inland dunes and semi-natural grasslands (S4, R1), Mesic grasslands (R2), Moist to wet grasslands (R3), Fresh water vegetation, springs and reeds (C3, D5; based on EUNIS 2012 version) , Linear and running surface waters (C2, C3; based on EUNIS 2012 version), Standing waters (C1, C3; based on EUNIS 2012 version), Bogs, transition mires, marshes and fens (D1, D2, D4, D5; based on EUNIS 2012 version), Scrubs, copses and field hedges (S3, T4,V4), Dry to moderately moist forests (T1, T3, V6), Moist to wet forests (T1, T3), Ruderal, fringe and tall forb communities, clearings (V3, T4, R5), Farmland (V1, V5, V6), Anthropogenic (V2, V3).

**Table S1.6** Mean and Median Beals trends of species grouped by their non-native status and by state.

| **Non-native status** | **State** | **n** | **Mean trend** | **Median trend** | **p** |
| --- | --- | --- | --- | --- | --- |
| Native | SH | 355 | -0.452 | -0.282 | < 0.001 |
| Native | HH | 253 | -0.066 | -0.184 | 0.023 |
| Native | BW | 702 | -0.223 | -0.122 | < 0.001 |
| Archaeophyte | SH | 3 | -0.142 | -0.079 | 0.5 |
| Archaeophyte | HH | 22 | -0.440 | -0.317 | < 0.001 |
| Archaeophyte | BW | 54 | -0.018 | -0.038 | < 0.001 |
| Neophyte | SH | 21 | 0.240 | 0.076 | 0.005 |
| Neophyte | HH | 34 | 0.190 | 0.174 | 0.138 |
| Neophyte | BW | 40 | 0.204 | 0.077 | < 0.001 |

.
 Notes: Only species were included per group that showed a significant trend in the respective state and that had a status
 assigned. *n* is given for number of species with a significant trend that are included in each group. p-values indicate
 whether a group’s trend deviated from zero change according to Wilcoxon signed rank tests.

**(a)	(b)**
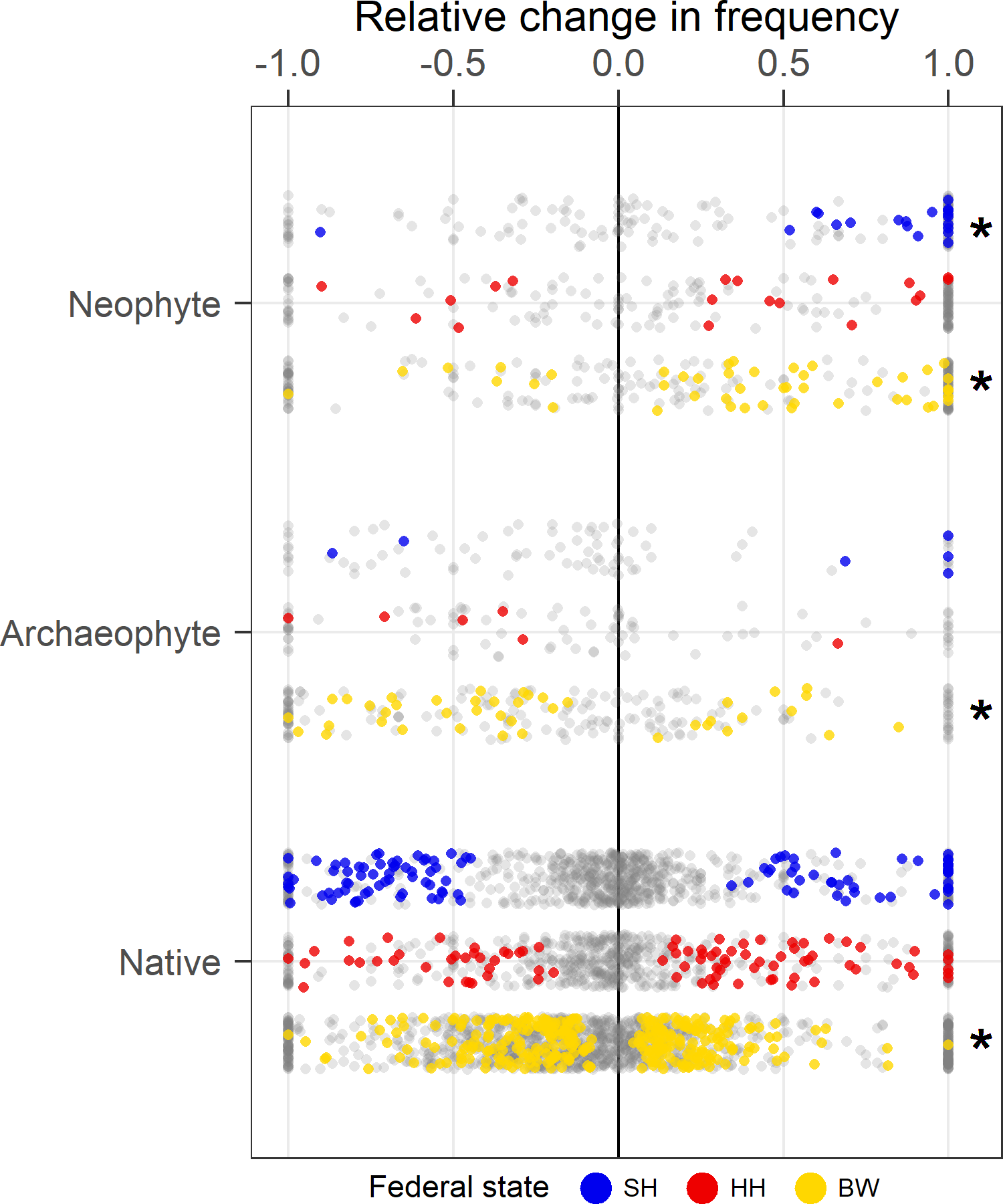

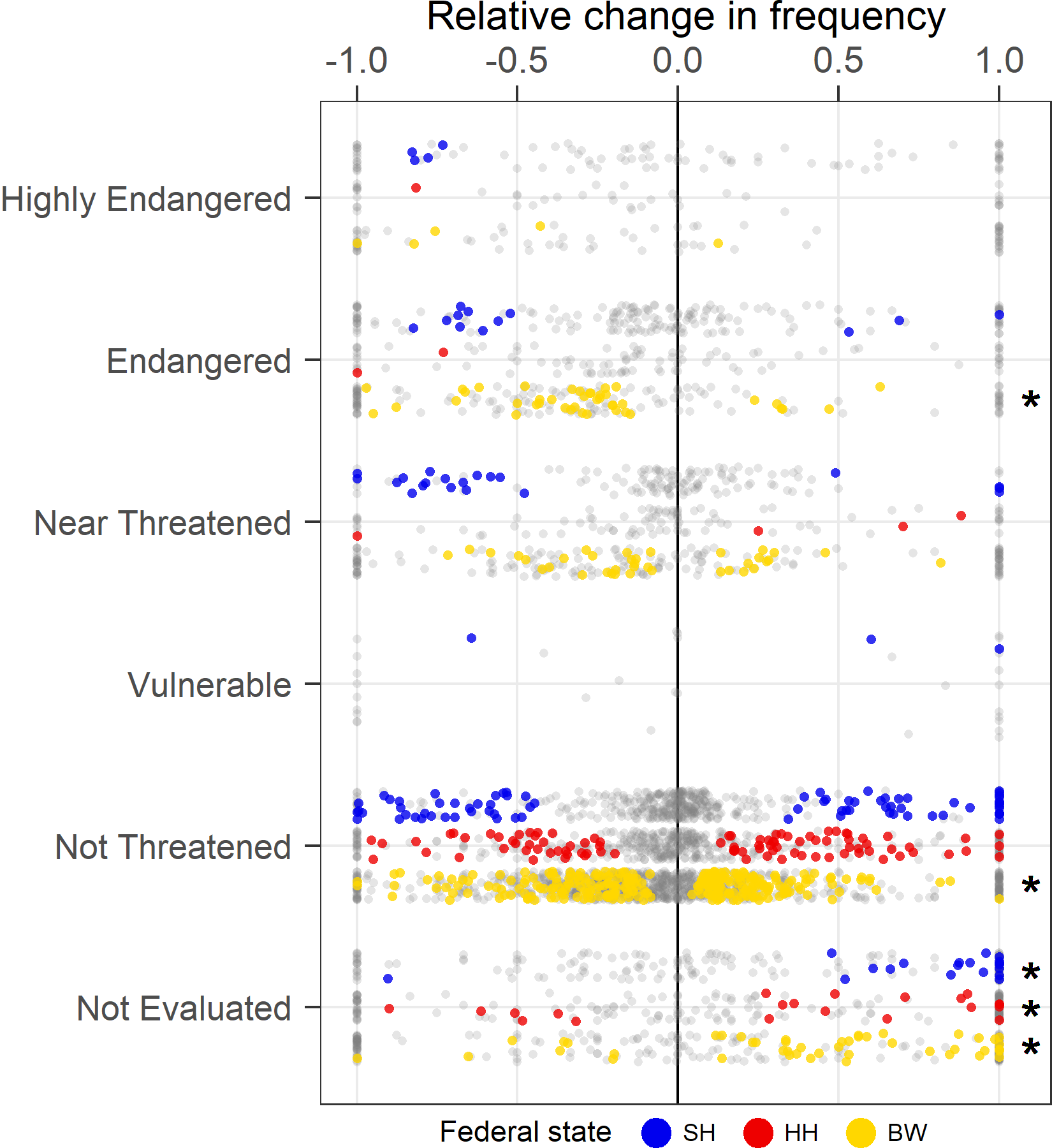

**Figure S1.11** Frequency trends grouped by states and **(a)** non-native and **(b)** Red List status. **(a)** Archaeophytes = non-natives introduced before 1492, neophytes = non-natives introduced after 1492. **(b)** There were no species with a significant trend of the Red List categories 0 (extinct or lost) or 1 (threatened with extinction). Significant trends are coloured, non-significant trends are displayed in grey. Asterisks indicate whether a group’s trend (only based on significant species trends) deviated from zero change according to Wilcoxon signed rank tests. Status according to German wide lists (Buttler et al., 2018; Klotz et al., 2002; Metzing et al., 2018; Wisskirchen & Haeupler, 1998).

**Table S1.7** Mean and Median Beals trends of species grouped by their Red List status and by state.

| **Red List status** | **State** | **n** | **Mean trend** | **Median trend** | **p** |
| --- | --- | --- | --- | --- | --- |
| Highly Endangered | SH | 11 | -0.126 | -0.120 | 0.001 |
| Highly Endangered | HH | 1 | -0.104 | -0.104 | 1 |
| Highly Endangered | BW | 29 | -0.045 | -0.030 | < 0.001 |
| Endangered | SH | 45 | -0.360 | -0.170 | < 0.001 |
| Endangered | BW | 132 | -0.151 | -0.071 | < 0.001 |
| Near Threatened | SH | 53 | -0.586 | -0.305 | < 0.001 |
| Near Threatened | HH | 12 | -0.069 | -0.152 | 0.092 |
| Near Threatened | BW | 119 | -0.447 | -0.223 | < 0.001 |
| Vulnerable | SH | 3 | -0.023 | 0.018 | 1 |
| Vulnerable | BW | 1 | -0.113 | -0.113 | 1 |
| Not Threatened | SH | 256 | -0.406 | -0.305 | < 0.001 |
| Not Threatened | HH | 270 | -0.089 | -0.209 | 0.005 |
| Not Threatened | BW | 486 | -0.147 | -0.129 | < 0.001 |
| Not Evaluated | SH | 21 | 0.244 | 0.080 | 0.011 |
| Not Evaluated | HH | 36 | 0.177 | 0.174 | 0.162 |
| Not Evaluated | BW | 42 | 0.153 | 0.065 | < 0.001 |

.
 Notes: Only species were included per group that showed a significant trend in the respective state and that had a status
 assigned. Not all states had species with significant trends of each Red List group. *n* is given for number of species with a
 significant trend that are included in each group. p-values indicate whether a group’s trend deviated from zero change
 according to Wilcoxon signed rank tests.

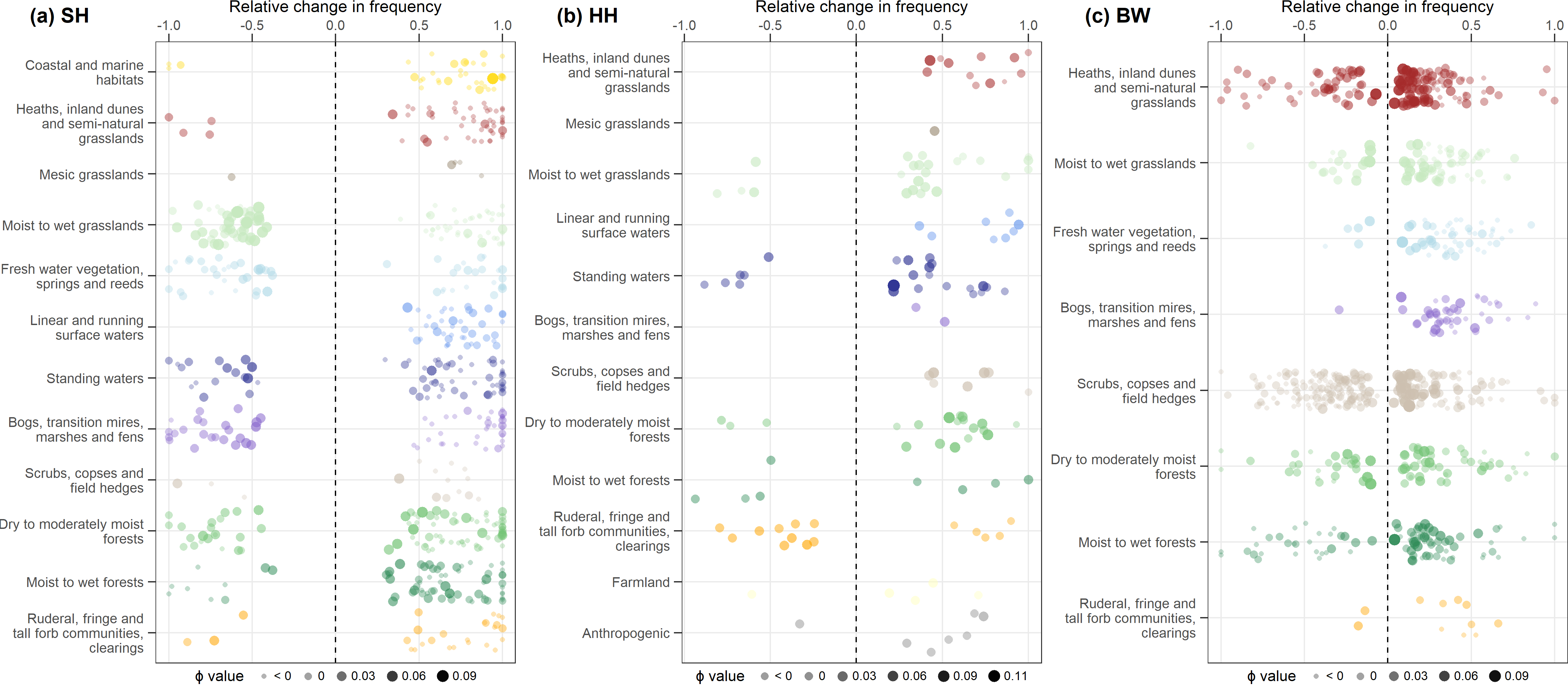

**Figure S1.12** Frequency trends within habitat types for **(a)** SH, **(b)** HH, and **(c)** BW. Species’ preferences to each habitat type (Φ value) are indicated by size and opacity. All Φ values below 0 are grouped into the category “< 0”. Only species are included that showed a significant trend. Note that not all habitat types were mapped in all states.

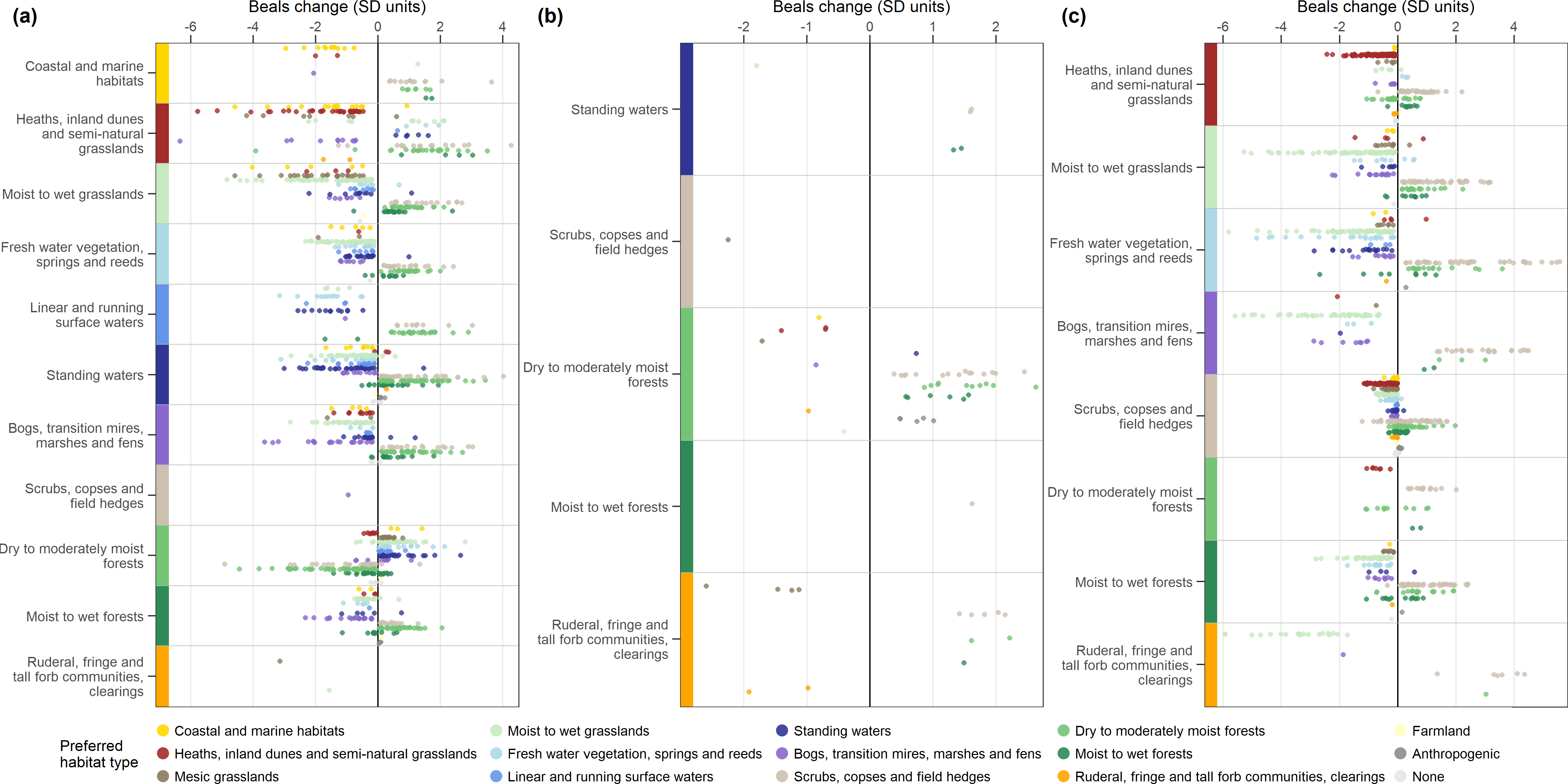

**Figure S1.13** Beals trends within habitat types, separately by the species’ preferred habitat type for **(a)** SH, **(b)** HH, and **(c)** BW. Colour bars on the left indicate the habitat type in which trends occurred (matches y-labels). Colours of points indicate the preferred habitat type of a species. Only species are included that showed a significant trend. Note that not all habitat types were mapped in all states.

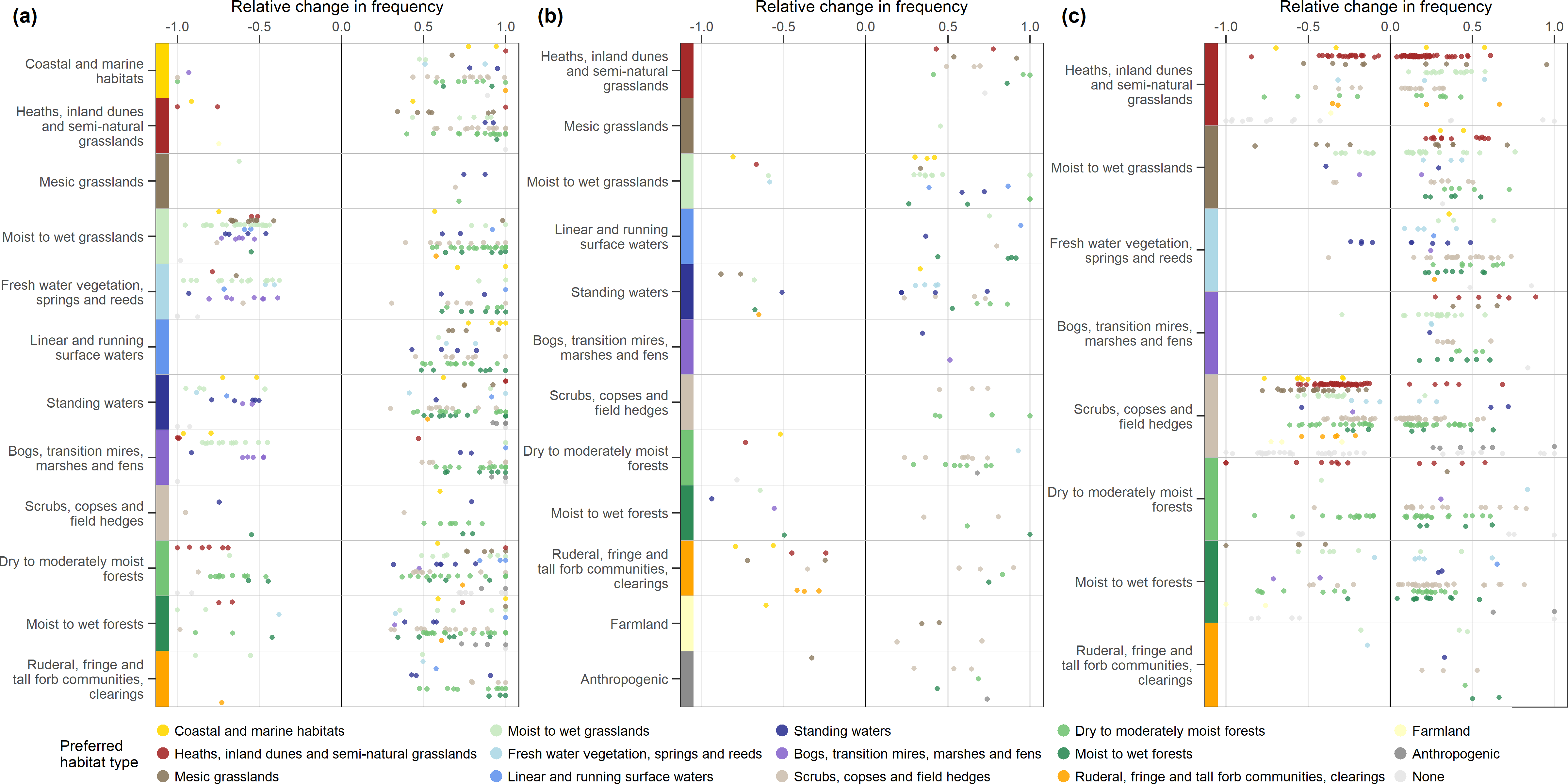

**Figure S1.14** Frequency trends within habitat types, separately by the species’ preferred habitat type for **(a)** SH, **(b)** HH, and **(c)** BW. Colour bars on the left indicate the habitat type in which trends occurred (matches y-labels). Colours of points indicate the preferred habitat type of a species. Only species are included that showed a significant trend. Note that not all habitat types were mapped in all states.
