## Appendix S2 for "Loss of characteristic species across German federal states detected by repeated mapping of protected habitats"

**Table S2** Beals trends of all species that showed a significant trend in at least one state.

| **Species** | **n (SH)** | **Beals trend (SH)** | **p (SH)** | **Beals trend scaled (SH)** | **n (HH)** | **Beals trend (HH)** | **p (HH)** | **Beals trend scaled (HH)** | **n (BW)** | **Beals trend (BW)** | **p (BW)** | **Beals trend scaled (BW)** | **Habitat type preferred** | **RL** | **NN** |
| --- | --- | --- | --- | --- | --- | --- | --- | --- | --- | --- | --- | --- | --- | --- | --- |
| *Festuca rubra* agg. | 2066 | -0.0189 | <.001 | -4.00 | 405 | -0.0165 | <.001 | -2.01 | 3200 | -0.0028 | <.001 | -1.28 | Coastal and marine habitats | Not Threatened | I |
| *Calluna vulgaris* | 1011 | -0.0183 | <.001 | -3.87 | 110 | -0.0071 | <.001 | -0.87 | 1834 | -0.0026 | <.001 | -1.21 | Bogs, transition mires, marshes and fens | Not Threatened | I |
| *Carex nigra* agg. | 1962 | -0.0181 | <.001 | -3.82 | 165 | -0.0037 | NA | NA | 2216 | -0.0048 | <.001 | -2.23 | Moist to wet grasslands | Not Threatened | I |
| *Eriophorum angustifolium* | 1150 | -0.0177 | <.001 | -3.74 | 48 | -0.0019 | NA | NA | 1249 | -0.0030 | <.001 | -1.37 | Bogs, transition mires, marshes and fens | Near Threatened | I |
| *Molinia caerulea* agg. | 2127 | -0.0168 | <.001 | -3.57 | 246 | -0.0041 | NA | NA | 2098 | -0.0031 | <.001 | -1.42 | Bogs, transition mires, marshes and fens | Not Threatened | I |
| *Erica tetralix* | 792 | -0.0152 | <.001 | -3.22 | 38 | -0.0035 | <.001 | -0.42 | 3 | 0.0000 | NA | NA | Bogs, transition mires, marshes and fens | Near Threatened | I |
| *Anthoxanthum odoratum* agg. | 1608 | -0.0143 | <.001 | -3.02 | 192 | -0.0090 | <.001 | -1.10 | 2466 | -0.0038 | <.001 | -1.77 | Moist to wet grasslands | Not Threatened | I |
| *Rumex acetosa* | 1890 | -0.0131 | NA | NA | 387 | -0.0164 | <.001 | -2.01 | 1376 | -0.0021 | <.001 | -0.95 | Mesic grasslands | Not Threatened | I |
| *Caltha palustris* | 1842 | -0.0130 | <.001 | -2.74 | 116 | -0.0007 | NA | NA | 6463 | -0.0080 | <.001 | -3.71 | Moist to wet grasslands | Near Threatened | I |
| *Hydrocotyle vulgaris* | 929 | -0.0129 | <.001 | -2.73 | 60 | -0.0017 | <.001 | -0.21 | 15 | 0.0000 | <.001 | -0.01 | Bogs, transition mires, marshes and fens | Not Threatened | I |
| *Lychnis flos-cuculi* | 1180 | -0.0127 | <.001 | -2.70 | 154 | -0.0056 | <.001 | -0.68 | 2890 | -0.0048 | <.001 | -2.23 | Moist to wet grasslands | Not Threatened | I |
| *Comarum palustre* | 1114 | -0.0127 | <.001 | -2.70 | 82 | -0.0018 | NA | NA | 648 | -0.0016 | <.001 | -0.72 | Bogs, transition mires, marshes and fens | Not Threatened | I |
| *Juncus articulatus* | 1290 | -0.0125 | <.001 | -2.64 | 176 | -0.0067 | <.001 | -0.82 | 1543 | -0.0027 | <.001 | -1.25 | Moist to wet grasslands | Not Threatened | I |
| *Cirsium palustre* | 2401 | -0.0124 | <.001 | -2.62 | 280 | -0.0057 | <.001 | -0.70 | 3916 | -0.0052 | <.001 | -2.41 | Moist to wet grasslands | Not Threatened | I |
| *Holcus lanatus* | 2922 | -0.0121 | NA | NA | 772 | -0.0266 | <.001 | -3.26 | 4185 | -0.0059 | <.001 | -2.76 | Moist to wet grasslands | Not Threatened | I |
| *Potentilla erecta* | 896 | -0.0111 | <.001 | -2.34 | 85 | -0.0032 | <.001 | -0.39 | 3204 | -0.0054 | <.001 | -2.49 | Moist to wet grasslands | Not Threatened | I |
| *Carex rostrata* | 1286 | -0.0111 | <.001 | -2.34 | 69 | -0.0011 | NA | NA | 1816 | -0.0036 | <.001 | -1.67 | Bogs, transition mires, marshes and fens | Not Threatened | I |
| *Luzula campestris* agg. | 978 | -0.0110 | <.001 | -2.33 | 134 | -0.0056 | <.001 | -0.69 | 1674 | -0.0024 | <.001 | -1.11 | Heaths, inland dunes and semi-natural grasslands | Not Threatened | I |
| *Lotus pedunculatus* | 1538 | -0.0109 | <.001 | -2.30 | 254 | -0.0087 | <.001 | -1.07 | 3692 | -0.0060 | <.001 | -2.79 | Moist to wet grasslands | Not Threatened | I |
| *Achillea millefolium* agg. | 1010 | -0.0105 | NA | NA | 346 | -0.0213 | <.001 | -2.61 | 4407 | -0.0043 | <.001 | -2.02 | Heaths, inland dunes and semi-natural grasslands | Not Threatened | I |
| *Cardamine pratensis* agg. | 1324 | -0.0100 | <.001 | -2.11 | 194 | -0.0050 | <.001 | -0.61 | 1424 | -0.0017 | <.001 | -0.77 | Moist to wet grasslands | Not Threatened | I |
| *Myosotis scorpioides* agg. | 2137 | -0.0100 | <.001 | -2.11 | 278 | -0.0036 | NA | NA | 4454 | -0.0076 | <.001 | -3.52 | Moist to wet grasslands | Not Threatened | I |
| *Rumex acetosella* s. l. | 866 | -0.0099 | <.001 | -2.09 | 221 | -0.0144 | <.001 | -1.76 | 965 | -0.0006 | <.001 | -0.27 | Heaths, inland dunes and semi-natural grasslands | Not Threatened | I |
| *Ranunculus acris* agg. | 1383 | -0.0099 | NA | NA | 324 | -0.0126 | <.001 | -1.54 | 2778 | -0.0042 | <.001 | -1.94 | Moist to wet grasslands | Not Threatened | I |
| *Betula pubescens* | 2632 | -0.0098 | NA | NA | 214 | 0.0025 | NA | NA | 644 | -0.0010 | <.001 | -0.45 | Bogs, transition mires, marshes and fens | Not Threatened | I |
| *Viola palustris* | 764 | -0.0097 | <.001 | -2.06 | 64 | -0.0015 | <.001 | -0.18 | 1558 | -0.0031 | <.001 | -1.42 | Moist to wet grasslands | Not Threatened | I |
| *Plantago lanceolata* | 1214 | -0.0095 | NA | NA | 382 | -0.0200 | <.001 | -2.44 | 2982 | -0.0043 | <.001 | -1.99 | Mesic grasslands | Not Threatened | A |
| *Hieracium pilosella* | 538 | -0.0095 | <.001 | -2.01 | 64 | -0.0058 | <.001 | -0.70 | 4046 | -0.0058 | <.001 | -2.70 | Heaths, inland dunes and semi-natural grasslands | Not Threatened | I |
| *Peucedanum palustre* | 1510 | -0.0095 | <.001 | -2.02 | 85 | -0.0012 | NA | NA | 220 | -0.0005 | <.001 | -0.22 | Bogs, transition mires, marshes and fens | Not Threatened | I |
| *Eriophorum vaginatum* | 752 | -0.0095 | <.001 | -2.01 | 22 | -0.0011 | NA | NA | 472 | -0.0013 | <.001 | -0.59 | Bogs, transition mires, marshes and fens | Near Threatened | I |
| *Deschampsia flexuosa* | 2633 | -0.0092 | <.001 | -1.95 | 350 | -0.0003 | NA | NA | 3024 | -0.0021 | <.001 | -0.96 | Dry to moderately moist forests | Not Threatened | I |
| *Agrostis capillaris* | 1528 | -0.0090 | NA | NA | 556 | -0.0194 | <.001 | -2.38 | 2474 | -0.0019 | <.001 | -0.90 | Mesic grasslands | Not Threatened | I |
| *Eleocharis palustris* agg. | 909 | -0.0089 | <.001 | -1.89 | 79 | -0.0023 | NA | NA | 605 | -0.0011 | <.001 | -0.52 | Standing waters | Not Threatened | I |
| *Trifolium repens* | 1138 | -0.0088 | NA | NA | 450 | -0.0208 | <.001 | -2.54 | 1492 | -0.0023 | <.001 | -1.04 | Mesic grasslands | Not Threatened | I |
| *Vaccinium oxycoccos* agg. | 434 | -0.0083 | <.001 | -1.76 | 14 | -0.0009 | NA | NA | 432 | -0.0012 | <.001 | -0.56 | Bogs, transition mires, marshes and fens | Endangered | I |
| *Ranunculus flammula* agg. | 964 | -0.0082 | <.001 | -1.73 | 158 | -0.0046 | <.001 | -0.56 | 1685 | -0.0029 | <.001 | -1.33 | Moist to wet grasslands | Not Threatened | I |
| *Typha latifolia* | 2367 | -0.0082 | <.001 | -1.74 | 236 | -0.0019 | NA | NA | 1736 | -0.0022 | <.001 | -1.01 | Standing waters | Not Threatened | I |
| *Carex panicea* | 486 | -0.0080 | <.001 | -1.70 | 25 | -0.0009 | <.001 | -0.11 | 2702 | -0.0056 | <.001 | -2.59 | Moist to wet grasslands | Near Threatened | I |
| *Carex arenaria* agg. | 492 | -0.0079 | <.001 | -1.66 | 36 | -0.0037 | <.001 | -0.46 | NA | NA | NA | NA | Coastal and marine habitats | Not Threatened | I |
| *Hypochaeris radicata* | 722 | -0.0078 | NA | NA | 125 | -0.0113 | <.001 | -1.38 | 444 | -0.0004 | <.001 | -0.18 | Mesic grasslands | Not Threatened | I |
| *Empetrum nigrum* agg. | 300 | -0.0075 | <.001 | -1.59 | 10 | -0.0009 | NA | NA | 24 | 0.0000 | <.001 | -0.02 | Coastal and marine habitats | Near Threatened | I |
| *Festuca ovina* agg. | 448 | -0.0073 | <.001 | -1.55 | 104 | -0.0073 | <.001 | -0.89 | 2152 | -0.0028 | <.001 | -1.30 | Heaths, inland dunes and semi-natural grasslands | Not Threatened | I |
| *Potentilla anserina* | 1218 | -0.0071 | <.001 | -1.49 | 360 | -0.0102 | <.001 | -1.25 | 585 | -0.0006 | <.001 | -0.28 | Coastal and marine habitats | Not Threatened | I |
| *Achillea ptarmica* agg. | 515 | -0.0070 | <.001 | -1.48 | 104 | -0.0043 | <.001 | -0.52 | 819 | -0.0012 | <.001 | -0.55 | Moist to wet grasslands | Not Threatened | I |
| *Carex disticha* | 951 | -0.0070 | <.001 | -1.48 | 77 | -0.0017 | NA | NA | 1583 | -0.0025 | <.001 | -1.18 | Moist to wet grasslands | Not Threatened | I |
| *Cerastium fontanum* agg. | 945 | -0.0069 | NA | NA | 291 | -0.0157 | <.001 | -1.92 | 462 | -0.0008 | <.001 | -0.39 | Mesic grasslands | Not Threatened | I |
| *Mentha aquatica* | 3102 | -0.0069 | <.001 | -1.47 | 138 | 0.0006 | NA | NA | 1196 | -0.0013 | <.001 | -0.58 | Fresh water vegetation, springs and reeds | Not Threatened | I |
| *Schoenoplectus lacustris* agg. | 652 | -0.0068 | <.001 | -1.44 | 40 | -0.0009 | NA | NA | 344 | -0.0005 | <.001 | -0.25 | Standing waters | Not Threatened | I |
| *Lycopus europaeus* | 2932 | -0.0068 | <.001 | -1.45 | 442 | 0.0001 | NA | NA | 1117 | -0.0008 | <.001 | -0.36 | Standing waters | Not Threatened | I |
| *Agrostis canina* agg. | 1095 | -0.0067 | <.001 | -1.41 | 99 | -0.0016 | NA | NA | 1076 | -0.0017 | <.001 | -0.78 | Moist to wet grasslands | Near Threatened | I |
| *Cirsium oleraceum* | 1890 | -0.0067 | <.001 | -1.42 | 178 | -0.0003 | NA | NA | 5762 | -0.0045 | <.001 | -2.11 | Fresh water vegetation, springs and reeds | Not Threatened | I |
| *Drosera rotundifolia* | 388 | -0.0065 | <.001 | -1.38 | 12 | -0.0008 | NA | NA | 238 | -0.0008 | <.001 | -0.36 | Bogs, transition mires, marshes and fens | Endangered | I |
| *Agrostis stolonifera* agg. | 2554 | -0.0063 | <.001 | -1.33 | 558 | -0.0115 | <.001 | -1.41 | 1227 | -0.0013 | <.001 | -0.61 | Coastal and marine habitats | Not Threatened | I |
| *Galium saxatile* | 738 | -0.0063 | <.001 | -1.33 | 112 | -0.0028 | <.001 | -0.34 | 973 | -0.0016 | <.001 | -0.76 | Heaths, inland dunes and semi-natural grasslands | Not Threatened | I |
| *Sparganium erectum* s. l. | 1236 | -0.0062 | <.001 | -1.31 | 116 | -0.0005 | NA | NA | 626 | -0.0008 | <.001 | -0.35 | Standing waters | Not Threatened | I |
| *Alisma plantago-aquatica* agg. | 826 | -0.0061 | <.001 | -1.29 | 134 | -0.0032 | NA | NA | 513 | -0.0007 | <.001 | -0.34 | Standing waters | Not Threatened | I |
| *Glyceria maxima* | 1644 | -0.0060 | <.001 | -1.28 | 532 | -0.0035 | NA | NA | 693 | -0.0006 | <.001 | -0.26 | Linear and running surface waters | Not Threatened | I |
| *Epilobium palustre* | 763 | -0.0060 | <.001 | -1.28 | 79 | -0.0015 | NA | NA | 790 | -0.0013 | <.001 | -0.59 | Bogs, transition mires, marshes and fens | Near Threatened | I |
| *Menyanthes trifoliata* | 396 | -0.0060 | <.001 | -1.27 | 13 | -0.0002 | NA | NA | 584 | -0.0015 | <.001 | -0.68 | Bogs, transition mires, marshes and fens | Endangered | I |
| *Juncus effusus* | 5956 | -0.0059 | NA | NA | 814 | -0.0085 | NA | NA | 6308 | -0.0068 | <.001 | -3.17 | Moist to wet grasslands | Not Threatened | I |
| *Berula erecta* | 1373 | -0.0059 | <.001 | -1.24 | 41 | 0.0005 | NA | NA | 374 | -0.0001 | NA | NA | Linear and running surface waters | Not Threatened | I |
| *Filipendula ulmaria* | 3089 | -0.0059 | <.001 | -1.24 | 370 | 0.0030 | NA | NA | 11815 | -0.0071 | <.001 | -3.31 | Moist to wet grasslands | Not Threatened | I |
| *Equisetum fluviatile* | 1386 | -0.0058 | <.001 | -1.24 | 111 | -0.0010 | NA | NA | 790 | -0.0012 | <.001 | -0.55 | Moist to wet grasslands | Not Threatened | I |
| *Alopecurus geniculatus* | 760 | -0.0057 | <.001 | -1.20 | 115 | -0.0055 | <.001 | -0.67 | 80 | -0.0001 | <.001 | -0.06 | Moist to wet grasslands | Not Threatened | I |
| *Galium palustre* agg. | 3112 | -0.0057 | <.001 | -1.20 | 248 | -0.0024 | NA | NA | 2848 | -0.0030 | <.001 | -1.40 | Moist to wet grasslands | Not Threatened | I |
| *Carex canescens* agg. | 803 | -0.0056 | <.001 | -1.18 | 64 | -0.0009 | NA | NA | 616 | -0.0012 | <.001 | -0.54 | Bogs, transition mires, marshes and fens | Not Threatened | I |
| *Andromeda polifolia* | 306 | -0.0056 | <.001 | -1.19 | 11 | -0.0006 | NA | NA | 157 | -0.0005 | <.001 | -0.21 | Bogs, transition mires, marshes and fens | Endangered | I |
| *Lysimachia vulgaris* | 3717 | -0.0056 | NA | NA | 428 | 0.0038 | NA | NA | 3284 | -0.0027 | <.001 | -1.27 | Fresh water vegetation, springs and reeds | Not Threatened | I |
| *Vicia cracca* agg. | 764 | -0.0055 | <.001 | -1.16 | 306 | -0.0132 | <.001 | -1.61 | 2634 | -0.0019 | <.001 | -0.87 | Mesic grasslands | Not Threatened | I |
| *Lathyrus pratensis* | 986 | -0.0055 | <.001 | -1.16 | 219 | -0.0086 | <.001 | -1.05 | 2433 | -0.0029 | <.001 | -1.33 | Moist to wet grasslands | Not Threatened | I |
| *Typha angustifolia* | 754 | -0.0055 | <.001 | -1.17 | 37 | -0.0006 | NA | NA | 178 | -0.0003 | <.001 | -0.14 | Standing waters | Not Threatened | I |
| *Lotus corniculatus* agg. | 507 | -0.0054 | <.001 | -1.14 | 84 | -0.0059 | <.001 | -0.72 | 4168 | -0.0048 | <.001 | -2.22 | Heaths, inland dunes and semi-natural grasslands | Not Threatened | I |
| *Geum rivale* | 1783 | -0.0054 | <.001 | -1.14 | 30 | 0.0008 | NA | NA | 3338 | -0.0029 | <.001 | -1.34 | Moist to wet grasslands | Not Threatened | I |
| *Primula elatior* | 1262 | -0.0053 | <.001 | -1.12 | 12 | 0.0009 | NA | NA | 3228 | -0.0005 | NA | NA | Moist to wet forests | Not Threatened | I |
| *Jasione montana* | 238 | -0.0052 | <.001 | -1.11 | 24 | -0.0035 | <.001 | -0.42 | 120 | -0.0002 | <.001 | -0.10 | Heaths, inland dunes and semi-natural grasslands | Not Threatened | I |
| *Salix repens* agg. | 256 | -0.0052 | <.001 | -1.10 | 7 | -0.0011 | NA | NA | 107 | -0.0003 | <.001 | -0.14 | Coastal and marine habitats | Not Threatened | I |
| *Myrica gale* | 553 | -0.0052 | <.001 | -1.10 | 22 | -0.0005 | NA | NA | NA | NA | NA | NA | Bogs, transition mires, marshes and fens | Endangered | I |
| *Juncus conglomeratus* | 524 | -0.0051 | <.001 | -1.09 | 82 | -0.0032 | <.001 | -0.39 | 1048 | -0.0017 | <.001 | -0.79 | Moist to wet grasslands | Not Threatened | I |
| *Dactylorhiza majalis* agg. | 294 | -0.0051 | <.001 | -1.08 | 16 | -0.0007 | <.001 | -0.09 | 1059 | -0.0023 | <.001 | -1.04 | Moist to wet grasslands | Near Threatened | I |
| *Campanula rotundifolia* agg. | 372 | -0.0050 | <.001 | -1.06 | 37 | -0.0021 | <.001 | -0.26 | 4148 | -0.0046 | <.001 | -2.12 | Heaths, inland dunes and semi-natural grasslands | Not Threatened | I |
| *Triglochin palustris* | 276 | -0.0050 | <.001 | -1.06 | 7 | -0.0003 | NA | NA | 46 | -0.0001 | <.001 | -0.05 | Coastal and marine habitats | Endangered | I |
| *Angelica sylvestris* | 1663 | -0.0050 | <.001 | -1.06 | 96 | 0.0003 | NA | NA | 6387 | -0.0041 | <.001 | -1.92 | Moist to wet grasslands | Not Threatened | I |
| *Crepis paludosa* | 961 | -0.0050 | <.001 | -1.05 | 50 | 0.0010 | NA | NA | 2080 | -0.0028 | <.001 | -1.31 | Moist to wet grasslands | Not Threatened | I |
| *Carex leporina* | 490 | -0.0049 | <.001 | -1.03 | 107 | -0.0048 | <.001 | -0.58 | 982 | -0.0016 | <.001 | -0.72 | Moist to wet grasslands | Not Threatened | I |
| *Poa pratensis* agg. | 716 | -0.0048 | NA | NA | 250 | -0.0091 | <.001 | -1.11 | 1888 | -0.0015 | <.001 | -0.72 | Mesic grasslands | Not Threatened | I |
| *Persicaria amphibia* | 1272 | -0.0048 | <.001 | -1.02 | 250 | -0.0067 | <.001 | -0.82 | 493 | -0.0006 | <.001 | -0.30 | Standing waters | Not Threatened | I |
| *Carex acuta* agg. | 1598 | -0.0048 | <.001 | -1.02 | 366 | -0.0019 | NA | NA | 2286 | -0.0027 | <.001 | -1.26 | Moist to wet grasslands | Not Threatened | I |
| *Rumex hydrolapathum* | 810 | -0.0048 | <.001 | -1.01 | 189 | -0.0012 | NA | NA | 92 | -0.0001 | <.001 | -0.04 | Linear and running surface waters | Not Threatened | I |
| *Carex hirta* | 883 | -0.0047 | NA | NA | 391 | -0.0126 | <.001 | -1.54 | 1818 | -0.0021 | <.001 | -0.99 | Moist to wet grasslands | Not Threatened | I |
| *Festuca pratensis* s. l. | 553 | -0.0046 | <.001 | -0.97 | 191 | -0.0074 | <.001 | -0.91 | 714 | -0.0010 | <.001 | -0.45 | Moist to wet grasslands | Not Threatened | I |
| *Lythrum salicaria* | 2042 | -0.0046 | <.001 | -0.98 | 394 | -0.0012 | NA | NA | 5010 | -0.0038 | <.001 | -1.77 | Fresh water vegetation, springs and reeds | Not Threatened | I |
| *Hypericum perforatum* | 468 | -0.0045 | NA | NA | 320 | -0.0161 | <.001 | -1.96 | 6006 | -0.0018 | <.001 | -0.85 | Heaths, inland dunes and semi-natural grasslands | Not Threatened | I |
| *Cynosurus cristatus* | 602 | -0.0045 | NA | NA | 61 | -0.0033 | <.001 | -0.40 | 902 | -0.0014 | <.001 | -0.66 | Mesic grasslands | Not Threatened | I |
| *Trifolium pratense* | 587 | -0.0044 | NA | NA | 239 | -0.0138 | <.001 | -1.69 | 2798 | -0.0042 | <.001 | -1.94 | Moist to wet grasslands | Not Threatened | I |
| *Rumex crispus* | 876 | -0.0044 | NA | NA | 260 | -0.0126 | <.001 | -1.54 | 382 | -0.0004 | <.001 | -0.20 | Mesic grasslands | Not Threatened | I |
| *Bolboschoenus maritimus* agg. | 392 | -0.0044 | <.001 | -0.94 | 9 | -0.0002 | NA | NA | 25 | 0.0000 | NA | NA | Coastal and marine habitats | Not Threatened | I |
| *Nardus stricta* | 264 | -0.0043 | <.001 | -0.92 | 35 | -0.0031 | <.001 | -0.38 | 1530 | -0.0031 | <.001 | -1.42 | Heaths, inland dunes and semi-natural grasslands | Near Threatened | I |
| *Succisa pratensis* | 216 | -0.0043 | <.001 | -0.92 | 11 | -0.0006 | <.001 | -0.07 | 1353 | -0.0027 | <.001 | -1.23 | Moist to wet grasslands | Near Threatened | I |
| *Juncus compressus* agg. | 248 | -0.0043 | <.001 | -0.91 | 6 | -0.0004 | NA | NA | 78 | -0.0001 | <.001 | -0.06 | Coastal and marine habitats | Not Threatened | I |
| *Trifolium arvense* | 268 | -0.0042 | <.001 | -0.90 | 59 | -0.0071 | <.001 | -0.87 | 107 | -0.0002 | <.001 | -0.08 | Mesic grasslands | Not Threatened | I |
| *Galium uliginosum* | 402 | -0.0042 | <.001 | -0.88 | 35 | -0.0009 | <.001 | -0.11 | 1536 | -0.0026 | <.001 | -1.22 | Moist to wet grasslands | Not Threatened | I |
| *Eupatorium cannabinum* | 1694 | -0.0042 | <.001 | -0.88 | 82 | 0.0004 | NA | NA | 2056 | 0.0000 | NA | NA | Fresh water vegetation, springs and reeds | Not Threatened | I |
| *Phragmites australis* | 3650 | -0.0042 | NA | NA | 504 | 0.0006 | NA | NA | 4628 | -0.0025 | <.001 | -1.15 | Fresh water vegetation, springs and reeds | Not Threatened | I |
| *Frangula alnus* | 2183 | -0.0042 | NA | NA | 284 | 0.0025 | NA | NA | 3332 | -0.0021 | <.001 | -0.98 | Moist to wet forests | Not Threatened | I |
| *Scorzoneroides autumnalis* | 436 | -0.0041 | NA | NA | 128 | -0.0080 | <.001 | -0.98 | 302 | -0.0004 | <.001 | -0.19 | Mesic grasslands | Not Threatened | I |
| *Bellis perennis* | 502 | -0.0041 | NA | NA | 109 | -0.0047 | <.001 | -0.57 | 398 | -0.0008 | <.001 | -0.38 | Mesic grasslands | Not Threatened | A |
| *Stellaria graminea* | 728 | -0.0040 | NA | NA | 186 | -0.0071 | <.001 | -0.87 | 797 | -0.0009 | <.001 | -0.39 | Mesic grasslands | Not Threatened | I |
| *Equisetum palustre* | 1162 | -0.0040 | <.001 | -0.85 | 263 | -0.0049 | <.001 | -0.60 | 1643 | -0.0023 | <.001 | -1.08 | Moist to wet grasslands | Not Threatened | I |
| *Prunella vulgaris* | 378 | -0.0040 | <.001 | -0.85 | 108 | -0.0049 | <.001 | -0.60 | 1226 | -0.0018 | <.001 | -0.83 | Heaths, inland dunes and semi-natural grasslands | Not Threatened | I |
| *Corynephorus canescens* | 167 | -0.0039 | <.001 | -0.82 | 21 | -0.0038 | <.001 | -0.47 | 39 | -0.0001 | NA | NA | Coastal and marine habitats | Not Threatened | I |
| *Ranunculus repens* | 3557 | -0.0037 | NA | NA | 763 | -0.0127 | <.001 | -1.55 | 4097 | -0.0037 | <.001 | -1.72 | Moist to wet grasslands | Not Threatened | I |
| *Juncus bufonius* agg. | 308 | -0.0036 | <.001 | -0.77 | 67 | -0.0028 | <.001 | -0.35 | 130 | -0.0003 | <.001 | -0.12 | Standing waters | Not Threatened | I |
| *Valeriana dioica* agg. | 306 | -0.0034 | <.001 | -0.73 | 13 | 0.0002 | NA | NA | 2259 | -0.0037 | <.001 | -1.73 | Moist to wet grasslands | Not Threatened | I |
| *Tanacetum vulgare* | 371 | -0.0033 | NA | NA | 245 | -0.0146 | <.001 | -1.79 | 252 | -0.0001 | <.001 | -0.05 | Ruderal, fringe and tall forb communities, clearings | Not Threatened | A |
| *Galium mollugo* agg. | 426 | -0.0033 | NA | NA | 114 | -0.0056 | <.001 | -0.68 | 7666 | -0.0033 | <.001 | -1.52 | Scrubs, copses and field hedges | Not Threatened | I |
| *Rhynchospora alba* | 220 | -0.0033 | <.001 | -0.71 | 10 | -0.0009 | NA | NA | 72 | -0.0002 | <.001 | -0.10 | Bogs, transition mires, marshes and fens | Endangered | I |
| *Epilobium hirsutum* | 1832 | -0.0033 | <.001 | -0.70 | 340 | -0.0007 | NA | NA | 2757 | -0.0013 | <.001 | -0.60 | Fresh water vegetation, springs and reeds | Not Threatened | I |
| *Lysimachia nummularia* | 792 | -0.0033 | <.001 | -0.70 | 150 | 0.0006 | NA | NA | 2292 | -0.0019 | <.001 | -0.87 | Moist to wet grasslands | Not Threatened | I |
| *Taraxacum* sect. *Ruderalia* | 1063 | -0.0032 | NA | NA | 430 | -0.0119 | <.001 | -1.46 | 1837 | -0.0019 | <.001 | -0.89 | Mesic grasslands | Not Threatened | I |
| *Carex echinata* | 201 | -0.0032 | <.001 | -0.67 | 21 | -0.0009 | <.001 | -0.11 | 1606 | -0.0035 | <.001 | -1.63 | Moist to wet grasslands | Not Threatened | I |
| *Genista anglica* | 155 | -0.0032 | <.001 | -0.68 | 6 | -0.0005 | NA | NA | 5 | 0.0000 | NA | NA | Heaths, inland dunes and semi-natural grasslands | Endangered | I |
| *Sanicula europaea* | 348 | -0.0032 | <.001 | -0.67 | 2 | 0.0006 | NA | NA | 456 | 0.0000 | NA | NA | Dry to moderately moist forests | Not Threatened | I |
| *Juncus squarrosus* | 176 | -0.0031 | <.001 | -0.67 | 24 | -0.0018 | <.001 | -0.21 | 284 | -0.0007 | <.001 | -0.32 | Heaths, inland dunes and semi-natural grasslands | Near Threatened | I |
| *Carex flava* agg. | 152 | -0.0031 | <.001 | -0.65 | 16 | -0.0008 | <.001 | -0.09 | 998 | -0.0022 | <.001 | -1.03 | Moist to wet grasslands | Not Threatened | I |
| *Ammophila arenaria* | 217 | -0.0031 | <.001 | -0.67 | NA | NA | NA | NA | NA | NA | NA | NA | Coastal and marine habitats | Not Threatened | I |
| *Armeria maritima* s. l. | 155 | -0.0030 | <.001 | -0.64 | 1 | -0.0003 | NA | NA | 2 | 0.0000 | NA | NA | Coastal and marine habitats | Near Threatened | I |
| *Aira praecox* | 151 | -0.0029 | <.001 | -0.61 | 14 | -0.0018 | <.001 | -0.22 | 6 | 0.0000 | NA | NA | Coastal and marine habitats | Near Threatened | I |
| *Lysimachia thyrsiflora* | 442 | -0.0029 | <.001 | -0.61 | 38 | 0.0000 | NA | NA | 86 | -0.0002 | <.001 | -0.07 | Standing waters | Near Threatened | I |
| *Carex paniculata* | 1720 | -0.0029 | NA | NA | 76 | 0.0009 | NA | NA | 568 | -0.0007 | <.001 | -0.33 | Standing waters | Not Threatened | I |
| *Calamagrostis canescens* agg. | 3466 | -0.0029 | NA | NA | 207 | 0.0009 | NA | NA | 151 | -0.0002 | <.001 | -0.09 | Bogs, transition mires, marshes and fens | Not Threatened | I |
| *Bromus hordeaceus* agg. | 326 | -0.0028 | NA | NA | 110 | -0.0063 | <.001 | -0.77 | 230 | -0.0003 | <.001 | -0.13 | Mesic grasslands | Not Threatened | I |
| *Galium verum* agg. | 172 | -0.0028 | <.001 | -0.60 | 10 | -0.0010 | NA | NA | 4762 | -0.0050 | <.001 | -2.34 | Heaths, inland dunes and semi-natural grasslands | Not Threatened | I |
| *Potamogeton natans* | 669 | -0.0028 | <.001 | -0.60 | 44 | -0.0004 | NA | NA | 610 | -0.0007 | <.001 | -0.34 | Standing waters | Not Threatened | I |
| *Pulmonaria officinalis* agg. | 381 | -0.0028 | <.001 | -0.60 | 1 | 0.0001 | NA | NA | 974 | 0.0001 | NA | NA | Dry to moderately moist forests | Not Threatened | I |
| *Nuphar lutea* | 559 | -0.0028 | <.001 | -0.59 | 66 | 0.0007 | NA | NA | 244 | -0.0001 | NA | NA | Standing waters | Not Threatened | I |
| *Cytisus scoparius* | 204 | -0.0027 | <.001 | -0.56 | 105 | -0.0026 | <.001 | -0.32 | 1189 | -0.0001 | NA | NA | Heaths, inland dunes and semi-natural grasslands | Not Threatened | I |
| *Bidens tripartita* | 228 | -0.0027 | <.001 | -0.58 | 73 | -0.0019 | <.001 | -0.23 | 26 | -0.0001 | <.001 | -0.03 | Standing waters | Not Threatened | I |
| *Juncus filiformis* | 180 | -0.0027 | <.001 | -0.58 | 17 | -0.0007 | <.001 | -0.08 | 272 | -0.0005 | <.001 | -0.25 | Moist to wet grasslands | Near Threatened | I |
| *Glaux maritima* | 234 | -0.0027 | <.001 | -0.56 | NA | NA | NA | NA | NA | NA | NA | NA | Coastal and marine habitats | Not Threatened | I |
| *Ornithopus perpusillus* | 150 | -0.0026 | <.001 | -0.55 | 31 | -0.0035 | <.001 | -0.42 | 15 | 0.0000 | NA | NA | Heaths, inland dunes and semi-natural grasslands | Not Threatened | I |
| *Phyteuma spicatum* agg. | 348 | -0.0026 | <.001 | -0.55 | 4 | 0.0004 | NA | NA | 1438 | -0.0003 | NA | NA | Dry to moderately moist forests | Not Threatened | I |
| *Triglochin maritima* | 194 | -0.0026 | <.001 | -0.54 | NA | NA | NA | NA | NA | NA | NA | NA | Coastal and marine habitats | Near Threatened | I |
| *Cirsium arvense* | 1800 | -0.0025 | NA | NA | 741 | -0.0253 | <.001 | -3.09 | 3304 | -0.0013 | <.001 | -0.62 | Mesic grasslands | Not Threatened | I |
| *Lolium perenne* | 705 | -0.0025 | NA | NA | 434 | -0.0161 | <.001 | -1.97 | 483 | -0.0003 | <.001 | -0.14 | Mesic grasslands | Not Threatened | I |
| *Plantago major* agg. | 372 | -0.0025 | NA | NA | 328 | -0.0102 | <.001 | -1.24 | 246 | -0.0003 | <.001 | -0.14 | Mesic grasslands | Not Threatened | I |
| *Stachys palustris* | 802 | -0.0025 | <.001 | -0.52 | 214 | -0.0024 | <.001 | -0.29 | 741 | -0.0004 | <.001 | -0.18 | Fresh water vegetation, springs and reeds | Not Threatened | I |
| *Bidens cernua* | 367 | -0.0025 | <.001 | -0.53 | 58 | -0.0017 | <.001 | -0.21 | 14 | 0.0000 | NA | NA | Standing waters | Not Threatened | I |
| *Sagina procumbens* | 135 | -0.0025 | <.001 | -0.54 | 14 | -0.0017 | <.001 | -0.21 | 7 | 0.0000 | NA | NA | None | Not Threatened | I |
| *Hieracium umbellatum* | 93 | -0.0025 | <.001 | -0.53 | 20 | -0.0010 | <.001 | -0.12 | 369 | -0.0002 | <.001 | -0.10 | Coastal and marine habitats | Not Threatened | I |
| *Mercurialis perennis* agg. | 836 | -0.0025 | NA | NA | 14 | 0.0009 | <.001 | 0.11 | 3572 | 0.0005 | NA | NA | Dry to moderately moist forests | Not Threatened | I |
| *Linaria vulgaris* agg. | 271 | -0.0024 | <.001 | -0.51 | 107 | -0.0050 | <.001 | -0.61 | 1532 | -0.0006 | <.001 | -0.26 | Scrubs, copses and field hedges | Not Threatened | I |
| *Danthonia decumbens* | 150 | -0.0024 | <.001 | -0.50 | 26 | -0.0023 | <.001 | -0.28 | 937 | -0.0012 | <.001 | -0.55 | Heaths, inland dunes and semi-natural grasslands | Near Threatened | I |
| *Ranunculus sceleratus* | 569 | -0.0024 | <.001 | -0.50 | 72 | -0.0023 | <.001 | -0.28 | 74 | -0.0001 | NA | NA | Standing waters | Not Threatened | I |
| *Knautia arvensis* agg. | 176 | -0.0024 | NA | NA | 10 | -0.0007 | NA | NA | 4754 | -0.0035 | <.001 | -1.62 | Heaths, inland dunes and semi-natural grasslands | Not Threatened | I |
| *Rhinanthus serotinus* s. l. | 168 | -0.0024 | <.001 | -0.52 | 25 | -0.0007 | NA | NA | 27 | 0.0000 | <.001 | -0.02 | Mesic grasslands | Endangered | I |
| *Narthecium ossifragum* | 114 | -0.0024 | <.001 | -0.51 | 6 | -0.0005 | NA | NA | NA | NA | NA | NA | Bogs, transition mires, marshes and fens | Endangered | I |
| *Cicuta virosa* | 202 | -0.0024 | <.001 | -0.50 | 1 | 0.0000 | NA | NA | 18 | 0.0000 | <.001 | -0.01 | Standing waters | Near Threatened | I |
| *Cirsium vulgare* | 464 | -0.0023 | NA | NA | 214 | -0.0105 | <.001 | -1.28 | 1370 | -0.0006 | <.001 | -0.26 | Mesic grasslands | Not Threatened | I |
| *Teesdalia nudicaulis* | 106 | -0.0023 | <.001 | -0.49 | 3 | -0.0005 | NA | NA | 16 | -0.0001 | NA | NA | Heaths, inland dunes and semi-natural grasslands | Not Threatened | I |
| *Persicaria hydropiper* | 650 | -0.0022 | <.001 | -0.47 | 226 | -0.0050 | <.001 | -0.61 | 226 | 0.0001 | NA | NA | Linear and running surface waters | Not Threatened | I |
| *Sedum acre* | 130 | -0.0022 | <.001 | -0.46 | 20 | -0.0024 | <.001 | -0.30 | 745 | -0.0012 | <.001 | -0.57 | Heaths, inland dunes and semi-natural grasslands | Not Threatened | I |
| *Epilobium parviflorum* | 487 | -0.0022 | <.001 | -0.46 | 55 | -0.0006 | NA | NA | 978 | -0.0007 | <.001 | -0.33 | Fresh water vegetation, springs and reeds | Not Threatened | I |
| *Briza media* | 78 | -0.0022 | <.001 | -0.46 | 4 | -0.0004 | NA | NA | 3954 | -0.0061 | <.001 | -2.83 | Heaths, inland dunes and semi-natural grasslands | Not Threatened | I |
| *Dactylorhiza maculata* agg. | 125 | -0.0022 | <.001 | -0.47 | 16 | -0.0004 | NA | NA | 777 | -0.0016 | <.001 | -0.74 | Moist to wet grasslands | Near Threatened | I |
| *Carex vulpina* agg. | 364 | -0.0022 | <.001 | -0.48 | 14 | -0.0004 | NA | NA | 324 | -0.0005 | <.001 | -0.22 | Coastal and marine habitats | Not Threatened | I |
| *Leymus arenarius* | 164 | -0.0022 | <.001 | -0.46 | 6 | -0.0003 | NA | NA | NA | NA | NA | NA | Coastal and marine habitats | Not Threatened | I |
| *Carex elata* | 1060 | -0.0022 | <.001 | -0.47 | 15 | 0.0000 | NA | NA | 983 | -0.0010 | <.001 | -0.44 | Fresh water vegetation, springs and reeds | Not Threatened | I |
| *Tripolium pannonicum* | 216 | -0.0022 | <.001 | -0.47 | 1 | 0.0000 | NA | NA | NA | NA | NA | NA | Coastal and marine habitats | Not Threatened | I |
| *Platanthera chlorantha* | 172 | -0.0022 | <.001 | -0.46 | 6 | 0.0001 | NA | NA | 340 | -0.0005 | <.001 | -0.25 | Heaths, inland dunes and semi-natural grasslands | Endangered | I |
| *Viola tricolor* agg. | 126 | -0.0021 | <.001 | -0.44 | 44 | -0.0044 | <.001 | -0.54 | 54 | -0.0001 | <.001 | -0.04 | Coastal and marine habitats | Not Threatened | NA |
| *Veronica chamaedrys* agg. | 404 | -0.0021 | NA | NA | 124 | -0.0028 | <.001 | -0.34 | 1398 | -0.0012 | <.001 | -0.54 | Mesic grasslands | Not Threatened | I |
| *Trichophorum cespitosum* agg. | 106 | -0.0021 | <.001 | -0.44 | 8 | -0.0010 | NA | NA | 76 | -0.0002 | <.001 | -0.11 | Bogs, transition mires, marshes and fens | Endangered | I |
| *Sonchus palustris* | 294 | -0.0021 | <.001 | -0.44 | 2 | -0.0001 | NA | NA | 1 | 0.0000 | NA | NA | Fresh water vegetation, springs and reeds | Not Threatened | I |
| *Plantago maritima* agg. | 192 | -0.0021 | <.001 | -0.44 | NA | NA | NA | NA | NA | NA | NA | NA | Coastal and marine habitats | Not Threatened | I |
| *Phleum pratense* agg. | 359 | -0.0020 | NA | NA | 263 | -0.0118 | <.001 | -1.44 | 1545 | -0.0015 | <.001 | -0.67 | Mesic grasslands | Not Threatened | I |
| *Calamagrostis epigejos* | 502 | -0.0020 | NA | NA | 476 | -0.0117 | <.001 | -1.43 | 1143 | -0.0003 | <.001 | -0.12 | Ruderal, fringe and tall forb communities, clearings | Not Threatened | I |
| *Daucus carota* | 191 | -0.0020 | NA | NA | 50 | -0.0045 | <.001 | -0.54 | 2854 | -0.0024 | <.001 | -1.10 | Heaths, inland dunes and semi-natural grasslands | Not Threatened | I |
| *Senecio aquaticus* agg. | 149 | -0.0020 | <.001 | -0.43 | 53 | -0.0009 | NA | NA | 1142 | -0.0019 | <.001 | -0.90 | Moist to wet grasslands | Near Threatened | I |
| *Ranunculus lingua* | 148 | -0.0020 | <.001 | -0.42 | 12 | 0.0000 | NA | NA | 46 | -0.0001 | <.001 | -0.03 | Standing waters | Endangered | I |
| *Nymphaea alba* | 477 | -0.0020 | <.001 | -0.43 | 56 | 0.0002 | NA | NA | 322 | -0.0004 | <.001 | -0.16 | Standing waters | Not Threatened | I |
| *Arum maculatum* agg. | 664 | -0.0020 | NA | NA | NA | NA | NA | NA | 2696 | 0.0010 | <.001 | 0.47 | Dry to moderately moist forests | Not Threatened | I |
| *Alopecurus pratensis* agg. | 1008 | -0.0019 | NA | NA | 326 | -0.0127 | <.001 | -1.55 | 2845 | -0.0029 | <.001 | -1.33 | Moist to wet grasslands | Not Threatened | I |
| *Trifolium campestre* | 138 | -0.0019 | NA | NA | 21 | -0.0028 | <.001 | -0.34 | 630 | -0.0009 | <.001 | -0.41 | Heaths, inland dunes and semi-natural grasslands | Not Threatened | I |
| *Vaccinium myrtillus* | 678 | -0.0019 | <.001 | -0.40 | 173 | -0.0017 | NA | NA | 2696 | -0.0026 | <.001 | -1.21 | Dry to moderately moist forests | Not Threatened | I |
| *Viola canina* agg. | 90 | -0.0019 | <.001 | -0.41 | 2 | -0.0004 | NA | NA | 207 | -0.0003 | <.001 | -0.12 | Heaths, inland dunes and semi-natural grasslands | Near Threatened | I |
| *Thelypteris palustris* | 518 | -0.0019 | <.001 | -0.41 | 17 | 0.0002 | NA | NA | 98 | -0.0001 | <.001 | -0.05 | Moist to wet forests | Near Threatened | I |
| *Veronica beccabunga* | 1168 | -0.0019 | <.001 | -0.41 | 76 | 0.0013 | NA | NA | 2502 | -0.0008 | NA | NA | Moist to wet forests | Not Threatened | I |
| *Trifolium dubium* agg. | 252 | -0.0018 | NA | NA | 82 | -0.0072 | <.001 | -0.88 | 243 | -0.0004 | <.001 | -0.17 | Mesic grasslands | Not Threatened | I |
| *Juncus acutiflorus* | 197 | -0.0018 | <.001 | -0.38 | 48 | -0.0015 | <.001 | -0.19 | 2752 | -0.0051 | <.001 | -2.34 | Moist to wet grasslands | Not Threatened | I |
| *Hydrocharis morsus-ranae* | 293 | -0.0018 | <.001 | -0.37 | 55 | -0.0007 | NA | NA | 31 | 0.0000 | NA | NA | Linear and running surface waters | Near Threatened | I |
| *Thalictrum flavum* | 194 | -0.0018 | <.001 | -0.38 | 26 | -0.0006 | NA | NA | 163 | -0.0002 | <.001 | -0.08 | Fresh water vegetation, springs and reeds | Near Threatened | I |
| *Potentilla recta* | 78 | -0.0018 | <.001 | -0.38 | 4 | -0.0005 | NA | NA | 104 | -0.0001 | <.001 | -0.04 | Heaths, inland dunes and semi-natural grasslands | Not Threatened | A |
| *Thymus chamaedrys* agg. | 82 | -0.0018 | <.001 | -0.38 | 2 | -0.0002 | NA | NA | 4266 | -0.0058 | <.001 | -2.69 | Heaths, inland dunes and semi-natural grasslands | Not Threatened | I |
| *Ranunculus lanuginosus* | 325 | -0.0018 | <.001 | -0.39 | NA | NA | NA | NA | 260 | 0.0001 | NA | NA | Dry to moderately moist forests | Not Threatened | I |
| *Elymus junceiformis* | 116 | -0.0018 | <.001 | -0.38 | NA | NA | NA | NA | NA | NA | NA | NA | Coastal and marine habitats | Not Threatened | I |
| *Honckenya peploides* | 126 | -0.0018 | <.001 | -0.38 | NA | NA | NA | NA | NA | NA | NA | NA | Coastal and marine habitats | Not Threatened | I |
| *Centaurea jacea* agg. | 146 | -0.0017 | <.001 | -0.36 | 12 | -0.0012 | <.001 | -0.14 | 3258 | -0.0035 | <.001 | -1.60 | Heaths, inland dunes and semi-natural grasslands | Not Threatened | I |
| *Juncus inflexus* | 326 | -0.0017 | <.001 | -0.36 | 29 | -0.0008 | <.001 | -0.10 | 1996 | -0.0026 | <.001 | -1.22 | Fresh water vegetation, springs and reeds | Not Threatened | I |
| *Arrhenatherum elatius* | 566 | -0.0016 | NA | NA | 225 | -0.0074 | <.001 | -0.91 | 9891 | 0.0001 | NA | NA | Scrubs, copses and field hedges | Not Threatened | I |
| *Tussilago farfara* | 198 | -0.0016 | <.001 | -0.35 | 148 | -0.0067 | <.001 | -0.82 | 334 | -0.0003 | <.001 | -0.15 | Ruderal, fringe and tall forb communities, clearings | Not Threatened | I |
| *Vicia sativa* agg. | 188 | -0.0016 | NA | NA | 74 | -0.0047 | <.001 | -0.57 | 243 | -0.0003 | <.001 | -0.13 | Mesic grasslands | Not Threatened | A |
| *Ononis spinosa* agg. | 122 | -0.0016 | <.001 | -0.35 | 3 | -0.0004 | NA | NA | 2722 | -0.0033 | <.001 | -1.51 | Heaths, inland dunes and semi-natural grasslands | Not Threatened | I |
| *Rhinanthus minor* | 83 | -0.0016 | <.001 | -0.33 | 4 | -0.0004 | NA | NA | 742 | -0.0011 | <.001 | -0.49 | Heaths, inland dunes and semi-natural grasslands | Not Threatened | I |
| *Carex flacca* | 125 | -0.0016 | <.001 | -0.33 | 2 | 0.0002 | NA | NA | 3846 | -0.0040 | <.001 | -1.83 | Heaths, inland dunes and semi-natural grasslands | Not Threatened | I |
| *Paris quadrifolia* | 302 | -0.0016 | <.001 | -0.34 | 2 | 0.0002 | NA | NA | 2012 | -0.0001 | NA | NA | Moist to wet forests | Not Threatened | I |
| *Senecio jacobaea* | 386 | -0.0015 | NA | NA | 43 | -0.0027 | <.001 | -0.33 | 870 | -0.0008 | <.001 | -0.35 | Mesic grasslands | Not Threatened | I |
| *Festuca arundinacea* | 371 | -0.0015 | <.001 | -0.32 | 126 | -0.0027 | <.001 | -0.33 | 378 | -0.0002 | <.001 | -0.10 | Coastal and marine habitats | Not Threatened | I |
| *Stellaria aquatica* | 316 | -0.0015 | <.001 | -0.31 | 44 | -0.0006 | NA | NA | 244 | -0.0001 | NA | NA | Moist to wet forests | Not Threatened | I |
| *Veronica scutellata* | 112 | -0.0015 | <.001 | -0.32 | 20 | -0.0005 | NA | NA | 120 | -0.0002 | <.001 | -0.10 | Moist to wet grasslands | Not Threatened | I |
| *Mycelis muralis* | 222 | -0.0015 | <.001 | -0.33 | 43 | 0.0015 | <.001 | 0.18 | 1400 | 0.0001 | NA | NA | Dry to moderately moist forests | Not Threatened | I |
| *Trifolium fragiferum* | 118 | -0.0015 | <.001 | -0.31 | NA | NA | NA | NA | 4 | 0.0000 | NA | NA | Coastal and marine habitats | Near Threatened | I |
| *Artemisia vulgaris* agg. | 286 | -0.0014 | NA | NA | 379 | -0.0133 | <.001 | -1.63 | 972 | -0.0001 | NA | NA | Ruderal, fringe and tall forb communities, clearings | Not Threatened | NA |
| *Epilobium angustifolium* | 370 | -0.0014 | <.001 | -0.29 | 272 | -0.0062 | <.001 | -0.76 | 1104 | -0.0007 | <.001 | -0.33 | Coastal and marine habitats | Not Threatened | I |
| *Arenaria serpyllifolia* agg. | 92 | -0.0014 | NA | NA | 28 | -0.0043 | <.001 | -0.53 | 384 | -0.0005 | <.001 | -0.22 | Heaths, inland dunes and semi-natural grasslands | Not Threatened | I |
| *Vicia sepium* | 246 | -0.0014 | NA | NA | 122 | -0.0039 | <.001 | -0.48 | 3648 | -0.0013 | <.001 | -0.61 | Scrubs, copses and field hedges | Not Threatened | I |
| *Erodium cicutarium* agg. | 97 | -0.0014 | <.001 | -0.29 | 22 | -0.0035 | <.001 | -0.43 | 83 | -0.0002 | <.001 | -0.09 | Heaths, inland dunes and semi-natural grasslands | Not Threatened | I |
| *Leucanthemum vulgare* agg. | 108 | -0.0014 | NA | NA | 30 | -0.0027 | <.001 | -0.33 | 1989 | -0.0027 | <.001 | -1.25 | Heaths, inland dunes and semi-natural grasslands | Not Threatened | I |
| *Potentilla reptans* | 283 | -0.0014 | NA | NA | 36 | -0.0020 | <.001 | -0.24 | 1480 | -0.0003 | <.001 | -0.16 | Coastal and marine habitats | Not Threatened | I |
| *Aira caryophyllea* s. l. | 58 | -0.0014 | <.001 | -0.31 | 18 | -0.0016 | <.001 | -0.19 | 15 | 0.0000 | NA | NA | Heaths, inland dunes and semi-natural grasslands | Near Threatened | I |
| *Sonchus arvensis* agg. | 290 | -0.0014 | <.001 | -0.30 | 30 | -0.0014 | <.001 | -0.18 | 48 | 0.0000 | <.001 | -0.02 | Coastal and marine habitats | Not Threatened | I |
| *Rorippa amphibia* | 320 | -0.0014 | <.001 | -0.31 | 110 | -0.0008 | NA | NA | 77 | 0.0000 | NA | NA | Linear and running surface waters | Not Threatened | I |
| *Sium latifolium* | 178 | -0.0014 | <.001 | -0.31 | 22 | -0.0005 | NA | NA | 4 | 0.0000 | NA | NA | Linear and running surface waters | Not Threatened | I |
| *Pimpinella saxifraga* agg. | 112 | -0.0014 | <.001 | -0.31 | 2 | -0.0003 | NA | NA | 4114 | -0.0050 | <.001 | -2.31 | Heaths, inland dunes and semi-natural grasslands | Not Threatened | I |
| *Drosera intermedia* | 61 | -0.0014 | <.001 | -0.30 | 2 | -0.0003 | NA | NA | 8 | 0.0000 | <.001 | -0.01 | Bogs, transition mires, marshes and fens | Endangered | I |
| *Carex vesicaria* | 496 | -0.0014 | <.001 | -0.30 | 32 | -0.0002 | NA | NA | 1054 | -0.0013 | <.001 | -0.61 | Fresh water vegetation, springs and reeds | Not Threatened | I |
| *Campanula trachelium* | 133 | -0.0014 | <.001 | -0.29 | 2 | 0.0000 | NA | NA | 1874 | 0.0005 | <.001 | 0.24 | Scrubs, copses and field hedges | Not Threatened | I |
| *Carex appropinquata* | 100 | -0.0014 | <.001 | -0.29 | 4 | 0.0001 | NA | NA | 102 | -0.0002 | <.001 | -0.09 | Fresh water vegetation, springs and reeds | Endangered | I |
| *Epipactis helleborine* agg. | 194 | -0.0014 | <.001 | -0.29 | 18 | 0.0010 | <.001 | 0.13 | 872 | -0.0004 | <.001 | -0.20 | Dry to moderately moist forests | Not Threatened | I |
| *Salix cinerea_Salix multinervis* | 4214 | -0.0014 | NA | NA | 380 | 0.0059 | <.001 | 0.72 | 5122 | -0.0020 | <.001 | -0.92 | Standing waters | Not Threatened | I |
| *Juncus subnodulosus* | 92 | -0.0014 | <.001 | -0.29 | NA | NA | NA | NA | 315 | -0.0005 | <.001 | -0.25 | Moist to wet grasslands | Endangered | I |
| *Carex lasiocarpa* | 126 | -0.0014 | <.001 | -0.30 | NA | NA | NA | NA | 69 | -0.0002 | <.001 | -0.11 | Bogs, transition mires, marshes and fens | Endangered | I |
| *Cerastium semidecandrum* | 106 | -0.0013 | <.001 | -0.27 | 14 | -0.0025 | <.001 | -0.31 | 40 | -0.0001 | NA | NA | Coastal and marine habitats | Not Threatened | I |
| *Potentilla argentea* s. l. | 86 | -0.0013 | NA | NA | 14 | -0.0021 | <.001 | -0.25 | 80 | -0.0001 | NA | NA | Heaths, inland dunes and semi-natural grasslands | Not Threatened | I |
| *Alchemilla vulgaris* agg. | 110 | -0.0013 | <.001 | -0.27 | 18 | -0.0008 | <.001 | -0.09 | 1198 | -0.0017 | <.001 | -0.79 | Moist to wet grasslands | Not Threatened | I |
| *Odontites vernus* agg. | 122 | -0.0013 | <.001 | -0.28 | 6 | -0.0005 | NA | NA | 164 | -0.0002 | <.001 | -0.11 | Coastal and marine habitats | Not Threatened | I |
| *Veronica officinalis* | 138 | -0.0013 | <.001 | -0.28 | 50 | -0.0002 | NA | NA | 1396 | -0.0015 | <.001 | -0.71 | Heaths, inland dunes and semi-natural grasslands | Not Threatened | I |
| *Melampyrum pratense* | 322 | -0.0013 | <.001 | -0.27 | 38 | -0.0002 | NA | NA | 980 | -0.0007 | <.001 | -0.34 | Dry to moderately moist forests | Not Threatened | I |
| *Carex pilulifera* | 746 | -0.0013 | NA | NA | 102 | 0.0000 | NA | NA | 890 | -0.0009 | <.001 | -0.42 | Heaths, inland dunes and semi-natural grasslands | Not Threatened | I |
| *Listera ovata* | 93 | -0.0013 | <.001 | -0.28 | 4 | 0.0001 | NA | NA | 711 | -0.0010 | <.001 | -0.48 | Heaths, inland dunes and semi-natural grasslands | Not Threatened | I |
| *Silene dioica* | 634 | -0.0013 | <.001 | -0.29 | 50 | 0.0008 | NA | NA | 2386 | -0.0002 | NA | NA | Moist to wet forests | Not Threatened | I |
| *Orchis mascula* | 98 | -0.0013 | <.001 | -0.28 | NA | NA | NA | NA | 260 | -0.0004 | <.001 | -0.16 | Heaths, inland dunes and semi-natural grasslands | Near Threatened | I |
| *Puccinellia maritima* | 183 | -0.0013 | <.001 | -0.28 | NA | NA | NA | NA | NA | NA | NA | NA | Coastal and marine habitats | Not Threatened | I |
| *Tripleurospermum maritimum* agg. | 130 | -0.0012 | <.001 | -0.26 | 102 | -0.0090 | <.001 | -1.10 | 46 | -0.0001 | <.001 | -0.04 | Coastal and marine habitats | Not Threatened | NA |
| *Poa palustris* | 190 | -0.0012 | <.001 | -0.25 | 138 | -0.0028 | <.001 | -0.34 | 122 | -0.0001 | <.001 | -0.05 | Linear and running surface waters | Not Threatened | I |
| *Rorippa islandica* agg. | 182 | -0.0012 | <.001 | -0.26 | 72 | -0.0019 | <.001 | -0.23 | 69 | -0.0001 | NA | NA | Standing waters | Not Threatened | I |
| *Juncus bulbosus* | 118 | -0.0012 | <.001 | -0.26 | 12 | -0.0007 | NA | NA | 104 | -0.0002 | <.001 | -0.09 | Standing waters | Not Threatened | I |
| *Pedicularis palustris* | 44 | -0.0012 | <.001 | -0.25 | 2 | -0.0001 | NA | NA | 48 | -0.0001 | <.001 | -0.05 | None | Highly Endangered | I |
| *Oenanthe fistulosa* | 81 | -0.0012 | <.001 | -0.25 | 3 | -0.0001 | NA | NA | NA | NA | NA | NA | None | Endangered | I |
| *Ranunculus auricomus* agg. | 307 | -0.0012 | NA | NA | 6 | 0.0005 | NA | NA | 861 | -0.0005 | <.001 | -0.21 | None | Not Threatened | I |
| *Stachys sylvatica* | 2002 | -0.0012 | NA | NA | 72 | 0.0038 | <.001 | 0.47 | 4972 | 0.0035 | <.001 | 1.63 | Dry to moderately moist forests | Not Threatened | I |
| *Dianthus deltoides* | 39 | -0.0012 | <.001 | -0.26 | NA | NA | NA | NA | 232 | -0.0003 | <.001 | -0.16 | Heaths, inland dunes and semi-natural grasslands | Near Threatened | I |
| *Euphrasia stricta* agg. | 52 | -0.0012 | <.001 | -0.25 | NA | NA | NA | NA | 98 | -0.0001 | <.001 | -0.06 | Heaths, inland dunes and semi-natural grasslands | Not Threatened | I |
| *Cakile maritima* | 86 | -0.0012 | <.001 | -0.25 | NA | NA | NA | NA | 2 | 0.0000 | NA | NA | Coastal and marine habitats | Not Threatened | I |
| *Spergularia marina* | 130 | -0.0012 | <.001 | -0.25 | NA | NA | NA | NA | NA | NA | NA | NA | Coastal and marine habitats | Not Threatened | I |
| *Medicago lupulina* | 125 | -0.0011 | NA | NA | 96 | -0.0077 | <.001 | -0.94 | 1096 | -0.0014 | <.001 | -0.66 | Heaths, inland dunes and semi-natural grasslands | Not Threatened | I |
| *Vicia hirsuta* | 98 | -0.0011 | NA | NA | 65 | -0.0059 | <.001 | -0.73 | 148 | -0.0002 | <.001 | -0.07 | Mesic grasslands | Not Threatened | I |
| *Scleranthus annuus* agg. | 62 | -0.0011 | <.001 | -0.24 | 16 | -0.0016 | <.001 | -0.19 | 16 | 0.0000 | NA | NA | Heaths, inland dunes and semi-natural grasslands | Not Threatened | NA |
| *Angelica archangelica* | 256 | -0.0011 | <.001 | -0.23 | 56 | -0.0006 | NA | NA | 20 | 0.0000 | NA | NA | Linear and running surface waters | Not Threatened | I |
| *Genista pilosa* | 48 | -0.0011 | <.001 | -0.23 | 4 | -0.0005 | NA | NA | 128 | 0.0000 | NA | NA | Heaths, inland dunes and semi-natural grasslands | Near Threatened | I |
| *Anthyllis vulneraria* s. l. | 34 | -0.0011 | <.001 | -0.23 | 2 | -0.0002 | NA | NA | 1374 | -0.0023 | <.001 | -1.05 | Heaths, inland dunes and semi-natural grasslands | Not Threatened | I |
| *Ranunculus aquatilis* agg. | 194 | -0.0011 | <.001 | -0.23 | 10 | -0.0002 | NA | NA | 110 | -0.0001 | <.001 | -0.04 | Standing waters | Not Threatened | I |
| *Carex distans* | 119 | -0.0011 | <.001 | -0.22 | 1 | -0.0001 | NA | NA | 78 | -0.0001 | <.001 | -0.07 | Coastal and marine habitats | Endangered | I |
| *Nasturtium officinale* agg. | 323 | -0.0011 | <.001 | -0.23 | 66 | 0.0001 | NA | NA | 571 | -0.0004 | <.001 | -0.18 | Linear and running surface waters | Not Threatened | I |
| *Polypodium vulgare* agg. | 186 | -0.0011 | <.001 | -0.23 | 6 | 0.0003 | NA | NA | 936 | 0.0003 | <.001 | 0.15 | Dry to moderately moist forests | Not Threatened | I |
| *Carex riparia* | 913 | -0.0011 | NA | NA | 46 | 0.0010 | NA | NA | 286 | -0.0002 | <.001 | -0.09 | Fresh water vegetation, springs and reeds | Not Threatened | I |
| *Festuca gigantea* | 936 | -0.0011 | NA | NA | 168 | 0.0092 | <.001 | 1.13 | 732 | 0.0004 | <.001 | 0.21 | Moist to wet forests | Not Threatened | I |
| *Sanguisorba officinalis* | 64 | -0.0011 | <.001 | -0.24 | NA | NA | NA | NA | 2707 | -0.0042 | <.001 | -1.96 | Moist to wet grasslands | Near Threatened | I |
| *Vaccinium uliginosum* s. l. | 76 | -0.0011 | <.001 | -0.24 | NA | NA | NA | NA | 484 | -0.0013 | <.001 | -0.60 | Bogs, transition mires, marshes and fens | Near Threatened | I |
| *Campanula latifolia* | 106 | -0.0011 | <.001 | -0.24 | NA | NA | NA | NA | 14 | 0.0000 | NA | NA | Dry to moderately moist forests | Not Threatened | I |
| *Atriplex littoralis* | 88 | -0.0011 | <.001 | -0.23 | NA | NA | NA | NA | NA | NA | NA | NA | Coastal and marine habitats | Not Threatened | I |
| *Heracleum sphondylium* | 378 | -0.0010 | NA | NA | 353 | -0.0046 | <.001 | -0.57 | 4150 | -0.0020 | <.001 | -0.93 | Scrubs, copses and field hedges | Not Threatened | I |
| *Crepis capillaris* | 76 | -0.0010 | NA | NA | 34 | -0.0027 | <.001 | -0.32 | 142 | 0.0000 | NA | NA | Mesic grasslands | Not Threatened | A |
| *Echium vulgare* | 55 | -0.0010 | NA | NA | 18 | -0.0017 | NA | NA | 722 | -0.0011 | <.001 | -0.50 | Heaths, inland dunes and semi-natural grasslands | Not Threatened | A |
| *Filago minima* | 62 | -0.0010 | <.001 | -0.21 | 4 | -0.0017 | NA | NA | 26 | -0.0001 | NA | NA | Heaths, inland dunes and semi-natural grasslands | Not Threatened | I |
| *Oenanthe aquatica* agg. | 431 | -0.0010 | <.001 | -0.21 | 68 | -0.0015 | <.001 | -0.18 | 22 | 0.0000 | <.001 | -0.02 | Standing waters | Not Threatened | I |
| *Agrimonia eupatoria* | 130 | -0.0010 | NA | NA | 25 | -0.0008 | NA | NA | 5536 | -0.0022 | <.001 | -1.02 | Heaths, inland dunes and semi-natural grasslands | Not Threatened | I |
| *Veronica anagallis-aquatica* agg. | 164 | -0.0010 | <.001 | -0.21 | 32 | -0.0006 | NA | NA | 238 | -0.0002 | <.001 | -0.08 | Linear and running surface waters | Not Threatened | I |
| *Elodea canadensis* | 378 | -0.0010 | NA | NA | 42 | -0.0005 | NA | NA | 166 | -0.0002 | <.001 | -0.07 | Linear and running surface waters | Not Evaluated | N |
| *Carex cespitosa* | 92 | -0.0010 | <.001 | -0.21 | 4 | 0.0001 | NA | NA | 101 | -0.0001 | <.001 | -0.06 | Fresh water vegetation, springs and reeds | Endangered | I |
| *Linum catharticum* | 44 | -0.0010 | <.001 | -0.21 | NA | NA | NA | NA | 1898 | -0.0027 | <.001 | -1.27 | Heaths, inland dunes and semi-natural grasslands | Not Threatened | I |
| *Helictotrichon pratense* | 38 | -0.0010 | <.001 | -0.22 | NA | NA | NA | NA | 1407 | -0.0018 | <.001 | -0.82 | Heaths, inland dunes and semi-natural grasslands | Near Threatened | I |
| *Gnaphalium uliginosum* | 84 | -0.0009 | <.001 | -0.19 | 37 | -0.0029 | <.001 | -0.35 | 28 | 0.0000 | <.001 | -0.02 | Standing waters | Not Threatened | I |
| *Tragopogon pratensis* s. l. | 90 | -0.0009 | NA | NA | 40 | -0.0026 | <.001 | -0.32 | 663 | -0.0009 | <.001 | -0.41 | Heaths, inland dunes and semi-natural grasslands | Not Threatened | I |
| *Veronica serpyllifolia* | 136 | -0.0009 | NA | NA | 24 | -0.0013 | <.001 | -0.16 | 56 | -0.0001 | <.001 | -0.05 | Mesic grasslands | Not Threatened | I |
| *Trifolium medium* | 86 | -0.0009 | NA | NA | 16 | -0.0008 | NA | NA | 2088 | -0.0016 | <.001 | -0.73 | Heaths, inland dunes and semi-natural grasslands | Not Threatened | I |
| *Centaurium erythraea* s. l. | 58 | -0.0009 | <.001 | -0.20 | 8 | -0.0006 | NA | NA | 193 | -0.0002 | <.001 | -0.10 | Heaths, inland dunes and semi-natural grasslands | Not Threatened | I |
| *Acorus calamus* | 162 | -0.0009 | <.001 | -0.19 | 57 | -0.0002 | NA | NA | 92 | -0.0001 | <.001 | -0.07 | Standing waters | Not Evaluated | N |
| *Arnica montana* | 30 | -0.0009 | <.001 | -0.20 | 1 | -0.0001 | NA | NA | 484 | -0.0012 | <.001 | -0.56 | Heaths, inland dunes and semi-natural grasslands | Endangered | I |
| *Polygala vulgaris* s. l. | 40 | -0.0009 | <.001 | -0.18 | 3 | -0.0001 | NA | NA | 886 | -0.0012 | <.001 | -0.56 | Heaths, inland dunes and semi-natural grasslands | Near Threatened | I |
| *Potamogeton crispus* | 165 | -0.0009 | NA | NA | 20 | -0.0001 | NA | NA | 72 | -0.0001 | <.001 | -0.04 | Linear and running surface waters | Not Threatened | I |
| *Saxifraga granulata* | 70 | -0.0009 | <.001 | -0.18 | NA | NA | NA | NA | 105 | -0.0001 | <.001 | -0.07 | Moist to wet grasslands | Near Threatened | I |
| *Erigeron canadensis* | 84 | -0.0008 | NA | NA | 85 | -0.0087 | <.001 | -1.06 | 158 | 0.0001 | NA | NA | Ruderal, fringe and tall forb communities, clearings | Not Evaluated | N |
| *Oenothera* | 68 | -0.0008 | NA | NA | 72 | -0.0072 | <.001 | -0.88 | 108 | -0.0001 | NA | NA | Ruderal, fringe and tall forb communities, clearings | Not Evaluated | N |
| *Poa annua* agg. | 201 | -0.0008 | NA | NA | 262 | -0.0064 | <.001 | -0.78 | 126 | -0.0002 | <.001 | -0.08 | Farmland | Not Threatened | I |
| *Mentha arvensis* | 84 | -0.0008 | <.001 | -0.16 | 82 | -0.0031 | <.001 | -0.37 | 234 | -0.0003 | <.001 | -0.12 | Moist to wet grasslands | Not Threatened | I |
| *Hypericum maculatum* agg. | 112 | -0.0008 | <.001 | -0.18 | 37 | -0.0009 | <.001 | -0.11 | 408 | -0.0004 | <.001 | -0.17 | Heaths, inland dunes and semi-natural grasslands | Not Threatened | I |
| *Silene vulgaris* | 54 | -0.0008 | NA | NA | 4 | -0.0007 | NA | NA | 2031 | -0.0010 | <.001 | -0.48 | Heaths, inland dunes and semi-natural grasslands | Not Threatened | I |
| *Gentiana pneumonanthe* | 30 | -0.0008 | <.001 | -0.17 | 5 | -0.0003 | NA | NA | 25 | -0.0001 | <.001 | -0.02 | None | Highly Endangered | I |
| *Geranium palustre* | 70 | -0.0008 | <.001 | -0.17 | 1 | -0.0001 | NA | NA | 1274 | -0.0005 | <.001 | -0.23 | Fresh water vegetation, springs and reeds | Not Threatened | I |
| *Dactylorhiza incarnata* agg. | 30 | -0.0008 | <.001 | -0.17 | 2 | -0.0001 | NA | NA | 182 | -0.0004 | <.001 | -0.20 | Moist to wet grasslands | Endangered | I |
| *Hieracium murorum* | 74 | -0.0008 | <.001 | -0.16 | 10 | -0.0001 | NA | NA | 1318 | -0.0004 | NA | NA | Dry to moderately moist forests | Not Threatened | I |
| *Dryopteris cristata* | 226 | -0.0008 | <.001 | -0.18 | 4 | 0.0001 | NA | NA | 18 | -0.0001 | <.001 | -0.02 | Bogs, transition mires, marshes and fens | Endangered | I |
| *Stratiotes aloides* | 90 | -0.0008 | <.001 | -0.16 | 14 | 0.0002 | NA | NA | 42 | -0.0001 | NA | NA | Standing waters | Endangered | I |
| *Chrysosplenium alternifolium* | 519 | -0.0008 | NA | NA | 4 | 0.0007 | NA | NA | 1266 | 0.0009 | <.001 | 0.43 | Moist to wet forests | Not Threatened | I |
| *Parnassia palustris* | 25 | -0.0008 | <.001 | -0.16 | NA | NA | NA | NA | 554 | -0.0014 | <.001 | -0.65 | Moist to wet grasslands | Endangered | I |
| *Eryngium maritimum* | 43 | -0.0008 | <.001 | -0.17 | NA | NA | NA | NA | NA | NA | NA | NA | Coastal and marine habitats | Highly Endangered | I |
| *Hierochloe odorata* agg. | 80 | -0.0008 | <.001 | -0.16 | NA | NA | NA | NA | NA | NA | NA | NA | Bogs, transition mires, marshes and fens | Highly Endangered | I |
| *Senecio viscosus* | 59 | -0.0007 | <.001 | -0.15 | 33 | -0.0028 | <.001 | -0.34 | 28 | 0.0000 | <.001 | -0.01 | Coastal and marine habitats | Not Threatened | I |
| *Convolvulus arvensis* | 139 | -0.0007 | NA | NA | 50 | -0.0024 | <.001 | -0.29 | 1052 | -0.0007 | <.001 | -0.33 | Scrubs, copses and field hedges | Not Threatened | I |
| *Bistorta officinalis* | 52 | -0.0007 | <.001 | -0.15 | 62 | -0.0015 | <.001 | -0.18 | 3288 | -0.0052 | <.001 | -2.41 | Moist to wet grasslands | Not Threatened | I |
| *Verbascum nigrum* | 80 | -0.0007 | NA | NA | 24 | -0.0015 | <.001 | -0.18 | 222 | -0.0002 | <.001 | -0.11 | Heaths, inland dunes and semi-natural grasslands | Not Threatened | I |
| *Pastinaca sativa* | 56 | -0.0007 | NA | NA | 12 | -0.0005 | NA | NA | 486 | -0.0004 | <.001 | -0.20 | Coastal and marine habitats | Not Threatened | I |
| *Mentha verticillata* agg. | 44 | -0.0007 | <.001 | -0.15 | 8 | -0.0003 | NA | NA | 14 | 0.0000 | NA | NA | None | Not Threatened | I |
| *Ranunculus bulbosus* agg. | 76 | -0.0007 | NA | NA | 1 | -0.0002 | NA | NA | 1486 | -0.0019 | <.001 | -0.86 | Heaths, inland dunes and semi-natural grasslands | Not Threatened | I |
| *Bromus racemosus* agg. | 62 | -0.0007 | <.001 | -0.14 | 10 | -0.0002 | NA | NA | 150 | -0.0002 | <.001 | -0.11 | Moist to wet grasslands | Vulnerable | NA |
| *Allium scorodoprasum* | 58 | -0.0007 | <.001 | -0.14 | 16 | -0.0002 | NA | NA | 40 | 0.0000 | NA | NA | Coastal and marine habitats | Not Threatened | I |
| *Osmunda regalis* | 93 | -0.0007 | <.001 | -0.16 | 13 | -0.0002 | NA | NA | 4 | 0.0000 | NA | NA | Moist to wet forests | Endangered | I |
| *Pedicularis sylvatica* | 18 | -0.0007 | <.001 | -0.14 | 2 | -0.0001 | NA | NA | 537 | -0.0012 | <.001 | -0.54 | Moist to wet grasslands | Endangered | I |
| *Ophioglossum vulgatum* | 37 | -0.0007 | <.001 | -0.15 | 1 | -0.0001 | NA | NA | 26 | 0.0000 | <.001 | -0.02 | Coastal and marine habitats | Endangered | I |
| *Salsola kali* s. l. | 46 | -0.0007 | <.001 | -0.16 | 1 | -0.0001 | NA | NA | 10 | 0.0000 | NA | NA | Coastal and marine habitats | Not Threatened | NA |
| *Viola riviniana* agg. | 756 | -0.0007 | NA | NA | 12 | 0.0014 | <.001 | 0.17 | 1551 | 0.0001 | NA | NA | Dry to moderately moist forests | Not Threatened | I |
| *Ajuga reptans* | 1841 | -0.0007 | NA | NA | 144 | 0.0037 | <.001 | 0.46 | 2862 | -0.0010 | <.001 | -0.47 | Moist to wet grasslands | Not Threatened | I |
| *Centaurea scabiosa* | 36 | -0.0007 | NA | NA | NA | NA | NA | NA | 3098 | -0.0027 | <.001 | -1.23 | Heaths, inland dunes and semi-natural grasslands | Not Threatened | I |
| *Thymus serpyllum* | 38 | -0.0007 | <.001 | -0.15 | NA | NA | NA | NA | 24 | -0.0001 | NA | NA | Coastal and marine habitats | Near Threatened | I |
| *Lycopodiella inundata* | 19 | -0.0007 | <.001 | -0.15 | NA | NA | NA | NA | 6 | 0.0000 | NA | NA | Coastal and marine habitats | Endangered | I |
| *Hordelymus europaeus* | 130 | -0.0007 | NA | NA | NA | NA | NA | NA | 1134 | 0.0006 | <.001 | 0.27 | Dry to moderately moist forests | Not Threatened | I |
| *Artemisia maritima* agg. | 76 | -0.0007 | <.001 | -0.15 | NA | NA | NA | NA | NA | NA | NA | NA | Coastal and marine habitats | Not Threatened | I |
| *Calammophila baltica* | 65 | -0.0007 | <.001 | -0.16 | NA | NA | NA | NA | NA | NA | NA | NA | Coastal and marine habitats | Not Threatened | I |
| *Crambe maritima* | 44 | -0.0007 | <.001 | -0.14 | NA | NA | NA | NA | NA | NA | NA | NA | Coastal and marine habitats | Endangered | I |
| *Suaeda maritima* | 72 | -0.0007 | <.001 | -0.15 | NA | NA | NA | NA | NA | NA | NA | NA | Coastal and marine habitats | Not Threatened | I |
| *Persicaria maculosa* | 101 | -0.0006 | <.001 | -0.13 | 82 | -0.0042 | <.001 | -0.51 | 56 | -0.0001 | NA | NA | Farmland | Not Threatened | I |
| *Geranium molle* agg. | 95 | -0.0006 | NA | NA | 24 | -0.0026 | <.001 | -0.31 | 80 | 0.0000 | NA | NA | Mesic grasslands | Not Threatened | A |
| *Cerastium arvense* | 59 | -0.0006 | NA | NA | 13 | -0.0012 | <.001 | -0.14 | 718 | -0.0010 | <.001 | -0.46 | Heaths, inland dunes and semi-natural grasslands | Not Threatened | I |
| *Chenopodium rubrum* agg. | 45 | -0.0006 | <.001 | -0.12 | 6 | -0.0011 | NA | NA | 4 | 0.0000 | NA | NA | Coastal and marine habitats | Not Threatened | I |
| *Carduus crispus* | 72 | -0.0006 | <.001 | -0.12 | 21 | -0.0010 | <.001 | -0.12 | 313 | 0.0000 | NA | NA | None | Not Threatened | I |
| *Solidago virgaurea* | 53 | -0.0006 | <.001 | -0.14 | 33 | -0.0007 | <.001 | -0.08 | 1683 | -0.0005 | NA | NA | Dry to moderately moist forests | Not Threatened | I |
| *Carex muricata* agg. | 112 | -0.0006 | NA | NA | 31 | -0.0007 | NA | NA | 634 | -0.0003 | <.001 | -0.14 | None | Not Threatened | I |
| *Epilobium tetragonum* s. l. | 192 | -0.0006 | <.001 | -0.13 | 26 | -0.0005 | NA | NA | 216 | -0.0002 | <.001 | -0.11 | Moist to wet grasslands | Not Threatened | I |
| *Hieracium sabaudum* | 61 | -0.0006 | <.001 | -0.14 | 50 | -0.0005 | NA | NA | 195 | -0.0001 | <.001 | -0.05 | Dry to moderately moist forests | Not Threatened | I |
| *Artemisia campestris* | 27 | -0.0006 | <.001 | -0.12 | 4 | -0.0004 | NA | NA | 170 | -0.0002 | <.001 | -0.10 | Heaths, inland dunes and semi-natural grasslands | Not Threatened | I |
| *Erigeron acris* s. l. | 24 | -0.0006 | <.001 | -0.12 | 3 | -0.0002 | NA | NA | 260 | -0.0003 | <.001 | -0.15 | Heaths, inland dunes and semi-natural grasslands | Not Evaluated | I |
| *Leontodon hispidus* | 58 | -0.0006 | NA | NA | 2 | -0.0001 | NA | NA | 1444 | -0.0020 | <.001 | -0.94 | Heaths, inland dunes and semi-natural grasslands | Not Threatened | I |
| *Astragalus glycyphyllos* | 41 | -0.0006 | <.001 | -0.12 | 2 | -0.0001 | NA | NA | 1037 | -0.0007 | <.001 | -0.32 | Heaths, inland dunes and semi-natural grasslands | Not Threatened | I |
| *Lathyrus palustris* | 48 | -0.0006 | <.001 | -0.14 | 2 | -0.0001 | NA | NA | 10 | 0.0000 | NA | NA | None | Endangered | I |
| *Ulmus glabra* | 690 | -0.0006 | NA | NA | 62 | 0.0026 | <.001 | 0.31 | 2963 | 0.0020 | <.001 | 0.92 | Dry to moderately moist forests | Not Threatened | I |
| *Scorzonera humilis* | 24 | -0.0006 | <.001 | -0.14 | NA | NA | NA | NA | 91 | -0.0002 | <.001 | -0.09 | Moist to wet grasslands | Endangered | I |
| *Actaea spicata* | 44 | -0.0006 | <.001 | -0.12 | NA | NA | NA | NA | 676 | 0.0000 | NA | NA | Dry to moderately moist forests | Not Threatened | I |
| *Centaurium pulchellum* | 56 | -0.0006 | <.001 | -0.13 | NA | NA | NA | NA | 17 | 0.0000 | <.001 | -0.02 | Coastal and marine habitats | Near Threatened | I |
| *Helichrysum arenarium* | 22 | -0.0006 | <.001 | -0.12 | NA | NA | NA | NA | 8 | 0.0000 | NA | NA | None | Endangered | I |
| *Radiola linoides* | 8 | -0.0006 | <.001 | -0.14 | NA | NA | NA | NA | 1 | 0.0000 | NA | NA | Coastal and marine habitats | Highly Endangered | I |
| *Equisetum telmateia* | 155 | -0.0006 | <.001 | -0.13 | NA | NA | NA | NA | 1108 | 0.0002 | NA | NA | Moist to wet forests | Not Threatened | I |
| *Carex extensa* | 45 | -0.0006 | <.001 | -0.12 | NA | NA | NA | NA | NA | NA | NA | NA | Coastal and marine habitats | Near Threatened | I |
| *Cotula coronopifolia* | 115 | -0.0006 | <.001 | -0.13 | NA | NA | NA | NA | NA | NA | NA | NA | Coastal and marine habitats | Not Evaluated | N |
| *Plantago coronopus* | 48 | -0.0006 | <.001 | -0.12 | NA | NA | NA | NA | NA | NA | NA | NA | Coastal and marine habitats | Not Threatened | I |
| *Puccinellia distans* agg. | 80 | -0.0006 | <.001 | -0.14 | NA | NA | NA | NA | NA | NA | NA | NA | Coastal and marine habitats | Not Threatened | I |
| *Sagina nodosa* | 22 | -0.0006 | <.001 | -0.12 | NA | NA | NA | NA | NA | NA | NA | NA | Coastal and marine habitats | Highly Endangered | I |
| *Polygonum aviculare* agg. | 124 | -0.0005 | NA | NA | 124 | -0.0066 | <.001 | -0.81 | 111 | 0.0000 | NA | NA | Mesic grasslands | Not Threatened | I |
| *Melilotus albus* | 34 | -0.0005 | NA | NA | 50 | -0.0041 | <.001 | -0.50 | 346 | -0.0002 | <.001 | -0.11 | Ruderal, fringe and tall forb communities, clearings | Not Threatened | A |
| *Veronica arvensis* | 54 | -0.0005 | NA | NA | 27 | -0.0026 | <.001 | -0.32 | 74 | -0.0001 | <.001 | -0.07 | Mesic grasslands | Not Threatened | A |
| *Juncus tenuis* | 82 | -0.0005 | NA | NA | 90 | -0.0015 | <.001 | -0.18 | 76 | -0.0001 | <.001 | -0.04 | Standing waters | Not Evaluated | N |
| *Rumex maritimus* | 94 | -0.0005 | <.001 | -0.11 | 28 | -0.0014 | <.001 | -0.17 | 7 | 0.0000 | NA | NA | Coastal and marine habitats | Not Threatened | I |
| *Draba verna* agg. | 31 | -0.0005 | NA | NA | 6 | -0.0007 | NA | NA | 108 | -0.0002 | <.001 | -0.11 | Heaths, inland dunes and semi-natural grasslands | Not Threatened | I |
| *Persicaria minor* | 32 | -0.0005 | <.001 | -0.11 | 23 | -0.0006 | NA | NA | 8 | 0.0000 | NA | NA | None | Not Threatened | I |
| *Salix pentandra* | 406 | -0.0005 | <.001 | -0.11 | 93 | -0.0005 | NA | NA | 146 | -0.0002 | <.001 | -0.09 | Moist to wet forests | Not Threatened | I |
| *Lathyrus sylvestris* | 42 | -0.0005 | NA | NA | 10 | -0.0005 | NA | NA | 298 | -0.0001 | <.001 | -0.06 | None | Not Threatened | I |
| *Eleocharis quinqueflora* | 10 | -0.0005 | <.001 | -0.11 | 1 | -0.0002 | NA | NA | 2 | 0.0000 | NA | NA | Coastal and marine habitats | Highly Endangered | I |
| *Utricularia minor* agg. | 26 | -0.0005 | <.001 | -0.10 | 2 | -0.0002 | NA | NA | 12 | 0.0000 | NA | NA | Bogs, transition mires, marshes and fens | Endangered | I |
| *Hippuris vulgaris* | 38 | -0.0005 | <.001 | -0.10 | 6 | -0.0001 | NA | NA | 82 | -0.0001 | NA | NA | Fresh water vegetation, springs and reeds | Near Threatened | I |
| *Genista tinctoria* | 18 | -0.0005 | <.001 | -0.10 | 1 | 0.0000 | NA | NA | 550 | -0.0007 | <.001 | -0.32 | Heaths, inland dunes and semi-natural grasslands | Near Threatened | I |
| *Cochlearia danica* | 24 | -0.0005 | <.001 | -0.10 | 1 | 0.0000 | NA | NA | NA | NA | NA | NA | Coastal and marine habitats | Not Threatened | I |
| *Stellaria nemorum* s. l. | 602 | -0.0005 | NA | NA | 30 | 0.0028 | <.001 | 0.35 | 968 | 0.0004 | <.001 | 0.17 | Moist to wet forests | Not Threatened | I |
| *Juncus anceps* | 14 | -0.0005 | <.001 | -0.10 | NA | NA | NA | NA | NA | NA | NA | NA | Coastal and marine habitats | Not Threatened | I |
| *Juncus maritimus* | 40 | -0.0005 | <.001 | -0.10 | NA | NA | NA | NA | NA | NA | NA | NA | Coastal and marine habitats | Not Threatened | I |
| *Lathyrus japonicus* | 46 | -0.0005 | <.001 | -0.10 | NA | NA | NA | NA | NA | NA | NA | NA | Coastal and marine habitats | Endangered | I |
| *Spergularia media* | 79 | -0.0005 | <.001 | -0.11 | NA | NA | NA | NA | NA | NA | NA | NA | Coastal and marine habitats | Not Threatened | I |
| *Melilotus officinalis* | 37 | -0.0004 | NA | NA | 37 | -0.0038 | <.001 | -0.47 | 438 | -0.0004 | <.001 | -0.18 | Ruderal, fringe and tall forb communities, clearings | Not Threatened | A |
| *Anthriscus sylvestris* agg. | 672 | -0.0004 | NA | NA | 312 | -0.0033 | NA | NA | 1688 | -0.0006 | <.001 | -0.30 | Scrubs, copses and field hedges | Not Threatened | I |
| *Sonchus asper* | 86 | -0.0004 | NA | NA | 60 | -0.0031 | <.001 | -0.38 | 140 | 0.0000 | NA | NA | Farmland | Not Threatened | I |
| *Senecio vulgaris* | 42 | -0.0004 | NA | NA | 65 | -0.0027 | <.001 | -0.33 | 30 | 0.0000 | <.001 | -0.02 | Farmland | Not Threatened | I |
| *Gnaphalium sylvaticum* | 24 | -0.0004 | <.001 | -0.08 | 44 | -0.0025 | <.001 | -0.31 | 52 | -0.0001 | <.001 | -0.03 | Heaths, inland dunes and semi-natural grasslands | Not Threatened | I |
| *Stellaria palustris* | 173 | -0.0004 | <.001 | -0.08 | 74 | -0.0017 | NA | NA | 24 | 0.0000 | <.001 | -0.02 | Linear and running surface waters | Endangered | I |
| *Spergula arvensis* | 25 | -0.0004 | <.001 | -0.08 | 10 | -0.0012 | <.001 | -0.15 | 4 | 0.0000 | NA | NA | Farmland | Not Threatened | A |
| *Leontodon saxatilis* | 24 | -0.0004 | <.001 | -0.08 | 6 | -0.0010 | NA | NA | 3 | 0.0000 | NA | NA | Mesic grasslands | Not Threatened | I |
| *Elymus caninus* | 48 | -0.0004 | <.001 | -0.08 | 18 | -0.0006 | <.001 | -0.07 | 616 | 0.0003 | <.001 | 0.13 | Moist to wet forests | Not Threatened | I |
| *Hylotelephium telephium* agg. | 66 | -0.0004 | NA | NA | 9 | -0.0004 | NA | NA | 1048 | -0.0007 | <.001 | -0.31 | Scrubs, copses and field hedges | Not Threatened | I |
| *Allium oleraceum* | 18 | -0.0004 | <.001 | -0.08 | 1 | -0.0001 | NA | NA | 646 | -0.0004 | <.001 | -0.17 | Scrubs, copses and field hedges | Not Threatened | I |
| *Epilobium obscurum* | 179 | -0.0004 | NA | NA | 6 | -0.0001 | NA | NA | 140 | -0.0002 | <.001 | -0.08 | Moist to wet grasslands | Near Threatened | I |
| *Lycopodium clavatum* | 17 | -0.0004 | <.001 | -0.08 | 2 | -0.0001 | NA | NA | 36 | -0.0001 | NA | NA | None | Endangered | I |
| *Utricularia vulgaris* agg. | 89 | -0.0004 | <.001 | -0.09 | 9 | -0.0001 | NA | NA | 94 | -0.0001 | <.001 | -0.04 | Standing waters | Near Threatened | I |
| *Symphytum asperum* agg. | 45 | -0.0004 | <.001 | -0.09 | 2 | -0.0001 | NA | NA | NA | NA | NA | NA | None | Not Evaluated | N |
| *Carex diandra* | 30 | -0.0004 | <.001 | -0.09 | 1 | 0.0000 | NA | NA | 26 | -0.0001 | <.001 | -0.03 | None | Highly Endangered | I |
| *Rosa spinosissima* | 15 | -0.0004 | <.001 | -0.08 | 4 | 0.0003 | NA | NA | 125 | -0.0001 | NA | NA | None | Endangered | I |
| *Equisetum hyemale* | 115 | -0.0004 | NA | NA | 1 | 0.0004 | NA | NA | 258 | 0.0001 | <.001 | 0.07 | Moist to wet forests | Not Threatened | I |
| *Carlina vulgaris* agg. | 20 | -0.0004 | NA | NA | NA | NA | NA | NA | 949 | -0.0014 | <.001 | -0.67 | Heaths, inland dunes and semi-natural grasslands | Not Threatened | I |
| *Blysmus compressus* | 22 | -0.0004 | <.001 | -0.08 | NA | NA | NA | NA | 8 | 0.0000 | NA | NA | None | Highly Endangered | I |
| *Primula vulgaris* | 42 | -0.0004 | <.001 | -0.08 | NA | NA | NA | NA | 3 | 0.0000 | NA | NA | None | Endangered | I |
| *Ranunculus sardous* | 40 | -0.0004 | <.001 | -0.08 | NA | NA | NA | NA | 1 | 0.0000 | NA | NA | Coastal and marine habitats | Endangered | I |
| *Centaurium littorale* | 36 | -0.0004 | <.001 | -0.09 | NA | NA | NA | NA | NA | NA | NA | NA | Coastal and marine habitats | Not Threatened | I |
| *Lepidium latifolium* | 36 | -0.0004 | <.001 | -0.08 | NA | NA | NA | NA | NA | NA | NA | NA | Coastal and marine habitats | Not Threatened | I |
| *Limonium vulgare* | 28 | -0.0004 | <.001 | -0.08 | NA | NA | NA | NA | NA | NA | NA | NA | Coastal and marine habitats | Not Threatened | I |
| *Capsella bursa-pastoris* | 58 | -0.0003 | NA | NA | 90 | -0.0064 | <.001 | -0.78 | 85 | -0.0001 | <.001 | -0.05 | Mesic grasslands | Not Threatened | I |
| *Persicaria lapathifolia* s. l. | 62 | -0.0003 | NA | NA | 70 | -0.0049 | <.001 | -0.60 | 66 | -0.0001 | <.001 | -0.04 | Mesic grasslands | Not Threatened | I |
| *Chenopodium album* agg. | 32 | -0.0003 | NA | NA | 52 | -0.0049 | <.001 | -0.60 | 118 | 0.0000 | NA | NA | Anthropogenic | Not Threatened | NA |
| *Silene latifolia* | 59 | -0.0003 | NA | NA | 43 | -0.0029 | <.001 | -0.36 | 842 | 0.0003 | <.001 | 0.13 | Scrubs, copses and field hedges | Not Threatened | I |
| *Poa compressa* | 33 | -0.0003 | NA | NA | 14 | -0.0017 | NA | NA | 236 | -0.0003 | <.001 | -0.15 | Heaths, inland dunes and semi-natural grasslands | Not Threatened | I |
| *Lupinus* | 50 | -0.0003 | NA | NA | 26 | -0.0014 | <.001 | -0.17 | 114 | -0.0001 | <.001 | -0.05 | Ruderal, fringe and tall forb communities, clearings | Not Evaluated | N |
| *Agrostis vinealis* | 24 | -0.0003 | <.001 | -0.06 | 4 | -0.0014 | NA | NA | 4 | 0.0000 | NA | NA | Heaths, inland dunes and semi-natural grasslands | Near Threatened | I |
| *Spergula pentandra* agg. | 30 | -0.0003 | <.001 | -0.06 | 3 | -0.0006 | NA | NA | 6 | 0.0000 | NA | NA | Heaths, inland dunes and semi-natural grasslands | Near Threatened | I |
| *Hieracium laevigatum* | 40 | -0.0003 | NA | NA | 30 | -0.0005 | NA | NA | 58 | -0.0001 | <.001 | -0.03 | Heaths, inland dunes and semi-natural grasslands | Not Threatened | I |
| *Hypericum humifusum* | 13 | -0.0003 | <.001 | -0.06 | 9 | -0.0004 | NA | NA | 30 | 0.0000 | <.001 | -0.01 | None | Not Threatened | I |
| *Cladium mariscus* | 20 | -0.0003 | <.001 | -0.05 | 7 | -0.0003 | NA | NA | 44 | -0.0001 | <.001 | -0.04 | Fresh water vegetation, springs and reeds | Endangered | I |
| *Epipactis palustris* | 14 | -0.0003 | <.001 | -0.07 | 1 | -0.0002 | NA | NA | 207 | -0.0005 | <.001 | -0.24 | Moist to wet grasslands | Endangered | I |
| *Festuca filiformis* | 12 | -0.0003 | <.001 | -0.07 | 4 | -0.0002 | NA | NA | 14 | 0.0000 | NA | NA | Heaths, inland dunes and semi-natural grasslands | Not Threatened | I |
| *Salix aurita* | 1329 | -0.0003 | NA | NA | 128 | -0.0001 | NA | NA | 1895 | -0.0021 | <.001 | -0.99 | Bogs, transition mires, marshes and fens | Not Threatened | I |
| *Carex pallescens* | 54 | -0.0003 | NA | NA | 8 | -0.0001 | NA | NA | 922 | -0.0016 | <.001 | -0.74 | Moist to wet grasslands | Not Threatened | I |
| *Juniperus communis* s. l. | 21 | -0.0003 | <.001 | -0.07 | 6 | 0.0000 | NA | NA | 2428 | -0.0037 | <.001 | -1.70 | Heaths, inland dunes and semi-natural grasslands | Near Threatened | I |
| *Clinopodium vulgare* | 14 | -0.0003 | NA | NA | 1 | 0.0000 | NA | NA | 2348 | -0.0008 | <.001 | -0.36 | Heaths, inland dunes and semi-natural grasslands | Not Threatened | I |
| *Hepatica nobilis* | 24 | -0.0003 | <.001 | -0.07 | 1 | 0.0000 | NA | NA | 345 | -0.0002 | NA | NA | Dry to moderately moist forests | Not Threatened | I |
| *Potentilla sterilis* | 25 | -0.0003 | <.001 | -0.06 | 1 | 0.0000 | NA | NA | 436 | 0.0001 | NA | NA | Dry to moderately moist forests | Not Threatened | I |
| *Pinus sylvestris* | 824 | -0.0003 | NA | NA | 251 | 0.0004 | NA | NA | 6027 | -0.0032 | <.001 | -1.47 | Heaths, inland dunes and semi-natural grasslands | Not Threatened | I |
| *Epilobium montanum* | 124 | -0.0003 | NA | NA | 82 | 0.0015 | <.001 | 0.18 | 914 | -0.0003 | <.001 | -0.12 | Dry to moderately moist forests | Not Threatened | I |
| *Scabiosa columbaria* agg. | 17 | -0.0003 | NA | NA | NA | NA | NA | NA | 2504 | -0.0035 | <.001 | -1.63 | Heaths, inland dunes and semi-natural grasslands | Not Threatened | I |
| *Platanthera bifolia* s. l. | 19 | -0.0003 | <.001 | -0.07 | NA | NA | NA | NA | 510 | -0.0010 | <.001 | -0.48 | Heaths, inland dunes and semi-natural grasslands | Endangered | I |
| *Acinos arvensis* | 8 | -0.0003 | NA | NA | NA | NA | NA | NA | 460 | -0.0008 | <.001 | -0.36 | Heaths, inland dunes and semi-natural grasslands | Near Threatened | I |
| *Silene nutans* | 8 | -0.0003 | NA | NA | NA | NA | NA | NA | 653 | -0.0006 | <.001 | -0.26 | Heaths, inland dunes and semi-natural grasslands | Not Threatened | I |
| *Lathyrus linifolius* | 16 | -0.0003 | <.001 | -0.06 | NA | NA | NA | NA | 344 | -0.0002 | <.001 | -0.09 | Heaths, inland dunes and semi-natural grasslands | Near Threatened | I |
| *Serratula tinctoria* s. l. | 4 | -0.0003 | NA | NA | NA | NA | NA | NA | 92 | -0.0002 | <.001 | -0.07 | None | Endangered | I |
| *Cynoglossum officinale* | 34 | -0.0003 | NA | NA | NA | NA | NA | NA | 85 | -0.0001 | <.001 | -0.03 | Coastal and marine habitats | Near Threatened | I |
| *Botrychium lunaria* | 6 | -0.0003 | NA | NA | NA | NA | NA | NA | 16 | 0.0000 | <.001 | -0.01 | None | Endangered | I |
| *Oenanthe lachenalii* | 26 | -0.0003 | <.001 | -0.07 | NA | NA | NA | NA | 13 | 0.0000 | <.001 | -0.01 | Coastal and marine habitats | Endangered | I |
| *Samolus valerandi* | 32 | -0.0003 | <.001 | -0.06 | NA | NA | NA | NA | 4 | 0.0000 | NA | NA | Coastal and marine habitats | Highly Endangered | I |
| *Scleranthus perennis* | 18 | -0.0003 | <.001 | -0.06 | NA | NA | NA | NA | 10 | 0.0000 | NA | NA | Heaths, inland dunes and semi-natural grasslands | Near Threatened | I |
| *Turritis glabra* | 11 | -0.0003 | NA | NA | NA | NA | NA | NA | 34 | 0.0000 | <.001 | -0.02 | None | Not Threatened | I |
| *Bromus ramosus* agg. | 34 | -0.0003 | NA | NA | NA | NA | NA | NA | 483 | 0.0002 | <.001 | 0.09 | Dry to moderately moist forests | Not Threatened | I |
| *Vicia tetrasperma* agg. | 42 | -0.0002 | NA | NA | 51 | -0.0040 | <.001 | -0.49 | 78 | -0.0001 | <.001 | -0.04 | Mesic grasslands | Not Threatened | I |
| *Myosotis arvensis* | 32 | -0.0002 | NA | NA | 52 | -0.0038 | <.001 | -0.46 | 203 | -0.0003 | <.001 | -0.15 | Farmland | Not Threatened | A |
| *Trifolium hybridum* | 14 | -0.0002 | NA | NA | 22 | -0.0024 | <.001 | -0.30 | 46 | -0.0001 | <.001 | -0.04 | Ruderal, fringe and tall forb communities, clearings | Not Evaluated | I |
| *Geranium pusillum* | 32 | -0.0002 | NA | NA | 12 | -0.0021 | <.001 | -0.26 | 58 | -0.0001 | <.001 | -0.03 | Mesic grasslands | Not Threatened | A |
| *Berteroa incana* | 11 | -0.0002 | NA | NA | 13 | -0.0016 | <.001 | -0.20 | 32 | -0.0001 | NA | NA | Ruderal, fringe and tall forb communities, clearings | Not Evaluated | N |
| *Senecio sylvaticus* | 23 | -0.0002 | <.001 | -0.04 | 43 | -0.0016 | <.001 | -0.20 | 14 | 0.0000 | NA | NA | None | Not Threatened | I |
| *Spergularia rubra* | 16 | -0.0002 | NA | NA | 14 | -0.0014 | <.001 | -0.17 | 18 | -0.0001 | <.001 | -0.03 | None | Not Threatened | A |
| *Medicago sativa* agg. | 8 | -0.0002 | NA | NA | 8 | -0.0011 | <.001 | -0.13 | 1776 | -0.0015 | <.001 | -0.69 | Heaths, inland dunes and semi-natural grasslands | Not Threatened | I |
| *Atriplex prostrata* agg. | 314 | -0.0002 | NA | NA | 13 | -0.0010 | <.001 | -0.12 | 6 | 0.0000 | NA | NA | Coastal and marine habitats | Not Threatened | I |
| *Sanguisorba minor* s. l. | 12 | -0.0002 | NA | NA | 6 | -0.0007 | NA | NA | 4481 | -0.0051 | <.001 | -2.36 | Heaths, inland dunes and semi-natural grasslands | Not Threatened | I |
| *Malva moschata* | 20 | -0.0002 | NA | NA | 8 | -0.0007 | NA | NA | 204 | -0.0001 | <.001 | -0.05 | None | Not Threatened | A |
| *Torilis japonica* agg. | 84 | -0.0002 | NA | NA | 46 | -0.0005 | NA | NA | 1256 | 0.0004 | <.001 | 0.17 | Scrubs, copses and field hedges | Not Threatened | I |
| *Cichorium intybus* | 10 | -0.0002 | NA | NA | 4 | -0.0004 | NA | NA | 442 | -0.0004 | <.001 | -0.18 | Heaths, inland dunes and semi-natural grasslands | Not Threatened | A |
| *Carduus nutans* agg. | 8 | -0.0002 | NA | NA | 4 | -0.0003 | NA | NA | 147 | -0.0002 | <.001 | -0.08 | Heaths, inland dunes and semi-natural grasslands | Not Threatened | A |
| *Rosa dumalis* agg. | 12 | -0.0002 | NA | NA | 1 | -0.0002 | NA | NA | 934 | -0.0003 | <.001 | -0.14 | Scrubs, copses and field hedges | Not Threatened | I |
| *Allium vineale* s. l. | 22 | -0.0002 | NA | NA | 2 | -0.0002 | NA | NA | 275 | -0.0001 | <.001 | -0.06 | None | Not Threatened | I |
| *Myosotis ramosissima* | 16 | -0.0002 | NA | NA | 2 | -0.0002 | NA | NA | 42 | -0.0001 | <.001 | -0.06 | Heaths, inland dunes and semi-natural grasslands | Not Threatened | A |
| *Pimpinella major* | 18 | -0.0002 | NA | NA | 2 | -0.0001 | NA | NA | 420 | -0.0006 | <.001 | -0.27 | Moist to wet grasslands | Not Threatened | I |
| *Selinum carvifolia* | 8 | -0.0002 | <.001 | -0.04 | 1 | -0.0001 | NA | NA | 128 | -0.0003 | <.001 | -0.14 | Moist to wet grasslands | Near Threatened | I |
| *Isolepis setacea* | 16 | -0.0002 | <.001 | -0.05 | 2 | -0.0001 | NA | NA | 38 | -0.0001 | <.001 | -0.03 | None | Near Threatened | I |
| *Myosotis sylvatica* agg. | 16 | -0.0002 | NA | NA | 1 | -0.0001 | NA | NA | 94 | -0.0001 | <.001 | -0.04 | None | Not Threatened | I |
| *Valerianella locusta* | 28 | -0.0002 | NA | NA | 1 | -0.0001 | NA | NA | 72 | -0.0001 | <.001 | -0.03 | Coastal and marine habitats | Not Threatened | A |
| *Alopecurus aequalis* | 18 | -0.0002 | NA | NA | 2 | -0.0001 | NA | NA | 30 | 0.0000 | <.001 | -0.02 | None | Not Threatened | I |
| *Catabrosa aquatica* | 12 | -0.0002 | <.001 | -0.05 | 1 | -0.0001 | NA | NA | 3 | 0.0000 | NA | NA | None | Highly Endangered | I |
| *Pulicaria dysenterica* | 64 | -0.0002 | NA | NA | 1 | 0.0000 | NA | NA | 234 | -0.0003 | <.001 | -0.12 | Fresh water vegetation, springs and reeds | Near Threatened | I |
| *Corydalis cava* | 98 | -0.0002 | NA | NA | 1 | 0.0001 | NA | NA | 252 | -0.0002 | <.001 | -0.09 | Dry to moderately moist forests | Not Threatened | I |
| *Scrophularia umbrosa* | 141 | -0.0002 | NA | NA | 6 | 0.0002 | NA | NA | 868 | -0.0004 | <.001 | -0.17 | Fresh water vegetation, springs and reeds | Not Threatened | I |
| *Lamium maculatum* | 116 | -0.0002 | NA | NA | 6 | 0.0002 | NA | NA | 3423 | 0.0016 | <.001 | 0.74 | Scrubs, copses and field hedges | Not Threatened | I |
| *Alnus incana* | 1186 | -0.0002 | NA | NA | 104 | 0.0024 | <.001 | 0.30 | 1870 | 0.0002 | NA | NA | Moist to wet forests | Not Threatened | I |
| *Valeriana officinalis* agg. | 1428 | -0.0002 | NA | NA | 138 | 0.0033 | <.001 | 0.40 | 6130 | -0.0006 | <.001 | -0.29 | Fresh water vegetation, springs and reeds | Not Threatened | I |
| *Viburnum opulus* | 1026 | -0.0002 | NA | NA | 94 | 0.0045 | <.001 | 0.55 | 7326 | 0.0018 | <.001 | 0.83 | Moist to wet forests | Not Threatened | I |
| *Cirsium acaulon* | 8 | -0.0002 | NA | NA | NA | NA | NA | NA | 1204 | -0.0017 | <.001 | -0.80 | Heaths, inland dunes and semi-natural grasslands | Near Threatened | I |
| *Ajuga genevensis* | 7 | -0.0002 | NA | NA | NA | NA | NA | NA | 540 | -0.0008 | <.001 | -0.37 | Heaths, inland dunes and semi-natural grasslands | Near Threatened | I |
| *Pinguicula vulgaris* | 6 | -0.0002 | NA | NA | NA | NA | NA | NA | 228 | -0.0007 | <.001 | -0.31 | Bogs, transition mires, marshes and fens | Endangered | I |
| *Campanula patula* | 9 | -0.0002 | NA | NA | NA | NA | NA | NA | 518 | -0.0005 | <.001 | -0.25 | Heaths, inland dunes and semi-natural grasslands | Near Threatened | I |
| *Neottia nidus-avis* | 11 | -0.0002 | NA | NA | NA | NA | NA | NA | 384 | -0.0004 | <.001 | -0.17 | Dry to moderately moist forests | Not Threatened | I |
| *Antennaria dioica* | 4 | -0.0002 | NA | NA | NA | NA | NA | NA | 112 | -0.0003 | <.001 | -0.14 | Heaths, inland dunes and semi-natural grasslands | Endangered | I |
| *Cuscuta epithymum* agg. | 6 | -0.0002 | NA | NA | NA | NA | NA | NA | 94 | -0.0001 | <.001 | -0.06 | Heaths, inland dunes and semi-natural grasslands | Endangered | I |
| *Hypericum montanum* | 6 | -0.0002 | NA | NA | NA | NA | NA | NA | 168 | -0.0001 | <.001 | -0.07 | Dry to moderately moist forests | Near Threatened | I |
| *Peucedanum oreoselinum* | 8 | -0.0002 | NA | NA | NA | NA | NA | NA | 114 | -0.0001 | <.001 | -0.05 | Heaths, inland dunes and semi-natural grasslands | Near Threatened | I |
| *Trifolium alpestre* | 4 | -0.0002 | NA | NA | NA | NA | NA | NA | 96 | -0.0001 | <.001 | -0.04 | None | Near Threatened | I |
| *Epipactis purpurata* | 10 | -0.0002 | <.001 | -0.05 | NA | NA | NA | NA | 48 | 0.0000 | <.001 | -0.02 | None | Near Threatened | I |
| *Solidago canadensis* | 90 | -0.0001 | NA | NA | 174 | -0.0045 | <.001 | -0.56 | 1306 | 0.0003 | <.001 | 0.13 | Scrubs, copses and field hedges | Not Evaluated | N |
| *Matricaria discoidea* | 30 | -0.0001 | NA | NA | 46 | -0.0036 | <.001 | -0.44 | 11 | 0.0000 | <.001 | -0.01 | Mesic grasslands | Not Evaluated | N |
| *Senecio inaequidens* | 40 | -0.0001 | NA | NA | 22 | -0.0029 | <.001 | -0.35 | 16 | 0.0000 | NA | NA | Coastal and marine habitats | Not Evaluated | N |
| *Symphytum officinale* agg. | 178 | -0.0001 | NA | NA | 143 | -0.0028 | <.001 | -0.34 | 1192 | -0.0006 | <.001 | -0.26 | Fresh water vegetation, springs and reeds | Not Threatened | I |
| *Sonchus oleraceus* | 36 | -0.0001 | NA | NA | 42 | -0.0025 | <.001 | -0.31 | 82 | 0.0000 | NA | NA | Farmland | Not Threatened | I |
| *Rorippa sylvestris* | 34 | -0.0001 | NA | NA | 57 | -0.0024 | <.001 | -0.29 | 15 | 0.0000 | NA | NA | None | Not Threatened | I |
| *Solidago gigantea* | 52 | -0.0001 | NA | NA | 191 | -0.0024 | NA | NA | 2103 | 0.0011 | <.001 | 0.50 | Fresh water vegetation, springs and reeds | Not Evaluated | N |
| *Solanum nigrum* s. l. | 34 | -0.0001 | NA | NA | 23 | -0.0018 | <.001 | -0.22 | 60 | 0.0001 | <.001 | 0.03 | None | Not Threatened | A |
| *Trientalis europaea* | 650 | -0.0001 | NA | NA | 136 | -0.0016 | NA | NA | 18 | 0.0000 | <.001 | -0.02 | Dry to moderately moist forests | Not Threatened | I |
| *Atriplex patula* | 8 | -0.0001 | NA | NA | 20 | -0.0013 | <.001 | -0.16 | 14 | 0.0000 | NA | NA | Ruderal, fringe and tall forb communities, clearings | Not Threatened | A |
| *Fallopia convolvulus* | 17 | -0.0001 | NA | NA | 18 | -0.0013 | <.001 | -0.16 | 90 | 0.0000 | NA | NA | Farmland | Not Threatened | A |
| *Papaver rhoeas* | 4 | -0.0001 | NA | NA | 8 | -0.0011 | NA | NA | 176 | -0.0002 | <.001 | -0.08 | None | Not Threatened | A |
| *Lactuca serriola* | 18 | -0.0001 | NA | NA | 10 | -0.0011 | NA | NA | 688 | 0.0002 | <.001 | 0.11 | Scrubs, copses and field hedges | Not Threatened | I |
| *Reseda lutea* | 6 | -0.0001 | NA | NA | 10 | -0.0009 | NA | NA | 221 | -0.0003 | <.001 | -0.13 | Heaths, inland dunes and semi-natural grasslands | Not Threatened | A |
| *Apera spica-venti* | 6 | -0.0001 | NA | NA | 8 | -0.0008 | <.001 | -0.10 | 12 | 0.0000 | NA | NA | Farmland | Not Threatened | I |
| *Saponaria officinalis* | 12 | -0.0001 | NA | NA | 19 | -0.0008 | NA | NA | 450 | 0.0002 | <.001 | 0.09 | Scrubs, copses and field hedges | Not Threatened | I |
| *Euphorbia cyparissias* | 11 | -0.0001 | NA | NA | 8 | -0.0007 | NA | NA | 8264 | -0.0047 | <.001 | -2.16 | Heaths, inland dunes and semi-natural grasslands | Not Threatened | I |
| *Persicaria mitis* | 22 | -0.0001 | NA | NA | 26 | -0.0006 | <.001 | -0.07 | 28 | 0.0000 | NA | NA | None | Not Threatened | I |
| *Dianthus carthusianorum* agg. | 8 | -0.0001 | NA | NA | 2 | -0.0005 | NA | NA | 2140 | -0.0029 | <.001 | -1.33 | Heaths, inland dunes and semi-natural grasslands | Near Threatened | I |
| *Hieracium lachenalii* | 46 | -0.0001 | NA | NA | 20 | -0.0005 | <.001 | -0.06 | 164 | -0.0002 | <.001 | -0.07 | Heaths, inland dunes and semi-natural grasslands | Not Threatened | I |
| *Anagallis arvensis* | 4 | -0.0001 | NA | NA | 8 | -0.0005 | NA | NA | 26 | -0.0001 | <.001 | -0.02 | None | Not Threatened | A |
| *Thlaspi arvense* | 12 | -0.0001 | NA | NA | 3 | -0.0005 | NA | NA | 32 | -0.0001 | <.001 | -0.03 | None | Not Threatened | A |
| *Sedum sexangulare* | 5 | -0.0001 | NA | NA | 2 | -0.0004 | NA | NA | 304 | -0.0002 | <.001 | -0.11 | Heaths, inland dunes and semi-natural grasslands | Not Threatened | I |
| *Cardamine hirsuta* | 62 | -0.0001 | NA | NA | 2 | -0.0004 | NA | NA | 66 | -0.0001 | <.001 | -0.04 | None | Not Threatened | I |
| *Eryngium campestre* | 5 | -0.0001 | NA | NA | 2 | -0.0004 | NA | NA | 72 | -0.0001 | <.001 | -0.04 | Heaths, inland dunes and semi-natural grasslands | Near Threatened | I |
| *Plantago media* | 24 | -0.0001 | NA | NA | 6 | -0.0003 | NA | NA | 1934 | -0.0027 | <.001 | -1.26 | Heaths, inland dunes and semi-natural grasslands | Not Threatened | I |
| *Hypericum tetrapterum* | 182 | -0.0001 | NA | NA | 24 | -0.0003 | NA | NA | 586 | -0.0008 | <.001 | -0.38 | Moist to wet grasslands | Not Threatened | I |
| *Centaurea paniculata* agg. | 7 | -0.0001 | NA | NA | 2 | -0.0002 | NA | NA | 83 | -0.0001 | <.001 | -0.05 | Heaths, inland dunes and semi-natural grasslands | Not Threatened | I |
| *Melilotus altissimus* | 15 | -0.0001 | NA | NA | 4 | -0.0002 | NA | NA | 43 | 0.0000 | <.001 | -0.02 | None | Not Threatened | I |
| *Salvia pratensis* | 2 | -0.0001 | NA | NA | 1 | -0.0001 | NA | NA | 3322 | -0.0038 | <.001 | -1.75 | Heaths, inland dunes and semi-natural grasslands | Near Threatened | I |
| *Geranium pratense* | 14 | -0.0001 | NA | NA | 8 | -0.0001 | NA | NA | 2230 | -0.0007 | <.001 | -0.32 | Scrubs, copses and field hedges | Not Threatened | I |
| *Campanula persicifolia* | 5 | -0.0001 | NA | NA | 1 | -0.0001 | NA | NA | 984 | -0.0006 | <.001 | -0.30 | Dry to moderately moist forests | Not Threatened | I |
| *Centaurea nigra* s. l. | 8 | -0.0001 | NA | NA | 1 | -0.0001 | NA | NA | 618 | -0.0006 | <.001 | -0.28 | Heaths, inland dunes and semi-natural grasslands | Not Threatened | I |
| *Crepis biennis* | 15 | -0.0001 | NA | NA | 2 | -0.0001 | NA | NA | 220 | -0.0002 | <.001 | -0.09 | None | Not Threatened | A |
| *Filipendula vulgaris* | 8 | -0.0001 | NA | NA | 1 | -0.0001 | NA | NA | 154 | -0.0002 | <.001 | -0.11 | Heaths, inland dunes and semi-natural grasslands | Endangered | I |
| *Verbascum densiflorum* | 4 | -0.0001 | NA | NA | 6 | -0.0001 | NA | NA | 71 | -0.0001 | <.001 | -0.05 | None | Not Threatened | I |
| *Bromus erectus* | 4 | -0.0001 | NA | NA | 2 | 0.0000 | NA | NA | 5444 | -0.0053 | <.001 | -2.46 | Heaths, inland dunes and semi-natural grasslands | Not Threatened | I |
| *Helictotrichon pubescens* | 8 | -0.0001 | NA | NA | 1 | 0.0000 | NA | NA | 721 | -0.0009 | <.001 | -0.41 | Heaths, inland dunes and semi-natural grasslands | Not Threatened | I |
| *Aconitum napellus* agg. | 6 | -0.0001 | NA | NA | 1 | 0.0000 | NA | NA | 253 | -0.0003 | <.001 | -0.14 | Ruderal, fringe and tall forb communities, clearings | Not Threatened | I |
| *Petrorhagia prolifera* | 2 | -0.0001 | NA | NA | 1 | 0.0000 | NA | NA | 105 | -0.0002 | <.001 | -0.08 | Heaths, inland dunes and semi-natural grasslands | Not Threatened | I |
| *Iris sibirica* | 2 | -0.0001 | NA | NA | 2 | 0.0000 | NA | NA | 52 | -0.0001 | <.001 | -0.04 | None | Endangered | I |
| *Lathraea squamaria* | 36 | -0.0001 | NA | NA | 2 | 0.0000 | NA | NA | 34 | 0.0000 | <.001 | -0.02 | None | Not Threatened | I |
| *Mimulus guttatus* | 4 | -0.0001 | NA | NA | 2 | 0.0000 | NA | NA | 57 | 0.0000 | <.001 | -0.02 | None | Not Evaluated | N |
| *Primula veris* | 52 | -0.0001 | NA | NA | 1 | 0.0001 | NA | NA | 2329 | -0.0025 | <.001 | -1.14 | Heaths, inland dunes and semi-natural grasslands | Near Threatened | I |
| *Agrimonia procera* | 29 | -0.0001 | NA | NA | 10 | 0.0001 | NA | NA | 84 | -0.0001 | <.001 | -0.04 | None | Not Threatened | I |
| *Senecio paludosus* | 5 | -0.0001 | NA | NA | 10 | 0.0001 | NA | NA | 107 | -0.0001 | <.001 | -0.06 | Fresh water vegetation, springs and reeds | Endangered | I |
| *Rosa canina* agg. | 693 | -0.0001 | NA | NA | 277 | 0.0026 | NA | NA | 17586 | 0.0043 | <.001 | 2.01 | Scrubs, copses and field hedges | Not Threatened | I |
| *Lonicera xylosteum* | 189 | -0.0001 | NA | NA | 46 | 0.0035 | <.001 | 0.43 | 11186 | 0.0030 | <.001 | 1.39 | Scrubs, copses and field hedges | Not Threatened | I |
| *Convallaria majalis* | 743 | -0.0001 | NA | NA | 92 | 0.0043 | <.001 | 0.52 | 1996 | 0.0000 | NA | NA | Dry to moderately moist forests | Not Threatened | I |
| *Potentilla verna* agg. | 2 | -0.0001 | NA | NA | NA | NA | NA | NA | 2544 | -0.0033 | <.001 | -1.54 | Heaths, inland dunes and semi-natural grasslands | Near Threatened | I |
| *Carex caryophyllea* | 10 | -0.0001 | NA | NA | NA | NA | NA | NA | 1242 | -0.0015 | <.001 | -0.71 | Heaths, inland dunes and semi-natural grasslands | Near Threatened | I |
| *Vaccinium vitis-idaea* | 13 | -0.0001 | <.001 | -0.03 | NA | NA | NA | NA | 670 | -0.0015 | <.001 | -0.69 | Heaths, inland dunes and semi-natural grasslands | Not Threatened | I |
| *Anthericum ramosum* | 2 | -0.0001 | NA | NA | NA | NA | NA | NA | 860 | -0.0014 | <.001 | -0.65 | Heaths, inland dunes and semi-natural grasslands | Near Threatened | I |
| *Pulsatilla vulgaris* s. l. | 2 | -0.0001 | NA | NA | NA | NA | NA | NA | 672 | -0.0014 | <.001 | -0.64 | Heaths, inland dunes and semi-natural grasslands | Endangered | I |
| *Euphrasia officinalis* agg. | 9 | -0.0001 | NA | NA | NA | NA | NA | NA | 873 | -0.0013 | <.001 | -0.59 | Heaths, inland dunes and semi-natural grasslands | Endangered | I |
| *Geranium sanguineum* | 8 | -0.0001 | NA | NA | NA | NA | NA | NA | 886 | -0.0012 | <.001 | -0.57 | Heaths, inland dunes and semi-natural grasslands | Near Threatened | I |
| *Carex montana* | 12 | -0.0001 | NA | NA | NA | NA | NA | NA | 1214 | -0.0011 | <.001 | -0.51 | Dry to moderately moist forests | Not Threatened | I |
| *Potentilla heptaphylla* | 4 | -0.0001 | NA | NA | NA | NA | NA | NA | 790 | -0.0011 | <.001 | -0.50 | Heaths, inland dunes and semi-natural grasslands | Near Threatened | I |
| *Campanula glomerata* | 8 | -0.0001 | NA | NA | NA | NA | NA | NA | 618 | -0.0009 | <.001 | -0.40 | Heaths, inland dunes and semi-natural grasslands | Endangered | I |
| *Viola hirta* | 4 | -0.0001 | NA | NA | NA | NA | NA | NA | 1829 | -0.0009 | <.001 | -0.42 | Heaths, inland dunes and semi-natural grasslands | Not Threatened | I |
| *Carex pulicaris* | 10 | -0.0001 | NA | NA | NA | NA | NA | NA | 260 | -0.0006 | <.001 | -0.30 | Moist to wet grasslands | Highly Endangered | I |
| *Pinus mugo* agg. | 18 | -0.0001 | NA | NA | NA | NA | NA | NA | 180 | -0.0005 | <.001 | -0.22 | Bogs, transition mires, marshes and fens | Not Threatened | I |
| *Arabis hirsuta* agg. | 2 | -0.0001 | NA | NA | NA | NA | NA | NA | 318 | -0.0004 | <.001 | -0.20 | Heaths, inland dunes and semi-natural grasslands | Not Threatened | I |
| *Cruciata laevipes* | 3 | -0.0001 | NA | NA | NA | NA | NA | NA | 466 | -0.0004 | <.001 | -0.18 | Scrubs, copses and field hedges | Not Threatened | I |
| *Geranium columbinum* | 4 | -0.0001 | NA | NA | NA | NA | NA | NA | 267 | -0.0003 | <.001 | -0.15 | None | Not Threatened | A |
| *Seseli libanotis* | 4 | -0.0001 | NA | NA | NA | NA | NA | NA | 140 | -0.0003 | <.001 | -0.13 | Heaths, inland dunes and semi-natural grasslands | Endangered | I |
| *Montia fontana* agg. | 6 | -0.0001 | NA | NA | NA | NA | NA | NA | 110 | -0.0002 | <.001 | -0.11 | None | Near Threatened | I |
| *Orchis morio* | 2 | -0.0001 | NA | NA | NA | NA | NA | NA | 120 | -0.0002 | <.001 | -0.10 | Heaths, inland dunes and semi-natural grasslands | Highly Endangered | I |
| *Alyssum alyssoides* | 1 | -0.0001 | NA | NA | NA | NA | NA | NA | 67 | -0.0001 | <.001 | -0.05 | Heaths, inland dunes and semi-natural grasslands | Endangered | I |
| *Carex limosa* | 4 | -0.0001 | NA | NA | NA | NA | NA | NA | 28 | -0.0001 | <.001 | -0.05 | Bogs, transition mires, marshes and fens | Highly Endangered | I |
| *Consolida regalis* | 10 | -0.0001 | NA | NA | NA | NA | NA | NA | 54 | -0.0001 | <.001 | -0.04 | None | Endangered | A |
| *Genista germanica* | 1 | -0.0001 | NA | NA | NA | NA | NA | NA | 112 | -0.0001 | <.001 | -0.07 | Heaths, inland dunes and semi-natural grasslands | Endangered | I |
| *Rubus saxatilis* | 14 | -0.0001 | NA | NA | NA | NA | NA | NA | 156 | -0.0001 | <.001 | -0.06 | Dry to moderately moist forests | Near Threatened | I |
| *Carex dioica* | 3 | -0.0001 | NA | NA | NA | NA | NA | NA | 12 | 0.0000 | <.001 | -0.02 | None | Highly Endangered | I |
| *Euphorbia stricta* | 9 | -0.0001 | NA | NA | NA | NA | NA | NA | 52 | 0.0000 | <.001 | -0.02 | None | Not Threatened | I |
| *Hypochaeris maculata* | 2 | -0.0001 | NA | NA | NA | NA | NA | NA | 20 | 0.0000 | <.001 | -0.02 | None | Highly Endangered | I |
| *Pyrola rotundifolia* | 6 | -0.0001 | NA | NA | NA | NA | NA | NA | 20 | 0.0000 | <.001 | -0.02 | None | Endangered | I |
| *Stellaria media* agg. | 342 | 0.0000 | NA | NA | 258 | -0.0048 | <.001 | -0.59 | 325 | -0.0001 | <.001 | -0.06 | Farmland | Not Threatened | NA |
| *Sisymbrium altissimum* | 2 | 0.0000 | NA | NA | 29 | -0.0026 | <.001 | -0.31 | 1 | 0.0000 | NA | NA | Ruderal, fringe and tall forb communities, clearings | Not Evaluated | N |
| *Matricaria chamomilla* | 21 | 0.0000 | NA | NA | 16 | -0.0025 | <.001 | -0.31 | 19 | 0.0000 | NA | NA | Mesic grasslands | Not Threatened | A |
| *Chenopodium polyspermum* | 3 | 0.0000 | NA | NA | 17 | -0.0020 | <.001 | -0.24 | 24 | 0.0000 | NA | NA | None | Not Threatened | I |
| *Erysimum cheiranthoides* | 2 | 0.0000 | NA | NA | 14 | -0.0014 | <.001 | -0.18 | 7 | 0.0000 | NA | NA | None | Not Threatened | I |
| *Galinsoga parviflora* | 4 | 0.0000 | NA | NA | 24 | -0.0014 | <.001 | -0.17 | 4 | 0.0000 | NA | NA | Anthropogenic | Not Evaluated | N |
| *Armoracia rusticana* | 8 | 0.0000 | NA | NA | 26 | -0.0013 | <.001 | -0.16 | 26 | 0.0000 | NA | NA | Ruderal, fringe and tall forb communities, clearings | Not Evaluated | N |
| *Lamium purpureum* s. str. | 37 | 0.0000 | NA | NA | 26 | -0.0010 | NA | NA | 333 | -0.0001 | <.001 | -0.04 | None | Not Threatened | A |
| *Verbascum thapsus* agg. | 11 | 0.0000 | NA | NA | 10 | -0.0010 | NA | NA | 206 | -0.0001 | <.001 | -0.06 | Heaths, inland dunes and semi-natural grasslands | Not Threatened | I |
| *Geranium dissectum* | 26 | 0.0000 | NA | NA | 10 | -0.0009 | <.001 | -0.11 | 132 | -0.0001 | NA | NA | Mesic grasslands | Not Threatened | A |
| *Lolium multiflorum* | 16 | 0.0000 | NA | NA | 10 | -0.0009 | <.001 | -0.11 | 107 | -0.0001 | <.001 | -0.05 | Moist to wet grasslands | Not Evaluated | N |
| *Galinsoga quadriradiata* s. str. | 1 | 0.0000 | NA | NA | 12 | -0.0009 | <.001 | -0.11 | 8 | 0.0000 | NA | NA | None | Not Evaluated | N |
| *Juncus tenageia* | 2 | 0.0000 | NA | NA | 23 | -0.0009 | <.001 | -0.10 | 2 | 0.0000 | NA | NA | None | Highly Endangered | I |
| *Sinapis arvensis* | 2 | 0.0000 | NA | NA | 6 | -0.0008 | NA | NA | 24 | 0.0000 | <.001 | -0.02 | None | Not Threatened | A |
| *Epilobium roseum* | 66 | 0.0000 | NA | NA | 26 | -0.0007 | <.001 | -0.09 | 93 | -0.0001 | NA | NA | None | Not Threatened | I |
| *Rumex pratensis* | 13 | 0.0000 | NA | NA | 17 | -0.0006 | <.001 | -0.07 | NA | NA | NA | NA | Mesic grasslands | Not Threatened | I |
| *Senecio erucifolius* | 8 | 0.0000 | NA | NA | 12 | -0.0005 | NA | NA | 1106 | -0.0010 | <.001 | -0.45 | Heaths, inland dunes and semi-natural grasslands | Not Threatened | I |
| *Bromus inermis* | 40 | 0.0000 | NA | NA | 27 | -0.0005 | NA | NA | 710 | 0.0004 | <.001 | 0.19 | Scrubs, copses and field hedges | Not Threatened | I |
| *Campanula rapunculoides* | 4 | 0.0000 | NA | NA | 12 | -0.0004 | NA | NA | 632 | -0.0003 | <.001 | -0.16 | None | Not Threatened | I |
| *Veronica persica* | 5 | 0.0000 | NA | NA | 4 | -0.0004 | NA | NA | 70 | -0.0001 | <.001 | -0.04 | None | Not Evaluated | N |
| *Dipsacus fullonum* | 20 | 0.0000 | NA | NA | 11 | -0.0003 | NA | NA | 520 | -0.0001 | <.001 | -0.06 | Scrubs, copses and field hedges | Not Threatened | A |
| *Erigeron annuus* | 11 | 0.0000 | NA | NA | 1 | -0.0003 | NA | NA | 430 | 0.0002 | <.001 | 0.09 | None | Not Evaluated | N |
| *Chaenorhinum minus* | 1 | 0.0000 | NA | NA | 1 | -0.0002 | NA | NA | 34 | -0.0001 | <.001 | -0.02 | None | Not Threatened | A |
| *Tragopogon dubius* | 1 | 0.0000 | NA | NA | 1 | -0.0002 | NA | NA | 73 | -0.0001 | <.001 | -0.03 | None | Not Threatened | I |
| *Polygala serpyllifolia* | 3 | 0.0000 | NA | NA | 1 | -0.0001 | NA | NA | 410 | -0.0007 | <.001 | -0.34 | Heaths, inland dunes and semi-natural grasslands | Endangered | I |
| *Carduus acanthoides* | 2 | 0.0000 | NA | NA | 4 | -0.0001 | NA | NA | 132 | -0.0002 | <.001 | -0.07 | None | Not Threatened | A |
| *Alopecurus myosuroides* | 3 | 0.0000 | NA | NA | 2 | -0.0001 | NA | NA | 38 | -0.0001 | <.001 | -0.03 | None | Not Threatened | A |
| *Malva alcea* | 4 | 0.0000 | NA | NA | 1 | -0.0001 | NA | NA | 130 | -0.0001 | <.001 | -0.04 | None | Not Threatened | A |
| *Saxifraga tridactylites* | 1 | 0.0000 | NA | NA | 1 | -0.0001 | NA | NA | 34 | -0.0001 | <.001 | -0.03 | None | Not Threatened | I |
| *Anthemis arvensis* | 4 | 0.0000 | NA | NA | 1 | -0.0001 | NA | NA | 12 | 0.0000 | <.001 | -0.01 | None | Near Threatened | A |
| *Setaria pumila* | 1 | 0.0000 | NA | NA | 2 | -0.0001 | NA | NA | 32 | 0.0000 | <.001 | 0.02 | None | Not Threatened | A |
| *Mentha spicata* agg. | 25 | 0.0000 | NA | NA | 6 | 0.0000 | NA | NA | 2325 | -0.0015 | <.001 | -0.71 | Fresh water vegetation, springs and reeds | Not Threatened | NA |
| *Sorbus aria* agg. | 14 | 0.0000 | NA | NA | 4 | 0.0000 | NA | NA | 3390 | -0.0015 | <.001 | -0.71 | Scrubs, copses and field hedges | Not Threatened | I |
| *Inula salicina* | 2 | 0.0000 | NA | NA | 1 | 0.0000 | NA | NA | 379 | -0.0006 | <.001 | -0.28 | Heaths, inland dunes and semi-natural grasslands | Near Threatened | I |
| *Silaum silaus* | 1 | 0.0000 | NA | NA | 1 | 0.0000 | NA | NA | 345 | -0.0006 | <.001 | -0.29 | Moist to wet grasslands | Near Threatened | I |
| *Aquilegia vulgaris* agg. | 2 | 0.0000 | NA | NA | 2 | 0.0000 | NA | NA | 578 | -0.0005 | <.001 | -0.24 | Heaths, inland dunes and semi-natural grasslands | Near Threatened | I |
| *Picris hieracioides* s. l. | 5 | 0.0000 | NA | NA | 4 | 0.0000 | NA | NA | 623 | -0.0004 | <.001 | -0.18 | Heaths, inland dunes and semi-natural grasslands | Not Threatened | I |
| *Melampyrum sylvaticum* | 8 | 0.0000 | NA | NA | 4 | 0.0000 | NA | NA | 224 | -0.0003 | <.001 | -0.12 | None | Not Threatened | I |
| *Rumex aquaticus* | 16 | 0.0000 | NA | NA | 1 | 0.0000 | NA | NA | 89 | -0.0002 | <.001 | -0.07 | Fresh water vegetation, springs and reeds | Near Threatened | I |
| *Lathyrus tuberosus* | 1 | 0.0000 | NA | NA | 1 | 0.0000 | NA | NA | 114 | -0.0001 | <.001 | -0.03 | None | Not Threatened | I |
| *Dipsacus pilosus* | 1 | 0.0000 | NA | NA | 1 | 0.0000 | NA | NA | 160 | 0.0001 | <.001 | 0.07 | Moist to wet forests | Not Threatened | I |
| *Sambucus ebulus* | 6 | 0.0000 | NA | NA | 1 | 0.0000 | NA | NA | 644 | 0.0003 | <.001 | 0.12 | Scrubs, copses and field hedges | Not Threatened | I |
| *Anemone ranunculoides* | 109 | 0.0000 | NA | NA | 1 | 0.0001 | NA | NA | 224 | -0.0002 | <.001 | -0.10 | Dry to moderately moist forests | Not Threatened | I |
| *Ornithogalum umbellatum* agg. | 6 | 0.0000 | NA | NA | 2 | 0.0001 | NA | NA | 35 | 0.0000 | <.001 | -0.02 | None | Not Threatened | A |
| *Acer negundo* | 2 | 0.0000 | NA | NA | 4 | 0.0001 | NA | NA | 58 | 0.0001 | <.001 | 0.04 | None | Not Evaluated | N |
| *Prunus mahaleb* | 2 | 0.0000 | NA | NA | 2 | 0.0001 | NA | NA | 180 | 0.0001 | <.001 | 0.06 | Scrubs, copses and field hedges | Not Threatened | I |
| *Bryonia dioica* | 5 | 0.0000 | NA | NA | 52 | 0.0001 | NA | NA | 1393 | 0.0008 | <.001 | 0.37 | Scrubs, copses and field hedges | Not Threatened | I |
| *Berberis vulgaris* | 2 | 0.0000 | NA | NA | 6 | 0.0002 | NA | NA | 1307 | -0.0009 | <.001 | -0.44 | Heaths, inland dunes and semi-natural grasslands | Not Threatened | I |
| *Gagea lutea* | 73 | 0.0000 | NA | NA | 2 | 0.0002 | NA | NA | 82 | -0.0001 | <.001 | -0.04 | Dry to moderately moist forests | Not Threatened | I |
| *Fallopia baldschuanica* | 3 | 0.0000 | NA | NA | 5 | 0.0002 | NA | NA | 14 | 0.0000 | <.001 | 0.01 | None | Not Evaluated | N |
| *Rhus typhina* | 1 | 0.0000 | NA | NA | 3 | 0.0002 | NA | NA | 189 | 0.0002 | <.001 | 0.10 | Scrubs, copses and field hedges | Not Evaluated | N |
| *Pinus nigra* | 50 | 0.0000 | NA | NA | 5 | 0.0003 | NA | NA | 307 | -0.0002 | <.001 | -0.10 | Coastal and marine habitats | Not Evaluated | N |
| *Parthenocissus quinquefolia* agg. | 6 | 0.0000 | NA | NA | 6 | 0.0005 | NA | NA | 175 | 0.0002 | <.001 | 0.10 | None | Not Evaluated | N |
| *Luzula sylvatica* | 112 | 0.0000 | NA | NA | 6 | 0.0005 | NA | NA | 1540 | 0.0005 | <.001 | 0.25 | Dry to moderately moist forests | Not Threatened | I |
| *Rubus* sect. *Corylifolii* | 7 | 0.0000 | NA | NA | 16 | 0.0005 | NA | NA | 475 | 0.0005 | <.001 | 0.23 | Scrubs, copses and field hedges | Not Threatened | I |
| *Clematis vitalba* | 22 | 0.0000 | NA | NA | 20 | 0.0008 | NA | NA | 7872 | 0.0057 | <.001 | 2.65 | Scrubs, copses and field hedges | Not Threatened | I |
| *Prunus cerasifera* | 2 | 0.0000 | NA | NA | 9 | 0.0009 | <.001 | 0.11 | 736 | 0.0011 | <.001 | 0.49 | Scrubs, copses and field hedges | Not Evaluated | N |
| *Ribes alpinum* | 1 | 0.0000 | NA | NA | 12 | 0.0013 | <.001 | 0.16 | 709 | 0.0000 | NA | NA | Scrubs, copses and field hedges | Not Threatened | I |
| *Maianthemum bifolium* | 1324 | 0.0000 | NA | NA | 98 | 0.0017 | NA | NA | 390 | -0.0003 | <.001 | -0.14 | Dry to moderately moist forests | Not Threatened | I |
| *Lapsana communis* | 199 | 0.0000 | NA | NA | 64 | 0.0019 | <.001 | 0.23 | 2211 | 0.0018 | <.001 | 0.83 | Scrubs, copses and field hedges | Not Threatened | I |
| *Sambucus racemosa* | 71 | 0.0000 | NA | NA | 108 | 0.0021 | <.001 | 0.26 | 2203 | -0.0005 | <.001 | -0.24 | Scrubs, copses and field hedges | Not Threatened | I |
| *Scrophularia nodosa* | 768 | 0.0000 | NA | NA | 222 | 0.0034 | <.001 | 0.42 | 1472 | 0.0000 | NA | NA | Dry to moderately moist forests | Not Threatened | I |
| *Helianthemum nummularium* s. l. | 1 | 0.0000 | NA | NA | NA | NA | NA | NA | 2745 | -0.0040 | <.001 | -1.84 | Heaths, inland dunes and semi-natural grasslands | Near Threatened | I |
| *Onobrychis viciifolia* agg. | 2 | 0.0000 | NA | NA | NA | NA | NA | NA | 1368 | -0.0020 | <.001 | -0.92 | Heaths, inland dunes and semi-natural grasslands | Endangered | NA |
| *Betonica officinalis* | 2 | 0.0000 | NA | NA | NA | NA | NA | NA | 1201 | -0.0015 | <.001 | -0.69 | Heaths, inland dunes and semi-natural grasslands | Near Threatened | I |
| *Trisetum flavescens* | 8 | 0.0000 | NA | NA | NA | NA | NA | NA | 1372 | -0.0015 | <.001 | -0.68 | Heaths, inland dunes and semi-natural grasslands | Not Threatened | I |
| *Vincetoxicum hirundinaria* | 1 | 0.0000 | NA | NA | NA | NA | NA | NA | 1506 | -0.0014 | <.001 | -0.65 | Heaths, inland dunes and semi-natural grasslands | Not Threatened | I |
| *Hieracium lactucella* | 1 | 0.0000 | NA | NA | NA | NA | NA | NA | 684 | -0.0012 | <.001 | -0.57 | Moist to wet grasslands | Endangered | I |
| *Rhinanthus alectorolophus* | 8 | 0.0000 | NA | NA | NA | NA | NA | NA | 1088 | -0.0012 | <.001 | -0.56 | Heaths, inland dunes and semi-natural grasslands | Not Threatened | I |
| *Melampyrum arvense* | 2 | 0.0000 | NA | NA | NA | NA | NA | NA | 818 | -0.0008 | <.001 | -0.39 | Heaths, inland dunes and semi-natural grasslands | Endangered | I |
| *Geranium sylvaticum* | 4 | 0.0000 | NA | NA | NA | NA | NA | NA | 1142 | -0.0007 | <.001 | -0.31 | Moist to wet grasslands | Not Threatened | I |
| *Galium pusillum* agg. | 3 | 0.0000 | NA | NA | NA | NA | NA | NA | 442 | -0.0006 | <.001 | -0.30 | Heaths, inland dunes and semi-natural grasslands | Near Threatened | I |
| *Trifolium montanum* | 1 | 0.0000 | NA | NA | NA | NA | NA | NA | 349 | -0.0006 | <.001 | -0.26 | Heaths, inland dunes and semi-natural grasslands | Near Threatened | I |
| *Cephalanthera damasonium* | 12 | 0.0000 | NA | NA | NA | NA | NA | NA | 465 | -0.0004 | <.001 | -0.17 | Dry to moderately moist forests | Not Threatened | I |
| *Eriophorum latifolium* | 1 | 0.0000 | NA | NA | NA | NA | NA | NA | 155 | -0.0004 | <.001 | -0.19 | Moist to wet grasslands | Endangered | I |
| *Lycopodium annotinum* | 10 | 0.0000 | NA | NA | NA | NA | NA | NA | 248 | -0.0004 | <.001 | -0.17 | Moist to wet forests | Near Threatened | I |
| *Phleum phleoides* | 1 | 0.0000 | NA | NA | NA | NA | NA | NA | 274 | -0.0004 | <.001 | -0.19 | Heaths, inland dunes and semi-natural grasslands | Near Threatened | I |
| *Carum carvi* | 10 | 0.0000 | NA | NA | NA | NA | NA | NA | 191 | -0.0003 | <.001 | -0.15 | Moist to wet grasslands | Not Threatened | I |
| *Carex hostiana* | 1 | 0.0000 | NA | NA | NA | NA | NA | NA | 104 | -0.0002 | <.001 | -0.11 | Moist to wet grasslands | Highly Endangered | I |
| *Carex tomentosa* | 2 | 0.0000 | NA | NA | NA | NA | NA | NA | 180 | -0.0002 | <.001 | -0.11 | Moist to wet grasslands | Endangered | I |
| *Cotoneaster integerrimus* | 1 | 0.0000 | NA | NA | NA | NA | NA | NA | 117 | -0.0002 | <.001 | -0.07 | Dry to moderately moist forests | Near Threatened | I |
| *Leucojum vernum* | 4 | 0.0000 | NA | NA | NA | NA | NA | NA | 161 | -0.0002 | <.001 | -0.07 | Dry to moderately moist forests | Near Threatened | I |
| *Melampyrum cristatum* | 1 | 0.0000 | NA | NA | NA | NA | NA | NA | 168 | -0.0002 | <.001 | -0.08 | Heaths, inland dunes and semi-natural grasslands | Endangered | I |
| *Phyteuma nigrum* | 1 | 0.0000 | NA | NA | NA | NA | NA | NA | 195 | -0.0002 | <.001 | -0.08 | Moist to wet grasslands | Near Threatened | I |
| *Ranunculus polyanthemos* s. l. | 6 | 0.0000 | NA | NA | NA | NA | NA | NA | 178 | -0.0002 | <.001 | -0.09 | Heaths, inland dunes and semi-natural grasslands | Near Threatened | I |
| *Thalictrum aquilegiifolium* | 2 | 0.0000 | NA | NA | NA | NA | NA | NA | 282 | -0.0002 | <.001 | -0.09 | None | Not Threatened | I |
| *Trichophorum alpinum* | 1 | 0.0000 | NA | NA | NA | NA | NA | NA | 46 | -0.0002 | <.001 | -0.07 | Bogs, transition mires, marshes and fens | Endangered | I |
| *Euphorbia palustris* | 1 | 0.0000 | NA | NA | NA | NA | NA | NA | 61 | -0.0001 | <.001 | -0.02 | Fresh water vegetation, springs and reeds | Endangered | I |
| *Gymnocarpium robertianum* | 1 | 0.0000 | NA | NA | NA | NA | NA | NA | 58 | -0.0001 | <.001 | -0.03 | Dry to moderately moist forests | Not Threatened | I |
| *Hypopitys monotropa* agg. | 2 | 0.0000 | NA | NA | NA | NA | NA | NA | 74 | -0.0001 | <.001 | -0.05 | None | Not Threatened | I |
| *Lepidium campestre* | 1 | 0.0000 | NA | NA | NA | NA | NA | NA | 47 | -0.0001 | <.001 | -0.03 | None | Not Threatened | A |
| *Primula farinosa* | 1 | 0.0000 | NA | NA | NA | NA | NA | NA | 40 | -0.0001 | <.001 | -0.06 | None | Endangered | I |
| *Salix myrsinifolia* | 2 | 0.0000 | NA | NA | NA | NA | NA | NA | 225 | -0.0001 | <.001 | -0.07 | None | Near Threatened | I |
| *Scheuchzeria palustris* | 3 | 0.0000 | NA | NA | NA | NA | NA | NA | 25 | -0.0001 | <.001 | -0.04 | Bogs, transition mires, marshes and fens | Highly Endangered | I |
| *Schoenus nigricans* | 1 | 0.0000 | NA | NA | NA | NA | NA | NA | 44 | -0.0001 | <.001 | -0.04 | None | Highly Endangered | I |
| *Thalictrum minus* agg. | 2 | 0.0000 | NA | NA | NA | NA | NA | NA | 69 | -0.0001 | <.001 | -0.05 | Heaths, inland dunes and semi-natural grasslands | Near Threatened | I |
| *Trifolium aureum* | 1 | 0.0000 | NA | NA | NA | NA | NA | NA | 40 | -0.0001 | <.001 | -0.03 | Heaths, inland dunes and semi-natural grasslands | Near Threatened | I |
| *Betula humilis* | 2 | 0.0000 | NA | NA | NA | NA | NA | NA | 9 | 0.0000 | <.001 | -0.01 | None | Highly Endangered | I |
| *Polemonium caeruleum* | 4 | 0.0000 | NA | NA | NA | NA | NA | NA | 16 | 0.0000 | <.001 | -0.01 | None | Endangered | I |
| *Thalictrum simplex* | 1 | 0.0000 | NA | NA | NA | NA | NA | NA | 25 | 0.0000 | <.001 | -0.01 | None | Highly Endangered | I |
| *Phytolacca americana* | 1 | 0.0000 | NA | NA | NA | NA | NA | NA | 74 | 0.0001 | <.001 | 0.06 | None | Not Evaluated | N |
| *Potentilla indica* | 1 | 0.0000 | NA | NA | NA | NA | NA | NA | 31 | 0.0001 | <.001 | 0.03 | None | Not Evaluated | N |
| *Holcus mollis* | 902 | 0.0001 | NA | NA | 288 | -0.0032 | <.001 | -0.40 | 872 | -0.0006 | <.001 | -0.29 | Dry to moderately moist forests | Not Threatened | I |
| *Sisymbrium officinale* | 14 | 0.0001 | NA | NA | 66 | -0.0032 | <.001 | -0.39 | 19 | 0.0000 | NA | NA | Anthropogenic | Not Threatened | A |
| *Lamium album* | 55 | 0.0001 | NA | NA | 146 | -0.0029 | <.001 | -0.35 | 1522 | -0.0004 | <.001 | -0.21 | Scrubs, copses and field hedges | Not Threatened | A |
| *Petasites hybridus* | 78 | 0.0001 | NA | NA | 78 | -0.0005 | NA | NA | 720 | -0.0002 | <.001 | -0.11 | Moist to wet forests | Not Threatened | I |
| *Bromus sterilis* | 22 | 0.0001 | NA | NA | 17 | -0.0005 | NA | NA | 1936 | 0.0019 | <.001 | 0.88 | Scrubs, copses and field hedges | Not Threatened | A |
| *Polygonatum odoratum* | 82 | 0.0001 | NA | NA | 2 | 0.0000 | NA | NA | 542 | -0.0007 | <.001 | -0.31 | Dry to moderately moist forests | Near Threatened | I |
| *Pyrus communis* agg. | 32 | 0.0001 | NA | NA | 74 | 0.0001 | NA | NA | 3988 | 0.0013 | <.001 | 0.61 | Scrubs, copses and field hedges | Not Threatened | NA |
| *Hyacinthoides non-scripta* | 16 | 0.0001 | <.001 | 0.02 | 2 | 0.0002 | NA | NA | 2 | 0.0000 | NA | NA | None | Vulnerable | I |
| *Narcissus pseudonarcissus* | 16 | 0.0001 | <.001 | 0.02 | 2 | 0.0002 | NA | NA | 9 | 0.0000 | NA | NA | None | Endangered | I |
| *Picea omorika* | 18 | 0.0001 | <.001 | 0.03 | 2 | 0.0002 | NA | NA | 18 | 0.0000 | NA | NA | None | Not Evaluated | N |
| *Abies alba* | 62 | 0.0001 | NA | NA | 6 | 0.0002 | NA | NA | 3006 | 0.0016 | <.001 | 0.73 | Dry to moderately moist forests | Not Threatened | I |
| *Castanea sativa* | 13 | 0.0001 | NA | NA | 10 | 0.0005 | NA | NA | 1085 | 0.0008 | <.001 | 0.37 | Scrubs, copses and field hedges | Not Threatened | A |
| *Cornus mas* | 16 | 0.0001 | <.001 | 0.03 | 8 | 0.0006 | <.001 | 0.07 | 286 | 0.0001 | NA | NA | Scrubs, copses and field hedges | Not Threatened | I |
| *Carex pendula* | 34 | 0.0001 | NA | NA | 10 | 0.0008 | <.001 | 0.10 | 1445 | 0.0018 | <.001 | 0.84 | Moist to wet forests | Not Threatened | I |
| *Vinca minor* | 54 | 0.0001 | NA | NA | 17 | 0.0009 | <.001 | 0.11 | 504 | 0.0002 | <.001 | 0.09 | Dry to moderately moist forests | Not Threatened | A |
| *Matteuccia struthiopteris* | 24 | 0.0001 | <.001 | 0.03 | 8 | 0.0011 | <.001 | 0.14 | 6 | 0.0000 | NA | NA | None | Near Threatened | I |
| *Mahonia aquifolium* | 3 | 0.0001 | NA | NA | 20 | 0.0015 | <.001 | 0.19 | 168 | 0.0002 | <.001 | 0.07 | Anthropogenic | Not Evaluated | N |
| *Prunus laurocerasus* | 5 | 0.0001 | NA | NA | 18 | 0.0015 | <.001 | 0.18 | 86 | 0.0002 | <.001 | 0.08 | Anthropogenic | Not Evaluated | N |
| *Prunus domestica* s. l. | 17 | 0.0001 | <.001 | 0.03 | 84 | 0.0015 | NA | NA | 5762 | 0.0042 | <.001 | 1.96 | Scrubs, copses and field hedges | Near Threatened | N |
| *Tilia europaea* | 6 | 0.0001 | NA | NA | 18 | 0.0016 | <.001 | 0.19 | 18 | 0.0000 | NA | NA | Anthropogenic | Not Evaluated | I |
| *Rosa multiflora* | 11 | 0.0001 | <.001 | 0.03 | 36 | 0.0018 | <.001 | 0.23 | 326 | 0.0004 | <.001 | 0.19 | Scrubs, copses and field hedges | Not Evaluated | N |
| *Ligustrum vulgare* | 12 | 0.0001 | NA | NA | 90 | 0.0049 | <.001 | 0.60 | 13798 | 0.0069 | <.001 | 3.21 | Scrubs, copses and field hedges | Not Threatened | I |
| *Poa chaixii* | 22 | 0.0001 | <.001 | 0.02 | NA | NA | NA | NA | 574 | -0.0002 | <.001 | -0.12 | Heaths, inland dunes and semi-natural grasslands | Not Threatened | I |
| *Carex pilosa* | 32 | 0.0001 | <.001 | 0.03 | NA | NA | NA | NA | 152 | 0.0001 | <.001 | 0.06 | Dry to moderately moist forests | Not Threatened | I |
| *Rubus spectabilis* | 20 | 0.0001 | <.001 | 0.02 | NA | NA | NA | NA | NA | NA | NA | NA | None | Not Evaluated | N |
| *Arctium lappa* | 102 | 0.0002 | NA | NA | 61 | -0.0006 | NA | NA | 396 | 0.0002 | <.001 | 0.11 | None | Not Threatened | A |
| *Abies grandis* | 52 | 0.0002 | <.001 | 0.05 | 5 | 0.0001 | NA | NA | 7 | 0.0000 | NA | NA | Dry to moderately moist forests | Not Evaluated | N |
| *Galanthus nivalis* | 31 | 0.0002 | <.001 | 0.04 | 2 | 0.0002 | NA | NA | 24 | 0.0000 | NA | NA | None | Near Threatened | I |
| *Chaerophyllum bulbosum* | 44 | 0.0002 | <.001 | 0.04 | 16 | 0.0002 | NA | NA | 602 | 0.0005 | <.001 | 0.21 | Scrubs, copses and field hedges | Not Threatened | I |
| *Ulmus laevis* | 256 | 0.0002 | NA | NA | 10 | 0.0004 | NA | NA | 317 | 0.0002 | <.001 | 0.10 | Dry to moderately moist forests | Near Threatened | I |
| *Populus balsamifera* | 38 | 0.0002 | <.001 | 0.05 | 16 | 0.0005 | NA | NA | 42 | 0.0000 | NA | NA | None | Not Evaluated | N |
| *Adoxa moschatellina* | 210 | 0.0002 | NA | NA | 2 | 0.0006 | NA | NA | 208 | -0.0002 | <.001 | -0.08 | Dry to moderately moist forests | Not Threatened | I |
| *Syringa vulgaris* | 15 | 0.0002 | NA | NA | 56 | 0.0015 | NA | NA | 586 | 0.0005 | <.001 | 0.21 | Scrubs, copses and field hedges | Not Evaluated | N |
| *Juglans regia* | 21 | 0.0002 | <.001 | 0.03 | 32 | 0.0022 | <.001 | 0.26 | 6497 | 0.0066 | <.001 | 3.04 | Scrubs, copses and field hedges | Not Threatened | A |
| *Fallopia dumetorum* | 64 | 0.0003 | <.001 | 0.07 | 4 | 0.0000 | NA | NA | 48 | 0.0000 | <.001 | 0.02 | None | Not Threatened | I |
| *Ceratocapnos claviculata* | 64 | 0.0003 | <.001 | 0.05 | 4 | 0.0001 | NA | NA | NA | NA | NA | NA | None | Not Threatened | I |
| *Veronica montana* | 248 | 0.0003 | NA | NA | 3 | 0.0004 | NA | NA | 98 | 0.0001 | <.001 | 0.04 | Dry to moderately moist forests | Not Threatened | I |
| *Sorbus intermedia* | 58 | 0.0003 | <.001 | 0.06 | 22 | 0.0006 | NA | NA | 138 | 0.0000 | NA | NA | None | Vulnerable | N |
| *Scutellaria galericulata* | 1689 | 0.0003 | NA | NA | 180 | 0.0007 | NA | NA | 770 | -0.0006 | <.001 | -0.26 | Standing waters | Not Threatened | I |
| *Salix triandra* | 112 | 0.0003 | <.001 | 0.06 | 83 | 0.0008 | NA | NA | 805 | -0.0002 | NA | NA | Moist to wet forests | Not Threatened | I |
| *Malus* | 122 | 0.0003 | NA | NA | 191 | 0.0010 | NA | NA | 5543 | 0.0019 | <.001 | 0.89 | Scrubs, copses and field hedges | Near Threatened | NA |
| *Polygonatum multiflorum* | 2347 | 0.0003 | NA | NA | 112 | 0.0071 | <.001 | 0.87 | 2764 | 0.0008 | NA | NA | Dry to moderately moist forests | Not Threatened | I |
| *Rumex obtusifolius* | 684 | 0.0004 | NA | NA | 514 | -0.0079 | <.001 | -0.97 | 974 | -0.0007 | <.001 | -0.32 | Mesic grasslands | Not Threatened | I |
| *Equisetum arvense* | 586 | 0.0004 | NA | NA | 205 | -0.0066 | <.001 | -0.80 | 1422 | -0.0008 | <.001 | -0.38 | Ruderal, fringe and tall forb communities, clearings | Not Threatened | I |
| *Epilobium ciliatum* | 131 | 0.0004 | <.001 | 0.08 | 99 | -0.0015 | <.001 | -0.19 | 90 | -0.0001 | NA | NA | Linear and running surface waters | Not Evaluated | N |
| *Ulmus minor* agg. | 114 | 0.0004 | NA | NA | 13 | 0.0009 | <.001 | 0.11 | 1140 | 0.0008 | <.001 | 0.38 | Scrubs, copses and field hedges | Not Threatened | I |
| *Heracleum mantegazzianum* | 83 | 0.0004 | <.001 | 0.09 | 54 | 0.0015 | <.001 | 0.18 | 194 | 0.0000 | NA | NA | None | Not Evaluated | N |
| *Spiraea* | 85 | 0.0004 | <.001 | 0.08 | 50 | 0.0020 | <.001 | 0.24 | 72 | 0.0000 | NA | NA | Anthropogenic | Not Evaluated | N |
| *Robinia pseudoacacia* | 70 | 0.0004 | NA | NA | 130 | 0.0041 | <.001 | 0.50 | 3430 | 0.0028 | <.001 | 1.32 | Scrubs, copses and field hedges | Not Evaluated | N |
| *Populus alba* agg. | 242 | 0.0005 | NA | NA | 88 | 0.0007 | NA | NA | 608 | 0.0003 | <.001 | 0.13 | None | Not Threatened | I |
| *Tilia platyphyllos* | 102 | 0.0005 | NA | NA | 36 | 0.0022 | <.001 | 0.27 | 2012 | 0.0010 | <.001 | 0.45 | Dry to moderately moist forests | Not Threatened | I |
| *Cornus sanguinea* | 168 | 0.0005 | NA | NA | 94 | 0.0042 | <.001 | 0.51 | 19774 | 0.0103 | <.001 | 4.79 | Scrubs, copses and field hedges | Not Threatened | I |
| *Salix caprea* | 787 | 0.0006 | NA | NA | 418 | 0.0014 | NA | NA | 10982 | 0.0012 | <.001 | 0.57 | Scrubs, copses and field hedges | Not Threatened | I |
| *Tilia cordata* | 217 | 0.0006 | NA | NA | 88 | 0.0044 | <.001 | 0.54 | 2866 | 0.0017 | <.001 | 0.81 | Scrubs, copses and field hedges | Not Threatened | I |
| *Symphoricarpos albus* | 78 | 0.0006 | <.001 | 0.13 | 122 | 0.0070 | <.001 | 0.85 | 400 | 0.0005 | <.001 | 0.21 | Anthropogenic | Not Evaluated | N |
| *Stellaria alsine* | 364 | 0.0007 | <.001 | 0.16 | 42 | 0.0001 | NA | NA | 942 | -0.0005 | <.001 | -0.25 | Moist to wet grasslands | Not Threatened | I |
| *Cardamine amara* | 1898 | 0.0007 | NA | NA | 38 | 0.0029 | <.001 | 0.36 | 1832 | 0.0002 | NA | NA | Moist to wet forests | Not Threatened | I |
| *Salix alba* | 1119 | 0.0007 | NA | NA | 366 | 0.0061 | <.001 | 0.75 | 3787 | 0.0006 | NA | NA | Moist to wet forests | Not Threatened | I |
| *Galium odoratum* | 1836 | 0.0008 | NA | NA | 24 | 0.0028 | <.001 | 0.34 | 6090 | 0.0032 | <.001 | 1.48 | Dry to moderately moist forests | Not Threatened | I |
| *Fallopia bohemica_Fallopia japonica_Fallopia sachalinensis* | 138 | 0.0008 | <.001 | 0.16 | 208 | 0.0056 | <.001 | 0.69 | 439 | 0.0005 | <.001 | 0.25 | Anthropogenic | Not Evaluated | N |
| *Taxus baccata* | 116 | 0.0008 | <.001 | 0.17 | 100 | 0.0091 | <.001 | 1.11 | 360 | 0.0004 | <.001 | 0.18 | Anthropogenic | Near Threatened | I |
| *Circaea intermedia* | 199 | 0.0009 | <.001 | 0.18 | 1 | 0.0001 | NA | NA | 76 | 0.0001 | <.001 | 0.03 | Dry to moderately moist forests | Not Threatened | I |
| *Chaerophyllum temulum* | 162 | 0.0009 | NA | NA | 45 | 0.0021 | <.001 | 0.26 | 352 | 0.0001 | NA | NA | Scrubs, copses and field hedges | Not Threatened | I |
| *Pseudotsuga menziesii* | 186 | 0.0009 | NA | NA | 68 | 0.0023 | <.001 | 0.28 | 861 | 0.0005 | <.001 | 0.25 | Dry to moderately moist forests | Not Evaluated | N |
| *Chelidonium majus* | 104 | 0.0009 | <.001 | 0.19 | 41 | 0.0033 | <.001 | 0.40 | 1488 | 0.0007 | <.001 | 0.32 | Scrubs, copses and field hedges | Not Threatened | I |
| *Aesculus hippocastanum* | 286 | 0.0010 | NA | NA | 135 | 0.0086 | <.001 | 1.05 | 1168 | 0.0008 | <.001 | 0.38 | Anthropogenic | Not Evaluated | N |
| *Carex strigosa* | 324 | 0.0010 | <.001 | 0.21 | NA | NA | NA | NA | 84 | 0.0001 | <.001 | 0.05 | Dry to moderately moist forests | Not Threatened | I |
| *Cardamine flexuosa* | 370 | 0.0011 | <.001 | 0.23 | 18 | 0.0008 | <.001 | 0.10 | 282 | 0.0003 | <.001 | 0.14 | Dry to moderately moist forests | Not Threatened | I |
| *Veronica hederifolia* agg. | 196 | 0.0012 | <.001 | 0.25 | 14 | 0.0009 | <.001 | 0.11 | 209 | -0.0001 | <.001 | -0.07 | Dry to moderately moist forests | Not Threatened | I |
| *Brachypodium sylvaticum* | 1228 | 0.0012 | NA | NA | 13 | 0.0012 | <.001 | 0.15 | 5442 | 0.0056 | <.001 | 2.58 | Dry to moderately moist forests | Not Threatened | I |
| *Larix kaempferi* | 166 | 0.0013 | <.001 | 0.28 | 49 | 0.0012 | NA | NA | 26 | 0.0000 | NA | NA | Dry to moderately moist forests | Not Evaluated | N |
| *Populus nigra_Populus canadensis* | 625 | 0.0013 | <.001 | 0.28 | 175 | 0.0027 | NA | NA | 1915 | 0.0009 | <.001 | 0.44 | Moist to wet forests | Not Evaluated | NA |
| *Quercus rubra* | 208 | 0.0013 | <.001 | 0.27 | 105 | 0.0048 | <.001 | 0.59 | 461 | 0.0004 | <.001 | 0.18 | Dry to moderately moist forests | Not Evaluated | N |
| *Picea sitchensis* | 296 | 0.0014 | <.001 | 0.29 | 10 | 0.0001 | NA | NA | 6 | 0.0000 | NA | NA | Dry to moderately moist forests | Not Evaluated | N |
| *Chrysosplenium oppositifolium* | 774 | 0.0014 | <.001 | 0.30 | 7 | 0.0012 | <.001 | 0.14 | 1250 | 0.0009 | <.001 | 0.40 | Moist to wet forests | Not Threatened | I |
| *Equisetum sylvaticum* | 664 | 0.0014 | NA | NA | 28 | 0.0015 | <.001 | 0.18 | 939 | -0.0009 | <.001 | -0.40 | Moist to wet forests | Not Threatened | I |
| *Quercus petraea* agg. | 318 | 0.0015 | <.001 | 0.33 | 148 | 0.0011 | NA | NA | 4276 | 0.0011 | <.001 | 0.51 | Dry to moderately moist forests | Not Threatened | I |
| *Carex sylvatica* | 1440 | 0.0015 | NA | NA | 20 | 0.0023 | <.001 | 0.28 | 3730 | 0.0035 | <.001 | 1.61 | Dry to moderately moist forests | Not Threatened | I |
| *Rumex sanguineus* | 550 | 0.0016 | <.001 | 0.33 | 54 | 0.0039 | <.001 | 0.47 | 162 | 0.0000 | NA | NA | Moist to wet forests | Not Threatened | I |
| *Rubus* sect. *Caesii* | 724 | 0.0016 | <.001 | 0.35 | 101 | 0.0062 | <.001 | 0.76 | 9583 | 0.0073 | <.001 | 3.38 | Scrubs, copses and field hedges | Not Threatened | I |
| *Acer platanoides* | 333 | 0.0016 | NA | NA | 233 | 0.0161 | <.001 | 1.96 | 4930 | 0.0041 | <.001 | 1.88 | Scrubs, copses and field hedges | Not Threatened | I |
| *Acer campestre* | 572 | 0.0017 | NA | NA | 190 | 0.0118 | <.001 | 1.44 | 15178 | 0.0091 | <.001 | 4.24 | Scrubs, copses and field hedges | Not Threatened | I |
| *Humulus lupulus* | 1240 | 0.0018 | <.001 | 0.39 | 193 | 0.0070 | <.001 | 0.86 | 3668 | 0.0024 | <.001 | 1.11 | Moist to wet forests | Not Threatened | I |
| *Galeopsis tetrahit* agg. | 1298 | 0.0019 | <.001 | 0.40 | 380 | 0.0009 | NA | NA | 5552 | 0.0006 | NA | NA | Scrubs, copses and field hedges | Not Threatened | I |
| *Prunus spinosa* agg. | 884 | 0.0019 | NA | NA | 128 | 0.0038 | <.001 | 0.46 | 21484 | 0.0059 | <.001 | 2.73 | Scrubs, copses and field hedges | Not Threatened | I |
| *Iris pseudacorus* | 3960 | 0.0019 | NA | NA | 418 | 0.0054 | NA | NA | 2523 | -0.0013 | <.001 | -0.58 | Standing waters | Not Threatened | I |
| *Anemone nemorosa* | 2328 | 0.0020 | NA | NA | 50 | 0.0037 | <.001 | 0.46 | 3798 | -0.0012 | <.001 | -0.58 | Dry to moderately moist forests | Not Threatened | I |
| *Impatiens glandulifera* | 388 | 0.0020 | <.001 | 0.42 | 99 | 0.0044 | <.001 | 0.54 | 3699 | 0.0029 | <.001 | 1.35 | Moist to wet forests | Not Evaluated | N |
| *Euonymus europaeus* | 978 | 0.0020 | NA | NA | 70 | 0.0061 | <.001 | 0.75 | 15076 | 0.0081 | <.001 | 3.74 | Scrubs, copses and field hedges | Not Threatened | I |
| *Populus tremula* | 1760 | 0.0022 | NA | NA | 359 | 0.0046 | <.001 | 0.56 | 5704 | 0.0005 | NA | NA | Scrubs, copses and field hedges | Not Threatened | I |
| *Elymus repens* s. str. | 727 | 0.0025 | <.001 | 0.53 | 594 | -0.0134 | <.001 | -1.63 | 3962 | 0.0018 | <.001 | 0.85 | Scrubs, copses and field hedges | Not Threatened | I |
| *Melica uniflora* | 2075 | 0.0025 | NA | NA | 12 | 0.0014 | <.001 | 0.18 | 664 | 0.0004 | <.001 | 0.17 | Dry to moderately moist forests | Not Threatened | I |
| *Carex elongata* | 1148 | 0.0026 | NA | NA | 42 | 0.0014 | <.001 | 0.18 | 326 | -0.0001 | NA | NA | Moist to wet forests | Not Threatened | I |
| *Ribes uva-crispa* | 688 | 0.0026 | NA | NA | 48 | 0.0031 | <.001 | 0.38 | 4266 | 0.0007 | NA | NA | Scrubs, copses and field hedges | Not Threatened | I |
| *Salix viminalis* | 780 | 0.0027 | <.001 | 0.57 | 302 | 0.0007 | NA | NA | 2312 | -0.0003 | NA | NA | Moist to wet forests | Not Threatened | I |
| *Larix decidua* | 680 | 0.0028 | <.001 | 0.58 | 60 | 0.0021 | <.001 | 0.26 | 730 | 0.0001 | NA | NA | Dry to moderately moist forests | Not Threatened | I |
| *Prunus avium* | 1236 | 0.0029 | NA | NA | 132 | 0.0069 | <.001 | 0.84 | 16675 | 0.0088 | <.001 | 4.07 | Scrubs, copses and field hedges | Not Threatened | I |
| *Festuca altissima* | 1049 | 0.0031 | NA | NA | 20 | 0.0005 | <.001 | 0.06 | 917 | 0.0007 | <.001 | 0.34 | Dry to moderately moist forests | Not Threatened | I |
| *Ribes nigrum* | 920 | 0.0032 | <.001 | 0.68 | 29 | 0.0020 | <.001 | 0.24 | 186 | 0.0000 | NA | NA | Moist to wet forests | Not Threatened | I |
| *Scirpus sylvaticus* | 2541 | 0.0035 | NA | NA | 222 | 0.0033 | NA | NA | 6744 | -0.0070 | <.001 | -3.27 | Moist to wet grasslands | Not Threatened | I |
| *Ilex aquifolium* | 1314 | 0.0035 | NA | NA | 138 | 0.0098 | <.001 | 1.20 | 1040 | 0.0008 | <.001 | 0.39 | Dry to moderately moist forests | Not Threatened | I |
| *Calystegia sepium* agg. | 1420 | 0.0036 | <.001 | 0.77 | 465 | 0.0044 | <.001 | 0.54 | 4864 | 0.0007 | NA | NA | Fresh water vegetation, springs and reeds | Not Threatened | NA |
| *Oxalis acetosella* | 2212 | 0.0039 | NA | NA | 144 | 0.0075 | <.001 | 0.92 | 2762 | 0.0012 | <.001 | 0.55 | Dry to moderately moist forests | Not Threatened | I |
| *Solanum dulcamara* | 3330 | 0.0040 | NA | NA | 298 | 0.0037 | <.001 | 0.45 | 1812 | -0.0004 | NA | NA | Standing waters | Not Threatened | I |
| *Ribes rubrum* agg. | 1397 | 0.0041 | <.001 | 0.87 | 98 | 0.0077 | <.001 | 0.95 | 1218 | 0.0008 | <.001 | 0.35 | Moist to wet forests | Not Threatened | I |
| *Prunus padus* | 1417 | 0.0042 | <.001 | 0.89 | 208 | 0.0170 | <.001 | 2.08 | 4686 | 0.0023 | <.001 | 1.06 | Moist to wet forests | Not Threatened | I |
| *Glyceria fluitans* agg. | 4003 | 0.0043 | NA | NA | 378 | -0.0034 | NA | NA | 3152 | -0.0023 | <.001 | -1.07 | Standing waters | Not Threatened | I |
| *Betula pendula* | 2650 | 0.0046 | NA | NA | 777 | 0.0108 | <.001 | 1.31 | 7156 | -0.0002 | NA | NA | Dry to moderately moist forests | Not Threatened | I |
| *Carex acutiformis* | 3831 | 0.0048 | NA | NA | 170 | 0.0045 | <.001 | 0.55 | 6115 | -0.0035 | <.001 | -1.63 | Fresh water vegetation, springs and reeds | Not Threatened | I |
| *Aegopodium podagraria* | 1926 | 0.0048 | NA | NA | 470 | 0.0154 | <.001 | 1.88 | 10437 | 0.0048 | <.001 | 2.20 | Scrubs, copses and field hedges | Not Threatened | I |
| *Deschampsia cespitosa* agg. | 4332 | 0.0049 | <.001 | 1.05 | 471 | 0.0036 | NA | NA | 6425 | -0.0026 | <.001 | -1.19 | Moist to wet forests | Not Threatened | I |
| *Sambucus nigra* | 3310 | 0.0050 | NA | NA | 740 | 0.0224 | <.001 | 2.74 | 22877 | 0.0105 | <.001 | 4.88 | Scrubs, copses and field hedges | Not Threatened | I |
| *Prunus serotina* | 1558 | 0.0051 | <.001 | 1.09 | 255 | 0.0108 | <.001 | 1.32 | 360 | 0.0004 | <.001 | 0.20 | Dry to moderately moist forests | Not Evaluated | N |
| *Athyrium filix-femina* | 1942 | 0.0052 | <.001 | 1.10 | 189 | 0.0093 | <.001 | 1.13 | 4008 | 0.0023 | <.001 | 1.08 | Moist to wet forests | Not Threatened | I |
| *Impatiens noli-tangere* | 1702 | 0.0053 | <.001 | 1.13 | 54 | 0.0032 | <.001 | 0.39 | 3682 | 0.0016 | <.001 | 0.72 | Moist to wet forests | Not Threatened | I |
| *Dactylis glomerata* agg. | 2290 | 0.0054 | NA | NA | 836 | -0.0042 | NA | NA | 13061 | 0.0031 | <.001 | 1.44 | Scrubs, copses and field hedges | Not Threatened | I |
| *Salix fragilis* agg. | 1150 | 0.0054 | <.001 | 1.14 | 208 | 0.0038 | <.001 | 0.47 | 5566 | 0.0018 | <.001 | 0.83 | Moist to wet forests | Not Threatened | I |
| *Milium effusum* | 2731 | 0.0054 | NA | NA | 104 | 0.0086 | <.001 | 1.06 | 4020 | 0.0020 | <.001 | 0.92 | Dry to moderately moist forests | Not Threatened | I |
| *Ficaria verna* s. l. | 2319 | 0.0055 | <.001 | 1.16 | 72 | 0.0046 | <.001 | 0.56 | 2699 | -0.0009 | <.001 | -0.44 | Dry to moderately moist forests | Not Threatened | I |
| *Poa nemoralis* agg. | 1591 | 0.0055 | NA | NA | 178 | 0.0097 | <.001 | 1.19 | 4182 | 0.0017 | <.001 | 0.81 | Dry to moderately moist forests | Not Threatened | I |
| *Alliaria petiolata* | 1180 | 0.0055 | <.001 | 1.15 | 250 | 0.0167 | <.001 | 2.05 | 9350 | 0.0079 | <.001 | 3.66 | Scrubs, copses and field hedges | Not Threatened | I |
| *Sorbus aucuparia* | 3506 | 0.0058 | <.001 | 1.23 | 533 | 0.0171 | <.001 | 2.09 | 6278 | -0.0017 | <.001 | -0.79 | Dry to moderately moist forests | Not Threatened | I |
| *Carpinus betulus* | 2266 | 0.0058 | NA | NA | 292 | 0.0172 | <.001 | 2.10 | 10810 | 0.0075 | <.001 | 3.47 | Dry to moderately moist forests | Not Threatened | I |
| *Dryopteris filix-mas* agg. | 1965 | 0.0059 | NA | NA | 344 | 0.0191 | <.001 | 2.33 | 7152 | 0.0056 | <.001 | 2.62 | Dry to moderately moist forests | Not Threatened | I |
| *Crataegus* | 3389 | 0.0060 | NA | NA | 478 | 0.0173 | <.001 | 2.11 | 18919 | 0.0065 | <.001 | 3.01 | Scrubs, copses and field hedges | Not Threatened | I |
| *Circaea lutetiana* | 2698 | 0.0063 | <.001 | 1.34 | 120 | 0.0108 | <.001 | 1.32 | 3252 | 0.0032 | <.001 | 1.47 | Dry to moderately moist forests | Not Threatened | I |
| *Poa trivialis* s. l. | 3006 | 0.0065 | <.001 | 1.38 | 578 | -0.0005 | NA | NA | 1723 | -0.0011 | <.001 | -0.53 | Moist to wet grasslands | Not Threatened | I |
| *Lonicera periclymenum* | 3324 | 0.0070 | NA | NA | 212 | 0.0104 | <.001 | 1.27 | 576 | 0.0003 | <.001 | 0.16 | Dry to moderately moist forests | Not Threatened | I |
| *Picea abies* | 1958 | 0.0073 | <.001 | 1.55 | 244 | 0.0071 | <.001 | 0.87 | 11900 | -0.0013 | NA | NA | Dry to moderately moist forests | Not Threatened | I |
| *Phalaris arundinacea* | 4222 | 0.0076 | NA | NA | 562 | -0.0031 | NA | NA | 4730 | -0.0021 | <.001 | -0.97 | Fresh water vegetation, springs and reeds | Not Threatened | I |
| *Geranium robertianum* agg. | 2120 | 0.0076 | <.001 | 1.60 | 90 | 0.0065 | <.001 | 0.80 | 11282 | 0.0063 | <.001 | 2.92 | Scrubs, copses and field hedges | Not Threatened | NA |
| *Corylus avellana* | 3294 | 0.0077 | NA | NA | 328 | 0.0221 | <.001 | 2.70 | 21638 | 0.0123 | <.001 | 5.69 | Scrubs, copses and field hedges | Not Threatened | I |
| *Hedera helix* | 2232 | 0.0080 | NA | NA | 219 | 0.0180 | <.001 | 2.20 | 9730 | 0.0102 | <.001 | 4.74 | Dry to moderately moist forests | Not Threatened | I |
| *Geum urbanum* | 2633 | 0.0083 | <.001 | 1.75 | 348 | 0.0263 | <.001 | 3.22 | 15057 | 0.0104 | <.001 | 4.82 | Scrubs, copses and field hedges | Not Threatened | I |
| *Quercus robur* | 4932 | 0.0092 | NA | NA | 710 | 0.0212 | <.001 | 2.60 | 16573 | 0.0074 | <.001 | 3.42 | Scrubs, copses and field hedges | Not Threatened | I |
| *Galium aparine* agg. | 2587 | 0.0093 | <.001 | 1.97 | 466 | 0.0071 | <.001 | 0.87 | 12650 | 0.0030 | <.001 | 1.38 | Scrubs, copses and field hedges | Not Threatened | I |
| *Dryopteris carthusiana* agg. | 4632 | 0.0099 | <.001 | 2.09 | 382 | 0.0151 | <.001 | 1.84 | 3582 | 0.0017 | <.001 | 0.81 | Dry to moderately moist forests | Not Threatened | I |
| *Fraxinus excelsior* | 3544 | 0.0100 | <.001 | 2.12 | 444 | 0.0216 | <.001 | 2.64 | 22786 | 0.0115 | <.001 | 5.34 | Scrubs, copses and field hedges | Not Threatened | I |
| *Stellaria holostea* | 3746 | 0.0103 | NA | NA | 120 | 0.0063 | <.001 | 0.76 | 2116 | 0.0007 | <.001 | 0.32 | Dry to moderately moist forests | Not Threatened | I |
| *Impatiens parviflora* | 1948 | 0.0109 | <.001 | 2.31 | 483 | 0.0241 | <.001 | 2.94 | 1208 | 0.0008 | <.001 | 0.37 | Dry to moderately moist forests | Not Evaluated | N |
| *Rubus idaeus* | 4244 | 0.0112 | <.001 | 2.38 | 518 | 0.0147 | <.001 | 1.79 | 9344 | -0.0003 | NA | NA | Scrubs, copses and field hedges | Not Threatened | I |
| *Acer pseudoplatanus* | 2928 | 0.0112 | NA | NA | 472 | 0.0268 | <.001 | 3.28 | 16524 | 0.0098 | <.001 | 4.54 | Dry to moderately moist forests | Not Threatened | I |
| *Alnus glutinosa* | 5080 | 0.0121 | NA | NA | 583 | 0.0176 | <.001 | 2.15 | 9470 | 0.0027 | <.001 | 1.24 | Moist to wet forests | Not Threatened | I |
| *Fagus sylvatica* | 2976 | 0.0123 | NA | NA | 356 | 0.0192 | <.001 | 2.35 | 11540 | 0.0035 | <.001 | 1.61 | Dry to moderately moist forests | Not Threatened | I |
| *Galeobdolon luteum* agg. | 3294 | 0.0130 | <.001 | 2.75 | 220 | 0.0184 | <.001 | 2.25 | 5666 | 0.0041 | <.001 | 1.92 | Dry to moderately moist forests | Not Threatened | NA |
| *Carex remota* | 2979 | 0.0145 | <.001 | 3.08 | 104 | 0.0080 | <.001 | 0.98 | 1942 | 0.0020 | <.001 | 0.94 | Standing waters | Not Threatened | I |
| *Glechoma hederacea* agg. | 4288 | 0.0150 | <.001 | 3.17 | 782 | 0.0188 | <.001 | 2.30 | 7063 | 0.0038 | <.001 | 1.78 | Moist to wet forests | Not Threatened | I |
| *Rubus* sect. *Rubus* | 5653 | 0.0209 | <.001 | 4.43 | 796 | 0.0184 | <.001 | 2.24 | 17402 | 0.0105 | <.001 | 4.86 | Scrubs, copses and field hedges | Not Threatened | NA |
| *Urtica dioica* s. l. | 7535 | 0.0279 | <.001 | 5.91 | 1270 | 0.0140 | <.001 | 1.72 | 28635 | 0.0093 | <.001 | 4.29 | Scrubs, copses and field hedges | Not Threatened | I |
| *Sedum album* | NA | NA | NA | NA | 1 | -0.0002 | NA | NA | 871 | -0.0009 | <.001 | -0.40 | Heaths, inland dunes and semi-natural grasslands | Not Threatened | I |
| *Anthemis tinctoria* agg. | NA | NA | NA | NA | 4 | -0.0002 | NA | NA | 110 | -0.0001 | <.001 | -0.06 | Heaths, inland dunes and semi-natural grasslands | Not Threatened | A |
| *Crepis vesicaria* | NA | NA | NA | NA | 1 | -0.0002 | NA | NA | 10 | 0.0000 | <.001 | -0.01 | None | Not Threatened | A |
| *Festuca pallens* | NA | NA | NA | NA | 1 | -0.0001 | NA | NA | 75 | -0.0001 | <.001 | -0.04 | Dry to moderately moist forests | Near Threatened | I |
| *Rosa gallica* | NA | NA | NA | NA | 3 | -0.0001 | NA | NA | 86 | -0.0001 | <.001 | -0.03 | None | Endangered | I |
| *Buddleja davidii* | NA | NA | NA | NA | 2 | -0.0001 | NA | NA | 36 | 0.0001 | <.001 | 0.03 | None | Not Evaluated | N |
| *Asplenium ruta-muraria* | NA | NA | NA | NA | 1 | 0.0000 | NA | NA | 619 | -0.0006 | <.001 | -0.30 | Heaths, inland dunes and semi-natural grasslands | Not Threatened | I |
| *Microthlaspi perfoliatum* | NA | NA | NA | NA | 1 | 0.0000 | NA | NA | 482 | -0.0006 | <.001 | -0.26 | Heaths, inland dunes and semi-natural grasslands | Not Threatened | I |
| *Verbascum lychnitis* | NA | NA | NA | NA | 1 | 0.0000 | NA | NA | 655 | -0.0006 | <.001 | -0.28 | Heaths, inland dunes and semi-natural grasslands | Not Threatened | I |
| *Hieracium piloselloides* | NA | NA | NA | NA | 1 | 0.0000 | NA | NA | 197 | -0.0003 | <.001 | -0.13 | Heaths, inland dunes and semi-natural grasslands | Not Threatened | I |
| *Muscari botryoides* | NA | NA | NA | NA | 1 | 0.0000 | NA | NA | 244 | -0.0003 | <.001 | -0.14 | Heaths, inland dunes and semi-natural grasslands | Endangered | I |
| *Cerastium brachypetalum* agg. | NA | NA | NA | NA | 1 | 0.0000 | NA | NA | 48 | -0.0001 | <.001 | -0.03 | Heaths, inland dunes and semi-natural grasslands | Not Threatened | A |
| *Medicago minima* | NA | NA | NA | NA | 1 | 0.0000 | NA | NA | 38 | -0.0001 | <.001 | -0.03 | Heaths, inland dunes and semi-natural grasslands | Near Threatened | I |
| *Asplenium scolopendrium* | NA | NA | NA | NA | 1 | 0.0000 | NA | NA | 94 | 0.0001 | <.001 | 0.04 | Dry to moderately moist forests | Not Threatened | I |
| *Ailanthus altissima* | NA | NA | NA | NA | 2 | 0.0001 | NA | NA | 47 | 0.0001 | <.001 | 0.03 | None | Not Evaluated | N |
| *Buxus sempervirens* | NA | NA | NA | NA | 3 | 0.0002 | NA | NA | 58 | 0.0001 | <.001 | 0.03 | None | Endangered | I |
| *Sorbaria sorbifolia* | NA | NA | NA | NA | 15 | 0.0006 | <.001 | 0.07 | 1 | 0.0000 | NA | NA | None | Not Evaluated | N |
| *Salix sepulcralis* | NA | NA | NA | NA | 78 | 0.0014 | <.001 | 0.17 | 10 | 0.0000 | NA | NA | Standing waters | Not Evaluated | N |
| *Amelanchier lamarckii* | NA | NA | NA | NA | 20 | 0.0015 | <.001 | 0.18 | 8 | 0.0000 | NA | NA | None | Not Evaluated | N |
| *Cornus sericea_Cornus alba* | NA | NA | NA | NA | 90 | 0.0038 | <.001 | 0.46 | 238 | 0.0002 | <.001 | 0.10 | Anthropogenic | Not Evaluated | N |
| *Philadelphus coronarius* | NA | NA | NA | NA | 70 | 0.0048 | <.001 | 0.59 | 56 | 0.0001 | <.001 | 0.03 | Anthropogenic | Not Evaluated | N |
| *Carlina acaulis* | NA | NA | NA | NA | NA | NA | NA | NA | 2255 | -0.0036 | <.001 | -1.68 | Heaths, inland dunes and semi-natural grasslands | Near Threatened | I |
| *Hippocrepis comosa* | NA | NA | NA | NA | NA | NA | NA | NA | 2273 | -0.0035 | <.001 | -1.63 | Heaths, inland dunes and semi-natural grasslands | Near Threatened | I |
| *Koeleria pyramidata* agg. | NA | NA | NA | NA | NA | NA | NA | NA | 2228 | -0.0034 | <.001 | -1.57 | Heaths, inland dunes and semi-natural grasslands | Near Threatened | I |
| *Stachys recta* | NA | NA | NA | NA | NA | NA | NA | NA | 2856 | -0.0031 | <.001 | -1.46 | Heaths, inland dunes and semi-natural grasslands | Near Threatened | I |
| *Asperula cynanchica* | NA | NA | NA | NA | NA | NA | NA | NA | 1772 | -0.0027 | <.001 | -1.27 | Heaths, inland dunes and semi-natural grasslands | Near Threatened | I |
| *Genista sagittalis* | NA | NA | NA | NA | NA | NA | NA | NA | 1634 | -0.0026 | <.001 | -1.20 | Heaths, inland dunes and semi-natural grasslands | Near Threatened | I |
| *Prunella grandiflora* | NA | NA | NA | NA | NA | NA | NA | NA | 1794 | -0.0026 | <.001 | -1.18 | Heaths, inland dunes and semi-natural grasslands | Near Threatened | I |
| *Teucrium chamaedrys* | NA | NA | NA | NA | NA | NA | NA | NA | 1616 | -0.0023 | <.001 | -1.05 | Heaths, inland dunes and semi-natural grasslands | Not Threatened | I |
| *Gymnadenia conopsea* s. l. | NA | NA | NA | NA | NA | NA | NA | NA | 1036 | -0.0020 | <.001 | -0.91 | Heaths, inland dunes and semi-natural grasslands | Near Threatened | I |
| *Colchicum autumnale* | NA | NA | NA | NA | NA | NA | NA | NA | 2224 | -0.0019 | <.001 | -0.89 | Moist to wet grasslands | Not Threatened | I |
| *Ranunculus aconitifolius* | NA | NA | NA | NA | NA | NA | NA | NA | 1364 | -0.0019 | <.001 | -0.90 | Moist to wet grasslands | Not Threatened | I |
| *Meum athamanticum* | NA | NA | NA | NA | NA | NA | NA | NA | 966 | -0.0017 | <.001 | -0.79 | Heaths, inland dunes and semi-natural grasslands | Near Threatened | I |
| *Veronica austriaca* agg. | NA | NA | NA | NA | NA | NA | NA | NA | 1640 | -0.0017 | <.001 | -0.81 | Heaths, inland dunes and semi-natural grasslands | Not Threatened | I |
| *Bupleurum falcatum* | NA | NA | NA | NA | NA | NA | NA | NA | 1513 | -0.0015 | <.001 | -0.70 | Heaths, inland dunes and semi-natural grasslands | Near Threatened | I |
| *Polygala comosa* | NA | NA | NA | NA | NA | NA | NA | NA | 976 | -0.0015 | <.001 | -0.71 | Heaths, inland dunes and semi-natural grasslands | Near Threatened | I |
| *Trollius europaeus* | NA | NA | NA | NA | NA | NA | NA | NA | 695 | -0.0014 | <.001 | -0.66 | Moist to wet grasslands | Endangered | I |
| *Cirsium rivulare* | NA | NA | NA | NA | NA | NA | NA | NA | 704 | -0.0013 | <.001 | -0.60 | Moist to wet grasslands | Endangered | I |
| *Euphorbia verrucosa* | NA | NA | NA | NA | NA | NA | NA | NA | 1164 | -0.0013 | <.001 | -0.63 | Heaths, inland dunes and semi-natural grasslands | Near Threatened | I |
| *Aster amellus* | NA | NA | NA | NA | NA | NA | NA | NA | 706 | -0.0012 | <.001 | -0.54 | Heaths, inland dunes and semi-natural grasslands | Endangered | I |
| *Chaerophyllum hirsutum* agg. | NA | NA | NA | NA | NA | NA | NA | NA | 2356 | -0.0012 | <.001 | -0.55 | Moist to wet grasslands | Not Threatened | I |
| *Gentianopsis ciliata* | NA | NA | NA | NA | NA | NA | NA | NA | 776 | -0.0012 | <.001 | -0.57 | Heaths, inland dunes and semi-natural grasslands | Near Threatened | I |
| *Buphthalmum salicifolium* | NA | NA | NA | NA | NA | NA | NA | NA | 584 | -0.0011 | <.001 | -0.51 | Heaths, inland dunes and semi-natural grasslands | Not Threatened | I |
| *Orchis militaris* | NA | NA | NA | NA | NA | NA | NA | NA | 678 | -0.0011 | <.001 | -0.50 | Heaths, inland dunes and semi-natural grasslands | Endangered | I |
| *Cirsium eriophorum* | NA | NA | NA | NA | NA | NA | NA | NA | 842 | -0.0010 | <.001 | -0.46 | Heaths, inland dunes and semi-natural grasslands | Not Threatened | I |
| *Teucrium montanum* | NA | NA | NA | NA | NA | NA | NA | NA | 458 | -0.0010 | <.001 | -0.45 | Heaths, inland dunes and semi-natural grasslands | Near Threatened | I |
| *Carex davalliana* | NA | NA | NA | NA | NA | NA | NA | NA | 358 | -0.0009 | <.001 | -0.41 | Moist to wet grasslands | Endangered | I |
| *Gentianella germanica* agg. | NA | NA | NA | NA | NA | NA | NA | NA | 408 | -0.0008 | <.001 | -0.36 | Heaths, inland dunes and semi-natural grasslands | Near Threatened | I |
| *Inula conyzae* | NA | NA | NA | NA | NA | NA | NA | NA | 1338 | -0.0008 | <.001 | -0.37 | Heaths, inland dunes and semi-natural grasslands | Not Threatened | I |
| *Peucedanum cervaria* | NA | NA | NA | NA | NA | NA | NA | NA | 498 | -0.0008 | <.001 | -0.38 | Heaths, inland dunes and semi-natural grasslands | Near Threatened | I |
| *Polygala amara* agg. | NA | NA | NA | NA | NA | NA | NA | NA | 424 | -0.0008 | <.001 | -0.37 | Heaths, inland dunes and semi-natural grasslands | Endangered | I |
| *Thesium bavarum* | NA | NA | NA | NA | NA | NA | NA | NA | 448 | -0.0008 | <.001 | -0.39 | Heaths, inland dunes and semi-natural grasslands | Endangered | I |
| *Tanacetum corymbosum* s. l. | NA | NA | NA | NA | NA | NA | NA | NA | 586 | -0.0007 | <.001 | -0.34 | Dry to moderately moist forests | Near Threatened | I |
| *Globularia bisnagarica* | NA | NA | NA | NA | NA | NA | NA | NA | 258 | -0.0006 | <.001 | -0.28 | Heaths, inland dunes and semi-natural grasslands | Endangered | I |
| *Rhinanthus aristatus* agg. | NA | NA | NA | NA | NA | NA | NA | NA | 420 | -0.0006 | <.001 | -0.26 | Heaths, inland dunes and semi-natural grasslands | Near Threatened | I |
| *Sesleria varia* agg. | NA | NA | NA | NA | NA | NA | NA | NA | 412 | -0.0006 | <.001 | -0.30 | Dry to moderately moist forests | Not Threatened | I |
| *Carex humilis* | NA | NA | NA | NA | NA | NA | NA | NA | 238 | -0.0005 | <.001 | -0.22 | Heaths, inland dunes and semi-natural grasslands | Near Threatened | I |
| *Gentiana verna* | NA | NA | NA | NA | NA | NA | NA | NA | 254 | -0.0005 | <.001 | -0.24 | Heaths, inland dunes and semi-natural grasslands | Endangered | I |
| *Laserpitium latifolium* | NA | NA | NA | NA | NA | NA | NA | NA | 430 | -0.0005 | <.001 | -0.24 | Dry to moderately moist forests | Near Threatened | I |
| *Ophrys insectifera* | NA | NA | NA | NA | NA | NA | NA | NA | 250 | -0.0005 | <.001 | -0.25 | Heaths, inland dunes and semi-natural grasslands | Endangered | I |
| *Epipactis atrorubens* | NA | NA | NA | NA | NA | NA | NA | NA | 214 | -0.0004 | <.001 | -0.21 | Dry to moderately moist forests | Near Threatened | I |
| *Gentiana lutea* | NA | NA | NA | NA | NA | NA | NA | NA | 199 | -0.0004 | <.001 | -0.18 | Heaths, inland dunes and semi-natural grasslands | Endangered | I |
| *Orobanche caryophyllacea* | NA | NA | NA | NA | NA | NA | NA | NA | 276 | -0.0004 | <.001 | -0.19 | Heaths, inland dunes and semi-natural grasslands | Endangered | I |
| *Teucrium botrys* | NA | NA | NA | NA | NA | NA | NA | NA | 206 | -0.0004 | <.001 | -0.17 | Heaths, inland dunes and semi-natural grasslands | Near Threatened | A |
| *Thesium pyrenaicum* | NA | NA | NA | NA | NA | NA | NA | NA | 184 | -0.0004 | <.001 | -0.17 | Heaths, inland dunes and semi-natural grasslands | Endangered | I |
| *Carex ornithopoda* s. str. | NA | NA | NA | NA | NA | NA | NA | NA | 265 | -0.0003 | <.001 | -0.15 | Heaths, inland dunes and semi-natural grasslands | Not Evaluated | I |
| *Cirsium tuberosum* | NA | NA | NA | NA | NA | NA | NA | NA | 161 | -0.0003 | <.001 | -0.14 | Heaths, inland dunes and semi-natural grasslands | Endangered | I |
| *Cytisus nigricans* | NA | NA | NA | NA | NA | NA | NA | NA | 147 | -0.0003 | <.001 | -0.15 | Heaths, inland dunes and semi-natural grasslands | Endangered | I |
| *Galium boreale* | NA | NA | NA | NA | NA | NA | NA | NA | 158 | -0.0003 | <.001 | -0.15 | Moist to wet grasslands | Near Threatened | I |
| *Galium glaucum* | NA | NA | NA | NA | NA | NA | NA | NA | 216 | -0.0003 | <.001 | -0.14 | Heaths, inland dunes and semi-natural grasslands | Near Threatened | I |
| *Lactuca perennis* | NA | NA | NA | NA | NA | NA | NA | NA | 156 | -0.0003 | <.001 | -0.14 | Heaths, inland dunes and semi-natural grasslands | Endangered | I |
| *Linum tenuifolium* | NA | NA | NA | NA | NA | NA | NA | NA | 130 | -0.0003 | <.001 | -0.12 | Heaths, inland dunes and semi-natural grasslands | Endangered | I |
| *Lotus maritimus* | NA | NA | NA | NA | NA | NA | NA | NA | 124 | -0.0003 | <.001 | -0.12 | Heaths, inland dunes and semi-natural grasslands | Endangered | I |
| *Melica ciliata* agg. | NA | NA | NA | NA | NA | NA | NA | NA | 209 | -0.0003 | <.001 | -0.14 | Heaths, inland dunes and semi-natural grasslands | Near Threatened | I |
| *Ophrys apifera* | NA | NA | NA | NA | NA | NA | NA | NA | 198 | -0.0003 | <.001 | -0.16 | Heaths, inland dunes and semi-natural grasslands | Not Threatened | I |
| *Orchis pyramidalis* | NA | NA | NA | NA | NA | NA | NA | NA | 205 | -0.0003 | <.001 | -0.12 | Heaths, inland dunes and semi-natural grasslands | Endangered | I |
| *Phyteuma orbiculare* s. l. | NA | NA | NA | NA | NA | NA | NA | NA | 165 | -0.0003 | <.001 | -0.16 | Heaths, inland dunes and semi-natural grasslands | Endangered | I |
| *Salvia verticillata* | NA | NA | NA | NA | NA | NA | NA | NA | 258 | -0.0003 | <.001 | -0.14 | Heaths, inland dunes and semi-natural grasslands | Not Threatened | N |
| *Trifolium rubens* | NA | NA | NA | NA | NA | NA | NA | NA | 179 | -0.0003 | <.001 | -0.13 | Heaths, inland dunes and semi-natural grasslands | Endangered | I |
| *Amelanchier ovalis* s. l. | NA | NA | NA | NA | NA | NA | NA | NA | 111 | -0.0002 | <.001 | -0.07 | Dry to moderately moist forests | Not Threatened | I |
| *Aster bellidiastrum* | NA | NA | NA | NA | NA | NA | NA | NA | 68 | -0.0002 | <.001 | -0.07 | None | Not Threatened | I |
| *Carduus defloratus* | NA | NA | NA | NA | NA | NA | NA | NA | 85 | -0.0002 | <.001 | -0.09 | Dry to moderately moist forests | Near Threatened | I |
| *Carex pauciflora* | NA | NA | NA | NA | NA | NA | NA | NA | 76 | -0.0002 | <.001 | -0.10 | Bogs, transition mires, marshes and fens | Endangered | I |
| *Centaurea phrygia* agg. | NA | NA | NA | NA | NA | NA | NA | NA | 154 | -0.0002 | <.001 | -0.08 | Moist to wet grasslands | Endangered | I |
| *Cephalanthera rubra* | NA | NA | NA | NA | NA | NA | NA | NA | 136 | -0.0002 | <.001 | -0.08 | Dry to moderately moist forests | Near Threatened | I |
| *Coronilla coronata* | NA | NA | NA | NA | NA | NA | NA | NA | 114 | -0.0002 | <.001 | -0.11 | Dry to moderately moist forests | Endangered | I |
| *Crepis alpestris* | NA | NA | NA | NA | NA | NA | NA | NA | 92 | -0.0002 | <.001 | -0.11 | Heaths, inland dunes and semi-natural grasslands | Endangered | I |
| *Crepis mollis* | NA | NA | NA | NA | NA | NA | NA | NA | 160 | -0.0002 | <.001 | -0.10 | Moist to wet grasslands | Endangered | I |
| *Dianthus superbus* | NA | NA | NA | NA | NA | NA | NA | NA | 118 | -0.0002 | <.001 | -0.10 | Moist to wet grasslands | Endangered | I |
| *Digitalis grandiflora* | NA | NA | NA | NA | NA | NA | NA | NA | 214 | -0.0002 | <.001 | -0.11 | Dry to moderately moist forests | Near Threatened | I |
| *Galatella linosyris* | NA | NA | NA | NA | NA | NA | NA | NA | 112 | -0.0002 | <.001 | -0.09 | Heaths, inland dunes and semi-natural grasslands | Endangered | I |
| *Gentiana asclepiadea* | NA | NA | NA | NA | NA | NA | NA | NA | 56 | -0.0002 | <.001 | -0.08 | None | Not Threatened | I |
| *Gentiana cruciata* | NA | NA | NA | NA | NA | NA | NA | NA | 118 | -0.0002 | <.001 | -0.10 | Heaths, inland dunes and semi-natural grasslands | Highly Endangered | I |
| *Hieracium maculatum* | NA | NA | NA | NA | NA | NA | NA | NA | 114 | -0.0002 | <.001 | -0.08 | Heaths, inland dunes and semi-natural grasslands | Not Threatened | I |
| *Himantoglossum hircinum* | NA | NA | NA | NA | NA | NA | NA | NA | 244 | -0.0002 | <.001 | -0.11 | Heaths, inland dunes and semi-natural grasslands | Not Threatened | I |
| *Inula hirta* | NA | NA | NA | NA | NA | NA | NA | NA | 92 | -0.0002 | <.001 | -0.08 | Heaths, inland dunes and semi-natural grasslands | Endangered | I |
| *Iris germanica* agg. | NA | NA | NA | NA | NA | NA | NA | NA | 147 | -0.0002 | <.001 | -0.09 | Heaths, inland dunes and semi-natural grasslands | Not Threatened | A |
| *Jasione laevis* | NA | NA | NA | NA | NA | NA | NA | NA | 104 | -0.0002 | <.001 | -0.09 | Heaths, inland dunes and semi-natural grasslands | Endangered | I |
| *Melittis melissophyllum* | NA | NA | NA | NA | NA | NA | NA | NA | 297 | -0.0002 | <.001 | -0.10 | Dry to moderately moist forests | Endangered | I |
| *Noccaea montana* | NA | NA | NA | NA | NA | NA | NA | NA | 112 | -0.0002 | <.001 | -0.12 | Dry to moderately moist forests | Endangered | I |
| *Ophrys holoserica* | NA | NA | NA | NA | NA | NA | NA | NA | 100 | -0.0002 | <.001 | -0.08 | Heaths, inland dunes and semi-natural grasslands | Endangered | I |
| *Orchis purpurea* | NA | NA | NA | NA | NA | NA | NA | NA | 166 | -0.0002 | <.001 | -0.09 | Heaths, inland dunes and semi-natural grasslands | Near Threatened | I |
| *Orobanche teucrii* | NA | NA | NA | NA | NA | NA | NA | NA | 70 | -0.0002 | <.001 | -0.07 | Heaths, inland dunes and semi-natural grasslands | Endangered | I |
| *Polygala chamaebuxus* | NA | NA | NA | NA | NA | NA | NA | NA | 66 | -0.0002 | <.001 | -0.09 | Heaths, inland dunes and semi-natural grasslands | Not Threatened | I |
| *Scorzoneroides helvetica* | NA | NA | NA | NA | NA | NA | NA | NA | 70 | -0.0002 | <.001 | -0.09 | Heaths, inland dunes and semi-natural grasslands | Not Threatened | I |
| *Allium angulosum* | NA | NA | NA | NA | NA | NA | NA | NA | 34 | -0.0001 | <.001 | -0.03 | None | Endangered | I |
| *Allium lusitanicum* | NA | NA | NA | NA | NA | NA | NA | NA | 72 | -0.0001 | <.001 | -0.07 | None | Endangered | I |
| *Allium rotundum* | NA | NA | NA | NA | NA | NA | NA | NA | 106 | -0.0001 | <.001 | -0.05 | None | Endangered | I |
| *Allium sativum* | NA | NA | NA | NA | NA | NA | NA | NA | 98 | -0.0001 | <.001 | -0.05 | None | Not Evaluated | N |
| *Allium sphaerocephalon* | NA | NA | NA | NA | NA | NA | NA | NA | 58 | -0.0001 | <.001 | -0.04 | Heaths, inland dunes and semi-natural grasslands | Endangered | I |
| *Alnus alnobetula* | NA | NA | NA | NA | NA | NA | NA | NA | 102 | -0.0001 | <.001 | -0.05 | None | Not Threatened | I |
| *Althaea hirsuta* | NA | NA | NA | NA | NA | NA | NA | NA | 50 | -0.0001 | <.001 | -0.04 | None | Endangered | A |
| *Alyssum montanum* | NA | NA | NA | NA | NA | NA | NA | NA | 40 | -0.0001 | <.001 | -0.04 | Heaths, inland dunes and semi-natural grasslands | Endangered | I |
| *Anemone sylvestris* | NA | NA | NA | NA | NA | NA | NA | NA | 66 | -0.0001 | <.001 | -0.03 | Heaths, inland dunes and semi-natural grasslands | Endangered | I |
| *Astragalus cicer* | NA | NA | NA | NA | NA | NA | NA | NA | 62 | -0.0001 | <.001 | -0.03 | None | Near Threatened | I |
| *Astrantia major* | NA | NA | NA | NA | NA | NA | NA | NA | 191 | -0.0001 | <.001 | -0.05 | None | Not Threatened | I |
| *Atocion rupestre* | NA | NA | NA | NA | NA | NA | NA | NA | 134 | -0.0001 | <.001 | -0.05 | None | Not Threatened | I |
| *Bartsia alpina* | NA | NA | NA | NA | NA | NA | NA | NA | 20 | -0.0001 | <.001 | -0.04 | None | Not Threatened | I |
| *Bupleurum longifolium* | NA | NA | NA | NA | NA | NA | NA | NA | 56 | -0.0001 | <.001 | -0.03 | Dry to moderately moist forests | Endangered | I |
| *Calamagrostis varia* | NA | NA | NA | NA | NA | NA | NA | NA | 80 | -0.0001 | <.001 | -0.04 | Dry to moderately moist forests | Near Threatened | I |
| *Centaurea montana* | NA | NA | NA | NA | NA | NA | NA | NA | 109 | -0.0001 | <.001 | -0.07 | None | Not Threatened | I |
| *Cephalanthera longifolia* | NA | NA | NA | NA | NA | NA | NA | NA | 90 | -0.0001 | <.001 | -0.03 | None | Near Threatened | I |
| *Cicerbita alpina* | NA | NA | NA | NA | NA | NA | NA | NA | 47 | -0.0001 | <.001 | -0.03 | None | Not Threatened | I |
| *Cypripedium calceolus* | NA | NA | NA | NA | NA | NA | NA | NA | 61 | -0.0001 | <.001 | -0.06 | None | Endangered | I |
| *Daphne cneorum* | NA | NA | NA | NA | NA | NA | NA | NA | 28 | -0.0001 | <.001 | -0.04 | None | Highly Endangered | I |
| *Dianthus gratianopolitanus* | NA | NA | NA | NA | NA | NA | NA | NA | 28 | -0.0001 | <.001 | -0.03 | None | Endangered | I |
| *Dianthus sylvaticus* | NA | NA | NA | NA | NA | NA | NA | NA | 86 | -0.0001 | <.001 | -0.06 | None | Endangered | I |
| *Dictamnus albus* | NA | NA | NA | NA | NA | NA | NA | NA | 37 | -0.0001 | <.001 | -0.02 | None | Endangered | I |
| *Drosera anglica* | NA | NA | NA | NA | NA | NA | NA | NA | 15 | -0.0001 | <.001 | -0.03 | None | Highly Endangered | I |
| *Euphorbia seguieriana* | NA | NA | NA | NA | NA | NA | NA | NA | 80 | -0.0001 | <.001 | -0.05 | Heaths, inland dunes and semi-natural grasslands | Endangered | I |
| *Galeopsis ladanum* agg. | NA | NA | NA | NA | NA | NA | NA | NA | 146 | -0.0001 | <.001 | -0.06 | None | Not Threatened | I |
| *Gymnadenia odoratissima* | NA | NA | NA | NA | NA | NA | NA | NA | 66 | -0.0001 | <.001 | -0.06 | Heaths, inland dunes and semi-natural grasslands | Endangered | I |
| *Juncus alpinoarticulatus* | NA | NA | NA | NA | NA | NA | NA | NA | 42 | -0.0001 | <.001 | -0.03 | None | Near Threatened | I |
| *Lathyrus heterophyllus* | NA | NA | NA | NA | NA | NA | NA | NA | 112 | -0.0001 | <.001 | -0.05 | Heaths, inland dunes and semi-natural grasslands | Endangered | I |
| *Odontites luteus* | NA | NA | NA | NA | NA | NA | NA | NA | 40 | -0.0001 | <.001 | -0.03 | None | Endangered | I |
| *Ophrys sphegodes* agg. | NA | NA | NA | NA | NA | NA | NA | NA | 38 | -0.0001 | <.001 | -0.04 | Heaths, inland dunes and semi-natural grasslands | Highly Endangered | I |
| *Orchis ustulata* | NA | NA | NA | NA | NA | NA | NA | NA | 76 | -0.0001 | <.001 | -0.06 | Heaths, inland dunes and semi-natural grasslands | Highly Endangered | I |
| *Orobanche lutea* | NA | NA | NA | NA | NA | NA | NA | NA | 67 | -0.0001 | <.001 | -0.04 | Heaths, inland dunes and semi-natural grasslands | Endangered | I |
| *Orthilia secunda* | NA | NA | NA | NA | NA | NA | NA | NA | 63 | -0.0001 | <.001 | -0.06 | None | Near Threatened | I |
| *Peucedanum alsaticum* | NA | NA | NA | NA | NA | NA | NA | NA | 45 | -0.0001 | <.001 | -0.04 | None | Endangered | I |
| *Potentilla aurea* | NA | NA | NA | NA | NA | NA | NA | NA | 18 | -0.0001 | <.001 | -0.02 | None | Not Threatened | I |
| *Pseudorchis albida* | NA | NA | NA | NA | NA | NA | NA | NA | 22 | -0.0001 | <.001 | -0.03 | None | Endangered | I |
| *Ranunculus montanus* agg. | NA | NA | NA | NA | NA | NA | NA | NA | 52 | -0.0001 | <.001 | -0.06 | None | Not Threatened | I |
| *Rhamnus saxatilis* | NA | NA | NA | NA | NA | NA | NA | NA | 23 | -0.0001 | <.001 | -0.03 | None | Not Threatened | I |
| *Saxifraga paniculata* | NA | NA | NA | NA | NA | NA | NA | NA | 57 | -0.0001 | <.001 | -0.05 | Dry to moderately moist forests | Not Threatened | I |
| *Scabiosa canescens* | NA | NA | NA | NA | NA | NA | NA | NA | 44 | -0.0001 | <.001 | -0.03 | Heaths, inland dunes and semi-natural grasslands | Endangered | I |
| *Schoenus ferrugineus* | NA | NA | NA | NA | NA | NA | NA | NA | 54 | -0.0001 | <.001 | -0.07 | None | Endangered | I |
| *Sempervivum tectorum* | NA | NA | NA | NA | NA | NA | NA | NA | 62 | -0.0001 | <.001 | -0.03 | None | Not Threatened | I |
| *Stachys germanica* | NA | NA | NA | NA | NA | NA | NA | NA | 66 | -0.0001 | <.001 | -0.03 | Heaths, inland dunes and semi-natural grasslands | Endangered | I |
| *Taraxacum* sect. *Erythrosperma* | NA | NA | NA | NA | NA | NA | NA | NA | 42 | -0.0001 | <.001 | -0.03 | Heaths, inland dunes and semi-natural grasslands | Not Threatened | I |
| *Tephroseris helenitis* | NA | NA | NA | NA | NA | NA | NA | NA | 58 | -0.0001 | <.001 | -0.05 | Moist to wet grasslands | Highly Endangered | I |
| *Thesium linophyllon* | NA | NA | NA | NA | NA | NA | NA | NA | 40 | -0.0001 | <.001 | -0.05 | Heaths, inland dunes and semi-natural grasslands | Endangered | I |
| *Tofieldia calyculata* | NA | NA | NA | NA | NA | NA | NA | NA | 42 | -0.0001 | <.001 | -0.07 | None | Endangered | I |
| *Trifolium ochroleucon* | NA | NA | NA | NA | NA | NA | NA | NA | 51 | -0.0001 | <.001 | -0.03 | Heaths, inland dunes and semi-natural grasslands | Highly Endangered | I |
| *Trifolium spadiceum* | NA | NA | NA | NA | NA | NA | NA | NA | 36 | -0.0001 | <.001 | -0.03 | None | Highly Endangered | I |
| *Veratrum album* s. l. | NA | NA | NA | NA | NA | NA | NA | NA | 60 | -0.0001 | <.001 | -0.03 | None | Not Threatened | I |
| *Adonis aestivalis* | NA | NA | NA | NA | NA | NA | NA | NA | 17 | 0.0000 | <.001 | -0.02 | None | Highly Endangered | A |
| *Allium suaveolens* | NA | NA | NA | NA | NA | NA | NA | NA | 20 | 0.0000 | <.001 | -0.02 | None | Endangered | I |
| *Anagallis foemina* | NA | NA | NA | NA | NA | NA | NA | NA | 16 | 0.0000 | <.001 | -0.02 | None | Endangered | I |
| *Anthericum liliago* | NA | NA | NA | NA | NA | NA | NA | NA | 52 | 0.0000 | <.001 | -0.02 | None | Near Threatened | I |
| *Asperula tinctoria* | NA | NA | NA | NA | NA | NA | NA | NA | 18 | 0.0000 | <.001 | -0.01 | None | Endangered | I |
| *Carex buxbaumii* agg. | NA | NA | NA | NA | NA | NA | NA | NA | 20 | 0.0000 | <.001 | -0.02 | Moist to wet grasslands | Highly Endangered | I |
| *Carex sempervirens* | NA | NA | NA | NA | NA | NA | NA | NA | 13 | 0.0000 | <.001 | -0.02 | None | Not Threatened | I |
| *Caucalis platycarpos* | NA | NA | NA | NA | NA | NA | NA | NA | 8 | 0.0000 | <.001 | -0.01 | None | Highly Endangered | A |
| *Corallorhiza trifida* | NA | NA | NA | NA | NA | NA | NA | NA | 19 | 0.0000 | <.001 | -0.02 | None | Endangered | I |
| *Coronilla vaginalis* | NA | NA | NA | NA | NA | NA | NA | NA | 11 | 0.0000 | <.001 | -0.01 | None | Endangered | I |
| *Cotoneaster tomentosus* | NA | NA | NA | NA | NA | NA | NA | NA | 14 | 0.0000 | <.001 | -0.01 | None | Near Threatened | I |
| *Dioscorea communis* | NA | NA | NA | NA | NA | NA | NA | NA | 39 | 0.0000 | <.001 | 0.02 | None | Near Threatened | I |
| *Draba aizoides* | NA | NA | NA | NA | NA | NA | NA | NA | 16 | 0.0000 | <.001 | -0.02 | None | Endangered | I |
| *Festuca amethystina* | NA | NA | NA | NA | NA | NA | NA | NA | 17 | 0.0000 | <.001 | -0.02 | None | Not Threatened | I |
| *Goodyera repens* | NA | NA | NA | NA | NA | NA | NA | NA | 20 | 0.0000 | <.001 | -0.02 | None | Endangered | I |
| *Herminium monorchis* | NA | NA | NA | NA | NA | NA | NA | NA | 10 | 0.0000 | <.001 | -0.01 | None | Highly Endangered | I |
| *Hieracium humile* | NA | NA | NA | NA | NA | NA | NA | NA | 22 | 0.0000 | <.001 | -0.02 | None | Not Threatened | I |
| *Juglans nigra* | NA | NA | NA | NA | NA | NA | NA | NA | 28 | 0.0000 | <.001 | 0.02 | None | Not Evaluated | N |
| *Lathyrus aphaca* | NA | NA | NA | NA | NA | NA | NA | NA | 38 | 0.0000 | <.001 | -0.02 | None | Endangered | A |
| *Linum perenne* agg. | NA | NA | NA | NA | NA | NA | NA | NA | 22 | 0.0000 | <.001 | -0.01 | None | Endangered | NA |
| *Listera cordata* | NA | NA | NA | NA | NA | NA | NA | NA | 12 | 0.0000 | <.001 | -0.01 | None | Endangered | I |
| *Moneses uniflora* | NA | NA | NA | NA | NA | NA | NA | NA | 18 | 0.0000 | <.001 | -0.02 | None | Highly Endangered | I |
| *Orchis anthropophora* | NA | NA | NA | NA | NA | NA | NA | NA | 27 | 0.0000 | <.001 | -0.02 | None | Endangered | I |
| *Orchis simia* | NA | NA | NA | NA | NA | NA | NA | NA | 29 | 0.0000 | <.001 | -0.02 | Heaths, inland dunes and semi-natural grasslands | Highly Endangered | I |
| *Orobanche minor* | NA | NA | NA | NA | NA | NA | NA | NA | 13 | 0.0000 | <.001 | -0.01 | None | Endangered | A |
| *Peucedanum officinale* | NA | NA | NA | NA | NA | NA | NA | NA | 24 | 0.0000 | <.001 | -0.02 | None | Endangered | I |
| *Ranunculus arvensis* | NA | NA | NA | NA | NA | NA | NA | NA | 13 | 0.0000 | <.001 | -0.01 | None | Endangered | A |
| *Rumex alpinus* | NA | NA | NA | NA | NA | NA | NA | NA | 19 | 0.0000 | <.001 | -0.02 | None | Not Threatened | I |
| *Rumex scutatus* | NA | NA | NA | NA | NA | NA | NA | NA | 14 | 0.0000 | <.001 | -0.01 | None | Not Threatened | I |
| *Schoenus intermedius* | NA | NA | NA | NA | NA | NA | NA | NA | 16 | 0.0000 | <.001 | -0.02 | None | Endangered | I |
| *Scutellaria minor* | NA | NA | NA | NA | NA | NA | NA | NA | 25 | 0.0000 | <.001 | -0.02 | None | Highly Endangered | I |
| *Seseli annuum* | NA | NA | NA | NA | NA | NA | NA | NA | 12 | 0.0000 | <.001 | -0.01 | None | Endangered | I |
| *Thymelaea passerina* | NA | NA | NA | NA | NA | NA | NA | NA | 11 | 0.0000 | <.001 | -0.01 | None | Highly Endangered | I |
| *Torilis arvensis* | NA | NA | NA | NA | NA | NA | NA | NA | 120 | 0.0001 | <.001 | 0.05 | None | Not Threatened | A |
| *Viscum album* agg. | NA | NA | NA | NA | NA | NA | NA | NA | 278 | 0.0003 | <.001 | 0.15 | None | Not Threatened | I |
| *Polystichum aculeatum* agg. | NA | NA | NA | NA | NA | NA | NA | NA | 430 | 0.0004 | <.001 | 0.20 | Dry to moderately moist forests | Not Threatened | I |
| *Rosa arvensis* | NA | NA | NA | NA | NA | NA | NA | NA | 2290 | 0.0013 | <.001 | 0.58 | Scrubs, copses and field hedges | Near Threatened | I |

.
 Notes: Given are raw and scaled Beals trends of species within each state, including n-, and p- values (after Holm adjustment) according to t-tests. In addition, each species
 preferred habitat type and its Red list (RL) and non-native (NN; I = Native (indigenous), N = Neophytes, A = Archaeophytes, NA = no status assigned) status in Germany are given.
 Habitat type preferences are based on the fidelity (Φ) of species to habitat types in all states taken together.
