## Appendix S3 for "Loss of characteristic species across German federal states detected by repeated mapping of protected habitats"

**Table S3** Frequency trends of all species that showed a significant trend in at least one state.

| **Species** | **n (SH)** | **Freq trend (SH)** | **p (SH)** | **n (HH)** | **Freq trend (HH)** | **p (HH)** | **n (BW)** | **Freq trend (BW)** | **p (BW)** | **Habitat type preferred** | **RL** | **NN** |
| --- | --- | --- | --- | --- | --- | --- | --- | --- | --- | --- | --- | --- |
| *Echinops sphaerocephalus* | 1 | -1.00 | NA | 2 | 0.00 | NA | 214 | 0.24 | <.001 | None | Not Evaluated | N |
| *Dipsacus pilosus* | 1 | -1.00 | NA | 1 | 1.00 | NA | 160 | 0.43 | <.001 | Moist to wet forests | Not Threatened | I |
| *Setaria pumila* | 1 | -1.00 | NA | 2 | 1.00 | NA | 32 | 0.85 | <.001 | None | Not Threatened | A |
| *Epipactis purpurata* | 10 | -1.00 | 0.002 | NA | NA | NA | 48 | -0.65 | <.001 | None | Near Threatened | I |
| *Trifolium aureum* | 1 | -1.00 | NA | NA | NA | NA | 40 | -0.47 | 0.005 | Heaths, inland dunes and semi-natural grasslands | Near Threatened | I |
| *Cruciata laevipes* | 3 | -1.00 | NA | NA | NA | NA | 466 | -0.33 | <.001 | Scrubs, copses and field hedges | Not Threatened | I |
| *Genista germanica* | 1 | -1.00 | NA | NA | NA | NA | 112 | -0.30 | <.001 | Heaths, inland dunes and semi-natural grasslands | Endangered | I |
| *Geranium pyrenaicum* | 2 | -1.00 | NA | NA | NA | NA | 300 | -0.20 | <.001 | Scrubs, copses and field hedges | Not Evaluated | N |
| *Galium sylvaticum* agg. | 8 | -1.00 | 0.008 | NA | NA | NA | 1208 | -0.18 | <.001 | Dry to moderately moist forests | Not Threatened | I |
| *Hypericum montanum* | 6 | -1.00 | 0.031 | NA | NA | NA | 168 | 0.06 | NA | Dry to moderately moist forests | Near Threatened | I |
| *Arabis hirsuta* agg. | 2 | -1.00 | NA | NA | NA | NA | 318 | 0.16 | 0.003 | Heaths, inland dunes and semi-natural grasslands | Not Threatened | I |
| *Salix myrsinifolia* | 2 | -1.00 | NA | NA | NA | NA | 225 | 0.20 | <.001 | None | Near Threatened | I |
| *Adenostyles alliariae* | 1 | -1.00 | NA | NA | NA | NA | 229 | 0.22 | <.001 | None | Not Threatened | I |
| *Festuca heterophylla* | 1 | -1.00 | NA | NA | NA | NA | 178 | 0.25 | <.001 | Dry to moderately moist forests | Near Threatened | I |
| *Sorbus latifolia* agg. | 1 | -1.00 | NA | NA | NA | NA | 35 | 1.00 | <.001 | None | Not Threatened | I |
| *Sedum telephium* | 10 | -1.00 | 0.002 | NA | NA | NA | NA | NA | NA | None | Not Threatened | I |
| *Hypericum humifusum* | 13 | -1.00 | <.001 | 9 | -0.62 | NA | 30 | 0.05 | NA | None | Not Threatened | I |
| *Rosa dumalis* agg. | 12 | -1.00 | <.001 | 1 | 1.00 | NA | 934 | -0.13 | NA | Scrubs, copses and field hedges | Not Threatened | I |
| *Clinopodium vulgare* | 14 | -0.98 | <.001 | 1 | -1.00 | NA | 2348 | 0.01 | NA | Heaths, inland dunes and semi-natural grasslands | Not Threatened | I |
| *Anagallis arvensis* | 4 | -0.97 | NA | 8 | -0.34 | NA | 26 | -0.75 | <.001 | None | Not Threatened | A |
| *Sinapis arvensis* | 2 | -0.95 | NA | 6 | -0.61 | NA | 24 | -0.88 | <.001 | None | Not Threatened | A |
| *Allium oleraceum* | 18 | -0.92 | <.001 | 1 | 1.00 | NA | 646 | -0.08 | NA | Scrubs, copses and field hedges | Not Threatened | I |
| *Symphytum asperum* agg. | 45 | -0.90 | <.001 | 2 | -0.50 | NA | NA | NA | NA | None | Not Evaluated | N |
| *Carex montana* | 12 | -0.90 | <.001 | NA | NA | NA | 1214 | 0.08 | NA | Dry to moderately moist forests | Not Threatened | I |
| *Tephroseris palustris* | 19 | -0.88 | <.001 | 2 | -1.00 | NA | NA | NA | NA | None | Near Threatened | I |
| *Lathyrus vernus* | 10 | -0.87 | 0.002 | 1 | 1.00 | NA | 874 | -0.14 | NA | Dry to moderately moist forests | Not Threatened | I |
| *Potentilla recta* | 78 | -0.87 | <.001 | 4 | -0.50 | NA | 104 | -0.10 | NA | Heaths, inland dunes and semi-natural grasslands | Not Threatened | A |
| *Antennaria dioica* | 4 | -0.87 | NA | NA | NA | NA | 112 | -0.39 | <.001 | Heaths, inland dunes and semi-natural grasslands | Endangered | I |
| *Gnaphalium sylvaticum* | 24 | -0.86 | <.001 | 44 | -0.66 | <.001 | 52 | 0.15 | NA | Heaths, inland dunes and semi-natural grasslands | Not Threatened | I |
| *Orchis mascula* | 98 | -0.86 | <.001 | NA | NA | NA | 260 | -0.36 | <.001 | Heaths, inland dunes and semi-natural grasslands | Near Threatened | I |
| *Campanula trachelium* | 133 | -0.85 | <.001 | 2 | 1.00 | NA | 1874 | -0.08 | NA | Scrubs, copses and field hedges | Not Threatened | I |
| *Pedicularis palustris* | 44 | -0.83 | <.001 | 2 | -0.50 | NA | 48 | -0.10 | NA | None | Highly Endangered | I |
| *Helictotrichon pratense* | 38 | -0.83 | <.001 | NA | NA | NA | 1407 | 0.05 | NA | Heaths, inland dunes and semi-natural grasslands | Near Threatened | I |
| *Platanthera chlorantha* | 172 | -0.82 | <.001 | 6 | -0.65 | NA | 340 | 0.11 | NA | Heaths, inland dunes and semi-natural grasslands | Endangered | I |
| *Sagina nodosa* | 22 | -0.82 | <.001 | NA | NA | NA | NA | NA | NA | Coastal and marine habitats | Highly Endangered | I |
| *Sagina procumbens* | 135 | -0.82 | <.001 | 14 | -0.39 | NA | 7 | 0.09 | NA | None | Not Threatened | I |
| *Achillea ptarmica* agg. | 515 | -0.80 | <.001 | 104 | -0.51 | <.001 | 819 | 0.03 | NA | Moist to wet grasslands | Not Threatened | I |
| *Thymus serpyllum* | 38 | -0.80 | <.001 | NA | NA | NA | 24 | -0.58 | 0.004 | Coastal and marine habitats | Near Threatened | I |
| *Campanula latifolia* | 106 | -0.79 | <.001 | NA | NA | NA | 14 | -0.62 | NA | Dry to moderately moist forests | Not Threatened | I |
| *Lathyrus linifolius* | 16 | -0.79 | 0.004 | NA | NA | NA | 344 | 0.24 | <.001 | Heaths, inland dunes and semi-natural grasslands | Near Threatened | I |
| *Gentiana pneumonanthe* | 30 | -0.78 | <.001 | 5 | -0.84 | NA | 25 | 0.02 | NA | None | Highly Endangered | I |
| *Dactylorhiza majalis* agg. | 294 | -0.77 | <.001 | 16 | -0.44 | NA | 1059 | -0.04 | NA | Moist to wet grasslands | Near Threatened | I |
| *Listera ovata* | 93 | -0.77 | <.001 | 4 | -0.71 | NA | 711 | -0.17 | <.001 | Heaths, inland dunes and semi-natural grasslands | Not Threatened | I |
| *Cochlearia danica* | 24 | -0.76 | <.001 | 1 | 1.00 | NA | NA | NA | NA | Coastal and marine habitats | Not Threatened | I |
| *Rhinanthus minor* | 83 | -0.74 | <.001 | 4 | 0.00 | NA | 742 | 0.18 | <.001 | Heaths, inland dunes and semi-natural grasslands | Not Threatened | I |
| *Centaurea nigra* s. l. | 8 | -0.74 | NA | 1 | 1.00 | NA | 618 | 0.40 | <.001 | Heaths, inland dunes and semi-natural grasslands | Not Threatened | I |
| *Cuscuta europaea* | 13 | -0.74 | NA | 18 | -0.09 | NA | 57 | -0.48 | <.001 | None | Not Threatened | I |
| *Myosotis sylvatica* agg. | 16 | -0.73 | NA | 1 | -1.00 | NA | 94 | -0.59 | <.001 | None | Not Threatened | I |
| *Blysmus compressus* | 22 | -0.73 | <.001 | NA | NA | NA | 8 | -0.42 | NA | None | Highly Endangered | I |
| *Succisa pratensis* | 216 | -0.73 | <.001 | 11 | -0.21 | NA | 1353 | -0.11 | NA | Moist to wet grasslands | Near Threatened | I |
| *Mycelis muralis* | 222 | -0.72 | <.001 | 43 | 0.53 | <.001 | 1400 | -0.06 | NA | Dry to moderately moist forests | Not Threatened | I |
| *Scorzonera humilis* | 24 | -0.72 | <.001 | NA | NA | NA | 91 | 0.10 | NA | Moist to wet grasslands | Endangered | I |
| *Arnica montana* | 30 | -0.71 | NA | 1 | -1.00 | NA | 484 | -0.19 | <.001 | Heaths, inland dunes and semi-natural grasslands | Endangered | I |
| *Cicuta virosa* | 202 | -0.71 | <.001 | 1 | 1.00 | NA | 18 | -0.28 | NA | Standing waters | Near Threatened | I |
| *Bidens tripartita* | 228 | -0.69 | <.001 | 73 | -0.39 | <.001 | 26 | -0.14 | NA | Standing waters | Not Threatened | I |
| *Sanicula europaea* | 348 | -0.69 | <.001 | 2 | 0.25 | NA | 456 | -0.20 | <.001 | Dry to moderately moist forests | Not Threatened | I |
| *Viola hirta* | 4 | -0.69 | NA | NA | NA | NA | 1829 | 0.18 | <.001 | Heaths, inland dunes and semi-natural grasslands | Not Threatened | I |
| *Dactylorhiza incarnata* agg. | 30 | -0.69 | <.001 | 2 | -0.25 | NA | 182 | -0.13 | NA | Moist to wet grasslands | Endangered | I |
| *Triglochin palustris* | 276 | -0.68 | <.001 | 7 | -0.27 | NA | 46 | -0.20 | NA | Coastal and marine habitats | Endangered | I |
| *Carex appropinquata* | 100 | -0.68 | <.001 | 4 | 0.80 | NA | 102 | 0.10 | NA | Fresh water vegetation, springs and reeds | Endangered | I |
| *Sanguisorba officinalis* | 64 | -0.67 | <.001 | NA | NA | NA | 2707 | 0.06 | NA | Moist to wet grasslands | Near Threatened | I |
| *Alopecurus myosuroides* | 3 | -0.67 | NA | 2 | 0.00 | NA | 38 | -0.70 | <.001 | None | Not Threatened | A |
| *Juncus filiformis* | 180 | -0.66 | <.001 | 17 | -0.15 | NA | 272 | -0.05 | NA | Moist to wet grasslands | Near Threatened | I |
| *Ophioglossum vulgatum* | 37 | -0.65 | <.001 | 1 | -1.00 | NA | 26 | -0.18 | NA | Coastal and marine habitats | Endangered | I |
| *Melilotus officinalis* | 37 | -0.65 | <.001 | 37 | -0.34 | NA | 438 | -0.30 | <.001 | Ruderal, fringe and tall forb communities, clearings | Not Threatened | A |
| *Oxalis stricta* | 10 | -0.65 | NA | NA | NA | NA | 24 | 0.53 | 0.023 | None | Not Evaluated | N |
| *Geranium palustre* | 70 | -0.64 | <.001 | 1 | -1.00 | NA | 1274 | 0.07 | NA | Fresh water vegetation, springs and reeds | Not Threatened | I |
| *Bromus racemosus* agg. | 62 | -0.64 | <.001 | 10 | 0.72 | NA | 150 | -0.18 | NA | Moist to wet grasslands | Vulnerable | NA |
| *Dactylorhiza maculata* agg. | 125 | -0.63 | <.001 | 16 | -0.52 | NA | 777 | -0.08 | NA | Moist to wet grasslands | Near Threatened | I |
| *Galinsoga parviflora* | 4 | -0.62 | NA | 24 | -0.48 | 0.019 | 4 | 0.27 | NA | Anthropogenic | Not Evaluated | N |
| *Viola palustris* | 764 | -0.62 | <.001 | 64 | -0.46 | <.001 | 1558 | -0.00 | NA | Moist to wet grasslands | Not Threatened | I |
| *Menyanthes trifoliata* | 396 | -0.61 | <.001 | 13 | -0.43 | NA | 584 | -0.07 | NA | Bogs, transition mires, marshes and fens | Endangered | I |
| *Chenopodium album* agg. | 32 | -0.60 | NA | 52 | -0.35 | 0.004 | 118 | 0.08 | NA | Anthropogenic | Not Threatened | NA |
| *Pulmonaria officinalis* agg. | 381 | -0.59 | <.001 | 1 | -1.00 | NA | 974 | -0.06 | NA | Dry to moderately moist forests | Not Threatened | I |
| *Rosa villosa* agg. | 6 | -0.58 | NA | 1 | -1.00 | NA | 199 | -0.27 | <.001 | Scrubs, copses and field hedges | Near Threatened | I |
| *Carex panicea* | 486 | -0.58 | <.001 | 25 | -0.13 | NA | 2702 | -0.09 | NA | Moist to wet grasslands | Near Threatened | I |
| *Briza media* | 78 | -0.58 | <.001 | 4 | -0.20 | NA | 3954 | 0.05 | NA | Heaths, inland dunes and semi-natural grasslands | Not Threatened | I |
| *Senecio paludosus* | 5 | -0.58 | NA | 10 | 0.45 | NA | 107 | -0.29 | <.001 | Fresh water vegetation, springs and reeds | Endangered | I |
| *Hieracium murorum* | 74 | -0.57 | <.001 | 10 | -0.78 | 0.008 | 1318 | 0.01 | NA | Dry to moderately moist forests | Not Threatened | I |
| *Astragalus glycyphyllos* | 41 | -0.57 | NA | 2 | -0.12 | NA | 1037 | 0.11 | <.001 | Heaths, inland dunes and semi-natural grasslands | Not Threatened | I |
| *Hydrocotyle vulgaris* | 929 | -0.57 | <.001 | 60 | -0.35 | NA | 15 | 0.01 | NA | Bogs, transition mires, marshes and fens | Not Threatened | I |
| *Carex flava* agg. | 152 | -0.56 | <.001 | 16 | -0.58 | 0.013 | 998 | -0.10 | NA | Moist to wet grasslands | Not Threatened | I |
| *Melilotus altissimus* | 15 | -0.56 | NA | 4 | -0.80 | NA | 43 | -0.39 | 0.017 | None | Not Threatened | I |
| *Ranunculus lingua* | 148 | -0.56 | <.001 | 12 | 0.01 | NA | 46 | -0.15 | NA | Standing waters | Endangered | I |
| *Neottia nidus-avis* | 11 | -0.56 | NA | NA | NA | NA | 384 | -0.20 | <.001 | Dry to moderately moist forests | Not Threatened | I |
| *Senecio aquaticus* agg. | 149 | -0.55 | <.001 | 53 | 0.27 | NA | 1142 | -0.11 | NA | Moist to wet grasslands | Near Threatened | I |
| *Salix repens* agg. | 256 | -0.54 | <.001 | 7 | -0.60 | NA | 107 | -0.31 | <.001 | Coastal and marine habitats | Not Threatened | I |
| *Sium latifolium* | 178 | -0.53 | <.001 | 22 | -0.13 | NA | 4 | 0.10 | NA | Linear and running surface waters | Not Threatened | I |
| *Hieracium sabaudum* | 61 | -0.53 | NA | 50 | 0.22 | NA | 195 | -0.20 | 0.002 | Dry to moderately moist forests | Not Threatened | I |
| *Lychnis flos-cuculi* | 1180 | -0.53 | <.001 | 154 | -0.24 | <.001 | 2890 | 0.05 | NA | Moist to wet grasslands | Not Threatened | I |
| *Trichophorum cespitosum* agg. | 106 | -0.52 | <.001 | 8 | -0.28 | NA | 76 | -0.12 | NA | Bogs, transition mires, marshes and fens | Endangered | I |
| *Mentha arvensis* | 84 | -0.51 | NA | 82 | -0.46 | <.001 | 234 | -0.15 | NA | Moist to wet grasslands | Not Threatened | I |
| *Anthyllis vulneraria* s. l. | 34 | -0.51 | NA | 2 | 0.50 | NA | 1374 | -0.16 | <.001 | Heaths, inland dunes and semi-natural grasslands | Not Threatened | I |
| *Stellaria aquatica* | 316 | -0.51 | <.001 | 44 | -0.18 | NA | 244 | -0.19 | NA | Moist to wet forests | Not Threatened | I |
| *Platanthera bifolia* s. l. | 19 | -0.50 | NA | NA | NA | NA | 510 | -0.32 | <.001 | Heaths, inland dunes and semi-natural grasslands | Endangered | I |
| *Hypopitys monotropa* agg. | 2 | -0.50 | NA | NA | NA | NA | 74 | -0.56 | <.001 | None | Not Threatened | I |
| *Diplotaxis tenuifolia* | 2 | -0.50 | NA | NA | NA | NA | 138 | 0.14 | 0.042 | None | Not Evaluated | N |
| *Filago arvensis* | 15 | -0.50 | NA | 6 | 0.69 | NA | 26 | -0.78 | <.001 | Heaths, inland dunes and semi-natural grasslands | Not Threatened | I |
| *Phyteuma spicatum* agg. | 348 | -0.49 | <.001 | 4 | 0.25 | NA | 1438 | -0.20 | <.001 | Dry to moderately moist forests | Not Threatened | I |
| *Caltha palustris* | 1842 | -0.48 | <.001 | 116 | -0.31 | NA | 6463 | -0.13 | <.001 | Moist to wet grasslands | Near Threatened | I |
| *Cladium mariscus* | 20 | -0.48 | NA | 7 | -1.00 | 0.016 | 44 | 0.17 | NA | Fresh water vegetation, springs and reeds | Endangered | I |
| *Alisma plantago-aquatica* agg. | 826 | -0.47 | <.001 | 134 | -0.16 | NA | 513 | -0.22 | <.001 | Standing waters | Not Threatened | I |
| *Geranium columbinum* | 4 | -0.47 | NA | NA | NA | NA | 267 | -0.38 | <.001 | None | Not Threatened | A |
| *Thalictrum minus* agg. | 2 | -0.46 | NA | NA | NA | NA | 69 | -0.22 | 0.023 | Heaths, inland dunes and semi-natural grasslands | Near Threatened | I |
| *Comarum palustre* | 1114 | -0.46 | <.001 | 82 | -0.22 | NA | 648 | -0.05 | NA | Bogs, transition mires, marshes and fens | Not Threatened | I |
| *Corynephorus canescens* | 167 | -0.46 | NA | 21 | -0.04 | NA | 39 | -0.31 | 0.021 | Coastal and marine habitats | Not Threatened | I |
| *Rhinanthus serotinus* s. l. | 168 | -0.45 | NA | 25 | -0.18 | NA | 27 | -0.66 | <.001 | Mesic grasslands | Endangered | I |
| *Veronica persica* | 5 | -0.45 | NA | 4 | -0.57 | NA | 70 | -0.65 | <.001 | None | Not Evaluated | N |
| *Juncus conglomeratus* | 524 | -0.45 | <.001 | 82 | -0.05 | NA | 1048 | -0.21 | <.001 | Moist to wet grasslands | Not Threatened | I |
| *Schoenoplectus lacustris* agg. | 652 | -0.44 | NA | 40 | -0.11 | NA | 344 | -0.16 | <.001 | Standing waters | Not Threatened | I |
| *Lycopodium clavatum* | 17 | -0.42 | NA | 2 | -0.25 | NA | 36 | -0.50 | 0.001 | None | Endangered | I |
| *Carex digitata* | 23 | -0.42 | NA | NA | NA | NA | 656 | 0.21 | <.001 | Dry to moderately moist forests | Not Threatened | I |
| *Apera spica-venti* | 6 | -0.41 | NA | 8 | -0.63 | NA | 12 | -0.89 | <.001 | Farmland | Not Threatened | I |
| *Heracleum sphondylium* | 378 | -0.41 | NA | 353 | -0.19 | NA | 4150 | -0.34 | <.001 | Scrubs, copses and field hedges | Not Threatened | I |
| *Galium uliginosum* | 402 | -0.41 | NA | 35 | -0.05 | NA | 1536 | 0.15 | <.001 | Moist to wet grasslands | Not Threatened | I |
| *Anthemis arvensis* | 4 | -0.40 | NA | 1 | -1.00 | NA | 12 | -0.72 | 0.012 | None | Near Threatened | A |
| *Melilotus albus* | 34 | -0.40 | NA | 50 | -0.35 | 0.014 | 346 | -0.29 | <.001 | Ruderal, fringe and tall forb communities, clearings | Not Threatened | A |
| *Solidago virgaurea* | 53 | -0.39 | NA | 33 | -0.54 | 0.002 | 1683 | 0.00 | NA | Dry to moderately moist forests | Not Threatened | I |
| *Tussilago farfara* | 198 | -0.39 | NA | 148 | -0.40 | <.001 | 334 | -0.36 | <.001 | Ruderal, fringe and tall forb communities, clearings | Not Threatened | I |
| *Fallopia convolvulus* | 17 | -0.38 | NA | 18 | -0.47 | NA | 90 | -0.32 | 0.001 | Farmland | Not Threatened | A |
| *Elymus caninus* | 48 | -0.38 | NA | 18 | -0.95 | <.001 | 616 | 0.04 | NA | Moist to wet forests | Not Threatened | I |
| *Teucrium scorodonia* | 4 | -0.38 | NA | 6 | -0.29 | NA | 2246 | 0.11 | <.001 | Heaths, inland dunes and semi-natural grasslands | Not Threatened | I |
| *Juncus articulatus* | 1290 | -0.37 | NA | 176 | -0.06 | NA | 1543 | -0.15 | <.001 | Moist to wet grasslands | Not Threatened | I |
| *Chondrilla juncea* | 8 | -0.36 | NA | NA | NA | NA | 38 | 0.34 | 0.019 | Coastal and marine habitats | Not Threatened | I |
| *Crepis paludosa* | 961 | -0.36 | NA | 50 | 0.04 | NA | 2080 | 0.13 | <.001 | Moist to wet grasslands | Not Threatened | I |
| *Drosera rotundifolia* | 388 | -0.35 | NA | 12 | -0.11 | NA | 238 | -0.22 | <.001 | Bogs, transition mires, marshes and fens | Endangered | I |
| *Eleocharis palustris* agg. | 909 | -0.35 | NA | 79 | -0.04 | NA | 605 | -0.20 | <.001 | Standing waters | Not Threatened | I |
| *Silene dioica* | 634 | -0.35 | NA | 50 | -0.05 | NA | 2386 | -0.17 | <.001 | Moist to wet forests | Not Threatened | I |
| *Cardamine pratensis* agg. | 1324 | -0.34 | NA | 194 | 0.17 | 0.005 | 1424 | -0.24 | <.001 | Moist to wet grasslands | Not Threatened | I |
| *Sisymbrium altissimum* | 2 | -0.33 | NA | 29 | -0.61 | <.001 | 1 | 1.00 | NA | Ruderal, fringe and tall forb communities, clearings | Not Evaluated | N |
| *Primula elatior* | 1262 | -0.33 | NA | 12 | 0.13 | NA | 3228 | -0.12 | <.001 | Moist to wet forests | Not Threatened | I |
| *Juncus bufonius* agg. | 308 | -0.33 | NA | 67 | 0.08 | NA | 130 | -0.37 | <.001 | Standing waters | Not Threatened | I |
| *Epipactis palustris* | 14 | -0.33 | NA | 1 | -0.50 | NA | 207 | -0.27 | <.001 | Moist to wet grasslands | Endangered | I |
| *Asparagus officinalis* | 10 | -0.32 | NA | 10 | -0.33 | NA | 157 | 0.23 | 0.002 | None | Not Threatened | A |
| *Carex disticha* | 951 | -0.32 | NA | 77 | 0.25 | 0.02 | 1583 | 0.03 | NA | Moist to wet grasslands | Not Threatened | I |
| *Vicia sepium* | 246 | -0.32 | NA | 122 | -0.12 | NA | 3648 | -0.22 | <.001 | Scrubs, copses and field hedges | Not Threatened | I |
| *Carex leporina* | 490 | -0.32 | NA | 107 | -0.20 | NA | 982 | 0.17 | <.001 | Moist to wet grasslands | Not Threatened | I |
| *Prunella vulgaris* | 378 | -0.32 | NA | 108 | -0.15 | NA | 1226 | -0.20 | <.001 | Heaths, inland dunes and semi-natural grasslands | Not Threatened | I |
| *Gnaphalium uliginosum* | 84 | -0.32 | NA | 37 | -0.45 | 0.011 | 28 | -0.03 | NA | Standing waters | Not Threatened | I |
| *Circaea alpina* | 13 | -0.31 | NA | NA | NA | NA | 116 | 0.35 | <.001 | None | Not Threatened | I |
| *Pastinaca sativa* | 56 | -0.31 | NA | 12 | -0.26 | NA | 486 | -0.48 | <.001 | Coastal and marine habitats | Not Threatened | I |
| *Phleum pratense* agg. | 359 | -0.30 | NA | 263 | -0.11 | NA | 1545 | -0.39 | <.001 | Mesic grasslands | Not Threatened | I |
| *Daphne mezereum* | 3 | -0.30 | NA | NA | NA | NA | 1508 | -0.15 | <.001 | Dry to moderately moist forests | Not Threatened | I |
| *Epilobium montanum* | 124 | -0.30 | NA | 82 | -0.07 | NA | 914 | -0.40 | <.001 | Dry to moderately moist forests | Not Threatened | I |
| *Carlina vulgaris* agg. | 20 | -0.30 | NA | NA | NA | NA | 949 | -0.22 | <.001 | Heaths, inland dunes and semi-natural grasslands | Not Threatened | I |
| *Chrysosplenium alternifolium* | 519 | -0.29 | NA | 4 | -0.40 | NA | 1266 | 0.10 | <.001 | Moist to wet forests | Not Threatened | I |
| *Senecio sylvaticus* | 23 | -0.29 | NA | 43 | -0.82 | <.001 | 14 | -0.43 | NA | None | Not Threatened | I |
| *Lapsana communis* | 199 | -0.29 | NA | 64 | 0.21 | NA | 2211 | 0.20 | <.001 | Scrubs, copses and field hedges | Not Threatened | I |
| *Bellis perennis* | 502 | -0.29 | NA | 109 | -0.08 | NA | 398 | -0.35 | <.001 | Mesic grasslands | Not Threatened | A |
| *Hordelymus europaeus* | 130 | -0.28 | NA | NA | NA | NA | 1134 | 0.32 | <.001 | Dry to moderately moist forests | Not Threatened | I |
| *Luzula campestris* agg. | 978 | -0.28 | NA | 134 | 0.01 | NA | 1674 | 0.25 | <.001 | Heaths, inland dunes and semi-natural grasslands | Not Threatened | I |
| *Pimpinella saxifraga* agg. | 112 | -0.28 | NA | 2 | -0.67 | NA | 4114 | -0.16 | <.001 | Heaths, inland dunes and semi-natural grasslands | Not Threatened | I |
| *Viola odorata* | 8 | -0.28 | NA | 2 | -0.50 | NA | 124 | -0.55 | <.001 | None | Not Threatened | A |
| *Poa pratensis* agg. | 716 | -0.28 | NA | 250 | -0.18 | NA | 1888 | -0.27 | <.001 | Mesic grasslands | Not Threatened | I |
| *Juncus tenageia* | 2 | -0.27 | NA | 23 | -0.82 | <.001 | 2 | 1.00 | NA | None | Highly Endangered | I |
| *Berula erecta* | 1373 | -0.27 | NA | 41 | -0.04 | NA | 374 | 0.23 | <.001 | Linear and running surface waters | Not Threatened | I |
| *Osmunda regalis* | 93 | -0.27 | NA | 13 | -0.73 | 0.006 | 4 | 0.25 | NA | Moist to wet forests | Endangered | I |
| *Veronica arvensis* | 54 | -0.27 | NA | 27 | 0.38 | NA | 74 | -0.48 | <.001 | Mesic grasslands | Not Threatened | A |
| *Cerastium fontanum* agg. | 945 | -0.25 | NA | 291 | 0.10 | NA | 462 | -0.31 | <.001 | Mesic grasslands | Not Threatened | I |
| *Typha latifolia* | 2367 | -0.25 | NA | 236 | -0.04 | NA | 1736 | -0.10 | <.001 | Standing waters | Not Threatened | I |
| *Armoracia rusticana* | 8 | -0.25 | NA | 26 | -0.51 | 0.004 | 26 | -0.22 | NA | Ruderal, fringe and tall forb communities, clearings | Not Evaluated | N |
| *Hesperis matronalis* agg. | 2 | -0.25 | NA | 3 | 0.00 | NA | 132 | -0.20 | 0.01 | None | Not Evaluated | N |
| *Senecio viscosus* | 59 | -0.24 | NA | 33 | -0.70 | <.001 | 28 | -0.30 | NA | Coastal and marine habitats | Not Threatened | I |
| *Tripleurospermum maritimum* agg. | 130 | -0.24 | NA | 102 | -0.56 | <.001 | 46 | -0.71 | <.001 | Coastal and marine habitats | Not Threatened | NA |
| *Rhynchospora alba* | 220 | -0.24 | NA | 10 | -0.02 | NA | 72 | -0.62 | <.001 | Bogs, transition mires, marshes and fens | Endangered | I |
| *Anthriscus sylvestris* agg. | 672 | -0.24 | NA | 312 | -0.12 | NA | 1688 | -0.14 | <.001 | Scrubs, copses and field hedges | Not Threatened | I |
| *Plantago major* agg. | 372 | -0.24 | NA | 328 | -0.04 | NA | 246 | -0.40 | <.001 | Mesic grasslands | Not Threatened | I |
| *Oenanthe aquatica* agg. | 431 | -0.23 | NA | 68 | -0.37 | 0.001 | 22 | -0.38 | NA | Standing waters | Not Threatened | I |
| *Lysimachia nummularia* | 792 | -0.23 | NA | 150 | 0.14 | NA | 2292 | -0.13 | <.001 | Moist to wet grasslands | Not Threatened | I |
| *Cirsium palustre* | 2401 | -0.23 | NA | 280 | -0.10 | NA | 3916 | 0.16 | <.001 | Moist to wet grasslands | Not Threatened | I |
| *Angelica sylvestris* | 1663 | -0.22 | NA | 96 | -0.04 | NA | 6387 | 0.09 | <.001 | Moist to wet grasslands | Not Threatened | I |
| *Sonchus oleraceus* | 36 | -0.22 | NA | 42 | -0.49 | <.001 | 82 | -0.08 | NA | Farmland | Not Threatened | I |
| *Persicaria maculosa* | 101 | -0.22 | NA | 82 | -0.44 | <.001 | 56 | -0.25 | NA | Farmland | Not Threatened | I |
| *Ranunculus auricomus* agg. | 307 | -0.22 | NA | 6 | 1.00 | NA | 861 | -0.39 | <.001 | None | Not Threatened | I |
| *Centaurium erythraea* s. l. | 58 | -0.21 | NA | 8 | -0.10 | NA | 193 | -0.28 | <.001 | Heaths, inland dunes and semi-natural grasslands | Not Threatened | I |
| *Eupatorium cannabinum* | 1694 | -0.21 | NA | 82 | -0.16 | NA | 2056 | 0.12 | <.001 | Fresh water vegetation, springs and reeds | Not Threatened | I |
| *Veronica anagallis-aquatica* agg. | 164 | -0.21 | NA | 32 | 0.47 | 0.016 | 238 | 0.01 | NA | Linear and running surface waters | Not Threatened | I |
| *Rumex acetosella* s. l. | 866 | -0.21 | NA | 221 | -0.30 | <.001 | 965 | 0.12 | <.001 | Heaths, inland dunes and semi-natural grasslands | Not Threatened | I |
| *Vicia cracca* agg. | 764 | -0.20 | NA | 306 | -0.09 | NA | 2634 | -0.14 | <.001 | Mesic grasslands | Not Threatened | I |
| *Rubus* sect. *Corylifolii* | 7 | -0.19 | NA | 16 | 0.54 | NA | 475 | 0.40 | <.001 | Scrubs, copses and field hedges | Not Threatened | I |
| *Achillea millefolium* agg. | 1010 | -0.19 | NA | 346 | -0.24 | <.001 | 4407 | -0.16 | <.001 | Heaths, inland dunes and semi-natural grasslands | Not Threatened | I |
| *Nasturtium officinale* agg. | 323 | -0.19 | NA | 66 | 0.18 | NA | 571 | -0.18 | <.001 | Linear and running surface waters | Not Threatened | I |
| *Gymnocarpium dryopteris* | 26 | -0.19 | NA | 8 | 1.00 | NA | 128 | -0.22 | 0.007 | Dry to moderately moist forests | Not Threatened | I |
| *Poa annua* agg. | 201 | -0.19 | NA | 262 | -0.11 | NA | 126 | -0.67 | <.001 | Farmland | Not Threatened | I |
| *Odontites vernus* agg. | 122 | -0.19 | NA | 6 | -0.83 | NA | 164 | -0.35 | <.001 | Coastal and marine habitats | Not Threatened | I |
| *Sambucus racemosa* | 71 | -0.19 | NA | 108 | -0.29 | 0.001 | 2203 | -0.13 | <.001 | Scrubs, copses and field hedges | Not Threatened | I |
| *Geranium pratense* | 14 | -0.18 | NA | 8 | -0.02 | NA | 2230 | -0.16 | <.001 | Scrubs, copses and field hedges | Not Threatened | I |
| *Persicaria hydropiper* | 650 | -0.18 | NA | 226 | 0.02 | NA | 226 | 0.41 | <.001 | Linear and running surface waters | Not Threatened | I |
| *Stachys sylvatica* | 2002 | -0.18 | NA | 72 | 0.14 | NA | 4972 | 0.10 | <.001 | Dry to moderately moist forests | Not Threatened | I |
| *Salix pentandra* | 406 | -0.17 | NA | 93 | -0.44 | <.001 | 146 | -0.16 | NA | Moist to wet forests | Not Threatened | I |
| *Epilobium tetragonum* s. l. | 192 | -0.17 | NA | 26 | 0.32 | NA | 216 | -0.17 | 0.01 | Moist to wet grasslands | Not Threatened | I |
| *Lamium purpureum* s. str. | 37 | -0.17 | NA | 26 | -0.45 | NA | 333 | -0.29 | <.001 | None | Not Threatened | A |
| *Nymphaea alba* | 477 | -0.17 | NA | 56 | 0.31 | 0.011 | 322 | -0.03 | NA | Standing waters | Not Threatened | I |
| *Ulmus glabra* | 690 | -0.16 | NA | 62 | -0.02 | NA | 2963 | 0.12 | <.001 | Dry to moderately moist forests | Not Threatened | I |
| *Viola tricolor* agg. | 126 | -0.16 | NA | 44 | -0.51 | NA | 54 | -0.49 | <.001 | Coastal and marine habitats | Not Threatened | NA |
| *Anthoxanthum odoratum* agg. | 1608 | -0.16 | NA | 192 | 0.07 | NA | 2466 | 0.13 | <.001 | Moist to wet grasslands | Not Threatened | I |
| *Lonicera xylosteum* | 189 | -0.16 | NA | 46 | 0.69 | <.001 | 11186 | 0.06 | <.001 | Scrubs, copses and field hedges | Not Threatened | I |
| *Maianthemum bifolium* | 1324 | -0.16 | NA | 98 | -0.33 | <.001 | 390 | -0.15 | NA | Dry to moderately moist forests | Not Threatened | I |
| *Sambucus nigra* | 3310 | -0.15 | NA | 740 | 0.03 | NA | 22877 | 0.04 | <.001 | Scrubs, copses and field hedges | Not Threatened | I |
| *Equisetum telmateia* | 155 | -0.15 | NA | NA | NA | NA | 1108 | 0.11 | <.001 | Moist to wet forests | Not Threatened | I |
| *Galium verum* agg. | 172 | -0.15 | NA | 10 | 0.15 | NA | 4762 | -0.09 | <.001 | Heaths, inland dunes and semi-natural grasslands | Not Threatened | I |
| *Vicia sylvatica* | 12 | -0.15 | NA | 1 | -1.00 | NA | 77 | -0.75 | <.001 | Dry to moderately moist forests | Not Threatened | I |
| *Sonchus arvensis* agg. | 290 | -0.15 | NA | 30 | -0.28 | NA | 48 | -0.38 | 0.01 | Coastal and marine habitats | Not Threatened | I |
| *Equisetum hyemale* | 115 | -0.15 | NA | 1 | 0.00 | NA | 258 | 0.18 | <.001 | Moist to wet forests | Not Threatened | I |
| *Solanum nigrum* s. l. | 34 | -0.15 | NA | 23 | -0.71 | <.001 | 60 | 0.57 | <.001 | None | Not Threatened | A |
| *Persicaria amphibia* | 1272 | -0.15 | NA | 250 | 0.17 | 0.004 | 493 | 0.01 | NA | Standing waters | Not Threatened | I |
| *Oxalis acetosella* | 2212 | -0.15 | NA | 144 | 0.30 | <.001 | 2762 | -0.04 | NA | Dry to moderately moist forests | Not Threatened | I |
| *Bromus tectorum* | 16 | -0.14 | NA | 8 | -0.33 | NA | 68 | 0.38 | 0.002 | None | Not Threatened | A |
| *Vicia hirsuta* | 98 | -0.13 | NA | 65 | -0.26 | NA | 148 | -0.39 | <.001 | Mesic grasslands | Not Threatened | I |
| *Scrophularia nodosa* | 768 | -0.13 | NA | 222 | -0.01 | NA | 1472 | -0.22 | <.001 | Dry to moderately moist forests | Not Threatened | I |
| *Malva alcea* | 4 | -0.13 | NA | 1 | 1.00 | NA | 130 | -0.27 | <.001 | None | Not Threatened | A |
| *Fragaria viridis* | 5 | -0.13 | NA | NA | NA | NA | 602 | 0.32 | <.001 | Heaths, inland dunes and semi-natural grasslands | Not Threatened | I |
| *Artemisia vulgaris* agg. | 286 | -0.13 | NA | 379 | -0.26 | <.001 | 972 | -0.23 | <.001 | Ruderal, fringe and tall forb communities, clearings | Not Threatened | NA |
| *Veronica serpyllifolia* | 136 | -0.12 | NA | 24 | 0.59 | 0.023 | 56 | -0.45 | <.001 | Mesic grasslands | Not Threatened | I |
| *Najas marina* s. l. | 9 | -0.12 | NA | NA | NA | NA | 19 | 0.60 | 0.003 | None | Not Threatened | I |
| *Veronica beccabunga* | 1168 | -0.12 | NA | 76 | 0.20 | NA | 2502 | 0.10 | <.001 | Moist to wet forests | Not Threatened | I |
| *Danthonia decumbens* | 150 | -0.12 | NA | 26 | 0.33 | NA | 937 | 0.28 | <.001 | Heaths, inland dunes and semi-natural grasslands | Near Threatened | I |
| *Carex pallescens* | 54 | -0.12 | NA | 8 | -0.05 | NA | 922 | 0.21 | <.001 | Moist to wet grasslands | Not Threatened | I |
| *Abies alba* | 62 | -0.12 | NA | 6 | 0.63 | NA | 3006 | 0.14 | <.001 | Dry to moderately moist forests | Not Threatened | I |
| *Tanacetum vulgare* | 371 | -0.12 | NA | 245 | -0.29 | <.001 | 252 | -0.05 | NA | Ruderal, fringe and tall forb communities, clearings | Not Threatened | A |
| *Veronica chamaedrys* agg. | 404 | -0.12 | NA | 124 | -0.04 | NA | 1398 | -0.22 | <.001 | Mesic grasslands | Not Threatened | I |
| *Festuca rubra* agg. | 2066 | -0.10 | NA | 405 | 0.13 | 0.002 | 3200 | 0.23 | <.001 | Coastal and marine habitats | Not Threatened | I |
| *Solidago gigantea* | 52 | -0.10 | NA | 191 | 0.09 | NA | 2103 | 0.37 | <.001 | Fresh water vegetation, springs and reeds | Not Evaluated | N |
| *Galium odoratum* | 1836 | -0.10 | NA | 24 | 0.32 | NA | 6090 | 0.13 | <.001 | Dry to moderately moist forests | Not Threatened | I |
| *Lactuca serriola* | 18 | -0.09 | NA | 10 | 0.03 | NA | 688 | 0.24 | <.001 | Scrubs, copses and field hedges | Not Threatened | I |
| *Galium mollugo* agg. | 426 | -0.09 | NA | 114 | -0.06 | NA | 7666 | -0.11 | <.001 | Scrubs, copses and field hedges | Not Threatened | I |
| *Taraxacum* sect. *Ruderalia* | 1063 | -0.09 | NA | 430 | 0.09 | NA | 1837 | -0.25 | <.001 | Mesic grasslands | Not Threatened | I |
| *Agrostis canina* agg. | 1095 | -0.09 | NA | 99 | 0.25 | 0.005 | 1076 | 0.13 | <.001 | Moist to wet grasslands | Near Threatened | I |
| *Hippophae rhamnoides* | 98 | -0.08 | NA | 34 | -0.16 | NA | 150 | -0.46 | <.001 | Coastal and marine habitats | Not Threatened | I |
| *Stellaria media* agg. | 342 | -0.08 | NA | 258 | -0.28 | <.001 | 325 | -0.34 | <.001 | Farmland | Not Threatened | NA |
| *Linaria vulgaris* agg. | 271 | -0.08 | NA | 107 | -0.03 | NA | 1532 | -0.14 | <.001 | Scrubs, copses and field hedges | Not Threatened | I |
| *Polypodium vulgare* agg. | 186 | -0.08 | NA | 6 | 1.00 | 0.031 | 936 | 0.05 | NA | Dry to moderately moist forests | Not Threatened | I |
| *Oenothera* | 68 | -0.08 | NA | 72 | -0.37 | <.001 | 108 | 0.08 | NA | Ruderal, fringe and tall forb communities, clearings | Not Evaluated | N |
| *Trifolium campestre* | 138 | -0.08 | NA | 21 | -0.30 | NA | 630 | -0.26 | <.001 | Heaths, inland dunes and semi-natural grasslands | Not Threatened | I |
| *Fraxinus excelsior* | 3544 | -0.08 | NA | 444 | 0.17 | NA | 22786 | 0.13 | <.001 | Scrubs, copses and field hedges | Not Threatened | I |
| *Alnus glutinosa* | 5080 | -0.08 | NA | 583 | 0.03 | NA | 9470 | 0.08 | <.001 | Moist to wet forests | Not Threatened | I |
| *Carex acuta* agg. | 1598 | -0.07 | NA | 366 | 0.21 | <.001 | 2286 | 0.09 | NA | Moist to wet grasslands | Not Threatened | I |
| *Adoxa moschatellina* | 210 | -0.07 | NA | 2 | 0.00 | NA | 208 | -0.47 | <.001 | Dry to moderately moist forests | Not Threatened | I |
| *Epilobium angustifolium* | 370 | -0.07 | NA | 272 | -0.42 | <.001 | 1104 | -0.15 | <.001 | Coastal and marine habitats | Not Threatened | I |
| *Quercus robur* | 4932 | -0.07 | NA | 710 | 0.12 | NA | 16573 | 0.09 | <.001 | Scrubs, copses and field hedges | Not Threatened | I |
| *Capsella bursa-pastoris* | 58 | -0.06 | NA | 90 | -0.25 | NA | 85 | -0.36 | <.001 | Mesic grasslands | Not Threatened | I |
| *Salix cinerea_Salix multinervis* | 4214 | -0.06 | NA | 380 | 0.28 | <.001 | 5122 | 0.12 | <.001 | Standing waters | Not Threatened | I |
| *Bromus sterilis* | 22 | -0.06 | NA | 17 | 0.18 | NA | 1936 | 0.47 | <.001 | Scrubs, copses and field hedges | Not Threatened | A |
| *Salix alba* | 1119 | -0.06 | NA | 366 | 0.25 | <.001 | 3787 | 0.01 | NA | Moist to wet forests | Not Threatened | I |
| *Gagea lutea* | 73 | -0.05 | NA | 2 | 1.00 | NA | 82 | -0.45 | <.001 | Dry to moderately moist forests | Not Threatened | I |
| *Cardamine amara* | 1898 | -0.05 | NA | 38 | 0.01 | NA | 1832 | 0.19 | <.001 | Moist to wet forests | Not Threatened | I |
| *Potentilla verna* agg. | 2 | -0.05 | NA | NA | NA | NA | 2544 | -0.09 | <.001 | Heaths, inland dunes and semi-natural grasslands | Near Threatened | I |
| *Alopecurus pratensis* agg. | 1008 | -0.05 | NA | 326 | -0.00 | NA | 2845 | 0.12 | <.001 | Moist to wet grasslands | Not Threatened | I |
| *Setaria viridis* | 7 | -0.04 | NA | 4 | -0.21 | NA | 60 | 0.52 | <.001 | None | Not Threatened | A |
| *Iris pseudacorus* | 3960 | -0.04 | NA | 418 | 0.15 | NA | 2523 | 0.12 | <.001 | Standing waters | Not Threatened | I |
| *Trientalis europaea* | 650 | -0.04 | NA | 136 | -0.35 | <.001 | 18 | 0.24 | NA | Dry to moderately moist forests | Not Threatened | I |
| *Geranium pusillum* | 32 | -0.04 | NA | 12 | 0.31 | NA | 58 | -0.36 | 0.006 | Mesic grasslands | Not Threatened | A |
| *Potentilla anserina* | 1218 | -0.04 | NA | 360 | 0.10 | NA | 585 | -0.22 | <.001 | Coastal and marine habitats | Not Threatened | I |
| *Phragmites australis* | 3650 | -0.03 | NA | 504 | 0.10 | NA | 4628 | 0.07 | <.001 | Fresh water vegetation, springs and reeds | Not Threatened | I |
| *Solidago canadensis* | 90 | -0.03 | NA | 174 | -0.32 | <.001 | 1306 | -0.05 | NA | Scrubs, copses and field hedges | Not Evaluated | N |
| *Melica uniflora* | 2075 | -0.03 | NA | 12 | -0.04 | NA | 664 | 0.25 | <.001 | Dry to moderately moist forests | Not Threatened | I |
| *Thlaspi arvense* | 12 | -0.03 | NA | 3 | 1.00 | NA | 32 | -0.69 | <.001 | None | Not Threatened | A |
| *Rosa rubiginosa* agg. | 26 | -0.03 | NA | 13 | 1.00 | <.001 | 862 | -0.06 | NA | Scrubs, copses and field hedges | Not Evaluated | I |
| *Fagus sylvatica* | 2976 | -0.02 | NA | 356 | 0.12 | NA | 11540 | 0.07 | <.001 | Dry to moderately moist forests | Not Threatened | I |
| *Corylus avellana* | 3294 | -0.02 | NA | 328 | 0.26 | NA | 21638 | 0.11 | <.001 | Scrubs, copses and field hedges | Not Threatened | I |
| *Carex sylvatica* | 1440 | -0.02 | NA | 20 | 0.19 | NA | 3730 | 0.28 | <.001 | Dry to moderately moist forests | Not Threatened | I |
| *Bromus inermis* | 40 | -0.02 | NA | 27 | -0.16 | NA | 710 | 0.23 | <.001 | Scrubs, copses and field hedges | Not Threatened | I |
| *Deschampsia cespitosa* agg. | 4332 | -0.01 | NA | 471 | 0.04 | NA | 6425 | 0.07 | <.001 | Moist to wet forests | Not Threatened | I |
| *Anemone nemorosa* | 2328 | -0.01 | NA | 50 | 0.14 | NA | 3798 | -0.14 | <.001 | Dry to moderately moist forests | Not Threatened | I |
| *Plantago lanceolata* | 1214 | -0.01 | NA | 382 | 0.04 | NA | 2982 | -0.15 | <.001 | Mesic grasslands | Not Threatened | A |
| *Rosa canina* agg. | 693 | -0.01 | NA | 277 | 0.20 | <.001 | 17586 | -0.03 | NA | Scrubs, copses and field hedges | Not Threatened | I |
| *Chrysosplenium oppositifolium* | 774 | -0.01 | NA | 7 | 0.33 | NA | 1250 | 0.20 | <.001 | Moist to wet forests | Not Threatened | I |
| *Lathraea squamaria* | 36 | -0.00 | NA | 2 | -0.88 | NA | 34 | -0.54 | 0.003 | None | Not Threatened | I |
| *Carpinus betulus* | 2266 | -0.00 | NA | 292 | 0.11 | NA | 10810 | 0.09 | <.001 | Dry to moderately moist forests | Not Threatened | I |
| *Milium effusum* | 2731 | -0.00 | NA | 104 | 0.30 | <.001 | 4020 | -0.00 | NA | Dry to moderately moist forests | Not Threatened | I |
| *Pulsatilla vulgaris* s. l. | 2 | 0.00 | NA | NA | NA | NA | 672 | -0.27 | <.001 | Heaths, inland dunes and semi-natural grasslands | Endangered | I |
| *Trisetum flavescens* | 8 | 0.01 | NA | NA | NA | NA | 1372 | -0.29 | <.001 | Heaths, inland dunes and semi-natural grasslands | Not Threatened | I |
| *Clematis vitalba* | 22 | 0.01 | NA | 20 | 0.41 | NA | 7872 | 0.10 | <.001 | Scrubs, copses and field hedges | Not Threatened | I |
| *Rumex maritimus* | 94 | 0.01 | NA | 28 | -0.44 | 0.023 | 7 | 0.67 | NA | Coastal and marine habitats | Not Threatened | I |
| *Cornus sanguinea* | 168 | 0.01 | NA | 94 | 0.36 | <.001 | 19774 | 0.16 | <.001 | Scrubs, copses and field hedges | Not Threatened | I |
| *Agrostis capillaris* | 1528 | 0.02 | NA | 556 | -0.10 | NA | 2474 | 0.21 | <.001 | Mesic grasslands | Not Threatened | I |
| *Hieracium lachenalii* | 46 | 0.02 | NA | 20 | -0.41 | NA | 164 | 0.23 | 0.003 | Heaths, inland dunes and semi-natural grasslands | Not Threatened | I |
| *Cirsium arvense* | 1800 | 0.02 | NA | 741 | -0.20 | <.001 | 3304 | -0.02 | NA | Mesic grasslands | Not Threatened | I |
| *Agrimonia eupatoria* | 130 | 0.02 | NA | 25 | 0.49 | 0.012 | 5536 | 0.03 | NA | Heaths, inland dunes and semi-natural grasslands | Not Threatened | I |
| *Corydalis cava* | 98 | 0.02 | NA | 1 | 1.00 | NA | 252 | -0.27 | <.001 | Dry to moderately moist forests | Not Threatened | I |
| *Lupinus* | 50 | 0.04 | NA | 26 | -0.27 | NA | 114 | 0.34 | <.001 | Ruderal, fringe and tall forb communities, clearings | Not Evaluated | N |
| *Galium palustre* agg. | 3112 | 0.04 | NA | 248 | 0.18 | 0.002 | 2848 | 0.12 | <.001 | Moist to wet grasslands | Not Threatened | I |
| *Athyrium filix-femina* | 1942 | 0.04 | NA | 189 | 0.28 | <.001 | 4008 | 0.14 | <.001 | Moist to wet forests | Not Threatened | I |
| *Thelypteris limbosperma* | 2 | 0.04 | NA | NA | NA | NA | 153 | 0.24 | 0.003 | None | Not Threatened | I |
| *Lysimachia nemorum* | 275 | 0.04 | NA | 14 | -0.63 | NA | 833 | 0.19 | <.001 | Moist to wet grasslands | Not Threatened | I |
| *Festuca arundinacea* | 371 | 0.04 | NA | 126 | -0.15 | NA | 378 | 0.25 | <.001 | Coastal and marine habitats | Not Threatened | I |
| *Carex acutiformis* | 3831 | 0.04 | NA | 170 | 0.34 | <.001 | 6115 | 0.13 | <.001 | Fresh water vegetation, springs and reeds | Not Threatened | I |
| *Carex pseudocyperus* | 1104 | 0.04 | NA | 116 | -0.02 | NA | 102 | 0.31 | 0.001 | Standing waters | Not Threatened | I |
| *Valeriana officinalis* agg. | 1428 | 0.04 | NA | 138 | 0.13 | NA | 6130 | 0.10 | <.001 | Fresh water vegetation, springs and reeds | Not Threatened | I |
| *Ilex aquifolium* | 1314 | 0.05 | NA | 138 | 0.72 | <.001 | 1040 | 0.26 | <.001 | Dry to moderately moist forests | Not Threatened | I |
| *Callitriche* | 770 | 0.05 | NA | 181 | 0.02 | NA | 646 | 0.12 | <.001 | Linear and running surface waters | Highly Endangered | I |
| *Lamium album* | 55 | 0.05 | NA | 146 | -0.22 | NA | 1522 | -0.23 | <.001 | Scrubs, copses and field hedges | Not Threatened | A |
| *Medicago lupulina* | 125 | 0.05 | NA | 96 | -0.26 | NA | 1096 | -0.15 | <.001 | Heaths, inland dunes and semi-natural grasslands | Not Threatened | I |
| *Hypochaeris radicata* | 722 | 0.06 | NA | 125 | 0.11 | NA | 444 | 0.35 | <.001 | Mesic grasslands | Not Threatened | I |
| *Prunus avium* | 1236 | 0.06 | NA | 132 | 0.41 | <.001 | 16675 | 0.12 | <.001 | Scrubs, copses and field hedges | Not Threatened | I |
| *Aegopodium podagraria* | 1926 | 0.07 | NA | 470 | 0.16 | NA | 10437 | 0.11 | <.001 | Scrubs, copses and field hedges | Not Threatened | I |
| *Holcus mollis* | 902 | 0.08 | NA | 288 | -0.24 | <.001 | 872 | 0.12 | NA | Dry to moderately moist forests | Not Threatened | I |
| *Poa compressa* | 33 | 0.08 | NA | 14 | -0.34 | NA | 236 | -0.20 | <.001 | Heaths, inland dunes and semi-natural grasslands | Not Threatened | I |
| *Dryopteris filix-mas* agg. | 1965 | 0.08 | NA | 344 | 0.32 | <.001 | 7152 | 0.21 | <.001 | Dry to moderately moist forests | Not Threatened | I |
| *Phalaris arundinacea* | 4222 | 0.09 | NA | 562 | 0.10 | NA | 4730 | 0.11 | <.001 | Fresh water vegetation, springs and reeds | Not Threatened | I |
| *Circaea lutetiana* | 2698 | 0.10 | NA | 120 | 0.64 | <.001 | 3252 | 0.26 | <.001 | Dry to moderately moist forests | Not Threatened | I |
| *Acer pseudoplatanus* | 2928 | 0.10 | NA | 472 | 0.29 | <.001 | 16524 | 0.20 | <.001 | Dry to moderately moist forests | Not Threatened | I |
| *Stellaria holostea* | 3746 | 0.11 | NA | 120 | 0.31 | <.001 | 2116 | -0.10 | <.001 | Dry to moderately moist forests | Not Threatened | I |
| *Ceratophyllum demersum* | 196 | 0.11 | NA | 52 | 0.43 | 0.002 | 158 | 0.16 | NA | Standing waters | Not Threatened | I |
| *Myosotis arvensis* | 32 | 0.11 | NA | 52 | -0.47 | <.001 | 203 | -0.65 | <.001 | Farmland | Not Threatened | A |
| *Euonymus europaeus* | 978 | 0.11 | NA | 70 | 0.37 | NA | 15076 | 0.12 | <.001 | Scrubs, copses and field hedges | Not Threatened | I |
| *Papaver rhoeas* | 4 | 0.12 | NA | 8 | -0.47 | NA | 176 | -0.43 | <.001 | None | Not Threatened | A |
| *Acer campestre* | 572 | 0.12 | NA | 190 | 0.32 | <.001 | 15178 | 0.17 | <.001 | Scrubs, copses and field hedges | Not Threatened | I |
| *Erigeron canadensis* | 84 | 0.12 | NA | 85 | -0.22 | NA | 158 | 0.38 | <.001 | Ruderal, fringe and tall forb communities, clearings | Not Evaluated | N |
| *Galeopsis tetrahit* agg. | 1298 | 0.13 | NA | 380 | -0.05 | NA | 5552 | -0.10 | <.001 | Scrubs, copses and field hedges | Not Threatened | I |
| *Rorippa sylvestris* | 34 | 0.13 | NA | 57 | -0.51 | <.001 | 15 | -0.23 | NA | None | Not Threatened | I |
| *Pseudotsuga menziesii* | 186 | 0.13 | NA | 68 | 0.01 | NA | 861 | 0.23 | <.001 | Dry to moderately moist forests | Not Evaluated | N |
| *Brachypodium sylvaticum* | 1228 | 0.14 | NA | 13 | 0.48 | NA | 5442 | 0.40 | <.001 | Dry to moderately moist forests | Not Threatened | I |
| *Chaerophyllum temulum* | 162 | 0.14 | NA | 45 | 0.52 | <.001 | 352 | -0.12 | NA | Scrubs, copses and field hedges | Not Threatened | I |
| *Anemone ranunculoides* | 109 | 0.15 | NA | 1 | -1.00 | NA | 224 | -0.52 | <.001 | Dry to moderately moist forests | Not Threatened | I |
| *Hylotelephium telephium* agg. | 66 | 0.15 | NA | 9 | -0.37 | NA | 1048 | -0.19 | <.001 | Scrubs, copses and field hedges | Not Threatened | I |
| *Dianthus carthusianorum* agg. | 8 | 0.15 | NA | 2 | 0.33 | NA | 2140 | -0.14 | <.001 | Heaths, inland dunes and semi-natural grasslands | Near Threatened | I |
| *Impatiens noli-tangere* | 1702 | 0.15 | NA | 54 | 0.08 | NA | 3682 | 0.08 | <.001 | Moist to wet forests | Not Threatened | I |
| *Carex pilulifera* | 746 | 0.15 | NA | 102 | 0.38 | <.001 | 890 | 0.33 | <.001 | Heaths, inland dunes and semi-natural grasslands | Not Threatened | I |
| *Geranium molle* agg. | 95 | 0.15 | NA | 24 | -0.16 | NA | 80 | 0.28 | 0.014 | Mesic grasslands | Not Threatened | A |
| *Allium vineale* s. l. | 22 | 0.18 | NA | 2 | 0.60 | NA | 275 | -0.21 | <.001 | None | Not Threatened | I |
| *Spergularia rubra* | 16 | 0.18 | NA | 14 | -0.48 | NA | 18 | -0.87 | <.001 | None | Not Threatened | A |
| *Carex elongata* | 1148 | 0.18 | NA | 42 | 0.26 | NA | 326 | 0.19 | <.001 | Moist to wet forests | Not Threatened | I |
| *Consolida regalis* | 10 | 0.18 | NA | NA | NA | NA | 54 | -0.43 | 0.003 | None | Endangered | A |
| *Cardamine flexuosa* | 370 | 0.18 | NA | 18 | 0.90 | <.001 | 282 | 0.28 | <.001 | Dry to moderately moist forests | Not Threatened | I |
| *Ficaria verna* s. l. | 2319 | 0.19 | NA | 72 | 0.53 | <.001 | 2699 | -0.28 | <.001 | Dry to moderately moist forests | Not Threatened | I |
| *Rumex sanguineus* | 550 | 0.19 | NA | 54 | 0.89 | <.001 | 162 | -0.07 | NA | Moist to wet forests | Not Threatened | I |
| *Salicornia europaea* agg. | 230 | 0.19 | NA | 12 | -0.92 | <.001 | NA | NA | NA | Coastal and marine habitats | Not Threatened | I |
| *Hedera helix* | 2232 | 0.20 | NA | 219 | 0.58 | <.001 | 9730 | 0.33 | <.001 | Dry to moderately moist forests | Not Threatened | I |
| *Lemna minor* | 3194 | 0.20 | NA | 287 | 0.06 | NA | 1549 | 0.10 | <.001 | Standing waters | Not Threatened | I |
| *Geum urbanum* | 2633 | 0.20 | NA | 348 | 0.59 | <.001 | 15057 | 0.14 | <.001 | Scrubs, copses and field hedges | Not Threatened | I |
| *Prunus padus* | 1417 | 0.20 | NA | 208 | 0.46 | <.001 | 4686 | 0.24 | <.001 | Moist to wet forests | Not Threatened | I |
| *Convolvulus arvensis* | 139 | 0.20 | NA | 50 | -0.16 | NA | 1052 | -0.25 | <.001 | Scrubs, copses and field hedges | Not Threatened | I |
| *Equisetum arvense* | 586 | 0.21 | NA | 205 | -0.17 | NA | 1422 | -0.24 | <.001 | Ruderal, fringe and tall forb communities, clearings | Not Threatened | I |
| *Urtica dioica* s. l. | 7535 | 0.21 | NA | 1270 | -0.01 | NA | 28635 | 0.06 | <.001 | Scrubs, copses and field hedges | Not Threatened | I |
| *Robinia pseudoacacia* | 70 | 0.23 | NA | 130 | 0.02 | NA | 3430 | 0.12 | <.001 | Scrubs, copses and field hedges | Not Evaluated | N |
| *Poa trivialis* s. l. | 3006 | 0.23 | NA | 578 | 0.18 | <.001 | 1723 | -0.08 | NA | Moist to wet grasslands | Not Threatened | I |
| *Digitalis purpurea* | 86 | 0.23 | NA | 117 | -0.33 | NA | 764 | 0.13 | <.001 | Dry to moderately moist forests | Not Threatened | I |
| *Carex muricata* agg. | 112 | 0.25 | NA | 31 | 0.56 | <.001 | 634 | -0.04 | NA | None | Not Threatened | I |
| *Acer platanoides* | 333 | 0.25 | NA | 233 | 0.38 | <.001 | 4930 | 0.26 | <.001 | Scrubs, copses and field hedges | Not Threatened | I |
| *Dryopteris carthusiana* agg. | 4632 | 0.25 | NA | 382 | 0.30 | <.001 | 3582 | 0.11 | <.001 | Dry to moderately moist forests | Not Threatened | I |
| *Lemna trisulca* | 496 | 0.27 | NA | 28 | 0.56 | 0.002 | 128 | 0.19 | NA | Standing waters | Not Threatened | I |
| *Elodea nuttallii* | 82 | 0.27 | NA | 22 | 0.91 | <.001 | 38 | 0.67 | <.001 | Linear and running surface waters | Not Evaluated | N |
| *Carex brizoides* | 24 | 0.31 | NA | 2 | 0.04 | NA | 2887 | 0.23 | <.001 | Moist to wet grasslands | Not Threatened | I |
| *Arctium lappa* | 102 | 0.32 | NA | 61 | 0.19 | NA | 396 | 0.33 | <.001 | None | Not Threatened | A |
| *Epilobium roseum* | 66 | 0.33 | NA | 26 | -0.68 | <.001 | 93 | -0.38 | <.001 | None | Not Threatened | I |
| *Geranium robertianum* agg. | 2120 | 0.33 | NA | 90 | 0.65 | <.001 | 11282 | 0.07 | <.001 | Scrubs, copses and field hedges | Not Threatened | NA |
| *Prunus mahaleb* | 2 | 0.33 | NA | 2 | 1.00 | NA | 180 | 0.34 | <.001 | Scrubs, copses and field hedges | Not Threatened | I |
| *Calystegia sepium* agg. | 1420 | 0.34 | NA | 465 | 0.16 | <.001 | 4864 | 0.06 | NA | Fresh water vegetation, springs and reeds | Not Threatened | NA |
| *Carex remota* | 2979 | 0.34 | NA | 104 | 0.43 | <.001 | 1942 | 0.28 | <.001 | Standing waters | Not Threatened | I |
| *Glechoma hederacea* agg. | 4288 | 0.34 | <.001 | 782 | 0.26 | <.001 | 7063 | 0.05 | NA | Moist to wet forests | Not Threatened | I |
| *Spirodela polyrhiza* | 784 | 0.36 | NA | 64 | 0.28 | NA | 269 | 0.39 | <.001 | Standing waters | Not Threatened | I |
| *Carex strigosa* | 324 | 0.37 | NA | NA | NA | NA | 84 | 0.38 | <.001 | Dry to moderately moist forests | Not Threatened | I |
| *Rubus* sect. *Rubus* | 5653 | 0.37 | <.001 | 796 | 0.17 | <.001 | 17402 | 0.16 | <.001 | Scrubs, copses and field hedges | Not Threatened | NA |
| *Dactylis glomerata* agg. | 2290 | 0.39 | <.001 | 836 | 0.03 | NA | 13061 | 0.09 | <.001 | Scrubs, copses and field hedges | Not Threatened | I |
| *Carex pendula* | 34 | 0.41 | NA | 10 | 1.00 | 0.002 | 1445 | 0.30 | <.001 | Moist to wet forests | Not Threatened | I |
| *Lemna gibba* | 68 | 0.44 | <.001 | 16 | -0.45 | NA | 8 | -0.48 | NA | Linear and running surface waters | Not Threatened | I |
| *Alliaria petiolata* | 1180 | 0.45 | <.001 | 250 | 0.53 | <.001 | 9350 | 0.24 | <.001 | Scrubs, copses and field hedges | Not Threatened | I |
| *Ribes rubrum* agg. | 1397 | 0.46 | <.001 | 98 | 0.47 | <.001 | 1218 | 0.09 | NA | Moist to wet forests | Not Threatened | I |
| *Salix smithiana* | 42 | 0.48 | 0.008 | 44 | 0.15 | NA | 238 | -0.34 | <.001 | None | Not Evaluated | I |
| *Polygonatum odoratum* | 82 | 0.49 | <.001 | 2 | -0.20 | NA | 542 | -0.08 | NA | Dry to moderately moist forests | Near Threatened | I |
| *Helianthus annuus* | 2 | 0.50 | NA | 10 | -0.90 | 0.008 | 9 | -0.11 | NA | Ruderal, fringe and tall forb communities, clearings | Not Evaluated | N |
| *Aethusa cynapium* | 5 | 0.50 | NA | 3 | -0.50 | NA | 92 | -0.42 | <.001 | None | Not Threatened | A |
| *Senecio jacobaea* | 386 | 0.51 | <.001 | 43 | 0.84 | <.001 | 870 | -0.05 | NA | Mesic grasslands | Not Threatened | I |
| *Castanea sativa* | 13 | 0.51 | NA | 10 | 0.10 | NA | 1085 | 0.12 | <.001 | Scrubs, copses and field hedges | Not Threatened | A |
| *Circaea intermedia* | 199 | 0.51 | <.001 | 1 | 1.00 | NA | 76 | 0.48 | <.001 | Dry to moderately moist forests | Not Threatened | I |
| *Taxus baccata* | 116 | 0.51 | NA | 100 | 0.70 | <.001 | 360 | 0.46 | <.001 | Anthropogenic | Near Threatened | I |
| *Juglans regia* | 21 | 0.52 | NA | 32 | 0.66 | <.001 | 6497 | 0.33 | <.001 | Scrubs, copses and field hedges | Not Threatened | A |
| *Quercus rubra* | 208 | 0.52 | <.001 | 105 | 0.12 | NA | 461 | 0.20 | <.001 | Dry to moderately moist forests | Not Evaluated | N |
| *Ribes uva-crispa* | 688 | 0.52 | <.001 | 48 | 0.17 | NA | 4266 | -0.09 | <.001 | Scrubs, copses and field hedges | Not Threatened | I |
| *Vicia tetrasperma* agg. | 42 | 0.53 | 0.008 | 51 | 0.36 | NA | 78 | -0.48 | <.001 | Mesic grasslands | Not Threatened | I |
| *Potamogeton polygonifolius* | 42 | 0.53 | 0.002 | 2 | 0.25 | NA | 2 | 1.00 | NA | Standing waters | Endangered | I |
| *Quercus petraea* agg. | 318 | 0.54 | <.001 | 148 | -0.11 | NA | 4276 | -0.05 | NA | Dry to moderately moist forests | Not Threatened | I |
| *Ribes nigrum* | 920 | 0.55 | <.001 | 29 | 0.29 | NA | 186 | 0.09 | NA | Moist to wet forests | Not Threatened | I |
| *Senecio erucifolius* | 8 | 0.56 | NA | 12 | -0.95 | <.001 | 1106 | -0.30 | <.001 | Heaths, inland dunes and semi-natural grasslands | Not Threatened | I |
| *Ligustrum vulgare* | 12 | 0.57 | NA | 90 | 0.15 | NA | 13798 | 0.14 | <.001 | Scrubs, copses and field hedges | Not Threatened | I |
| *Elymus athericus* | 125 | 0.59 | <.001 | NA | NA | NA | NA | NA | NA | Coastal and marine habitats | Not Threatened | I |
| *Sorbus intermedia* | 58 | 0.60 | <.001 | 22 | -0.01 | NA | 138 | -0.00 | NA | None | Vulnerable | N |
| *Impatiens parviflora* | 1948 | 0.61 | <.001 | 483 | 0.27 | <.001 | 1208 | 0.14 | <.001 | Dry to moderately moist forests | Not Evaluated | N |
| *Galeobdolon luteum* agg. | 3294 | 0.63 | <.001 | 220 | 0.52 | <.001 | 5666 | 0.09 | <.001 | Dry to moderately moist forests | Not Threatened | NA |
| *Rosa multiflora* | 11 | 0.63 | NA | 36 | 0.46 | 0.002 | 326 | 0.35 | <.001 | Scrubs, copses and field hedges | Not Evaluated | N |
| *Veronica hederifolia* agg. | 196 | 0.64 | <.001 | 14 | 0.23 | NA | 209 | -0.44 | <.001 | Dry to moderately moist forests | Not Threatened | I |
| *Festuca brevipila* | 28 | 0.65 | 0.002 | 5 | -0.46 | NA | 2 | -0.50 | NA | None | Not Threatened | I |
| *Salix fragilis* agg. | 1150 | 0.66 | <.001 | 208 | 0.06 | NA | 5566 | 0.12 | <.001 | Moist to wet forests | Not Threatened | I |
| *Fallopia bohemica_Fallopia japonica_Fallopia sachalinensis* | 138 | 0.66 | <.001 | 208 | 0.17 | NA | 439 | 0.44 | <.001 | Anthropogenic | Not Evaluated | N |
| *Melica nutans* agg. | 48 | 0.66 | <.001 | NA | NA | NA | 577 | -0.15 | <.001 | Dry to moderately moist forests | Not Threatened | I |
| *Elymus repens* s. str. | 727 | 0.67 | <.001 | 594 | -0.05 | NA | 3962 | 0.06 | NA | Scrubs, copses and field hedges | Not Threatened | I |
| *Matricaria chamomilla* | 21 | 0.69 | 0.001 | 16 | -0.49 | NA | 19 | -0.08 | NA | Mesic grasslands | Not Threatened | A |
| *Stellaria palustris* | 173 | 0.69 | <.001 | 74 | 0.01 | NA | 24 | 0.16 | NA | Linear and running surface waters | Endangered | I |
| *Plantago media* | 24 | 0.69 | 0.002 | 6 | -0.40 | NA | 1934 | -0.01 | NA | Heaths, inland dunes and semi-natural grasslands | Not Threatened | I |
| *Picea sitchensis* | 296 | 0.70 | <.001 | 10 | -0.27 | NA | 6 | 1.00 | NA | Dry to moderately moist forests | Not Evaluated | N |
| *Rubus* sect. *Caesii* | 724 | 0.71 | <.001 | 101 | 0.73 | <.001 | 9583 | 0.26 | <.001 | Scrubs, copses and field hedges | Not Threatened | I |
| *Stellaria alsine* | 364 | 0.71 | <.001 | 42 | 0.37 | NA | 942 | 0.13 | <.001 | Moist to wet grasslands | Not Threatened | I |
| *Erigeron annuus* | 11 | 0.74 | NA | 1 | -1.00 | NA | 430 | 0.34 | <.001 | None | Not Evaluated | N |
| *Brachypodium pinnatum* agg. | 9 | 0.78 | NA | 1 | 1.00 | NA | 10162 | 0.07 | <.001 | Scrubs, copses and field hedges | Not Threatened | I |
| *Atriplex prostrata* agg. | 314 | 0.79 | <.001 | 13 | -0.67 | NA | 6 | -0.38 | NA | Coastal and marine habitats | Not Threatened | I |
| *Galeopsis pubescens* | 8 | 0.80 | NA | 4 | 1.00 | NA | 18 | -0.72 | 0.001 | None | Not Threatened | I |
| *Fallopia dumetorum* | 64 | 0.83 | <.001 | 4 | 0.75 | NA | 48 | 0.24 | NA | None | Not Threatened | I |
| *Prunus serotina* | 1558 | 0.85 | <.001 | 255 | 0.49 | <.001 | 360 | 0.41 | <.001 | Dry to moderately moist forests | Not Evaluated | N |
| *Hypericum tetrapterum* | 182 | 0.86 | <.001 | 24 | 0.48 | NA | 586 | -0.25 | <.001 | Moist to wet grasslands | Not Threatened | I |
| *Larix kaempferi* | 166 | 0.87 | <.001 | 49 | 0.09 | NA | 26 | -0.07 | NA | Dry to moderately moist forests | Not Evaluated | N |
| *Epilobium ciliatum* | 131 | 0.87 | <.001 | 99 | 0.33 | <.001 | 90 | 0.12 | NA | Linear and running surface waters | Not Evaluated | N |
| *Luzula luzuloides* | 20 | 0.91 | <.001 | 4 | -0.75 | NA | 1884 | 0.05 | NA | Dry to moderately moist forests | Not Threatened | I |
| *Symphoricarpos albus* | 78 | 0.91 | <.001 | 122 | 0.15 | NA | 400 | 0.34 | <.001 | Anthropogenic | Not Evaluated | N |
| *Impatiens glandulifera* | 388 | 0.95 | <.001 | 99 | 0.65 | <.001 | 3699 | 0.53 | <.001 | Moist to wet forests | Not Evaluated | N |
| *Carex elytroides* | 118 | 0.96 | <.001 | 42 | 1.00 | <.001 | NA | NA | NA | Moist to wet grasslands | Not Evaluated | I |
| *Lilium martagon* | 1 | 1.00 | NA | 1 | -1.00 | NA | 880 | -0.14 | <.001 | Dry to moderately moist forests | Not Threatened | I |
| *Melampyrum sylvaticum* | 8 | 1.00 | 0.008 | 4 | -1.00 | NA | 224 | -0.14 | NA | None | Not Threatened | I |
| *Verbena officinalis* | 26 | 1.00 | <.001 | 1 | -1.00 | NA | 184 | 0.08 | NA | Heaths, inland dunes and semi-natural grasslands | Not Threatened | A |
| *Picea omorika* | 18 | 1.00 | <.001 | 2 | -0.83 | NA | 18 | -0.12 | NA | None | Not Evaluated | N |
| *Amaranthus retroflexus* | 1 | 1.00 | NA | 1 | -0.50 | NA | 13 | 0.86 | 0.003 | None | Not Evaluated | N |
| *Acer negundo* | 2 | 1.00 | NA | 4 | -0.36 | NA | 58 | 0.52 | <.001 | None | Not Evaluated | N |
| *Barbarea stricta* | 7 | 1.00 | 0.016 | 8 | -0.05 | NA | 1 | -1.00 | NA | Linear and running surface waters | Not Threatened | I |
| *Populus balsamifera* | 38 | 1.00 | <.001 | 16 | 0.06 | NA | 42 | 0.03 | NA | None | Not Evaluated | N |
| *Rumex pratensis* | 13 | 1.00 | <.001 | 17 | 0.12 | NA | NA | NA | NA | Mesic grasslands | Not Threatened | I |
| *Mahonia aquifolium* | 3 | 1.00 | NA | 20 | 0.17 | NA | 168 | 0.59 | <.001 | Anthropogenic | Not Evaluated | N |
| *Hordeum murinum* s. l. | 12 | 1.00 | <.001 | 5 | 0.20 | NA | 14 | 0.65 | NA | Mesic grasslands | Not Threatened | A |
| *Bidens frondosa* | 15 | 1.00 | <.001 | 55 | 0.20 | NA | 30 | 0.87 | <.001 | Standing waters | Not Evaluated | N |
| *Cornus mas* | 16 | 1.00 | <.001 | 8 | 0.26 | NA | 286 | -0.11 | NA | Scrubs, copses and field hedges | Not Threatened | I |
| *Prunus domestica* s. l. | 17 | 1.00 | <.001 | 84 | 0.29 | NA | 5762 | 0.04 | NA | Scrubs, copses and field hedges | Near Threatened | N |
| *Agrostis vinealis* | 24 | 1.00 | <.001 | 4 | 0.30 | NA | 4 | -0.27 | NA | Heaths, inland dunes and semi-natural grasslands | Near Threatened | I |
| *Ornithogalum umbellatum* agg. | 6 | 1.00 | 0.031 | 2 | 0.38 | NA | 35 | -0.52 | 0.005 | None | Not Threatened | A |
| *Hieracium aurantiacum* | 8 | 1.00 | 0.008 | 4 | 0.42 | NA | 22 | 0.27 | NA | Mesic grasslands | Not Threatened | I |
| *Chaerophyllum bulbosum* | 44 | 1.00 | <.001 | 16 | 0.47 | NA | 602 | 0.28 | <.001 | Scrubs, copses and field hedges | Not Threatened | I |
| *Populus trichocarpa* | 10 | 1.00 | 0.002 | 10 | 0.48 | NA | 2 | 1.00 | NA | None | Not Evaluated | N |
| *Ribes alpinum* | 1 | 1.00 | NA | 12 | 0.53 | NA | 709 | 0.12 | <.001 | Scrubs, copses and field hedges | Not Threatened | I |
| *Ceratocapnos claviculata* | 64 | 1.00 | <.001 | 4 | 0.58 | NA | NA | NA | NA | None | Not Threatened | I |
| *Prunus laurocerasus* | 5 | 1.00 | NA | 18 | 0.88 | <.001 | 86 | 1.00 | <.001 | Anthropogenic | Not Evaluated | N |
| *Matteuccia struthiopteris* | 24 | 1.00 | <.001 | 8 | 0.88 | 0.016 | 6 | 1.00 | NA | None | Near Threatened | I |
| *Prunus cerasifera* | 2 | 1.00 | NA | 9 | 0.90 | 0.008 | 736 | 0.56 | <.001 | Scrubs, copses and field hedges | Not Evaluated | N |
| *Agrimonia procera* | 29 | 1.00 | <.001 | 10 | 1.00 | 0.002 | 84 | -0.57 | <.001 | None | Not Threatened | I |
| *Narcissus pseudonarcissus* | 16 | 1.00 | <.001 | 2 | 1.00 | NA | 9 | -0.45 | NA | None | Endangered | I |
| *Sorbus torminalis* | 2 | 1.00 | NA | 2 | 1.00 | NA | 1587 | 0.10 | <.001 | Dry to moderately moist forests | Not Threatened | I |
| *Lathyrus latifolius* | 8 | 1.00 | 0.008 | 2 | 1.00 | NA | 89 | 0.15 | NA | None | Not Evaluated | N |
| *Parthenocissus quinquefolia* agg. | 6 | 1.00 | NA | 6 | 1.00 | NA | 175 | 0.51 | <.001 | None | Not Evaluated | N |
| *Fallopia baldschuanica* | 3 | 1.00 | NA | 5 | 1.00 | NA | 14 | 0.94 | <.001 | None | Not Evaluated | N |
| *Hyacinthoides non-scripta* | 16 | 1.00 | <.001 | 2 | 1.00 | NA | 2 | 1.00 | NA | None | Vulnerable | I |
| *Senecio inaequidens* | 40 | 1.00 | <.001 | 22 | 1.00 | <.001 | 16 | 1.00 | <.001 | Coastal and marine habitats | Not Evaluated | N |
| *Schoenus nigricans* | 1 | 1.00 | NA | NA | NA | NA | 44 | -0.43 | <.001 | None | Highly Endangered | I |
| *Euphorbia stricta* | 9 | 1.00 | 0.004 | NA | NA | NA | 52 | -0.34 | 0.015 | None | Not Threatened | I |
| *Euphorbia palustris* | 1 | 1.00 | NA | NA | NA | NA | 61 | -0.24 | 0.024 | Fresh water vegetation, springs and reeds | Endangered | I |
| *Phleum phleoides* | 1 | 1.00 | NA | NA | NA | NA | 274 | -0.19 | <.001 | Heaths, inland dunes and semi-natural grasslands | Near Threatened | I |
| *Asarum europaeum* | 1 | 1.00 | NA | NA | NA | NA | 3488 | -0.19 | <.001 | Scrubs, copses and field hedges | Not Threatened | I |
| *Helianthemum nummularium* s. l. | 1 | 1.00 | NA | NA | NA | NA | 2745 | -0.15 | <.001 | Heaths, inland dunes and semi-natural grasslands | Near Threatened | I |
| *Geranium sylvaticum* | 4 | 1.00 | NA | NA | NA | NA | 1142 | 0.20 | <.001 | Moist to wet grasslands | Not Threatened | I |
| *Poa chaixii* | 22 | 1.00 | <.001 | NA | NA | NA | 574 | 0.21 | <.001 | Heaths, inland dunes and semi-natural grasslands | Not Threatened | I |
| *Hieracium lactucella* | 1 | 1.00 | NA | NA | NA | NA | 684 | 0.24 | <.001 | Moist to wet grasslands | Endangered | I |
| *Phyteuma nigrum* | 1 | 1.00 | NA | NA | NA | NA | 195 | 0.30 | <.001 | Moist to wet grasslands | Near Threatened | I |
| *Carex pilosa* | 32 | 1.00 | <.001 | NA | NA | NA | 152 | 0.30 | <.001 | Dry to moderately moist forests | Not Threatened | I |
| *Potamogeton nodosus* | 2 | 1.00 | NA | NA | NA | NA | 30 | 0.82 | <.001 | None | Near Threatened | I |
| *Potentilla indica* | 1 | 1.00 | NA | NA | NA | NA | 31 | 0.94 | <.001 | None | Not Evaluated | N |
| *Phytolacca americana* | 1 | 1.00 | NA | NA | NA | NA | 74 | 0.99 | <.001 | None | Not Evaluated | N |
| *Conopodium majus* | 20 | 1.00 | <.001 | NA | NA | NA | NA | NA | NA | None | Not Threatened | N |
| *Rubus spectabilis* | 20 | 1.00 | <.001 | NA | NA | NA | NA | NA | NA | None | Not Evaluated | N |
| *Rosa gallica* | NA | NA | NA | 3 | -1.00 | NA | 86 | -0.35 | <.001 | None | Endangered | I |
| *Anthemis cotula* | NA | NA | NA | 7 | -1.00 | 0.016 | NA | NA | NA | None | Near Threatened | A |
| *Raphanus raphanistrum* agg. | NA | NA | NA | 6 | -0.41 | NA | 8 | -1.00 | 0.008 | None | Not Threatened | NA |
| *Salix sepulcralis* | NA | NA | NA | 78 | 0.15 | NA | 10 | 1.00 | 0.002 | Standing waters | Not Evaluated | N |
| *Philadelphus coronarius* | NA | NA | NA | 70 | 0.28 | 0.002 | 56 | 0.25 | NA | Anthropogenic | Not Evaluated | N |
| *Ailanthus altissima* | NA | NA | NA | 2 | 0.33 | NA | 47 | 0.78 | <.001 | None | Not Evaluated | N |
| *Cornus sericea_Cornus alba* | NA | NA | NA | 90 | 0.36 | <.001 | 238 | 0.33 | <.001 | Anthropogenic | Not Evaluated | N |
| *Pterocarya fraxinifolia* | NA | NA | NA | 10 | 0.71 | 0.008 | 1 | 1.00 | NA | Anthropogenic | Not Evaluated | N |
| *Lathyrus hirsutus* | NA | NA | NA | 1 | 1.00 | NA | 12 | -0.88 | 0.002 | None | Endangered | A |
| *Salix rubra* | NA | NA | NA | 2 | 1.00 | NA | 55 | -0.64 | <.001 | None | Not Threatened | I |
| *Rosa glauca* | NA | NA | NA | 1 | 1.00 | NA | 33 | -0.48 | 0.004 | None | Endangered | I |
| *Cerastium tomentosum* | NA | NA | NA | 1 | 1.00 | NA | 58 | -0.36 | 0.005 | None | Not Evaluated | N |
| *Muscari botryoides* | NA | NA | NA | 1 | 1.00 | NA | 244 | -0.24 | <.001 | Heaths, inland dunes and semi-natural grasslands | Endangered | I |
| *Senecio nemorensis* agg. | NA | NA | NA | 2 | 1.00 | NA | 2504 | -0.11 | <.001 | Dry to moderately moist forests | Not Threatened | I |
| *Asplenium scolopendrium* | NA | NA | NA | 1 | 1.00 | NA | 94 | 0.35 | <.001 | Dry to moderately moist forests | Not Threatened | I |
| *Cotoneaster horizontalis* | NA | NA | NA | 2 | 1.00 | NA | 52 | 0.56 | <.001 | None | Not Evaluated | N |
| *Buddleja davidii* | NA | NA | NA | 2 | 1.00 | NA | 36 | 0.85 | <.001 | None | Not Evaluated | N |
| *Pyracantha coccinea* | NA | NA | NA | 2 | 1.00 | NA | 9 | 0.95 | 0.008 | None | Not Evaluated | N |
| *Amelanchier lamarckii* | NA | NA | NA | 20 | 1.00 | <.001 | 8 | 1.00 | 0.008 | None | Not Evaluated | N |
| *Caucalis platycarpos* | NA | NA | NA | NA | NA | NA | 8 | -1.00 | 0.008 | None | Highly Endangered | A |
| *Erigeron strigosus* | NA | NA | NA | NA | NA | NA | 10 | -1.00 | 0.002 | None | Not Evaluated | N |
| *Poa bulbosa* | NA | NA | NA | NA | NA | NA | 8 | -1.00 | 0.008 | None | Not Threatened | I |
| *Ranunculus arvensis* | NA | NA | NA | NA | NA | NA | 13 | -0.97 | <.001 | None | Endangered | A |
| *Corallorhiza trifida* | NA | NA | NA | NA | NA | NA | 19 | -0.95 | <.001 | None | Endangered | I |
| *Carex sempervirens* | NA | NA | NA | NA | NA | NA | 13 | -0.88 | <.001 | None | Not Threatened | I |
| *Adonis aestivalis* | NA | NA | NA | NA | NA | NA | 17 | -0.82 | 0.002 | None | Highly Endangered | A |
| *Thymelaea passerina* | NA | NA | NA | NA | NA | NA | 11 | -0.76 | 0.008 | None | Highly Endangered | I |
| *Ranunculus montanus* agg. | NA | NA | NA | NA | NA | NA | 52 | -0.70 | <.001 | None | Not Threatened | I |
| *Dianthus sylvaticus* | NA | NA | NA | NA | NA | NA | 86 | -0.69 | <.001 | None | Endangered | I |
| *Lathyrus aphaca* | NA | NA | NA | NA | NA | NA | 38 | -0.67 | <.001 | None | Endangered | A |
| *Festuca amethystina* | NA | NA | NA | NA | NA | NA | 17 | -0.65 | 0.003 | None | Not Threatened | I |
| *Aster bellidiastrum* | NA | NA | NA | NA | NA | NA | 68 | -0.56 | <.001 | None | Not Threatened | I |
| *Allium sativum* | NA | NA | NA | NA | NA | NA | 98 | -0.52 | <.001 | None | Not Evaluated | N |
| *Orobanche teucrii* | NA | NA | NA | NA | NA | NA | 70 | -0.50 | <.001 | Heaths, inland dunes and semi-natural grasslands | Endangered | I |
| *Astragalus cicer* | NA | NA | NA | NA | NA | NA | 62 | -0.50 | <.001 | None | Near Threatened | I |
| *Gentiana asclepiadea* | NA | NA | NA | NA | NA | NA | 56 | -0.47 | <.001 | None | Not Threatened | I |
| *Orchis pallens* | NA | NA | NA | NA | NA | NA | 56 | -0.44 | 0.003 | None | Endangered | I |
| *Ophrys insectifera* | NA | NA | NA | NA | NA | NA | 250 | -0.43 | <.001 | Heaths, inland dunes and semi-natural grasslands | Endangered | I |
| *Gentianopsis ciliata* | NA | NA | NA | NA | NA | NA | 776 | -0.42 | <.001 | Heaths, inland dunes and semi-natural grasslands | Near Threatened | I |
| *Gentianella germanica* agg. | NA | NA | NA | NA | NA | NA | 408 | -0.40 | <.001 | Heaths, inland dunes and semi-natural grasslands | Near Threatened | I |
| *Lonicera caprifolium* | NA | NA | NA | NA | NA | NA | 44 | -0.40 | 0.011 | None | Not Threatened | I |
| *Forsythia suspensa* | NA | NA | NA | NA | NA | NA | 36 | -0.37 | 0.009 | None | Not Evaluated | N |
| *Centaurea montana* | NA | NA | NA | NA | NA | NA | 109 | -0.36 | <.001 | None | Not Threatened | I |
| *Noccaea montana* | NA | NA | NA | NA | NA | NA | 112 | -0.33 | <.001 | Dry to moderately moist forests | Endangered | I |
| *Orobanche caryophyllacea* | NA | NA | NA | NA | NA | NA | 276 | -0.33 | <.001 | Heaths, inland dunes and semi-natural grasslands | Endangered | I |
| *Galeopsis ladanum* agg. | NA | NA | NA | NA | NA | NA | 146 | -0.31 | <.001 | None | Not Threatened | I |
| *Trifolium rubens* | NA | NA | NA | NA | NA | NA | 179 | -0.31 | <.001 | Heaths, inland dunes and semi-natural grasslands | Endangered | I |
| *Epipactis atrorubens* | NA | NA | NA | NA | NA | NA | 214 | -0.30 | <.001 | Dry to moderately moist forests | Near Threatened | I |
| *Ophrys apifera* | NA | NA | NA | NA | NA | NA | 198 | -0.29 | <.001 | Heaths, inland dunes and semi-natural grasslands | Not Threatened | I |
| *Orchis purpurea* | NA | NA | NA | NA | NA | NA | 166 | -0.29 | <.001 | Heaths, inland dunes and semi-natural grasslands | Near Threatened | I |
| *Gentiana verna* | NA | NA | NA | NA | NA | NA | 254 | -0.28 | <.001 | Heaths, inland dunes and semi-natural grasslands | Endangered | I |
| *Scilla bifolia* | NA | NA | NA | NA | NA | NA | 70 | -0.28 | 0.01 | None | Not Threatened | I |
| *Salvia verticillata* | NA | NA | NA | NA | NA | NA | 258 | -0.25 | <.001 | Heaths, inland dunes and semi-natural grasslands | Not Threatened | N |
| *Crepis alpestris* | NA | NA | NA | NA | NA | NA | 92 | -0.25 | 0.003 | Heaths, inland dunes and semi-natural grasslands | Endangered | I |
| *Cytisus nigricans* | NA | NA | NA | NA | NA | NA | 147 | -0.23 | <.001 | Heaths, inland dunes and semi-natural grasslands | Endangered | I |
| *Gymnadenia conopsea* s. l. | NA | NA | NA | NA | NA | NA | 1036 | -0.21 | <.001 | Heaths, inland dunes and semi-natural grasslands | Near Threatened | I |
| *Cirsium rivulare* | NA | NA | NA | NA | NA | NA | 704 | -0.20 | <.001 | Moist to wet grasslands | Endangered | I |
| *Trollius europaeus* | NA | NA | NA | NA | NA | NA | 695 | -0.20 | <.001 | Moist to wet grasslands | Endangered | I |
| *Teucrium botrys* | NA | NA | NA | NA | NA | NA | 206 | -0.20 | <.001 | Heaths, inland dunes and semi-natural grasslands | Near Threatened | A |
| *Orchis militaris* | NA | NA | NA | NA | NA | NA | 678 | -0.19 | <.001 | Heaths, inland dunes and semi-natural grasslands | Endangered | I |
| *Aster amellus* | NA | NA | NA | NA | NA | NA | 706 | -0.17 | <.001 | Heaths, inland dunes and semi-natural grasslands | Endangered | I |
| *Thesium bavarum* | NA | NA | NA | NA | NA | NA | 448 | -0.16 | <.001 | Heaths, inland dunes and semi-natural grasslands | Endangered | I |
| *Koeleria pyramidata* agg. | NA | NA | NA | NA | NA | NA | 2228 | -0.16 | <.001 | Heaths, inland dunes and semi-natural grasslands | Near Threatened | I |
| *Polygala amara* agg. | NA | NA | NA | NA | NA | NA | 424 | -0.15 | <.001 | Heaths, inland dunes and semi-natural grasslands | Endangered | I |
| *Carlina acaulis* | NA | NA | NA | NA | NA | NA | 2255 | -0.15 | <.001 | Heaths, inland dunes and semi-natural grasslands | Near Threatened | I |
| *Asperula cynanchica* | NA | NA | NA | NA | NA | NA | 1772 | -0.13 | <.001 | Heaths, inland dunes and semi-natural grasslands | Near Threatened | I |
| *Stachys recta* | NA | NA | NA | NA | NA | NA | 2856 | -0.09 | <.001 | Heaths, inland dunes and semi-natural grasslands | Near Threatened | I |
| *Hippocrepis comosa* | NA | NA | NA | NA | NA | NA | 2273 | -0.08 | <.001 | Heaths, inland dunes and semi-natural grasslands | Near Threatened | I |
| *Chaerophyllum aureum* | NA | NA | NA | NA | NA | NA | 3942 | 0.07 | <.001 | Scrubs, copses and field hedges | Not Threatened | I |
| *Euphorbia verrucosa* | NA | NA | NA | NA | NA | NA | 1164 | 0.13 | <.001 | Heaths, inland dunes and semi-natural grasslands | Near Threatened | I |
| *Inula conyzae* | NA | NA | NA | NA | NA | NA | 1338 | 0.14 | <.001 | Heaths, inland dunes and semi-natural grasslands | Not Threatened | I |
| *Meum athamanticum* | NA | NA | NA | NA | NA | NA | 966 | 0.16 | <.001 | Heaths, inland dunes and semi-natural grasslands | Near Threatened | I |
| *Chaerophyllum hirsutum* agg. | NA | NA | NA | NA | NA | NA | 2356 | 0.16 | <.001 | Moist to wet grasslands | Not Threatened | I |
| *Polystichum aculeatum* agg. | NA | NA | NA | NA | NA | NA | 430 | 0.19 | <.001 | Dry to moderately moist forests | Not Threatened | I |
| *Rosa arvensis* | NA | NA | NA | NA | NA | NA | 2290 | 0.22 | <.001 | Scrubs, copses and field hedges | Near Threatened | I |
| *Carex alba* | NA | NA | NA | NA | NA | NA | 129 | 0.24 | 0.001 | Dry to moderately moist forests | Not Threatened | I |
| *Viola mirabilis* | NA | NA | NA | NA | NA | NA | 228 | 0.26 | <.001 | Dry to moderately moist forests | Near Threatened | I |
| *Isatis tinctoria* | NA | NA | NA | NA | NA | NA | 141 | 0.27 | <.001 | Heaths, inland dunes and semi-natural grasslands | Not Threatened | A |
| *Rhinanthus aristatus* agg. | NA | NA | NA | NA | NA | NA | 420 | 0.28 | <.001 | Heaths, inland dunes and semi-natural grasslands | Near Threatened | I |
| *Dictamnus albus* | NA | NA | NA | NA | NA | NA | 37 | 0.31 | 0.027 | None | Endangered | I |
| *Centaurea phrygia* agg. | NA | NA | NA | NA | NA | NA | 154 | 0.32 | <.001 | Moist to wet grasslands | Endangered | I |
| *Allium sphaerocephalon* | NA | NA | NA | NA | NA | NA | 58 | 0.32 | 0.003 | Heaths, inland dunes and semi-natural grasslands | Endangered | I |
| *Orchis pyramidalis* | NA | NA | NA | NA | NA | NA | 205 | 0.47 | <.001 | Heaths, inland dunes and semi-natural grasslands | Endangered | I |
| *Athyrium distentifolium* | NA | NA | NA | NA | NA | NA | 30 | 0.48 | 0.009 | None | Not Threatened | I |
| *Lonicera nigra* | NA | NA | NA | NA | NA | NA | 146 | 0.48 | <.001 | None | Not Threatened | I |
| *Juglans nigra* | NA | NA | NA | NA | NA | NA | 28 | 0.50 | 0.019 | None | Not Evaluated | N |
| *Rumex arifolius* | NA | NA | NA | NA | NA | NA | 35 | 0.50 | 0.008 | None | Not Threatened | I |
| *Torilis arvensis* | NA | NA | NA | NA | NA | NA | 120 | 0.57 | <.001 | None | Not Threatened | A |
| *Himantoglossum hircinum* | NA | NA | NA | NA | NA | NA | 244 | 0.59 | <.001 | Heaths, inland dunes and semi-natural grasslands | Not Threatened | I |
| *Viscum album* agg. | NA | NA | NA | NA | NA | NA | 278 | 0.62 | <.001 | None | Not Threatened | I |
| *Crepis mollis* | NA | NA | NA | NA | NA | NA | 160 | 0.63 | <.001 | Moist to wet grasslands | Endangered | I |
| *Portulaca oleracea* | NA | NA | NA | NA | NA | NA | 21 | 0.64 | 0.004 | None | Not Evaluated | A |
| *Crepis pulchra* | NA | NA | NA | NA | NA | NA | 46 | 0.82 | <.001 | None | Not Threatened | I |
| *Aubrieta deltoidea* | NA | NA | NA | NA | NA | NA | 7 | 1.00 | 0.016 | None | Not Evaluated | N |

.
 Notes: n-, and p- values according to two-tailed binomial tests. In addition, each species preferred habitat type and its Red list (RL) and non-native (NN; I = Native (indigenous), N =
 Neophytes, A = Archaeophytes, NA = no status assigned) status in Germany are given. Habitat type preferences are based on the fidelity (Φ) of species to habitat types in all states
 taken together.
